# Dynamic decomposition of transcriptome responses during plant effector-triggered immunity revealed conserved responses in two distinct cell populations

**DOI:** 10.1101/2022.12.30.522333

**Authors:** Xiaotong Liu, Daisuke Igarashi, Rachel A. Hillmer, Thomas Stoddard, You Lu, Kenichi Tsuda, Chad L. Myers, Fumiaki Katagiri

## Abstract

- Rapid plant immune responses in the appropriate cells are needed for effective defense against pathogens. Although transcriptome analysis is often used to describe overall immune responses, collecting transcriptome data with sufficient resolution in both space and time is challenging.
- We reanalyzed public Arabidopsis time-course transcriptome data obtained after a low-dose inoculation of a *Pseudomonas syringae* strain expressing the effector AvrRpt2, which induces Effector-Triggered Immunity (ETI) in Arabidopsis. Double-peak time-course patterns were prevalent among thousands of upregulated genes. We implemented a multi-compartment modeling approach to decompose the double-peak pattern into two single-peak patterns for each gene.
- The decomposed peaks revealed an “echoing” pattern: the peak times of the first and second peaks correlated well across most upregulated genes. We demonstrated that two peaks likely represent responses of two distinct cell populations, which respond either cell-autonomously or indirectly to AvrRpt2. Thus, the peak decomposition extracted spatial information from the time-course data.
- The echoing pattern also indicated a conserved transcriptome response between two cell populations despite different elicitor types. WRKY transcription factors appeared to underlie the conserved transcriptome response. Activation of a WRKY network via different entry-point WRKYs could explain the conserved transcriptome response elicited by different elicitor types.

## Introduction

Two modes of inducible immunity have been well characterized in plants: Pattern-Triggered Immunity (PTI) and Effector-Triggered Immunity (ETI) (Jones & Dangl, 2006; Dodds & Rathjen, 2010). PTI signaling is initiated when molecular patterns in the apoplastic space are recognized by Pattern-Recognition Receptors (PRRs) on the cell surface (Albert *et al*., 2020). Such molecular patterns include Microbe-Associated Molecular Patterns (MAMPs) and Damage-Associated Molecular Patterns (DAMPs). For example, a 22-amino acid peptide of the bacterial flagellin, flg22, is a MAMP recognized by an Arabidopsis PRR, FLS2 (Gómez-Gómez & Boller, 2000). Similarly, Arabidopsis DAMPs, Plant Elicitor Peptides (PEPs), are recognized by functionally redundant Arabidopsis PRRs, PEPR1 and PEPR2 (Krol *et al*., 2010; Yamaguchi *et al*., 2010). Pathogens well-adapted to plant hosts deliver effectors into the plant cell that interfere with PTI signaling (Khan *et al*., 2018). ETI signaling is initiated when some of the pathogen effectors or their impacts are recognized by cognate intracellular resistance (R) proteins (van der Hoorn & Kamoun, 2008). For example, impacts of effectors from the bacterial pathogen *Pseudomonas syringae*, AvrRpt2 and AvrRpm1, on a plant protein, RIN4, are recognized by Arabidopsis R proteins, RPS2 and RPM1, respectively (Mackey *et al*., 2002; Axtell & Staskawicz, 2003; Mackey *et al*., 2003).

Plant immunity is expensive as it requires large amounts of energy and resources (Bolton, 2009). Furthermore, plant immunity includes responses harmful to host cells as well as pathogens. For example, during ETI, plant cells autonomously recognizing a pathogen effector undergo programmed cell death (PCD), called a hypersensitive response (HR) (Coll *et al*., 2011). The cost of immunity is likely the reason plants evolved to have inducible, rather than constitutive immunity (Heidel *et al*., 2004). It is important for plants to have immunity induced rapidly at the right place when it is needed. Thus, tracking when and where particular immune responses are induced with respect to pathogen infection is crucial in understanding plant immunity.

For measurements of many different responses, transcriptome analysis is a standard, relatively economical approach. For example, Mine et al. collected time-course transcriptome data during ETI in six Arabidopsis genotypes inoculated with *P*. *syringae* pv *tomato* DC3000 (*Pto*) strains carrying an empty vector or expressing AvrRpt2 or AvrRpm1 effectors (Mine *et al*., 2018). The study revealed the importance of immune signaling sectors (Tsuda *et al*., 2009) in rapid transcriptome responses during ETI. There are other studies in which transcriptome analysis was performed at the cellular level of spatial resolution during plant immunity, e.g., (Chandran *et al*., 2010; Zhu *et al*., 2022). However, transcriptome analysis with sufficiently high resolution in both time and space is technically and economically challenging, particularly with plant tissues.

Time-course transcriptomes are typically analyzed assuming independence across time points using profile-based approaches such as clustering or statistical models such as ANOVA (Eisen *et al*., 1998; Ben-Dor *et al*., 1999; Cui & Churchill, 2003). While these approaches are useful in classification of genes into different expression pattern groups and detection of differentially expressed genes, respectively, the temporal continuity of the data is not fully appreciated. Mathematical models that can capture temporal continuity information have been used, such as regression and spline models (Ramsay & Silverman, 2005). However, these models are not built upon specific mechanistic hypotheses and generally fail to provide mechanistic interpretations.

A Multi-Compartment Model (MCM) is a type of mathematical model used to describe some quantity (signals, in our study) transmitted among compartments of a system (Godfrey, 1983). It has been applied to many fields of biology, including pharmacokinetics and epidemiology (Kermack & McKendrick, 1927; Rescigno, 1960; Prabakaran *et al*., 2021). Each compartment has an input and an output and a rule defining the output. A simple implementation of a single compartment has a first-order self-decay as the rule (Fig. 1A). The output of a compartment (compartment Y in Fig. 1A) is specified by the input (output of compartment X), the input amplification ratio (𝑎_𝑌_), and the self-decay rate (𝑘_𝑌_). An MCM formed by a network of such simple compartments is a tractable linear ordinary differential equation (ODE) system, and the mechanistic parameters of each compartment, the input amplification ratio and self-decay rate, are readily interpretable.

**Fig. 1.**
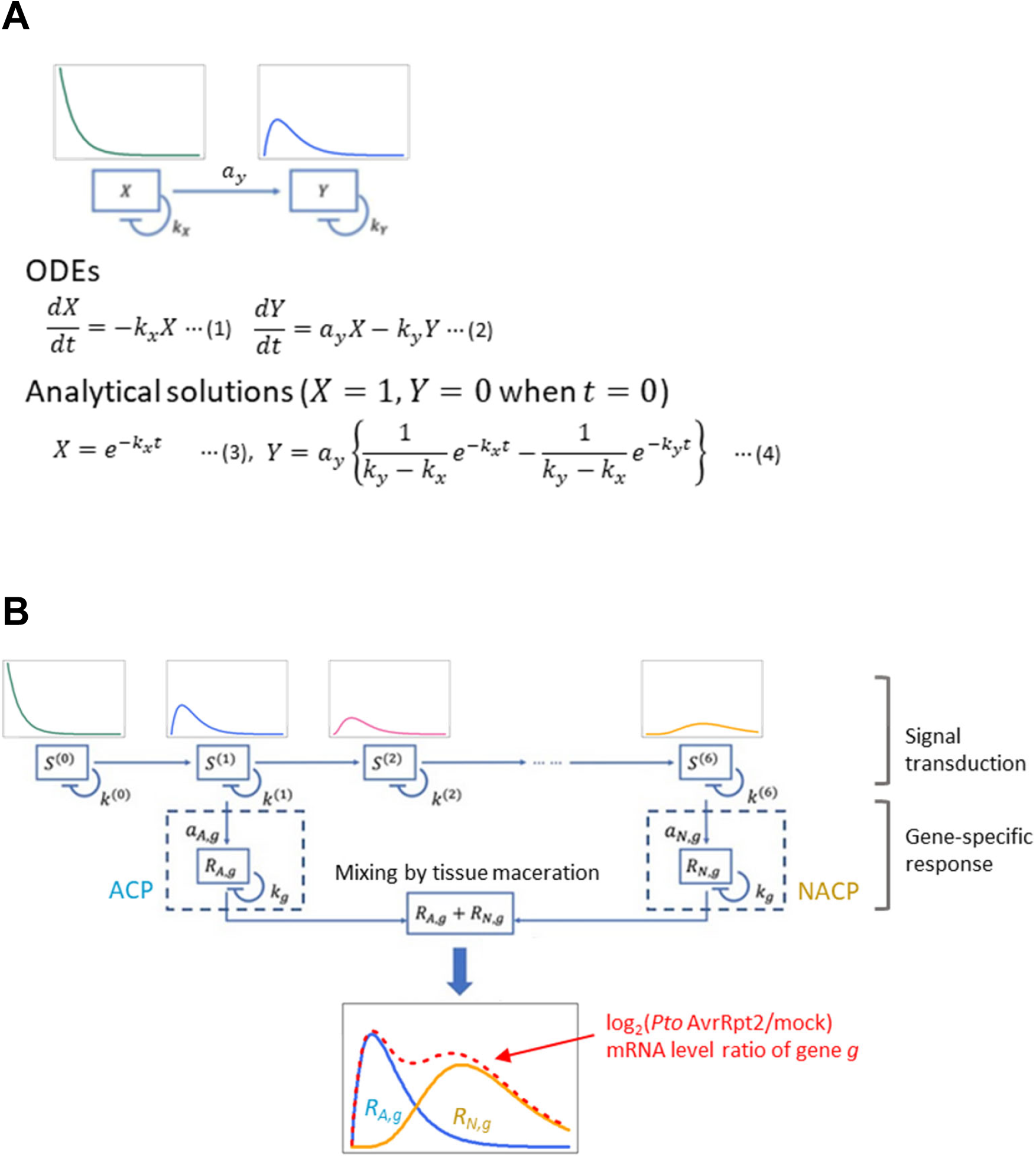
The design of the MCM. **A**. A two-compartment model. Each of the compartments, *X* and *Y*, has a parameter that determines the first-order self-decay rate (𝑘_𝑥_ and 𝑘_𝑦_). The input level to compartment *Y* is proportional to the input amplification rate parameter 𝑎_𝑦_. The output of each compartment, *X* and *Y*, has a first-order decay. The output of X has a first order decay, while the output of Y has a single-peak time course (see 𝑆^(0)^ and 𝑆^(1)^ in **B**). The model is defined by the ordinary differential equations (ODEs) (1) and (2) for each compartment. The analytical solutions for the output of the compartments are generally in the form of linear combinations of first order decays in MCM. Solutions for the two-compartment model under the initial conditions of 𝑋 = 1 and 𝑌 = 0 at 𝑡 = 0 are shown. **B**. The MCM structure used in the study. A series of signaling compartments (*S*^(1)^ to *S*^(6)^, shown as an example) generate single-peak time-course patterns with delayed peak times (shown above each compartment) when 𝑆^(𝑛)^(0) = 0, 𝑛 = 1,2, …, 6. The signaling compartment series is common among all the genes we modeled, and the parameter values for the signaling compartments were predetermined. For each gene, the output from a signaling compartment (*S*^(1)^ as an example in the figure) was used as the input to the first-peak (ACP) response compartment (*R_A,g_*), and the output from another signaling compartment (*S*^(6)^ as an example in the figure) was used as the input to the second-peak (NACP) response compartment (*R_N,g_*). A common gene-specific decay rate, *k_g_*, was assumed for the ACP and NACP response compartments of a gene *g*. The gene-specific input amplification ratio parameters of the response compartments are *a_A,g_*. and *a_N,g_*. Thus, for each gene, the selection of the two signaling compartments and the parameters *k_g_*, *a_A,g_*. and *a_N,g_*, were fit to the mRNA ratio time-course data corresponding to log_2_(*Pto* AvrRpt2/mock).

In this study, we reanalyzed the transcriptome data set generated by Mine et al. (Mine *et al*., 2018). We noticed that double-peak time-course patterns were prevalent among the genes upregulated in Arabidopsis leaves upon inoculation with a relatively low dose of *Pto* AvrRpt2. We implemented an MCM to decompose the double-peak time-course into two single-peak time-courses for each gene. This double-peak decomposition revealed an “echoing” pattern between two single peaks in approximately 1400 upregulated genes: when the first peak for a gene is early, its second peak is also early; when the first peak for a gene is late, its second peak is also late. We demonstrated that the first and second peak responses likely represent responses of two distinct plant cell populations: the cells responding cell-autonomously to AvrRpt2 (autonomous cell population, ACP) and those surrounding the ACP and responding indirectly (non-autonomous cell population, NACP). The echoing pattern of the transcriptome responses also showed that there is a well-conserved transcriptome response between two cell populations whereas their eliciting signal molecule types are different. The echoing mRNA response pattern between two cell populations appeared to be mainly regulated by WRKY transcription factors. We propose WRKY network activation through different entry-point WRKYs as a mechanism for the conserved transcriptome response.

## Materials and Methods

Detailed methods are provided in Text S1.

### RNA-seq data analysis, general

The data analysis was performed in R (R_Core_Team, 2021) using custom codes. All R scripts are available from https://github.com/fumikatagiri/MCM_At_AvrRpt2_time_course.

### RNA-seq data set, mRNA level value estimates, and gene selection

We used a subset of the RNA-seq read count data set from Mine et al. (Mine *et al*., 2018) (GEO accession GSE88798). The subset included the data collected from the leaf tissues of Arabidopsis Col-0 at 1, 2, 3, 4, 6, 9, 12, 16, 20, 24 hours post inoculation (hpi) of mock (water) or *Pto* AvrRpt2 with three biological replicates. We used the entire data set, which contains 366 libraries, for the pre-selection of 18442 genes which had at least 39 read counts for the 15th highest among 366 read count values for each gene.

We fit a negative binomial generalized linear model (GLM-NB) to the read count data, while normalizing the 90^th^ percentile read count of each library (Hillmer *et al*., 2017). All values for the mRNA level and level ratio in this study are expressed as log_2_-transformed values.

We selected 3039 significantly upregulated genes by comparing the mean values with *Pto* AvrRpt2 to those in mock after 3 hpi (Storey FDR = 0.05, log_2_FC > 1; Fig. S1A). We defined the signal-to-noise ratio as (maximum of log_2_FC across the time points) / (mean of the SEs for the log_2_FC across the time points) and selected 2435 high-precision upregulated genes with signal-to-noise ratio > 6.5 (Figs. S1B and S1C). A spline model was fit to the log_2_FC data of each of 2435 genes (Fig. S2).

### An MCM for a double-peak time-course

The output change rate of a single compartment is defined by ODE (2) in Fig. 1A. Our MCM was composed of a chain of 12 signaling compartments, 𝑆^(𝑛)^(𝑡), 𝑛 = 0,1,2, …, 11, which generated single-peak time-courses with delaying peak times. Then two gene-specific response compartments 𝑅_𝐴,𝑔_ and 𝑅_𝑁,𝑔_ linearly combined the outputs of two single-peak signaling compartments with different peak times to generate a double-peak time-course pattern for gene 𝑔 (Fig. 1B). Two sets, Sets 1 and 2, of the predetermined decay rates for the signaling compartments were used to generate the single-peak patterns with the predetermined peak times at signaling compartments 𝑆^(𝑛)^, 𝑛 = 1,3,5,7,9,11: Set 1, ([4, 6, 9, 12, 16, 20] × 0.95) hpi and Set 2, ([3.5, 5, 7.5, 10.5, 14, 18] × 0.95) hpi, which were relatively orthogonal.

For each gene, the outputs of two signaling compartments were used as the inputs to the response compartments, and the best of 10 combinations (indicated by the 𝑛 value of 𝑆^(𝑛)^: (1, 5), (1, 7), (1, 9), (1, 11), (3, 7), (3, 9), (3, 11), (5, 9), (5, 11), and (7, 11)) was selected through MCM fitting by considering the combination as a discrete variable of the model. We assigned a common decay rate parameter, 𝑘_𝑔_, to both response compartments, 𝑅_𝐴,𝑔_ and 𝑅_𝑁,𝑔_, for each gene (Fig. 1B) to avoid overfitting. Thus, for gene 𝑔, four parameters, 𝑘_𝑔_, two input amplification ratios of the response compartments, 𝑎_𝐴,𝑔_ and 𝑎_𝑁,𝑔_ (Fig. 1B), and the signaling compartment combination were fit in the MCM, given one of Sets 1 or 2 for the signaling compartment parameter values. The MCM was fit to the log-transformed mRNA ratio (log_2_(*Pto* AvrRpt2/mock), which is equal to log_2_(*Pto* AvrRpt2) – log_2_(mock)). This was because mRNA species in Arabidopsis commonly follow a double-exponential decay time course pattern (Sorenson *et al*., 2018). Consequently, the mRNA level log-ratio of a gene should follow a first-order decay, which is assumed at each compartment.

### Altered MCMs for the genes that had time-course patterns of single peak with a shoulder

The modeled values were clearly higher than data values at 6 hpi in 191 genes (the gene selection criteria are in Text S1), suggesting that they have early second peak responses and their time-courses are like a single peak with a shoulder. To impose early double peaks in model fitting, only the read count data up to 16 hpi were used in the “altered MCM” for 191 genes (Fig. S3).

### TF-binding sites enrichment analysis

We used the Arabidopsis cistrome data set for 349 TFs across > 30000 genes in the analysis (O’Malley *et al*., 2016). For each gene, the DNA sequence 1000 bp upstream to 200 bp downstream of the transcription start site was defined as the promoter region and searched for TF binding sites. Fisher’s exact test (1-sided) was used to calculate the *p*-value for enrichment of genes with each TF-binding site among ACP-specific, Echoing, or NACP-specific genes compared to the proportions in the genome. This yielded 100 TFs with Benjamini-Hochberg FDR-corrected *p* < 0.01.

We examined whether the proportion of genes with a binding site for each of 100 significant TFs temporally changes. Subsets of (ACP-specific + Echoing) or (Echoing + NACP-specific) gene sets were made using a sliding window of 150 genes with a sliding step of 15 genes along the peak time of the first-peak or the second-peak response, respectively. For each gene subset, the *p*-value was calculated using Fisher’s exact test (2-sided) compared to the proportion in the gene sets. We selected 38 TFs with *p* < 0.001 in at least one window subset for further study. Similar analysis was performed for consolidated TF families and Echoing genes only.

## Results

### Double-peak time-course patterns are prevalent in the mRNA levels of *Pto* AvrRpt2-upregulated genes

From the dataset (Mine *et al*., 2018), we selected 3039 genes significantly upregulated at one or more time points after 3 hours post inoculation (hpi) with *Pto* AvrRpt2 compared to mock (water). We focused on the time range after 3 hpi because major responses of most genes were confined to this time range (Fig. S1A). We further selected 2435 high-precision upregulated genes (Figs. S1B and S1C). A preliminary visual inspection using a spline model of the mRNA level log-ratio of *Pto* AvrRpt2 over mock for each gene suggested widespread double-peak time-course patterns (Fig. S2A).

We hypothesized that widespread double-peak responses are superimposed responses of two distinct cell populations (Lu & Tsuda, 2021), each of which has single-peak responses (Fig. 2). In fact, the experimental system with a low inoculation dose of *Pto* AvrRpt2 (OD_600_ = 0.001, (Mine *et al*., 2018)) contains at least two plant cell populations with distinct responses, e.g., (Hatsugai *et al*., 2016). One population of plant cells, ACP, receives and cell-autonomously responds to the ETI-eliciting effector AvrRpt2. Most ACP cells undergo HR cell death by 9 hpi (Hatsugai *et al*., 2017). Another population of plant cells, NACP, surround the ACP cells and do not undergo HR cell death. Based on the cell death timing, we reasoned that any responses with peak times after 9 hpi must be attributed to NACP cells. Thus, a better-defined form of the two-cell population hypothesis is that responses with peak times before 9 hpi are largely attributable to ACP cells.

**Fig. 2.**
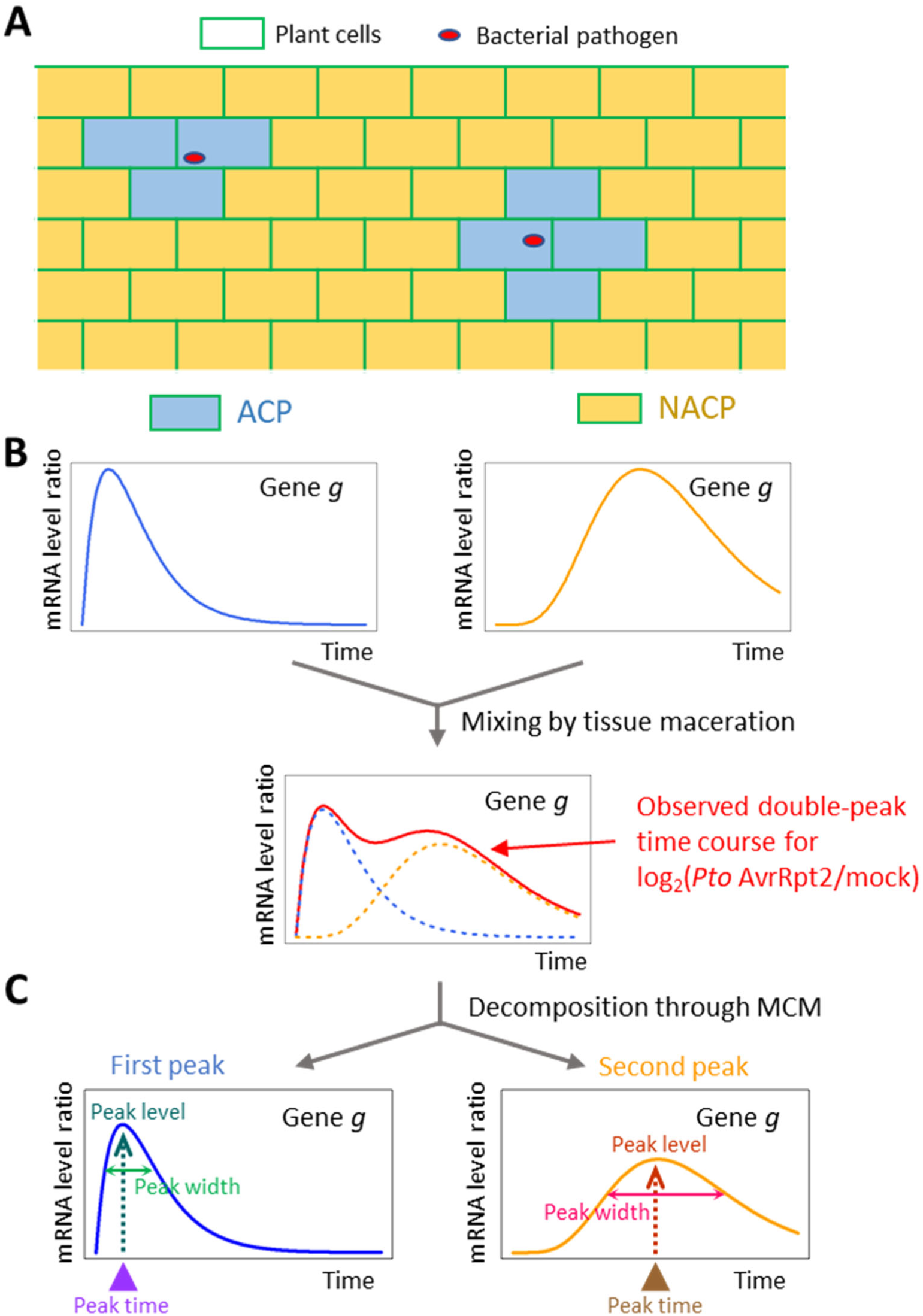
Two plant cell populations spatially defined relative to the locations of bacterial cells can explain the double-peak time-courses of the mRNA levels. **A**. A schematic diagram of two plant cell populations. The plant cells in the close vicinity of bacterial cells receive the effector AvrRpt2 from the bacterial cells and the ETI response is induced cell-autonomously (Autonomous Cell Population, ACP; blue cells). Most cells in ACP undergo the HR cell death by 9 hpi. The plant cells surrounding ACP respond to AvrRpt2 in an indirect manner (Non-Autonomous Cell Population, NACP; orange cells). NACP cells do not die. **B**. A schematic diagram explaining how a single-peak response in each of ACP and NACP (blue and orange traces, respectively) can be observed as a double-peak response (red trace) in the log(mRNA level ratio) time course for gene 𝑔. The mRNA molecules from two cell populations are mixed when the tissue is macerated in the RNA preparation procedure. **C**. We decomposed the double-peak time-course pattern into two single-peak patterns (first and second peaks) using MCM. Each peak can be characterized by the peak level, peak width, and peak time.

### The MCM used in the study

To test the hypothesis of two cell populations, we designed an MCM to decompose the double-peak time-course mRNA level log-ratio pattern of each gene into two single-peak patterns. A chain of compartments generates a single-peak time-course pattern at each compartment with a delaying peak time along the chain (signaling compartments labeled as 𝑆^(𝑖)^, 𝑖 = 1,2, …, 6 in Fig. 1B). This signaling compartment chain may be considered as molecular components in a conceptualized signal transduction pathway. For practicality of model fitting, the parameters of the signaling compartments, including their numbers, were predetermined.

For regulation of a particular gene 𝑔, we modeled as if the outputs from some upstream and downstream signaling compartments (𝑆^(1)^ and 𝑆^(6)^in Fig. 1B as an example) were used in ACP or NACP, respectively. The mRNA level log-ratios of gene 𝑔 in ACP and NACP were modeled as the outputs of response compartments 𝑅_𝐴,𝑔_ and 𝑅_𝑁,𝑔_, respectively. When the leaf tissue was macerated for RNA preparation, mRNA molecules from ACP and NACP were mixed, and the mixed mRNA level log-ratio was modeled as the double-peak pattern, 𝑅_𝐴,𝑔_ + 𝑅_𝑁,𝑔_. The input amplification parameters 𝑎_𝐴,𝑔_ and 𝑎_𝑁,𝑔_ of the response compartments depend on regulation of gene 𝑔 in ACP and NACP and the ratio of ACP and NACP in the tissue. 𝑅_𝐴,𝑔_ and 𝑅_𝑁,𝑔_ represent two decomposed single-peak time-courses for the gene (Figs 1B and 2C).

### The MCM explains the double-peak transcriptome response well

To avoid bias potentially introduced by using the predetermined signal compartment parameters, consistency in the model fit between two sets of the parameter values, Sets 1 and 2, (Text S1; the fitted parameter values are in Table S1) was used as the criterion for a good model. The fitted values for the two peaks with Sets 1 and 2 across 2435 selected upregulated genes were generally well-correlated (Fig. S4). We selected 1889 genes with Pearson correlation ≥ 0.9 as the genes stably modeled by the MCM, and the MCMs with Set 1 for these genes were used subsequently.

The MCM compared to the data over time for each gene is given in Fig. S5. The median across the genes of the Pearson correlations between the mean estimates and the MCM-modeled values across the time points was 0.93 (Fig. S6). The heatmaps in Fig. 3A show the discrepancies across the genes between the mean estimates of the data and the MCM-modeled values at each time point. The discrepancies were small and had no obvious pattern, except for the 191 genes shown at the bottom of the figure. These 191 genes had very early first and second peaks, and their third peaks were not captured by the model. We concluded that our MCM explains the double-peak transcriptome response well.

**Fig. 3.**
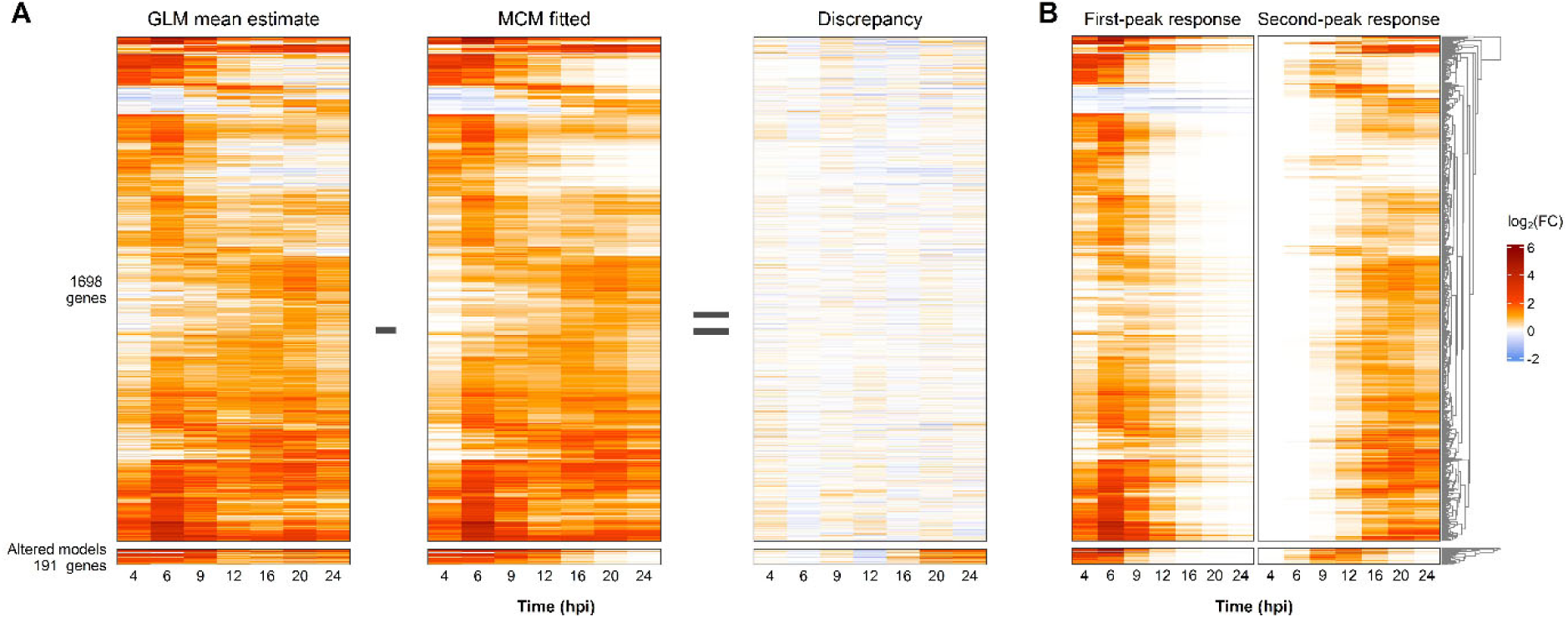
MCM decomposed the double-peak time-course pattern. **A**. The fit of the MCM. Heatmaps of the GLM mean estimates, MCM fitted values, and the discrepancies between them (left to right panels) are shown for 1889 (=1698 + 191) upregulated genes that were consistently modeled between Set 1 and Set 2 of the signaling compartment decay rate parameter values across 7 time points (hpi). The bottom 191 genes had single-peak-with-a-shoulder time-course patterns, so MCM was fit to the data at 4-16 hpi time points (“Altered models”). The genes were clustered according to the MCM fitted values using the Euclidean distance and the average linkage for 1698 and 191 genes separately. **B**. Decomposition into two single-peak time course patterns. Heatmaps of the first-peak and second-peak responses modeled in MCM are shown for 1889 genes across 7 time points (hpi). The gene order is the same as in **A**. The color scale of the heatmap value is common between **A** and **B** and shown at the right.

### The second-peak response echoes the first-peak response

Using the outputs of the response compartments, 𝑅_𝐴,𝑔_ and 𝑅_𝑁,𝑔_, the double-peak pattern was decomposed into two single-peak patterns for each gene (Figs. 3B and S5). We characterized the single-peak patterns by their peak times and peak levels (Fig. 2C). We classified genes into three groups, 214 “First peak-specific”, 1366 “Echoing”, and 227 “Second peak-specific” genes, according to the ratio of the first peak level over the second peak level (1807 modeled-and-classified upregulated genes; Fig. S7 and Table S2), sorted the genes in each group according to their peak times, and visualized them using heatmaps (Fig. 4A). Among the 1366 Echoing genes, it was strikingly evident that the first peak time-course patterns were well replicated in the second peak time-course patterns: when the first peak of a gene was relatively early, its second peak tended to be early; when the first peak was late, its second peak tended to be late. We call this replicated pattern an echoing of transcriptome responses (hence the gene group name). The second-peak responses were time-stretched as well as delayed compared to the first-peak responses in the Echoing genes (Note the time interval size differences in Fig. 4A). The mean ratio of the second peak time over the first peak time, which indicates the mean time-delaying rate, was 2.8. The mean ratio of the second peak width over the first peak width (width at 75% peak level, Fig. 2C), which indicates the time-stretching ratio, was 2.2 (Fig. S8). Since these values are similar, the second-peak response is approximately time-scaled from the first-peak response with a scaling factor of about 2.5.

**Fig. 4.**
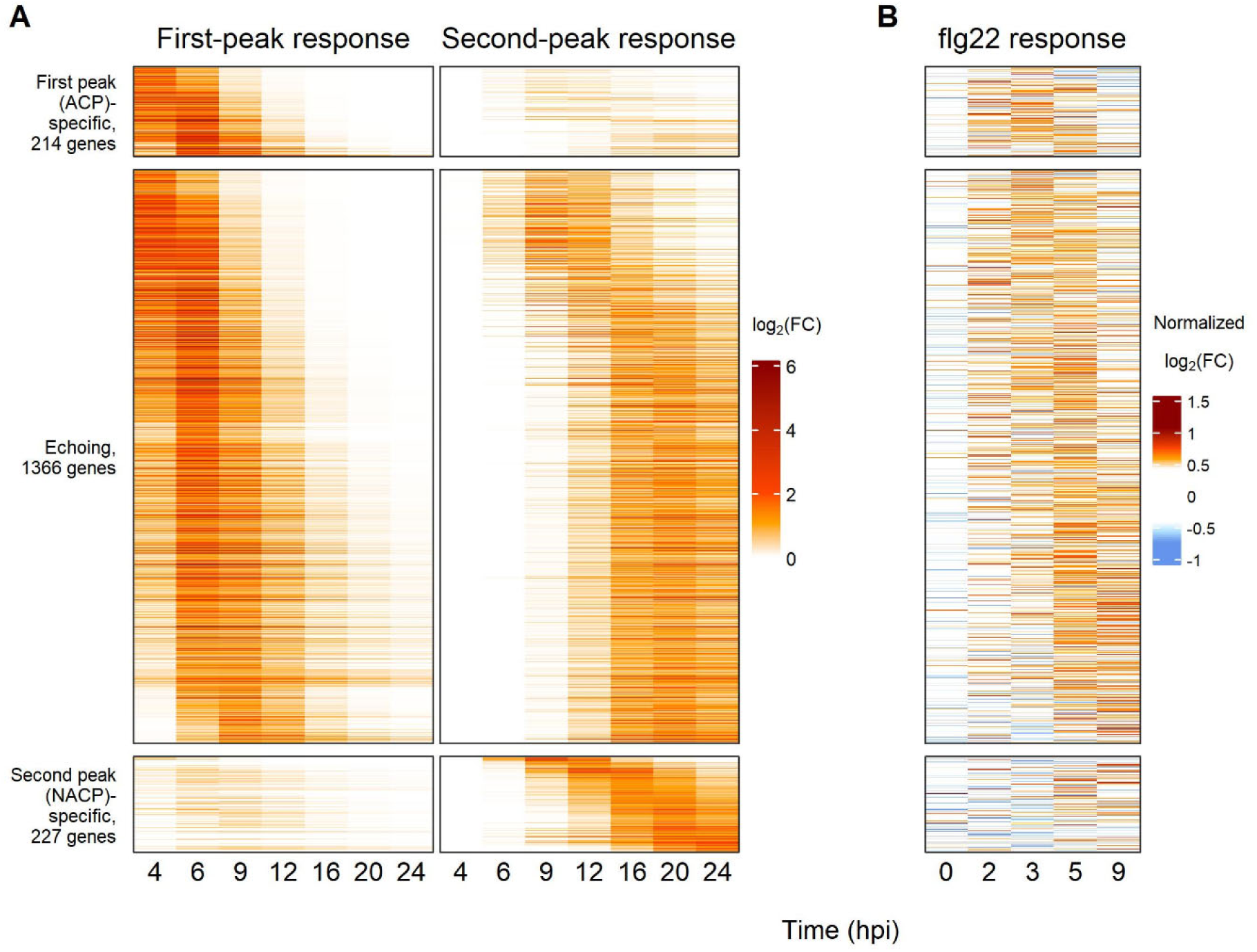
The second-peak response “echoes” the first-peak response across about 1400 genes. **A**. Heatmaps of the first-peak and second-peak responses modeled by MCM (left and right panels, respectively) for 1807 modeled-and-classified upregulated genes. The genes were classified to three groups, “First peak (ACP)-specific”, “Echoing”, and “Second peak (NACP)-specific” genes, according to the peak level ratio value between the first and second peaks of the gene. The number of genes in each group is also shown. Within each group, the genes were ordered according to the modeled peak time values of the first peaks of the genes for First peak-specific and Echoing genes and those of the second peaks for Second peak-specific genes. The values used in the heatmaps are the log_2_ mRNA level ratio of *Pto* AvrRpt2-innoculated over mock at the indicated hpi. Among the 1889 modeled upregulated genes, 82 genes were unclassified and not included in this figure. **B**. A heatmap of the flg22 response mean estimates (Hillmer et al., 2017). The gene groups and the gene orders are the same as in **A**. The log_2_ mRNA level ratio of flg22- infiltrated Arabidopsis Col-0 over flg22-infiltrated Arabidopsis *fls2* mutant was normalized to the vector size of 1 for each gene across the time points. These normalized log_2_-fold change values were used in the heatmaps. Although the original data included a 1-hpi time point, those data were removed from this figure because such early responses largely represent non-immune-specific, general stress responses (Bjornson *et al*., 2021).

We interrogated our hypothesis that the first-peak responses, whose peak times were mostly earlier than 9 hpi, are largely attributable to ACP. First, we compared the transcriptome response after *Pto* AvrRpt2 inoculation to that after *Pto* inoculation in the same dataset (Mine *et al*., 2018). With *Pto* AvrRpt2 inoculation, ACP cells received the ETI-eliciting effector AvrRpt2, in addition to immune-suppressing effectors, and underwent the autonomous ETI response, including the HR. With *Pto* inoculation, the cells spatially equivalent to ACP cells only receive immune-suppressing effectors, and their immune response is thus predicted to be strongly suppressed during the comparable time range. Consistent with this prediction, in the time range for the first-peak response with the *Pto* AvrRpt2-inoculated samples (4 to 9 hpi), very little response was observed in the *Pto* inoculated samples (Figure 1D in Mine et al., 2018). Thus, this observation suggests that the transcriptome responses in the early time range are mainly responses of cells spatially equivalent to ACP.

Second, we studied RNA-seq data from Arabidopsis Col-0 plants in which AvrRpt2 protein was transgenically and conditionally expressed in plant cells (“*in planta*-expressed AvrRpt2” data) (Hillmer *et al*., 2023). Since AvrRpt2 protein was expressed upon induction in virtually all cells, resulting in macroscopic HR cell death (Tsuda *et al*., 2012; Hatsugai *et al*., 2017), the transcriptome response before cell death in the *in planta*-expressed AvrRpt2 data should mostly represent responses in ACP. When the peak times and the peak levels of the first and second peak responses to *Pto* AvrRpt2 and *in planta*-expressed AvrRpt2 were compared, the first-peak response was more similar than the second-peak response to the response to *in planta*-expressed AvrRpt2 (Figs. 5A-5D). Next, we sorted 1807 modeled-and-classified upregulated genes according to the ratio of the first peak level over the second peak level, and subsets of the sorted genes were made using a sliding window of 150 genes. When the number of overlapping genes between each of the gene subsets and the 1972 genes for *in planta*-expressed AvrRpt2 upregulated genes were visualized, the overall trend was clear. The higher the ratio of the first peak level over the second peak level, the larger the overlapping gene number was (Fig. 5E). Since the genes with higher peak ratios are more first peak-specific/preferential, this trend also indicates that the first-peak response is more similar to the response to the *in planta*-expressed AvrRpt2. In summary, two lines of evidence support the hypothesis that the first peak response is largely from ACP.

**Fig. 5.**
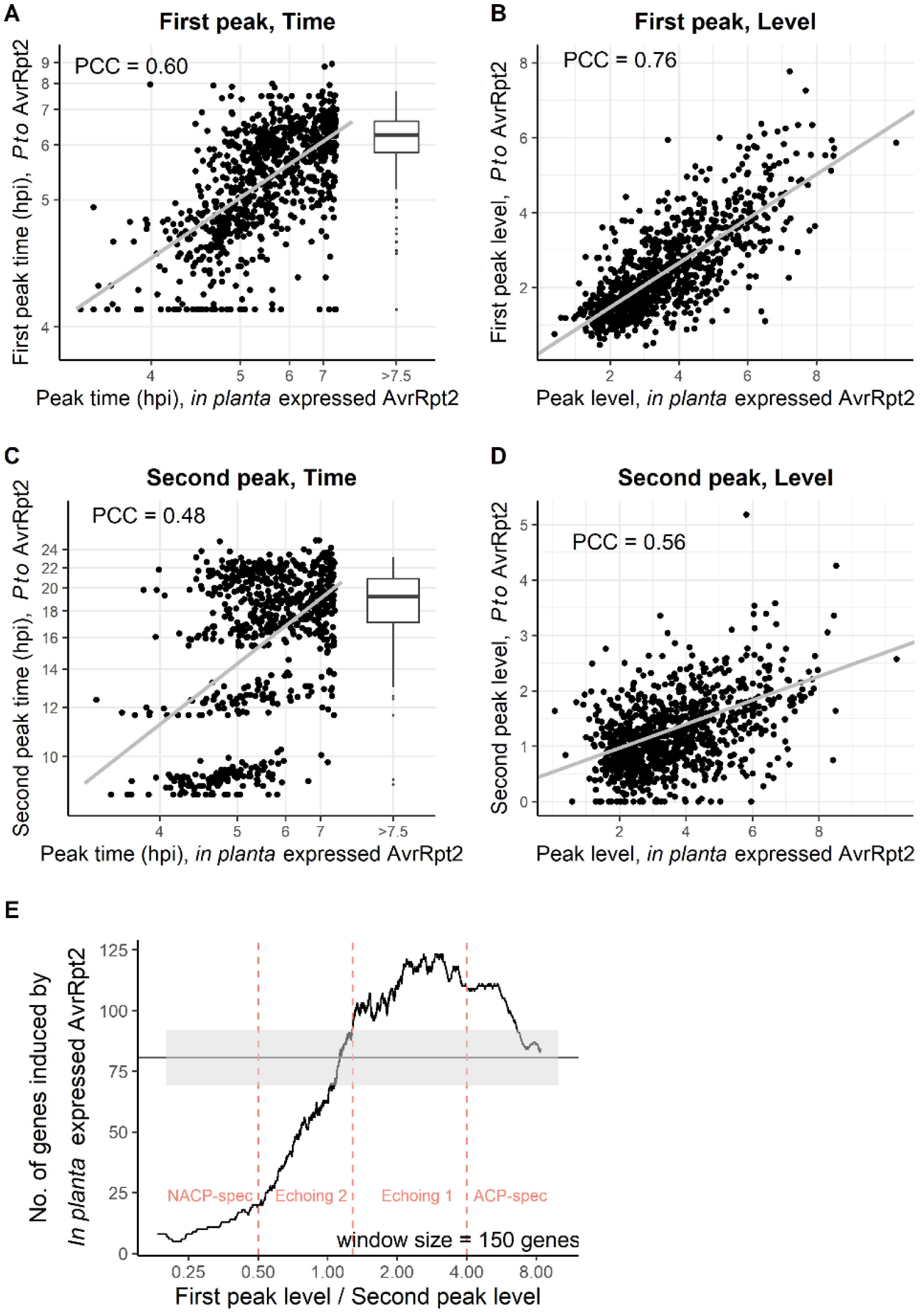
The first-peak response is more similar than the second-peak response to the ACP response induced by *in planta*-expressed AvrRpt2. **A-D**. The peak times (**A**, **C**) and the peak levels (**B**, **D**) of the first-peak (**A**, **B**) and the second-peak (**C**, **D**) responses were compared to the peak times (**A**, **C**) and the peak levels (**B**, **D**) of the response to *in planta*-expressed AvrRpt2 across the 1092 (**A**, **B**) and 967 (**C**, **D**) overlapping genes. Since late peak time estimates for the *in planta*-expressed AvrRpt2 response are not precise, the genes with the *in planta*-expressed AvrRpt2 peak times later than 7.5 hpi are aggregated into a group and a box plot of the *Pto* AvrRpt2-inoculated peak times for the group is shown (“> 7.5” hpi; **A**, **C**). A linear regression line (gray line) and Pearson correlation coefficient (PCC) are shown in each panel. In (**A**, **C**), the linear regression and the correlation coefficients were calculated for the genes excluding the “> 7.5” hpi group after the peak time values were offset-subtracted and log-transformed, as seen in the scaling of the axes. **E**. The level of the overlap in the modeled upregulated genes between the *Pto* AvrRpt2-innoculated and the *in planta*-expressed AvrRpt2 along the ratio of the peak levels between the first and second peaks in the *Pto* AvrRpt2-innoculated is shown using a sliding window method (window size, 150 genes). The dashed horizontal line and the gray shade indicate the mean and its 95% confidence interval (Fisher’s exact test, 2-sided), respectively, for the entire 1807 modeled and classified upregulated genes. Pink vertical dashed lines show the boundary first-peak level/second-peak level ratio values of four gene groups, ACP-specific, Echoing 1, Echoing 2, and NACP-specific genes. Echoing genes shown in Fig. 4 were divided into Echoing 1 and Echoing 2 genes at the first-peak level/second-peak level ratio for the upper boundary of the 95% confidence interval.

### Two-peak decomposition sorted genes into functionally distinct groups

If our two-peak decomposition captures characteristics of actual biological mechanisms, gene groups defined by their two-peak decomposition characteristics should contain genes with different functions. We tested this idea using GO term enrichment analysis. As classifiers, we used the first- and second-peak level ratio and the peak times: four peak-level-ratio categories, ACP-specific, Echoing 1 (ACP-preferential), Echoing 2 (NACP-preferential), and NACP- specific (Fig. 5E), and early and late peak time categories; 4 x 2 = 8 groups. Note that the early and late peak times are determined relatively within each peak-level-ratio category and are not directly comparable across the peak-level-ratio categories.

The results of biological process GO term enrichment analysis are summarized in Table 1 (details in Table S2). While general defense-related terms (“defense response to …”) were enriched in ACP-preferential and early groups, this trend was weaker in the Echoing 2 late group, and no defense-related term was significantly enriched in NACP-specific groups (Table S2). This observation may suggest that the ETI transcriptome responses specific to NACP are quite different from what we think of as immune-related transcriptome responses.

**Table 1.** Enrichment of biological process GO terms specific to gene groups defined by double peak characteristics. The values in the gene group columns shows fold enrichment compared to the genome mean proportions. The higher the fold enrichment values are, the darker the orange color shades. The sizes of the eight gene groups are shown at the second row. The “whole.genome” column shows the number of genes in the genome for each GO terms.

|  | whole.genome | ACP.spec.early | Echo1.early | Echo2.early | NACP.spec.early | ACP.spec.late | Echo1.late | Echo2.late | NACP.spec.late |
| --- | --- | --- | --- | --- | --- | --- | --- | --- | --- |
| Number of genes | 27430 | 107 | 428 | 264 | 104 | 107 | 428 | 265 | 104 |
| response to oxidative stress (GO:0006979) | 743 | 5 | 3.7 | 3.1 | - | 4.5 | 3.9 | - | - |
| defense response to fungus (GO:0050832) | 930 | 6.6 | 7.4 | 4.2 | - | - | 4.7 | - | - |
| defense response to bacterium (GO:0042742) | 1105 | 6.8 | 7 | 3.9 | - | - | - | 3 | - |
| cellular response to hypoxia (GO:0071456) | 239 | 14.5 | 8.6 | - | - | 9.7 | 4.3 | - | - |
| response to inorganic substance (GO:0010035) | 2250 | 3 | - | 2.8 | - | 3.3 | - | - | - |
| response to water deprivation (GO:0009414) | 1250 | - | 2.3 | - | - | - | 3.3 | - | 3.8 |
| response to abscisic acid (GO:0009737) | 1181 | - | 3 | - | - | - | 3.1 | 2.5 | - |
| response to wounding (GO:0009611) | 943 | - | 4.5 | 2.7 | - | - | 4.1 | - | - |
| response to molecule of bacterial origin (GO:0002237) | 160 | - | 13.7 | 7.3 | - | - | 6.5 | - | - |
| response to salicylic acid (GO:0009751) | 418 | - | - | 6.3 | - | - | 4.5 | 4 | - |
| regulation of defense response (GO:0031347) | 885 | 4.2 | - | - | - | - | 5.5 | - | - |
| protein phosphorylation (GO:0006468) | 1070 | - | 3.2 | 3.1 | - | - | - | - | - |
| cellular response to lipid (GO:0071396) | 837 | - | 3.4 | - | - | - | 3.9 | - | - |
| defense response to other organism (GO:0098542) | 2299 | - | 4.9 | - | - | 3.1 | - | - | - |
| organonitrogen compound catabolic process (GO:1901565) | 1200 | - | - | 2.6 | - | - | - | 3.6 | - |
| cellular response to heat (GO:0034605) | 77 | 24.2 | - | - | - | - | - | - | - |
| heat acclimation (GO:0010286) | 71 | 30 | - | - | - | - | - | - | - |
| regulation of signal transduction (GO:0009966) | 453 | - | 4.8 | - | - | - | - | - | - |
| salicylic acid mediated signaling pathway (GO:0009863) | 83 | - | 11.2 | - | - | - | - | - | - |
| plant-type hypersensitive response (GO:0009626) | 58 | - | 11.4 | - | - | - | - | - | - |
| regulation of plant-type hypersensitive response (GO:0010363) | 28 | - | 14.2 | - | - | - | - | - | - |
| chlorophyll catabolic process (GO:0015996) | 38 | - | - | - | 42.5 | - | - | - | - |
| cellular response to oxygen-containing compound (GO:1901701) | 1258 | - | - | - | - | - | 3.9 | - | - |
| fatty acid catabolic process (GO:0009062) | 43 | - | - | - | - | - | - | 14.7 | - |
| fatty acid oxidation (GO:0019395) | 41 | - | - | - | - | - | - | 15.4 | - |
| sulfate assimilation (GO:0000103) | 22 | - | - | - | - | - | - | 23.9 | - |

HR-related terms were specifically enriched in the Echoing 1 early group, which is consistent with the fact that ACP undergoes the HR. These terms were not significantly enriched in the ACP-specific early group. This trend that HR-related genes tend to be also expressed, but relatively weakly, in NACP suggests that the HR program could be activated in NACP as well as ACP by default but that an HR-suppressing program could also be activated in NACP to limit spread of the HR (Jabs *et al*., 1996). HR-associated reactive oxygen species (ROS) accumulation (Su *et al*., 2018) appears to stress both ACP and NACP as the “response to oxidative stress” term is enriched in gene groups for both cell populations. Hypoxic response terms were strongly enriched in the ACP-specific early group. These observations suggest that O_2_ is converted to ROS mainly in the HR-undergoing ACP, and consequently O_2_ is limiting in ACP, inducing the hypoxic response. ACP-generated ROS probably diffuses and stresses both ACP and NACP. The sulfate assimilation term is enriched in the Echoing 2 late group, suggesting that reducing compounds, such as glutathione, accumulate in NACP, which might be part of the HR- suppressing program (Király *et al*., 2012).

Terms related to water-deprivation response, including response to abscisic acid, were enriched in the Echoing 1 early and late groups and the Echoing 2 and NACP-specific late groups. Limiting water accessibility in the apoplastic space is an effective ETI response against apoplastic pathogens, such as *Pto* (Xin *et al*., 2016). Plants may activate a water-deprivation response to draw water from the apoplast into the cytosol, thereby limiting apoplastic water availability, rapidly in ACP and later in NACP.

It is common for the mRNA levels of photosynthesis-related genes to be downregulated during immune responses (Bilgin *et al*., 2010) (as in this data set; Table S3), which would decrease synthesis of new photosynthetic apparatus. The enrichment of chlorophyll-catabolic genes in the NACP-specific early group suggests active downregulation of photosynthetic apparatus specifically in NACP. Fatty acid catabolism-related terms were enriched in the Echoing 2 late group, which suggests mobilization of an alternative energy resource in NACP while photosynthesis is rapidly downregulated. In summary, gene grouping based on two-peak decomposition characteristics revealed group-specific/preferential GO term enrichment, demonstrating the biological relevance of our two-peak decomposition approach.

### Echoing genes with the binding sites of WRKYs, NACs, HSFs, and CAMTA1 are associated with certain peak times

To gain insight into the regulatory mechanisms underlying the echoing transcriptome response, we performed TF-binding site enrichment analysis among 1807 modeled-and-classified upregulated genes. For 38 highly significantly enriched TF-binding sites, the distributions of the over- or under-representation of genes with each TF-binding site along the peak times for first- and second-peak responses were visualized (Fig. 6A). Most of the TFs for the binding sites in the figure belong to three large TF families, the NAC (Mathew & Agarwal, 2018), WRKY (Birkenbihl *et al*., 2018; Chen *et al*., 2019), and HSF (Guo *et al*., 2016) families. The time-dependent representation patterns within each of the WRKY and HSF families were similar, while there appear to be two different patterns within the NAC family (the heatmaps and the dendrogram on the left in Fig. 6A). While members of both NAC groups (NAC1s and NAC2s, blue and cyan for the TF names) had overrepresentation around 8 hpi, only NAC2s had overrepresentation around 6.5 hpi and 14 hpi and underrepresentation around 20 hpi. WRKYs (orange TF names) had a common pattern in the first- and second-peak responses: strong overrepresentation around 4.5 and 10 hpi, followed by overrepresentation around 6.3 and 18.5 hpi, and further followed by strong underrepresentation around 8 and 20 hpi. Thus, the representation patterns of the genes with WRKY-binding sites show an echoing pattern. HSFs (red TF names) had overrepresentation around 5 and 16.5 hpi, but the number of sliding windows with overrepresentation in the first-peak response was higher, suggesting higher overrepresentation in ACP-specific genes. Indeed, the proportions of the genes with binding sites for any of the HSF members are 26%, 16%, and 11% for ACP-specific, Echoing, and NACP-specific genes, respectively. This higher representation of genes with HSF-binding sites among ACP-specific genes is consistent with the heat response GO term enrichment in the ACP-specific early gene group (Table 1).

**Fig. 6.**
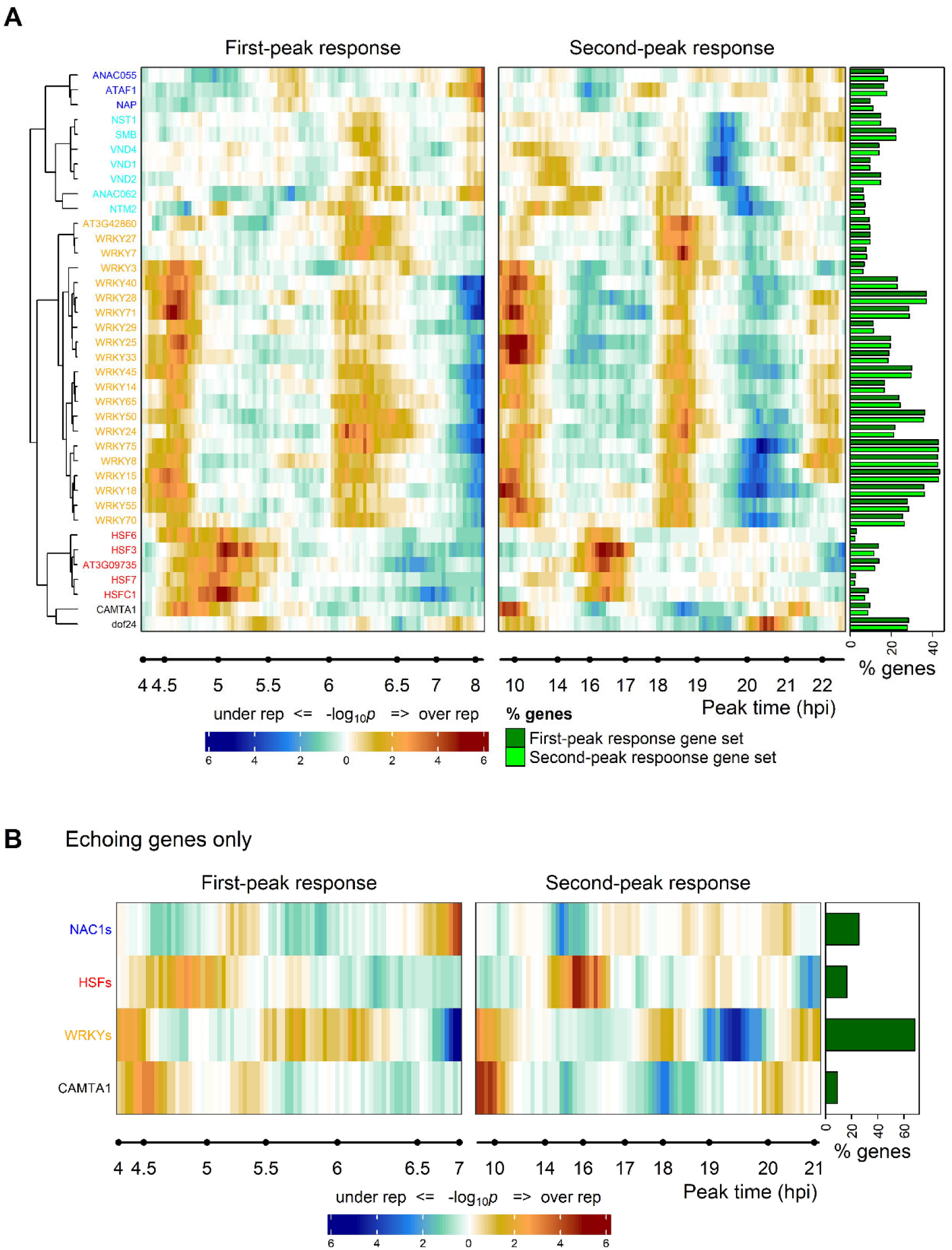
Genes with WRKY-binding sites are abundant among 1366 Echoing genes and show an echoing distribution pattern across the first- and second-peak responses. **A**. Enrichment of genes with the indicated TF-binding sites along the peak time of the first- and second-peak responses. The significance in a -log_10_ scale and whether the genes with the TF-binding sites are over-represented or under-represented are shown in the heatmaps. The color scale bar is shown at the bottom. The members of the NAC1, NAC2, WRKY, and HSF families are indicated by blue, cyan, orange, and red font colors. CAMTA1 and dof24 are singletons and shown in a black font color. Structurally, AT3G42860 and AT3G09735 do not belong to these TF families, but they were included in the WRKY and HSF families, respectively, due to their similar distribution patterns along the peak times. The horizontal bar plot on the right shows the proportion of the genes with the indicated TF-binding sites in each of the first- and second-peak response gene sets (214 ACP-specific plus 1366 Echoing genes and 1366 Echoing genes plus 227 NACP-specific genes, respectively). **B**. Heatmaps similar to **A** with consolidated NAC1s, HSFs, and WRKYs and CAMTA1 for 1366 Echoing genes only. The horizontal bar plot on the right shows the proportion of Echoing genes with the TF-binding sites.

We consolidated each of the TF families into one for better visualization of singletons, CAMTA1 and dof24: for one consolidated TF family, genes with binding sites for any of the family members were counted. The consolidated TF-binding sites heatmap generally conserved the features of the patterns for the TF families described above (Fig. S9).

To further explore the echoing transcriptome response, we selected NAC1s, HSFs, WRKYs, and CAMTA1, which showed significant temporal variation in representation (Fig. 6B). Genes with the CAMTA1-binding site have an early echoing overrepresentation around 4.5 hpi and 10 hpi. CAMTAs are transcription activators for general stress responses, likely responding to increased intracellular Ca^2+^ concentration (Doherty *et al*., 2009; Benn *et al*., 2014; Bjornson *et al*., 2021). Intracellular Ca^2+^ concentration increase also occurs in ACP early during ETI (Grant *et al*., 2000), which explains overrepresentation of genes with the CAMTA1-binding site around 4.5 hpi. Their overrepresentation around 10 hpi strongly suggests that elicitation of responses in NACP is also associated with increased intracellular Ca^2+^ concentration. Since the proportion of Echoing genes with CAMTA1-binding sites was only 9% and since its echoing pattern was limited to early times, CAMTA1 cannot explain the response behaviors of most Echoing genes. In this light, WRKYs were remarkable: 68% of Echoing genes had WRKY-binding sites, and the echoing representation patterns of the genes cover most of the time range of interest (Fig. 6B). We conclude that it is highly likely that WRKYs are central to the echoing transcriptome response during ETI.

### Diverse WRKY genes are transcriptionally upregulated in the time ranges of interest in both cell populations

We examined transcriptional regulation of 35 *WRKY* genes among 3039 genes upregulated after *Pto* AvrRpt2 inoculation. Their temporal mRNA levels are shown in a heatmap (Fig. 7A), and the profiles of binding sites for the WRKY proteins in the data set (columns) (O’Malley *et al*., 2016) in the 35 *WRKY* genes (rows) are also shown (Fig. 7C). Different subgroups of WRKYs (Eulgem *et al*., 2000) are indicated by different colors of the WRKY gene or protein names. We speculate that the binding specificities of the WRKY proteins are more similar within a subgroup than across the subgroups. First, many *WRKY* genes were transcriptionally upregulated from 4 to 24 hpi, strongly suggesting that many WRKY TFs were activated in the time ranges of interest in both cell populations (Fig. 7A). Second, there is no obvious correlation between the mRNA level time-course pattern and certain WRKY subgroups (i.e., the colors of the WRKY gene names are well mixed in Fig. 7C), suggesting that some members of most WRKY subgroups were upregulated in the time ranges for both cell populations. This means that any genes that have at least one WRKY-binding site could be transcriptionally regulated by WRKYs at any time in the time range in either cell population. Third, 28 out of the 35 upregulated *WRKY* genes have at least one WRKY-binding site (Fig. 7C), suggesting that the majority of the 35 *WRKY* genes are regulated by WRKY TFs. While WRKY TFs can be transcription activators or repressors (Birkenbihl *et al*., 2018), these observations are consistent with the notion that WKRY TFs likely play a major role in mediating the echoing transcriptome responses.

**Fig. 7.**
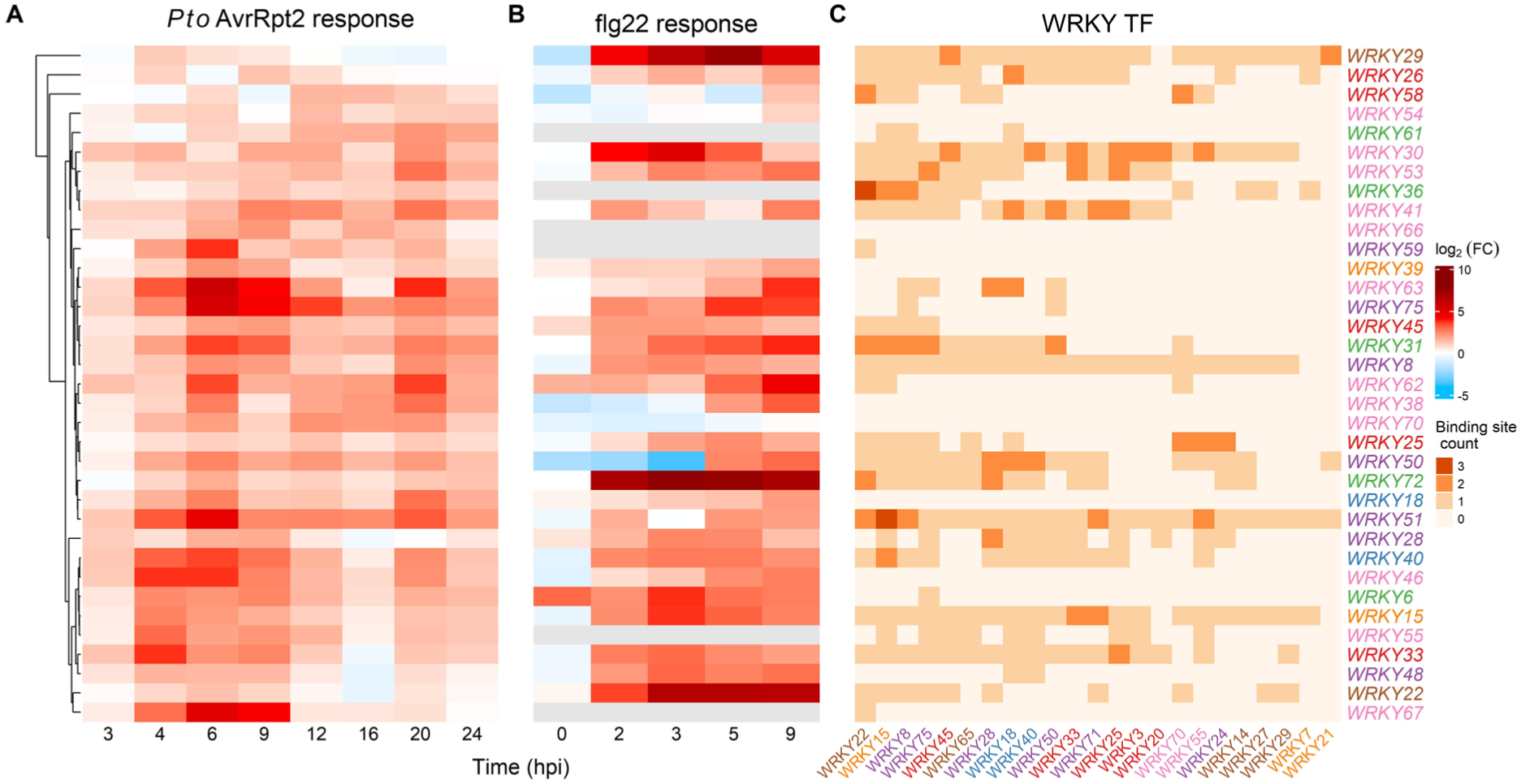
Diverse WRKY genes, most of which have WRKY-binding sites, are transcriptionally activated during the times of interest. **A**. A heatmap of the log mRNA level ratio (*Pto* AvrRpt2 over mock) time courses for 35 WRKY genes after inoculation of *Pto* AvrRpt2. **B**. A heatmap of the log mRNA level ratio (wild type over *fls2*) time courses for 35 WRKY genes after infiltration of flg22. **C**. The number of binding sites for diverse WRKY TFs (labeled at the bottom) in the 35 WRKY genes (labeled at the right). Different font colors for the names of WRKY genes (right) or TFs (bottom) represent different subgroups defined by (Eulgem *et al*., 2000). The WRKY gene orders in the row are the same across **A**-**C**. The color scale bars for the log_2_-fold change for **A** and **B** and for the number of binding sites per gene for **C** are shown at the right of C.

## Discussion

### MCM is a powerful modeling framework for temporal responses

MCM is a general mathematical framework that is highly applicable to a wide range of systems. The structure of MCM is a network of compartments. Thus, MCM can be applied to any system that can be described by a directed graph. Any quantity flowing through the network can be modeled, e.g., concentrations of a drug in different parts of the body in pharmocokinetics (compartments represent the body parts) (Rescigno, 1960), the numbers of people in different infectious stages in epidemiology (compartments represent the stages) (Kermack & McKendrick, 1927; Prabakaran *et al*., 2021), intensities of signals in different parts of a regulatory network in our study (compartments represent the regulatory parts). Each compartment is defined by the input amplification ratio and self-decay rate parameters, and the behavior of each compartment is described by an ODE with the parameters. Typically, the ODEs describing an MCM are linear, and their analytical solutions can be parametrically expressed.

We designed an MCM structure with: (1) a chain of Signaling compartments, representing a conceptualized signaling pathway common for transcriptional regulation of all genes (2) two gene-specific Response compartments, each of which draws a signal from a Signaling compartment and represents the mRNA response of the gene in one of the two cell populations (Fig. 1B). The outputs of the two Response compartments were combined to represent the mixing of mRNA from two cell populations during the mRNA sample preparation (Fig. 2). With parametrically described analytical solutions available, we fit the analytical solutions to the data, which was more computationally efficient than fitting the corresponding ODEs. This made it practical for us to fit the MCM to the RNA-seq read count data assuming a negative binomial residual distribution.

We predetermined the decay rate parameters of eleven Signaling compartments in our MCM. This was because the data to constrain the MCM were available only for gene-specific outputs of the MCM and did not have sufficient power to constrain these parameters. We fit the outputs of which Signaling compartments to use (a discrete variable), two input amplification ratio and one common decay rate parameters of two Response compartments to the data for each gene. A consequence of predetermining the Signaling compartment parameters was that the Response compartment parameters were not mechanistically interpretable. This is the reason we analyzed the kinetic characteristics we learned from the MCM (e.g., peak levels and times) instead. If more biological information about the Signaling compartments were available, such as the output of each Signaling compartment, we could build a more realistic model for the chain of Signaling compartments, which would allow mechanistic interpretations of the Response compartment parameter values.

### Elicitor(s) for the NACP transcriptome response likely originate from the responding ACP

While recognition of the effector AvrRpt2 by the cognate R protein RPS2 through the guardee/decoy RIN4 initiates the signaling event inside the ACP cells (Axtell & Staskawicz, 2003; Mackey *et al*., 2003), it is not clear what elicitor molecule(s) is recognized for initiation of signaling events in the NACP cells. There are two potential sources of elicitor(s) for transcriptome responses in NACP, which are not mutually exclusive: some signaling molecule(s) from the responding ACP; and some diffusible molecule(s) from the pathogen *Pto* AvrRpt2. Potential molecules derived from the bacterium are limited to diffusible ones because the spreading of bacterial cells from initial sites in the plant leaf is limited before 24 hpi (Zhu *et al*., 2022).

One candidate molecule class for signaling molecules originated from responding ACP cells is DAMPs (Lu & Tsuda, 2021). For example, PEP peptides are DAMPs, whose precursors are encoded by *PROPEP1* to *6* (Huffaker *et al*., 2006). The mRNA time-course pattern of *PROPEP3* showed early echoing patterns (AT5G64905 in Fig. S5). Thus, both ACP and NACP produced PEP very quickly. It is possible that the amount of PEP produced by ACP cells may not be sufficient to elicit all cells in NACP. Thus, a wave of PEP may be radially propagated through NACP from ACP cells at the center. This wave of PEP propagation could explain the time-stretched nature of the echoing transcriptome response pattern in NACP (Fig. S8B).

DAMPs are a class of molecules that can induce PTI (Yamaguchi *et al*., 2010; Liu *et al*., 2013). Another major class of molecules that can induce PTI are MAMPs, such as flg22 (Gómez-Gómez & Boller, 2000). Thus, we compared the echoing transcriptome response patterns of the two cell populations upon *Pto* AvrRpt2 inoculation with the flg22-induced transcriptome response (Hillmer *et al*., 2017) (Figs. 4A and 4B). Remarkably, the time-course patterns of the 1366 Echoing genes identified in the *Pto* AvrRpt2 response were similar in flg22-induced responses although the flg22-induced responses were generally weaker and variable (for flg22 response, the mRNA level ratio values were normalized for each gene). The similarity in the time-course patterns of NACP and flg22 responses further supports the notion that the elicitor(s) for the responses in NACP in the *Pto* AvrRpt2 response may be DAMPs. Note that the actual peak times of the flg22 response, in which all cells received flg22 at 0 hpi, is more like the peak times of the ACP response rather than the NACP response, which is consistent with the above PEP propagation wave hypothesis for the time-stretched response in NACP. Furthermore, the similarity in the time course patterns among all ACP, NACP, and flg22 responses implies that major parts of transcriptome responses in ETI and PTI share a common regulatory mechanism.

### Activation of the WRKY network may be a common signaling mechanism in two distinct cell populations

What could be the molecular mechanism that induces transcriptome responses with similar relative timing in two cell populations during ETI and in cells during PTI? We found that WRKY TFs may be important for such transcriptome responses in ACP and NACP during ETI (Figs. 6B and 6C). Eighty percent of WRKY genes upregulated upon *Pto* AvrRpt2 inoculation have WRKY-binding sites (Fig. 7C), suggesting that once some WRKY TFs are activated transcriptionally or post-transcriptionally, their encoded WRKY TF proteins bind and transcriptionally activate other WRKY genes. Many diverse WRKY genes form a transcriptionally interconnected network, and a substantial part of the WRKY network could eventually be transcriptionally activated once one or a few “entry-point” WRKYs are activated. Although the sets of WRKYs activated through different entry-point WRKYs are unlikely to be identical, such activated WRKY sets contain members of many WRKY subgroups, which could result in similar transcriptome responses downstream (Fig. 8). Any other TF genes with WRKY-binding sites and the TFs that can bind to some WRKY genes could also be part of the WRKY network (e.g., “TF-A” in Fig. 8). In fact, many WRKY genes have binding sites for diverse TFs (Fig. S10). Such “TF-A” and “TF-B” (see below) TFs could be responsible for upregulation of the approximately 30% of the Echoing genes that lack WRKY-binding sites.

**Fig. 8.**
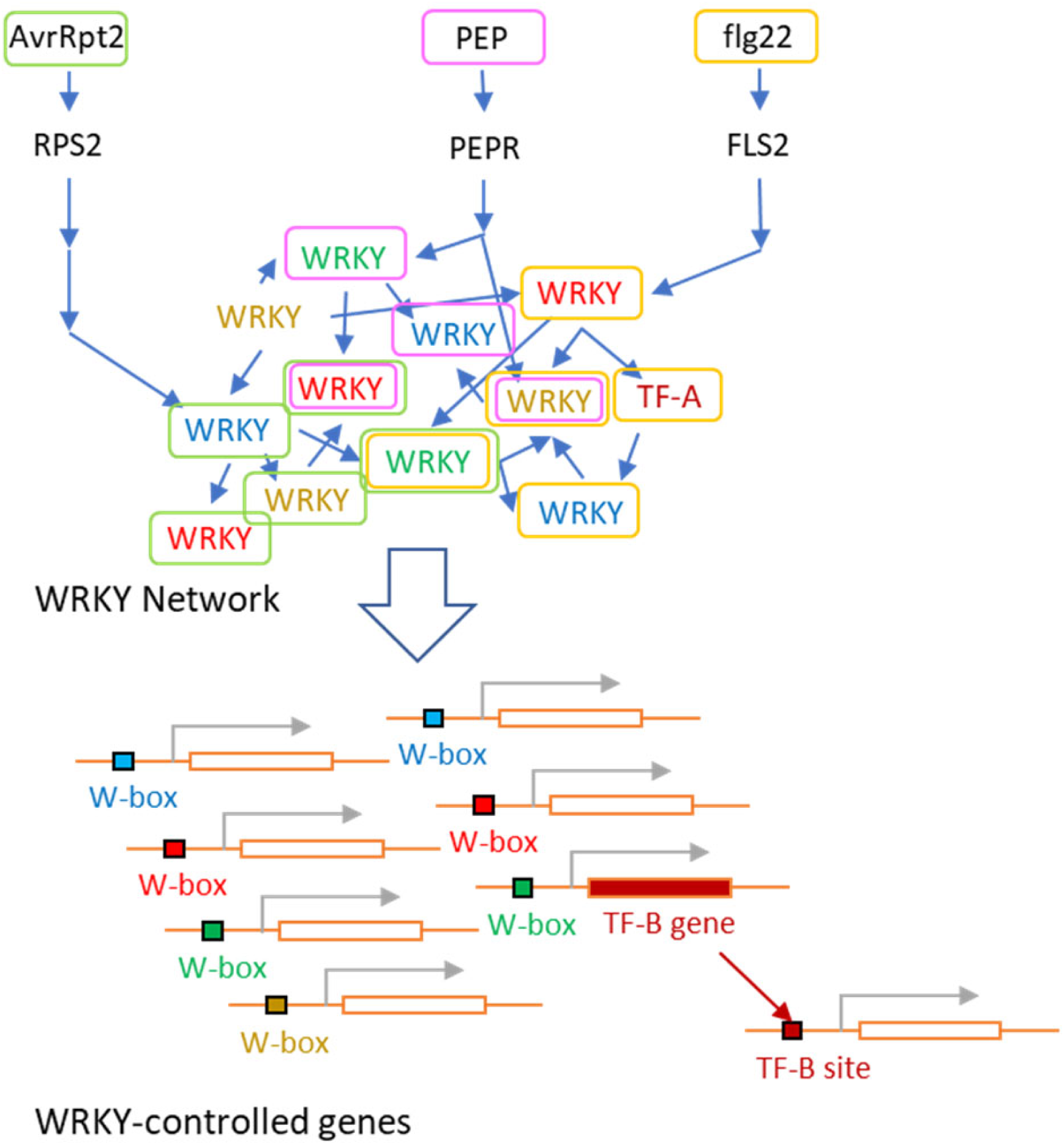
A schematic diagram of the WRKY network activation hypothesis. (Top and Middle, “WRKY Network”) The signals initiated by distinct elicitors, AvrRpt2 (an ETI elicitor), PEP (a DAMP), and flg22 (a MAMP) are fed into a transcriptional activation network of WRKY TFs through different entry-point WRKYs. Different colors of “WRKY” letters represent WRKY subgroups with somewhat different binding specificities. The elicitors and WRKYs that are activated by the elicitors are indicated by the same colors of enclosing boxes (green, pink, and orange for AvrRpt2, PEP, and flg22). Note that although different elicitors do not activate the same set of WRKYs, every elicitor activates members of all WRKY subgroups. Non-WRKY TFs (e.g., TF-A) can be part of the network if their genes have WRKY-binding sites and some WRKY genes have binding sites for the non-WRKY TFs. (Bottom “WRKY-controlled genes”) Genes with somewhat different WRKY-binding sites (W-boxes). Different W-box colors represent the binding sites for the WRKY subgroups of the same colors. Thus, as long as some members of all subgroups are activated, very similar sets of WRKY-controlled genes are upregulated, which explains the conserved transcriptome responses across the different elicitors. WRKY-controlled non-WRKY TFs (e.g., TF-B) can expand the WRKY-controlled gene set.

Consistent with this WRKY network activation hypothesis, we observed that at any time point from 4 to 24 hpi of *Pto* AvrRpt2, *WRKY* genes representing most WRKY subgroups were transcriptionally upregulated (Fig. 7A). The activation of large parts of the WRKY network during the time range in two cell populations suggests that any genes with some types of WRKY-binding sites can potentially be upregulated in either of the cell populations. Since 68% of 1366 Echoing genes have WRKY-binding sites, this potential WRKY network activation by different elicitors can explain similar gene sets for the echoing responses. Members of most WRKY subgroups were transcriptionally upregulated between 4 and 24 hpi of flg22 whereas the specific WRKY genes upregulated in ETI and PTI were not the same (Figs. 7A and B). Thus, a similar but not identical set of upregulated genes during PTI can also be explained by the WRKY network activation hypothesis (Fig. 8).

How could the conserved echoing timing in two cell populations be achieved under this WRKY network activation hypothesis? There are two possible mechanisms, which are not mutually exclusive: indirect activation through other TFs and intrinsic diversity among response genes. First, many of the upregulated genes with WRKY-binding sites encode TFs themselves. Genes that can be upregulated by such TFs would be upregulated with a delay (indirect activation through other TFs; “TF-B” in Fig. 8). Some NAC1s may be examples of “TF-B” and accumulate with a delay, which could explain a role of NAC1s in very late responses in both cell populations (Figs. 6A and 6B). Second, the time course of transcriptional activation of a particular gene could be affected by characteristics of the gene itself, which we call intrinsic diversity among response genes. For example, transcriptional activation of a particular gene with a WRKY-binding site could be modulated by other TFs and/or epigenetic mechanisms while such modulating mechanisms are conserved between ACP and NACP. Note that there is no difference between cells in ACP and NACP before inoculation. Differences in the mRNA degradation rate would also result in different peak times. In short, indirect activation and intrinsic diversity among response genes could explain conserved response timing across the Echoing genes.

The structural organization of the proposed WRKY network-based signaling mechanism can provide an advantage to plants in the evolutionary race against fast-evolving pathogens. First, the WRKY network is highly interconnected, which would provide resilience to the network against targeting by pathogen effectors. Second, the resilience of the WRKY network leaves only small parts of the immune signaling mechanism, namely the pathways specific to each signaling event, vulnerable to targeting by pathogen effectors. Such limited, vulnerable parts of the signaling mechanisms could be protected by, for example, using parallel signaling pathways connecting to different entry-point WRKYs or R proteins guarding the pathway. Third, the massive and conserved immune transcriptome response can be relatively easily repurposed or reutilized for new signaling pathways. For example, recognition of a new pathogen-associated molecule can be rapidly connected to the massive immune transcriptome response through evolution. Also, in case a specific pathway is compromised by a pathogen effector, the signal could be rapidly rerouted through a different entry-point WRKY.

It should be emphasized that in addition to this massive transcriptome response conserved across different types of immunity, there are specific transcriptome responses. We observed upregulation of the ACP- and NACP-specific genes (Fig. 4). There are upregulated genes specific to flg22-elicited PTI (Fig. S11). It appears that the massive, conserved transcriptome response provides a resilient core response across different types of immunity while different types of immunity have further adapted to their specific needs.

### Concluding remarks

In this study, we implemented an MCM to decompose the double-peak pattern, which was prevalent among the mRNA level time-courses of ETI-upregulated genes, into two single-peak patterns. We demonstrated that the decomposed first and second single-peak responses largely represent responses in two cell populations, ACP and NACP, respectively. This temporal mRNA accumulation pattern of the genes was relatively conserved during PTI as well. We propose that activation of the WRKY network is central in generation of this massive and conserved immune transcriptome response across different types of immune responses.

## Acknowledgements

This work was supported by grants from the National Science Foundation (grant nos. MCB- 0918908 and MCB-1518058 to FK and CLM and IOS-1645460 to FK) and by a grant from Ajinomoto Co., Inc. to FK. We thank Jane Glazebrook for editing the manuscript.

## Author Contributions

XL, CLM, and FK conceived the project, developed the MCM approach, analyzed the data, and wrote the manuscript. DI, RAH, TS, YL, and KT generated the unpublished RNA-seq data set (Hillmer *et al*., 2023), which formed important supporting evidence for this study.

## Supporting Information

Text S1. Detailed Materials and Methods

Fig. S1. Selection of the model time range and the high-precision genes for modeling.

Fig. S2. Spline models of 2435 high-precision upregulated genes

Fig. S3. Altered models were fit for 191 genes

Fig. S4. Selection of 1889 genes that were stably modeled with both signaling compartment decay rate parameter value Set 1 and Set 2

Fig. S5. MCM with decompositions and data points for 1889 modeled upregulated genes

Fig. S6. The distribution of the Pearson correlations between the mean estimates from the data and MCM-modeled values across 1889 modeled upregulated genes

Fig. S7. Explanations and examples of gene classification

Fig. S8. Time-scaling relationships between the first- and second-peak responses.

Fig. S9. Heatmaps of genes with consolidated TF binding sites, organized along the peak times of the first- and second-peak responses.

Fig. S10. WRKY genes have binding sites for various TFs.

Fig. S11. Core immune transcriptome response and responses specific to different elicitor types

Table S1. The fitted parameter values in MCMs with Set 1 or Set 2 of signaling compartment decay parameters.

Table S2. List of ACP-specific, Echoing 1, Echoing 2, and NACP-specific genes (each divided into early and late gene groups) and results of their GO term enrichment analysis

Table S3. List of ETI-downregulated genes (4848 genes) and results of their GO biological process term enrichment analysis.

