## Supplementary material for "Dynamic decomposition of transcriptome responses during plant effector-triggered immunity revealed conserved responses in two distinct cell populations": Text S1

Text S1. Supplemental Methods

### Data sets

Subsets of three previously published RNA-seq data sets from NCBI GEO were used: accessions GSE88798 (Mine *et al.*, 2018), GSE78735 (Hillmer *et al.*, 2017), and GSE196892 (Hillmer *et al.*, 2023). The Arabidopsis cistrome data set (O'Malley *et al.*, 2016) was also used.

### Data analysis scripts

All R scripts used to obtain results from the above-mentioned data sets are available from Github: https://github.com/fumikatagiri/MCM_At_AvrRpt2_time_course. The GLM-NB mean estimates and the upregulated gene sets from (Hillmer *et al.*, 2017; Hillmer *et al.*, 2023) were obtained according to these citations and used directly in some scripts in this study.

### Major R functions and packages

Generally, R version 4.0.2 was used for analysis, including the glm.nb function in the MASS package for negative binomial-generalized linear models (GLM-NB) with the log link function; the splinefun function for the spline model fit; the polyval function in the pracma packages for polynomial regression model value calculation; the qvalue package for the Storey FDR correction; the ComplexHeatmap package for heatmap generation; and the ggplot2 package for plot generation in general. The Curry function in the functional package was used in MCM fitting scripts for partly implementing functional programming. The mpfr function in the Rmpfr package, which is for high-precision calculation, was used in MCM fitting because the calculation of the MCM analytical solution requires high precision when the self-decay rate of the response compartment, $k_g$, is very close to one of the self-decay rates for the signaling compartment. The actual fitted $k_g$ values across the 1989 modeled upregulated genes were not too close to any of the self-decay rates for the signaling compartment, and thus, the Rmpfr package was not used in calculation of MCM-modeled values in the interest of computing time. Since execution of MCM fitting with high precision required a long computing time, the fitting procedure was parallelized. The R packages, doParallel and foreach were used for this purpose. Shell scripts (.sh) were also used when parallelized R scripts were executed.

### Spline time-course model for each of the 2435 high-precision upregulated genes

For each gene at each time point, the $\log_2$-transformed GLM mean estimate for mock was subtracted from the $\log_2$-transformed RNA-seq read count data after between-libraries normalization based on the $90^{th}$ percentile read count, which was used in GLM fitting. These values were used as the $\log_2$FC data to fit a spline model. Since time-course curvatures were generally sharper at early hpi, the square-root hpi values were used in the model to make the curvature relatively even. The data points and the fitted spline models for 2435 genes are shown in Fig. S2.

**The structure and the parameters of the MCM**

As our MCM had an acyclic structure and was a linear ODE system, the output (analytical solution) of each compartment can be parametrically expressed, and the analytical solution for $R_{A,g} + R_{N,g}$ was fit to the data for gene $g$.

Since we did not observe much response before 3 hpi (Fig. S1A), we set 3 hpi as 0 hour of the model time.

The output for each of the signaling compartments, $S^{(n)}(t)$, $n = 0,1, \dots ,11$, is expressed as ((13) and (26) of Appendix 1 at the end of this Text S1):

$$S^{(n)}(t) \ = \prod_{i=1}^{n} a_i \cdot \left\{ \sum_{i=0}^{n} \left( e^{-k_i t} \cdot \prod_{j=0, j \neq i}^{n} \frac{1}{k_j - k_i} \right) \right\} \dots (1)$$

where $a_i$ is the input amplification ratio and $k_i$ is the first-order self-decay rate of the $i$th compartment. For simplicity, we normalized the output of each signaling compartment to have the area-under-curve of 1, i.e., $\int_0^\infty S^{(n)}(t)dt = 1$. This can be achieved by setting the input amplification ratio parameter, $a_n = k_n$ (Appendix 2 at the end of Text S1). Note that: (1) the shape and the time scale, including the peak time, of the single-peak time-course output are not affected by the input amplification rates of the signaling compartments; (2) if the signaling compartment input amplification ratio are not fixed, the model cannot be fit due to the singularity.

We used two sets of the predetermined decay rate values for the signaling compartments, Set 1 and Set 2, which generated two sets of predetermined peak times of the signaling compartment outputs. The decay-rate parameter values for Sets 1 and 2 in the order of signaling compartments 0 to 11 were: (Set 1) 1.8, 0.5487, 1.1733, 1.1253, 0.6492, 0.6853, 0.6957, 0.6987, 0.4949, 0.5065, 0.513, and 0.5174; (Set 2) 1.8, 2.4573, 1.0775, 1.3624, 0.6624, 0.7903, 0.646, 0.6757, 0.5669, 0.5814, 0.5003, and 0.5094. The generated peak times of the outputs of signaling compartments 1, 3, 5, 7, 9, and 11 were: Set 1, ($[4, 6, 9, 12, 16, 20] * 0.95$) hpi and Set 2, ($[3.5, 5, 7.5, 10.5, 14, 18] * 0.95$) hpi. We used the outputs from every second signaling compartment because a linear combination of the outputs from two consecutive signaling compartments cannot generate a double-peak pattern but only a single-peak pattern (Appendix 3 at the end of Text S1). However, we found that the data did not have sufficient power to efficiently discriminate 15 possible combinations of two from signaling compartments 1, 3, 5, 7, 9, and 11. This is the reason we used combinations of two signaling compartments with further spacing : (1, 5), (1, 7), (1, 9), (1, 11), (3, 7), (3, 9), (3, 11), (5, 9), (5, 11), and (7, 11). The peak times for Set 1 were chosen according to the data time points, and those for Set 2 were chosen to take middle time points relative to those for Set 1. In this way, the two sets of peak times were relatively orthogonal, and we thought that model fitting results consistent between Sets 1and 2 were unlikely to have strong biases caused by the predetermination of the parameter values.

As our MCM has an acyclic structure and is a linear ODE system, the model can be considered as a linear combination of two chains of MCMs. For example, if the outputs of signaling compartments $S^{(2)}$ and $S^{(4)}$ are used as the inputs to response compartments $R_{A,g}$ and $R_{N,g}$, respectively, the model structure can be considered as a linear combination of two MCM chains, $S^{(0)} \rightarrow S^{(1)} \rightarrow S^{(2)} \rightarrow R_{A,g}$ and $S^{(0)} \rightarrow S^{(1)} \rightarrow S^{(2)} \rightarrow S^{(3)} \rightarrow S^{(4)} \rightarrow R_{N,g}$, where the linear coefficients for the chains are $a_{A,g}$, and $a_{N,g}$, respectively. Thus, the analytical solution of the MCM can be expressed with $k_g$, $a_{A,g}$, and $a_{N,g}$ using equation (1) for each chain, while the parameter values for $S^{(n)}$ were predetermined (either Set 1 or Set 2).

**Fitting polynomial regression model to mock read count data**

For each gene a polynomial regression was fit to the mock time-course read count data using GLM-NB with the between-libraries normalization offset. The polynomial regression for mock was included as an offset in the MCM. In the polynomial regression model, the time scale was first square-root-transformed as was done with the spline model described above, and the transformed time values were further linearly transformed to the range [-1,1] for stable polynomial model fitting. Regression models up to the $5^{th}$-order polynomial were fit, and the best model was chosen according to Akaike's information criterion (AIC).

**Fitting the MCM**

For the $g$th gene, four parameters, $k_g$, two input amplification rates of the response compartments, $a_{A,g}$ and $a_{N,g}$ (Fig. 2B), and a discreate signaling compartment combination variable were fit in the MCM, given one of Sets 1 or 2 for the signaling compartment decay rate parameter values. The MCM had two offsets, log(mock polynomial fitted values) and the between-libraries normalization term based on the $90^{th}$ percentile read counts. The response of the MCM was the *Pto* AvrRpt2 time-course read count data.

Since the MCM models used log(*Pto* AvrRpt2/mock), we expected $R_{A,g} + R_{N,g} = 0$ at model times $t = 0$ and $t = \infty$. To satisfy these conditions, we assumed the output of each compartment to be 0 at $t = 0$ and $t = \infty$. Since we did not observe many responses before 3 hpi across the genes (Fig. S1A), we set 3 hpi as the 0 hour for the model times.

Model fitting was based on the maximum likelihood method on the log scale as we expected negative binomial residual distributions with the read count data. In practice, for each gene, models with different signaling compartment combinations were fit separately, and the best fit model among them was chosen as the fitted value for the discreate signaling compartment combination variable.

**Altered MCMs for the genes that had time-course patterns of a single peak with a shoulder**

We used two different empirical methods to select the genes that have clearly higher modeled values than data values at 6 hpi. First, 176 genes were selected that had MCM value – GLM mean estimate > 0.55 at 6 hpi. Second, an additional 15 genes were selected that were the intersect of: (i) all three read count data are lower than 0.85 * MCM value at 6 hpi; (ii) the mean of three read count data > 0.6 at 6 hpi; (iii) the second-peak time > 15 hpi; (iv) the first-peak time was 5 to 7 hpi; (v) 0.9 * first-peak level > second-peak level; (vi) mean of three read count data at 12 hpi < 0.75 * mean of three read count data at 6 hpi. To impose MCM to capture these very early second peaks, an altered model was fit to only the data up to 16 hpi. Fig. S3 shows how the altered MCMs changed from the original MCMs for these genes. These early second-peak genes often had third peaks, which were not captured by the MCM designed for double peaks (the bottom 191 genes in Fig. 3A). Some of the genes had relatively early third peaks, which already started rising at 16 hpi, and the fit of the altered MCM was not ideal. However, the number of such genes was low, and we did not try to fit another special model for these genes.

**Double-peak patterns were not model artifacts**

As our MCM imposes double-peak pattern interpretations of the data, the wide-spread double-peak phenomenon detected by the MCM approach might be a model artifact. To demonstrate that it is not an artifact for most of the modeled genes, the spline model described above, which does not impose particular patterns, was used. For each gene, the peak times were determined by the derivative of the spline model = 0 and the second derivative < 0. If there is only one such peak time for a gene, it was considered as a single-peak time course. If there are more than two such peak times for a gene, two time points that gave the largest log-ratio fitted values were chosen. Then, the time-course pattern was decomposed as follows: (1) the part earlier than the first peak time and that later than the second peak time were assigned to the first-peak and second-peak responses; (2) the part between the first and second peak times was linearly decomposed according to the ratio of time differences from the first and second peak times. The two peak responses decomposed in this manner were similar to those obtained by MCM for most of the 1366 Echoing genes, indicating that, for most genes, the MCMs did not introduce artifactual double peak patterns (Fig. S2B).

**Time-scaling relationships between the first- and second-peak responses**

To test whether the first-peak and second-peak responses are approximately time-scaled from each other, the ratio of the second-peak time over the first-peak time and the ratio of the second-peak width over the first-peak width were examined for 1366 Echoing genes. For the width of a peak, the width at 75% peak level was used. We also tried the width at 50% peak level and obtained very close ratio values.

**GO term enrichment analysis**

Two classifiers were used for the gene groups, the first-peak level/second-peak level ratio and the peak times, so this is a 2-dimentional classification: four first-peak level/second-peak level ratio categories, ACP-specific, Echoing 1, Echoing 2, NACP-specific genes; and two peak time categories, early and late. For the peak time, the first-peak time was used for ACP-specific, Echoing 1, and Echoing 2 genes. The second-peak time was used for NACP-specific genes. Then the genes in each level ratio category were divided into halves according to the peak time. Thus, we generated eight gene groups: ACP-specific early and late, Echoing 1 early and late, Echoing 2 early and late, and NACP-specific early and late. The list of genes for each gene group is included in Table S2.

Panther Classification System (http://go.pantherdb.org/) version 17.0 with GO Ontology database DOI: 10.5281/zenodo.7942786 was used for the GO term enrichment analysis (Mi *et al.*, 2019; Thomas *et al.*, 2022). For each of the eight gene groups with each GO term category, terms with Bonferroni-corrected $p < 0.05$ were returned. The analysis results for 8 gene groups x 3 GO term categories = 24 results are included in Table S2. For further processing of the results with biological process GO terms, only the lowest term in each hierarchy was selected ("hierarchy" column value, 1, in Table S2), as hierarchically higher terms are often too broad. Among the hierarchically lowest terms, those with fold enrichment > 10-fold or Bonferroni-corrected $p < 10^{-8}$ in at least one gene group were selected, as shown in Table 1. If the terms were selected in multiple gene groups under the initial conditions of Bonferroni-corrected $p < 0.05$, their fold enrichment values are included in Table 1. Otherwise, the cells in the table show "-".

**TF-binding sites enrichment analysis along the peak times**

This analysis used the cistrome data set consisting of experimentally determined binding sites for 349 TFs across more than 30,000 genes in Arabidopsis (http://neomorph.salk.edu/dap_web/pages/index.php) (O'Malley *et al.*, 2016). For each gene, the DNA sequence 1000 bp upstream to 200 bp downstream of the transcription start site was defined as the promoter region and searched for TF binding sites. In Fig. 6A, (214 ACP-specific + 1366 Echoing) or (1366 Echoing + 227 NACP-specific) gene sets for the first-peak response and the second-peak sets were used. Fisher's exact test (1-sided) was used to calculate the *p*-

value for enrichment of genes with each TF-binding site among ACP-specific, Echoing, or NACP-specific genes compared to the proportions of such genes in the genome. The union of the TFs with Benjamini-Hochberg FDR-corrected $p < 0.01$ contained 100 TFs.

For these 100 TFs, over- or under-representation of genes with their binding sites along the peak time compared to the proportion of the genes with their binding sites in each corresponding gene set was evaluated. Subsets of each gene set defined by a sliding window of 150 genes with a sliding step of 15 genes along the peak time were made. The *p*-value was calculated using Fisher's exact test (2-sided) for the number of the genes with binding sites for each TF in each gene subset, compared to the proportion of the genes with binding-sites for the TF in the gene set. We selected 38 TFs with $p < 0.001$ in at least one window subset for further study.

The over- and under-representation results for each of the TF families, NAC1s, NAC2s, WRKYs, and HSFs were consolidated in Fig. S9. For each consolidated TF family, the genes with binding sites for any of the consolidated TF family members were counted as genes with the binding site for the consolidated TF family.

Over- and under-representation analysis along the peak time, which was essentially the same as the analysis above, was performed for the consolidated TF families and singleton TFs with the first-peak and second-peak response gene sets in Fig. S9 and for the consolidated TF families and singleton TFs with 1366 Echoing genes only in Fig. 6B. In Fig. 6B, only the TF families and the singleton TF that had $p < 0.001$ in at least one window subset in each of the first- and the second-peak responses were selected.

**Appendix 1. Analytical solutions for the outputs of the signaling compartments.**

The signaling compartments are a series of multiple compartments. The compartments are controlled by the self-decay rate and the input amplification ratio parameters.

We set the self-decay rate of the $i$th compartment in a series of compartments, $k_i$ as follows:

$k_i > 0$ for $\forall i \in \{0,1,2,\ldots\}$ … (1)

And $k_i \neq k_j$ for any $i \neq j$ … (2)

When the $n$th compartment ($n > 0$) has the input amplification ratio $a_n$, the output of the compartment $S^{(n)}(t)$ is expressed by an ODE:

$$\frac{dS^{(n)}}{dt}(t) = a_n S^{(n-1)}(t) - k_n S^{(n)}(t) \ldots (3)$$

(3) can be varied to:

$$\frac{dS^{(n)}}{dt}(t) + k_n S^{(n)}(t) = a_n S^{(n-1)}(t) \ldots (3')$$

Solve the homogeneous equation of (3'):

$$\frac{dS^{(n)}}{dt}(t) + k_n S^{(n)}(t) = 0 \ldots (4)$$

The general solution of (4) is:

$S^{(n)}(t) = C^{(n)} e^{-k_n t}$, where $C^{(n)}$ is an integration constant … (5)

Apply a variation of constant to (3') and (5)

$$\frac{dC^{(n)}}{dt}(t) = a_n e^{k_n t} S^{(n-1)}(t) \ldots (6)$$

If we hypothesize that $S^{(n-1)}(t)$ can be expressed as a linear combination of first-order decays specified by the decay rate values $k_0, k_1, \ldots, k_{n-1}$, and the linear coefficients, $b_1^{(n-1)}, b_2^{(n-1)}, \ldots, b_{n-1}^{(n-1)}$:

$$S^{(n-1)}(t) = \sum_{i=0}^{n-1} b_i^{(n-1)} e^{-k_i t} \ldots (7)$$

Assign (7) to (6)

$$\frac{dC^{(n)}}{dt}(t) = a_n e^{k_n t} \sum_{i=0}^{n-1} b_i^{(n-1)} e^{-k_i t} = a_n \sum_{i=0}^{n-1} b_i^{(n-1)} e^{-(k_i - k_n)t} \ldots (8)$$

Integrate (8)

$$C^{(n)}(t) = a_n \int \sum_{i=0}^{n-1} b_i^{(n-1)} e^{-(k_i - k_n)t}\, dt = a_n \sum_{i=0}^{n-1} b_i^{(n-1)} \int e^{-(k_i - k_n)t} dt$$

$= a_n \sum_{i=0}^{n-1} \frac{b_i^{(n-1)}}{k_n - k_i} e^{-(k_i - k_n)t} + P^{(n)}$, where $P^{(n)}$ is an integration constant … (9)

Assign (9) to (5),

$$S^{(n)}(t) = \sum_{i=0}^{n-1} \frac{a_n b_i^{(n-1)}}{k_n - k_i} e^{-k_i t} + P^{(n)} e^{-k_n t} \quad \ldots (10)$$

Thus, if hypothesis (7), that ($S^{(n-1)}(t)$ can be expressed as a linear combination of first-order decays specified by the decay rate values $k_0, k_1, \ldots, k_{n-1}$) is true, $S^{(n)}(t)$ is also a linear combination of first-order decays specified by the decay rate values $k_0, k_1, \ldots, k_n$. … (11)

We assume that the $0^{th}$ compartment, which does not have any input, has the output of:

$$S^{(0)}(t) = a_0 e^{-k_0 t} \quad \ldots (12)$$

Then by induction from (12) and (11), $S^{(n)}(t)$ is indeed a linear combination of first-order decays specified by the decay rate values $k_0, k_1, \ldots, k_n$.

i.e., $S^{(n)}(t)$ can be generally expressed as:

$$S^{(n)}(t) = \sum_{i=0}^{n} b_i^{(n)} e^{-k_i t} \quad \ldots (13)$$

We also assume the initial condition, $S^{(n)}(t) = 0$ for $\forall n > 0$ … (14)

From (14) and (10)

$$P^{(n)} = -\sum_{i=0}^{n-1} \frac{a_n b_i^{(n-1)}}{k_n - k_i} \quad \ldots (15)$$

Thus, (10) becomes

$$S^{(n)}(t) = \sum_{i=0}^{n-1} \frac{a_n b_i^{(n-1)}}{k_n - k_i} e^{-k_i t} - \sum_{i=0}^{n-1} \frac{a_n b_i^{(n-1)}}{k_n - k_i} e^{-k_n t} \quad \ldots (16)$$

Comparison of (13) and (16) leads to:

$b_i^{(n)} = \frac{a_n b_i^{(n-1)}}{k_n - k_i}$, when $i < n$ … (17)

(17) can be expressed as:

$$b_i^{(n)} = b_i^{(i)} \prod_{j=i+1}^{n} \frac{a_j}{k_j - k_i} \quad \ldots (18)$$

$b_i^{(i)}$ is the integration constant to make $f^{(i)}(0) = 0$, so from (16):

$$b_i^{(i)} = -\sum_{j=0}^{i-1} \frac{a_i b_j^{(i-1)}}{k_i - k_j} \quad \ldots (19)$$

From (12), $b_0^{(0)} = a_0$ … (20)

Assign $i = 0$ to (18) and further assign (20) to it. When $n > 0$

$$b_0^{(n)} = a_0 \prod_{j=1}^{n} \frac{a_j}{k_j - k_0} = \prod_{j=0}^{n} a_j \prod_{j=0, j\neq 0}^{n} \frac{1}{k_j - k_0} \quad \dots (21)$$

From (19) and (21),

$$b_1^{(1)} = -b_0^{(1)} = -a_0 a_1 \frac{1}{k_1 - k_0} \quad \dots (22)$$

Assign $i = 1$ and (22) to (18). When $n > 1$,

$$b_1^{(n)} = -a_0 \frac{a_1}{k_1 - k_0} \prod_{j=2}^{n} \frac{a_j}{k_j - k_1} = \prod_{j=0}^{n} a_j \prod_{j=0, j\neq 1}^{n} \frac{1}{k_j - k_1} \quad \dots (23)$$

Similarly to (22) and (23), assign (21) and (23) to (19):

$$b_2^{(2)} = -b_0^{(2)} - b_1^{(2)} = -\prod_{j=0}^{2} a_j \prod_{j=0, j\neq 0}^{2} \frac{1}{k_j - k_0} - \prod_{j=0}^{2} a_j \prod_{j=0, j\neq 1}^{2} \frac{1}{k_j - k_1}$$

$$= -\prod_{j=0}^{2} a_j \left( \frac{1}{(k_1 - k_0)(k_2 - k_0)} + \frac{1}{(k_0 - k_1)(k_2 - k_1)} \right) = \prod_{j=0}^{2} a_j \left( \frac{1}{(k_0 - k_2)(k_1 - k_2)} \right)$$

$$= \prod_{j=0}^{2} a_j \prod_{j=0, j\neq 2}^{2} \frac{1}{k_j - k_2} \quad \dots (24)$$

(24) suggests in general:

$$b_i^{(i)} = \prod_{j=0}^{i} a_j \prod_{j=0, j\neq i}^{i} \frac{1}{k_j - k_i}, \text{ when } i > 0 \quad \dots (25)$$

Assign (25) to (18)

$$b_i^{(n)} = \prod_{j=0}^{i} a_j \prod_{j=0, j\neq i}^{i} \frac{1}{k_j - k_i} \prod_{j=i+1}^{n} \frac{a_j}{k_j - k_i}$$

$$= \prod_{j=0}^{n} a_j \prod_{j=0, j\neq i}^{n} \frac{1}{k_j - k_i} \quad \dots (26)$$

We will prove (26).

If (26) is true for $n-1, i$ when $0 < i \leq n-1$, is it true for $n$ and $i$ when $0 < i \leq n$? … (27)

Assign $n = n - 1$ to (26) then to (17)

$$b_i^{(n)} = \frac{a_n b_i^{(n-1)}}{k_n - k_i} = \frac{a_n}{k_n - k_i} \prod_{j=0}^{n-1} a_j \prod_{j=0, j\neq i}^{n-1} \frac{1}{k_j - k_i} = \prod_{j=0}^{n} a_j \prod_{j=0, j\neq i}^{n} \frac{1}{k_j - k_i}$$

This is the same as (26). Thus, (27) is true when $0 < i \leq n - 1$ … (28)

From (19), $b_n^{(n)}$ is:

$$b_n^{(n)} = -\sum_{j=0}^{n-1} \frac{a_n b_j^{(n-1)}}{k_n - k_j} \quad \dots (29)$$

Assign $n = n - 1$ to (26) then to $b_j^{(n-1)}$ of (29) (indices were changed to avoid an overlap of $j$)

$$b_n^{(n)} = -\sum_{j=0}^{n-1} \left( \frac{a_n}{k_n - k_j} \prod_{m=0}^{n-1} a_m \prod_{m=0, m \neq j}^{n-1} \frac{1}{k_m - k_j} \right) = -\prod_{m=0}^{n} a_m \sum_{j=0}^{n-1} \left( \prod_{m=0, m \neq j}^{n-1} \frac{1}{k_n - k_j} \frac{1}{k_m - k_j} \right)$$

$$= -\prod_{m=0}^{n} a_m \sum_{j=0}^{n-1} \left( \prod_{m=0, m \neq j}^{n} \frac{1}{k_m - k_j} \right) \quad \dots (30)$$

Dividing (25) and (30) by $\prod_{m=0}^{n} a_m$, now the question is whether the following (31) is true.

$$\prod_{j=0, j \neq n}^{n} \frac{1}{k_j - k_n} = -\sum_{j=0}^{n-1} \left( \prod_{m=0, m \neq j}^{n} \frac{1}{k_m - k_j} \right) \quad \dots (31)$$

Change the indices in (31) for clarity.

$$\prod_{m=0, m \neq n}^{n} \frac{1}{k_m - k_n} = -\sum_{j=0}^{n-1} \left( \prod_{m=0, m \neq j}^{n} \frac{1}{k_m - k_j} \right)$$

Move the term on the right hand to the left hand.

$$\prod_{m=0, m \neq n}^{n} \frac{1}{k_m - k_n} + \sum_{j=0}^{n-1} \left( \prod_{m=0, m \neq j}^{n} \frac{1}{k_m - k_j} \right) = 0$$

$$\sum_{j=0}^{n} \left( \prod_{m=0, m \neq j}^{n} \frac{1}{k_m - k_j} \right) = 0 \quad \dots (31')$$

It was manually confirmed that (31') was true for $n \leq 4$. Because the left side of (31') is symmetric to any interchanges among $k_i$, $i = 0, 1, \dots, n$, (31') must be generally true. Because (31') is true, (25) and (30) are equivalents, and thus (25) is true. Since (25) is true, (26) is true as well.

**Conclusions**:

When $S^{(n)}(t)$ is

(i) expressed by the ODE $\frac{dS^{(n)}}{dt}(t) = a_n S^{(n-1)}(t) - k_n S^{(n)}(t)$ … (3)
(ii) We set $S^{(0)}(t) = a_0 e^{-k_0 t}$ … (12)
(iii) We use the initial conditions, $S^{(i)}(0) = 0$ for $i = 1, \dots, n$ … (14)

Then $S^{(n)}(t)$ is expressed by

(iv) a linear combination of first-order decays specified by the decay rate values $k_0, k_1, \dots, k_n$. i.e., $S^{(n)}(t) = \sum_{i=0}^{n} b_i^{(n)} e^{-k_i t}$ … (13)
(v) where $b_i^{(n)} = \prod_{j=0}^{n} a_j \prod_{j=0, j \neq i}^{n} \frac{1}{k_j - k_i}$ … (26)

**Appendix 2. Normalization of the signaling compartment output to AUC = 1**

In the following, we explain how to set the input amplification ratio parameter value to make the area under curve (AUC) of the time-course function for the output of each compartment in a series of compartments 1, as we did in the signaling compartments we used.

GOAL: Normalize $S^{(n)}(t)$ defined in Appendix 1. to have $\int_0^\infty S^{(n)}(t)dt = 1$ … (32), i.e., AUC=1.

If $S^{(n-1)}$ is already normalized to $\int_0^\infty S^{(n-1)}(t)dt = 1$ … (32):

From (3)

$S^{(n)}(t) = \frac{1}{k_n}\left\{a_n S^{(n-1)}(t) - \frac{dS^{(n)}(t)}{dt}\right\}$ … (3'')

Integrate (3'') from 0 to ∞,

$\int_0^\infty S^{(n)}(t)dt = \frac{1}{k_n}\left\{a_n \int_0^\infty S^{(n-1)}(t)dt - S^{(n)}(\infty) + S^{(n)}(0)\right\}$

From (32), $S^{(n)}(\infty) = 0$ (from (13), the intrinsic characteristics of the function) and $S^{(n)}(0) = 0$ (from (14), the initial condition)

$\int_0^\infty S^{(n)}(t)dt = \frac{a_n}{k_n}$ ... (33)

Thus, if $\int_0^\infty S^{(n-1)}(t)dt = 1$, $\int_0^\infty S^{(n)}(t)dt = 1$ by setting $a_n = k_n$ … (34)

For $S^{(0)}(t) = a_0 e^{-k_0 t}$ … (12), if we set $a_0 = k_0$, $\int_0^\infty S^{(0)}(t)dt = 1$ … (35)

Using induction from (35) and (34), if we set the input amplification ratio parameter to be the same as the decay rate parameter for each compartment, $a_i = k_i$, $i = 0,1,\ldots,n$,

$\int_0^\infty S^{(n)}(t)dt = 1$ …. (36)

**Appendix 3. Two consecutive signaling compartments cannot generate a double-peak time-course pattern when their outputs are linearly combined.**

We consider the outputs of two consecutive signaling compartments in a chain of multiple compartments, $S^{(n)}(t)$ and $S^{(n+1)}(t)$, $(n > 0)$. Then by definition:

$$\frac{dS^{(n+1)}(t)}{dt} = S^{(n)}(t) - k_{n+1}S^{(n+1)}(t) \ldots (1)$$

At the peak time for $S^{(n+1)}(t)$, which is designed as $t_p^{(n+1)}$,

$$\frac{dS^{(n+1)}}{dt}\left(t_p^{(n+1)}\right) = 0, \text{ thus,}$$

$$S^{(n)}\left(t_p^{(n+1)}\right) = k_{n+1}S^{(n+1)}\left(t_p^{(n+1)}\right) \ldots (2)$$

(2) means that the curves, $S^{(n)}(t)$ and $k_{n+1}S^{(n+1)}(t)$ cross at $t = t_p^{(n+1)}$, i.e., at the peak of $k_{n+1}S^{(n+1)}(t)$ (Fig ST1). Therefore,

When $t > t_p^{(n+1)}$, $S^{(n)}(t) < k_{n+1}S^{(n+1)}(t) \ldots (3)$

When $t < t_p^{(n+1)}$, $S^{(n)}(t) > k_{n+1}S^{(n+1)}(t) \ldots (4)$

$k_{n+1}S^{(n+1)}(t)$ decreases slower than $S^{(n)}(t)$ at any time points in $t > t_p^{(n+1)}$ (Fig ST1), which is expressed as:

When $t > t_p^{(n+1)}$, $\frac{dS^{(n)}}{dt}(t) - k_{n+1}\frac{dS^{(n+1)}}{dt}(t) < 0 \ldots (5)$

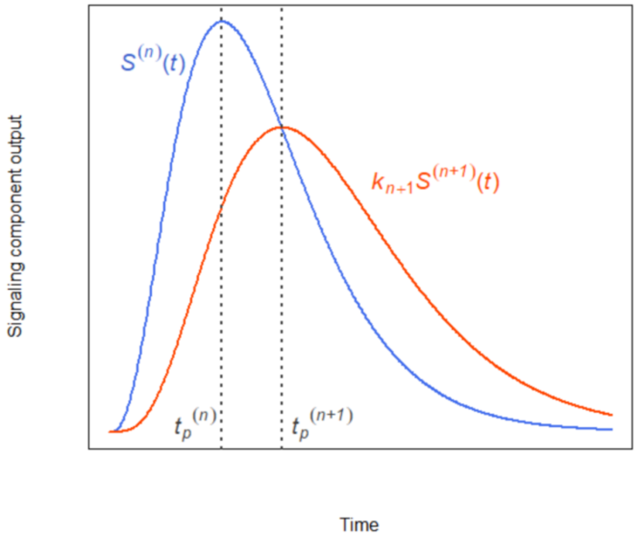


Fig ST1. $S^{(n)}(t)$ and $k_{n+1}S^{(n+1)}(t)$ take the same value at $t_p^{(n+1)}$, which is the peak time for $S^{(n+1)}(t)$. An example of $S^{(n)}(t)$ (blue) and $k_{n+1}S^{(n+1)}(t)$ (red) and their peak times $t_p^{(n)}$ and $t_p^{(n+1)}$ are shown.

Although we summed the outputs of two response compartments, $R_{A,g}$ and $R_{N,g}$, in our MCM (Fig. 2B), it is equivalent to feeding a linear combination of two signaling compartment outputs as the input of a single response compartment $R_g$ with the decay rate of $k_g$, using the amplitude parameters $a_{A,g}$ and $a_{N,g}$ as linear coefficients (Fig ST2B). This is because the MCM we used is a linear ODE system and because $R_{A,g}$ and $R_{N,g}$ share the same decay rate of $k_g$.

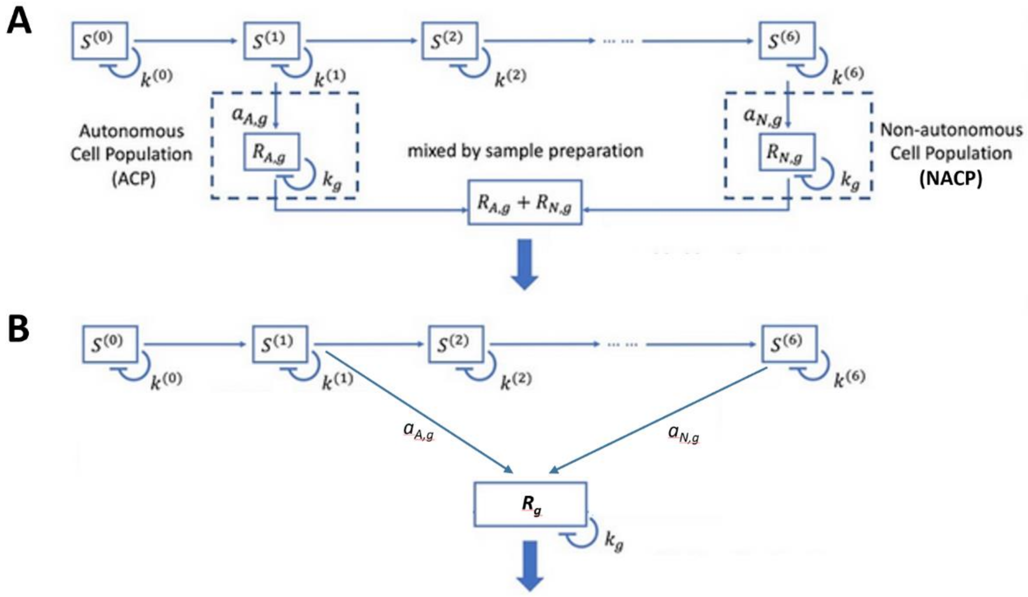


Fig ST2. Two equivalent MCM structures. (A) The MCM structure in Fig. 1B. (B) An equivalent MCM structure, in which the inputs to a single response compartment $R_g$ are linearly combined before the compartment.

Thus, to obtain a double-peak time-course output of the response compartment, the linear combination of two signaling compartment outputs must have a double-peak time-course. We denote a linear combination of outputs of two consecutive signaling compartments, $S^{(n+1)}(t)$ and $S^{(n+2)}(t)$, as $G^{(n+2)}(t)$:

$$G^{(n+2)}(t) = a_{n+1}S^{(n+1)}(t) + a_{n+2}S^{(n+2)}(t) \ldots (6)$$

where $a_{n+1}$ and $a_{n+2}$ ($a_{n+1}, a_{n+2} > 0$) are linear coefficients.

By taking the derivative of (6):

$$\frac{dG^{(n+2)}}{dt}(t) = a_{n+1}\frac{dS^{(n+1)}}{dt}(t) + a_{n+2}\frac{dS^{(n+2)}}{dt}(t) \ldots (7)$$

We consider the time range, $t_p^{(n+1)} \le t \le t_p^{(n+2)} \ldots (8)$, which should contain all the peak(s) of $G^{(n+2)}(t)$.

From (3)

For $t_p^{(n+1)} < \forall t \leq t_p^{(n+2)}$, $\frac{dS^{(n+1)}}{dt}(t) = S^{(n)}(t) - k_{n+1}S^{(n+1)}(t) < 0 \ldots$ (9)

From (4) (by adding 1 to the index)

For $t_p^{(n+1)} \leq \forall t < t_p^{(n+2)}$, $\frac{dS^{(n+2)}}{dt}(t) = S^{(n+1)}(t) - k_{n+1}S^{(n+2)}(t) > 0 \ldots$ (10)

Furthermore,

$\frac{dS^{(n+1)}}{dt}\left(t_p^{(n+1)}\right) = 0 \ldots$ (11) AND

$\frac{dS^{(n+2)}}{dt}\left(t_p^{(n+2)}\right) = 0 \ldots$ (12)

From (7), (10), and (11),

$\frac{dG^{(n+2)}}{dt}\left(t_p^{(n+1)}\right) > 0 \ldots$ (13)

From (7), (9), and (12),

$\frac{dG^{(n+2)}}{dt}\left(t_p^{(n+2)}\right) < 0 \ldots$ (14)

From (13) and (14), $G^{(n+2)}(t)$ has at least one peak in $t_p^{(n+1)} < t < t_p^{(n+2)}$.

If $\frac{dG^{(n+2)}}{dt}(t)$ is monotonically decreasing in $t_p^{(n+1)} < t < t_p^{(n+2)}$ (i.e., $\frac{d^2G^{(n+2)}}{dt^2}(t) < 0$ for $t_p^{(n+1)} < \forall t < t_p^{(n+2)} \ldots$ (15)), $G^{(n+2)}(t)$ has only one peak in $t_p^{(n+1)} < t < t_p^{(n+2)}$. In other words, condition (15) is sufficient for $G^{(n+2)}(t)$ to have only one peak.

By taking the derivative of (7):

$\frac{d^2G^{(n+2)}}{dt^2}(t) = a_{n+1}\frac{d^2S^{(n+1)}}{dt^2}(t) + a_{n+2}\frac{d^2S^{(n+2)}}{dt^2}(t) \ldots$ (16)

By taking the derivative of (1) (and by adding 1 to the index), together with (5):

In $t_p^{(n+1)} < t < t_p^{(n+2)}$, $\frac{d^2S^{(n+1)}}{dt^2}(t) = \frac{dS^{(n)}}{dt}(t) - k_{n+1}\frac{dS^{(n+1)}}{dt}(t) < 0 \ldots$ (17)

Also, in $t_p^{(n+1)} < t < t_p^{(n+2)}$, $\frac{dS^{(n+1)}}{dt}(t) < 0 \ldots$ (18) (after the peak of $S^{(n+1)}(t)$) and $\frac{dS^{(n+2)}}{dt}(t) > 0 \ldots$ (19) (before the peak of $S^{(n+2)}(t)$). From (18) and (19) with the derivative of (1) (by adding 2 to the index),

When $t_p^{(n+1)} < t < t_p^{(n+2)}$, $\frac{d^2S^{(n+2)}}{dt^2}(t) = \frac{dS^{(n+1)}}{dt}(t) - k_{n+1}\frac{dS^{(n+2)}}{dt}(t) < 0 \ldots$ (20)

By applying (17) and (20) to (16):

$\frac{d^2G^{(n+2)}}{dt^2}(t) < 0 \ldots$ (21)

(21) demonstrates that the condition (15) is satisfied. Therefore, $G^{(n+2)}(t)$ always has only one peak for any $a_{n+1}$ and $a_{n+2}$. In other words, a linear combination of the outputs of two consecutive signaling compartments cannot generate a double-peak time-course pattern.
