## Supplemental Figures for "Dynamic decomposition of transcriptome responses during plant effector-triggered immunity revealed conserved responses in two distinct cell populations"

**Fig. S1. Selection of the model time range and the high-precision genes for modeling.**

**A**. Relative to other time points, very little response was observed at 3 hpi and earlier for most of the 18442 significantly responding genes. The $log_2$ mRNA level ratios of *Pto* AvrRpt2 over mock are shown in a heatmap.  Thus, we decided to use the data at 4 hpi and later and set the model time 0 of the MCM at 3 hpi.

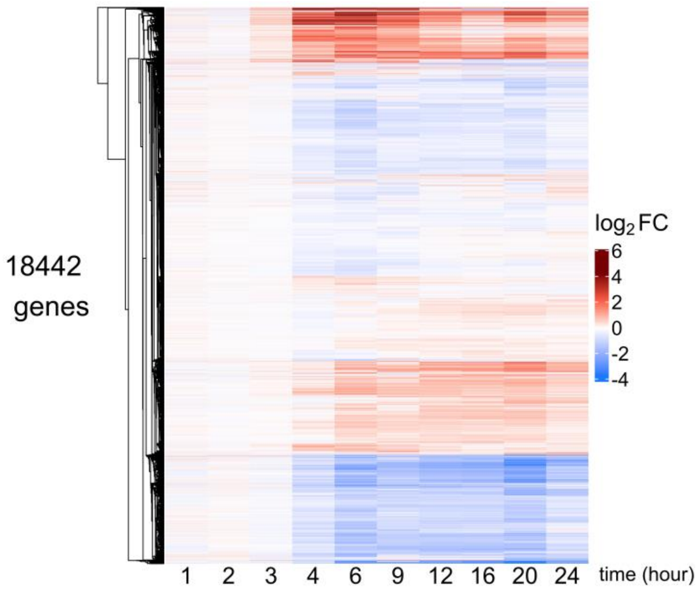


**B. The distribution of the signal-to-noise ratio across 3039 significantly upregulated genes.** The signal-to-noise ratio was defined as (maximum of $log_2$FC across the time points) / (mean of the SEs for the $log_2$FC across the time points). The vertical red dashed line shows signal-to-noise-ratio = 6.5. We selected 2435 genes that had a signal-to-noise ratio higher than 6.5 as the high-precision upregulated genes.

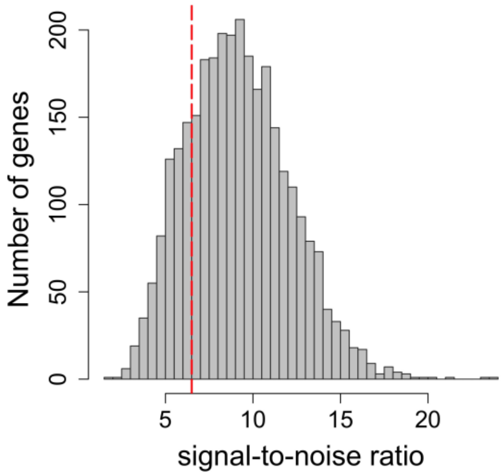

**C.** A heatmap of 2435 high-precision upregulated genes. Note that the absolute values at 3 hpi are much lower compared to the peak values for the genes.

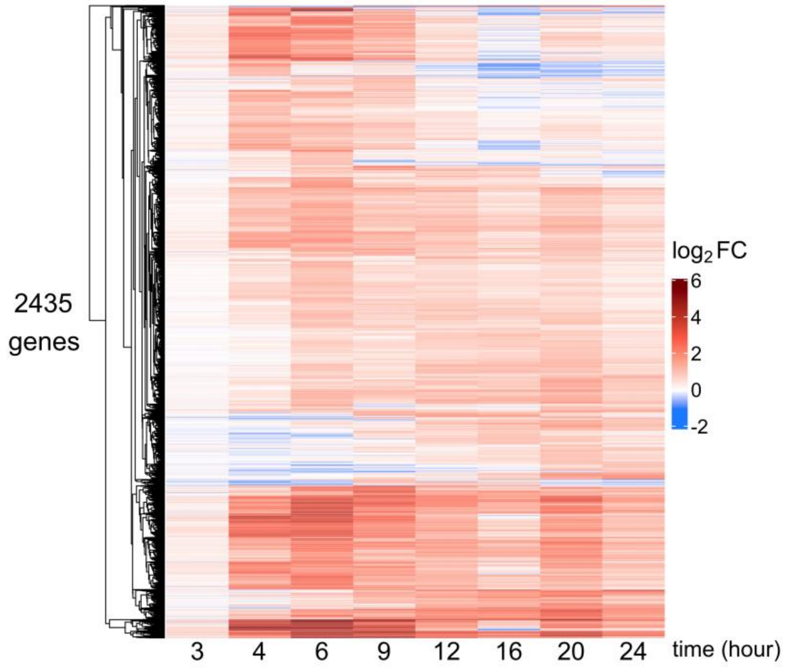

**Fig. S2. Spline models of 2435 high-precision upregulated genes. A.** The spline model (curve) and the normalized read count data for the $\log_2$-scale (circle points) are shown for each indicated gene. This large figure is provided in a separate file.

**B.** Double-peak decomposition based on the spline models (Spline) is compared to double-peak decomposition by MCM with Set 1 or Set 2 (MCM (Set1) and MCM (Set2)). The decomposition results across 1366 Echoing genes (Fig. 4) are shown. The gene order is the same as in Fig. 4. The decomposed single-peak patterns were generally similar between MCM and spline models, indicating that the double-peak patterns were generally not artifacts of MCM fitting.

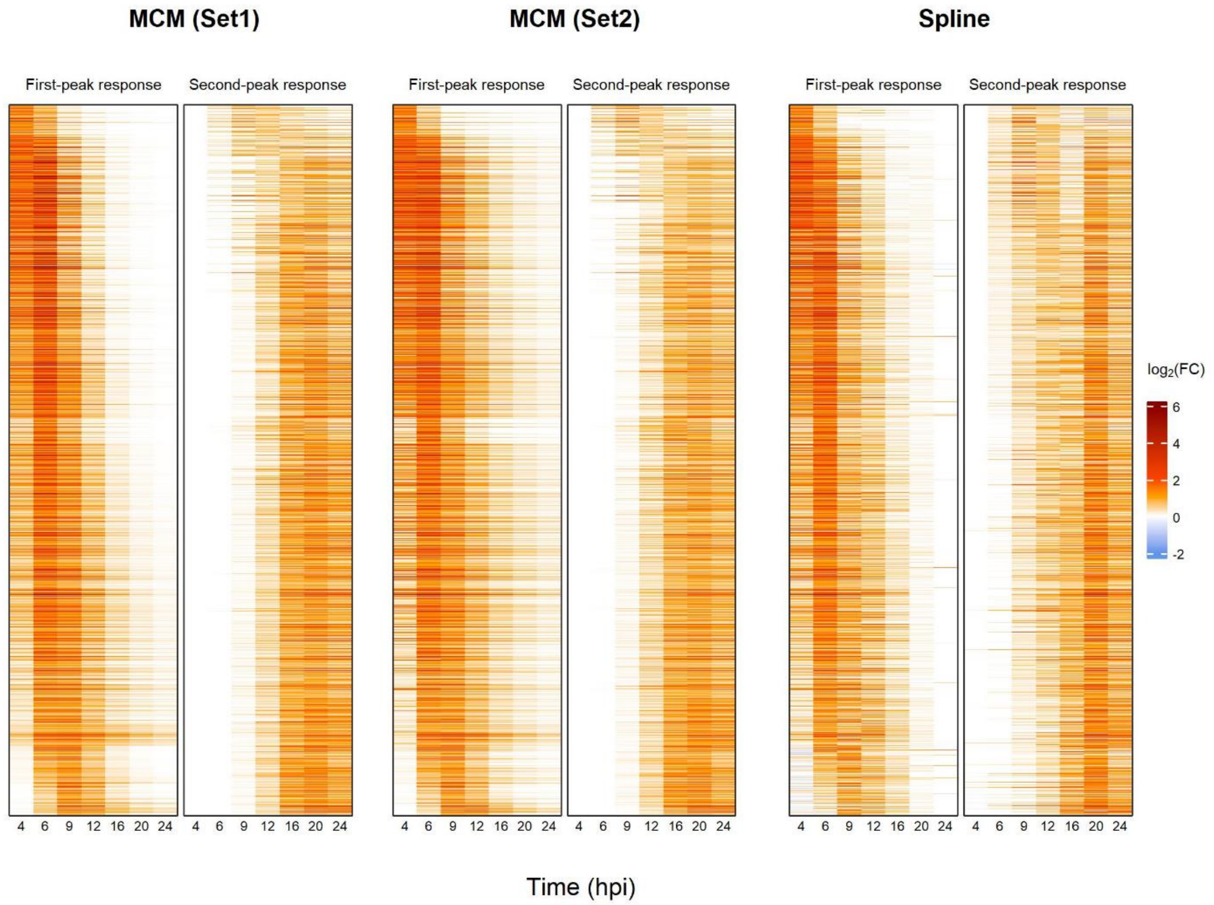

**Fig. S3. Altered models were fit for 191 genes.** These genes have time-course patterns with a single peak and a shoulder because of early second peaks. To capture this trend, the MCM was fit to the read count data up to 16 hpi (altered MCM). For each gene, the original MCM (black) and the altered MCM (red) curves are shown. The circle points show the normalized read count data for the $\log_2$-scale. These genes tend to have third peaks, which were not captured by the altered models. Some of the genes had relatively early third peaks, which already started rising at 16 hpi, and the fit of the altered MCM for these genes was not ideal. However, the number of such genes was low, and we did not try to fit another special model for the genes. For seven genes, only the altered MCM curves (red) are visible because the altered MCMs were almost identical to the original MCMs. This large figure is provided in a separate file.

**Fig. S4. Selection of 1889 genes that were stably modeled with both signaling compartment decay rate parameter value Set 1 and Set 2.**

The distribution of the Pearson correlation coefficients between the MCMs fitted using Set 1 and Set 2 of the signaling compartment decay rate parameter values across 2435 high-precision upregulated genes is shown. Among them 1889 genes had a Pearson correlation higher than 0.9. We designated 1889 genes as the modeled upregulated genes and used their MCMs for the analysis of the model results. For each gene, the outputs of which two signaling compartments were used, the input amplification ratio parameter values for two response compartments, and the decay rate parameter values of the response compartments are given in Table S1 for Set1 and Set 2.

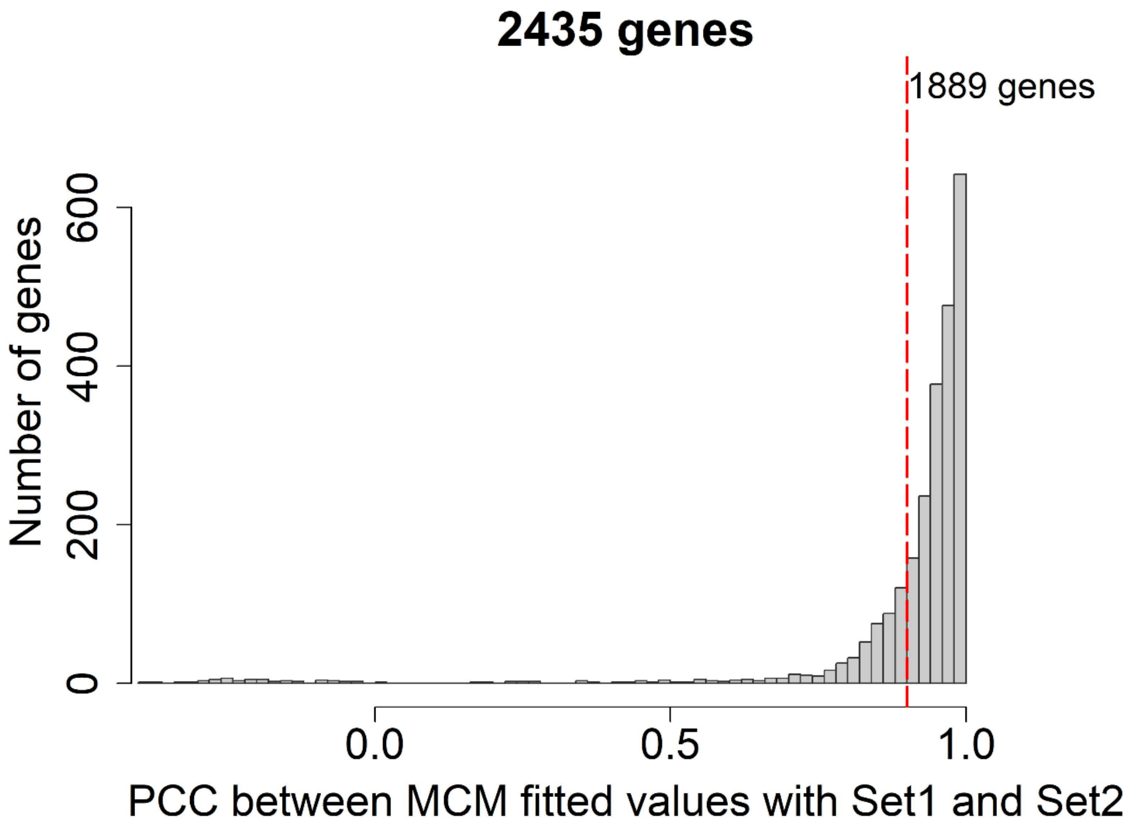

**Fig. S5. MCM with decompositions and data points for 1889 modeled upregulated genes.** Set 1 signaling compartment decay rate parameter values were used. Altered models (Fig. S3) were fit for 176 genes with green asterisks after the gene names and 15 genes with turquoise squares with "x" after the gene names. The MCM, the decomposed first peak, and the decomposed second peak curves are shown in red, blue, and orange, respectively. Points show actual observations of the between-libraries-normalized $\log_2$ mRNA level ratio of *Pto* AvrRpt2 over mock. The color of each gene name shows the classification of the gene according to the decomposition patterns: ACP-specific genes (class 5, red), echoing genes (class 6, orange), NACP-specific genes (classes 1, 7, and 8, with turquoise, blue, and cyan, respectively), and unclassified genes (classes 2, 3, and 4, black). See Fig. S7 for details of gene classification. This large figure is provided in a separate file.

For the MCM for each gene, the outputs of which two signaling compartments were used as the inputs to the response compartment, the input amplification rate parameter values for two response compartments, and the decay rate parameter values of the response compartments are shown in Table S1.

**Fig. S6. The distribution of the Pearson correlations between the mean estimates from the data and MCM-modeled values across 1889 modeled upregulated genes.** The values were compared at 4, 6, 9, 12, 16, 20, and 24 hpi for most genes and at 4, 6, 9, 12, and 16 hpi for 104 altered model genes. The median of the Pearson correlation was 0.93 as shown by a vertical red dashed line.

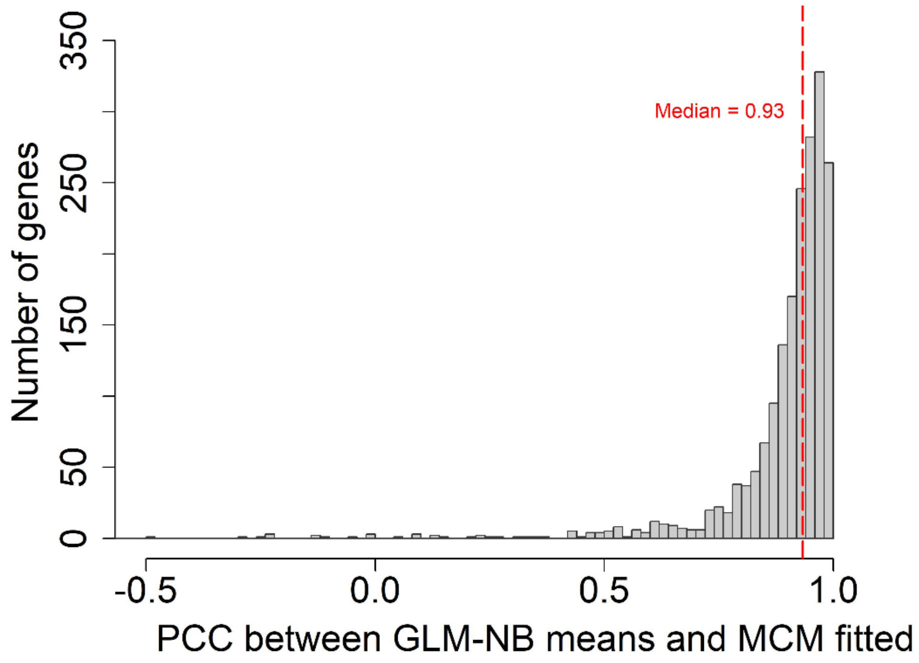

**Fig. S7. Explanations and examples of gene classification**

This explains how genes were classified into four categories of I. ACP-specific genes, II. Echoing genes, III. NACP-specific genes, and IV. Unclassified genes. First, genes were grouped into 8 classes based on (first peak time), (first peak level), and (second peak level) estimated by the MCM. For each of the eight classes, three example gene models are shown.

**I. ACP-specific genes (214 genes)**

Class 5. 214 genes. (first peak time) ≤ 10 hpi AND

0 < (second peak level) * 4 < (first peak level) AND (first peak level) ≥ 0.5

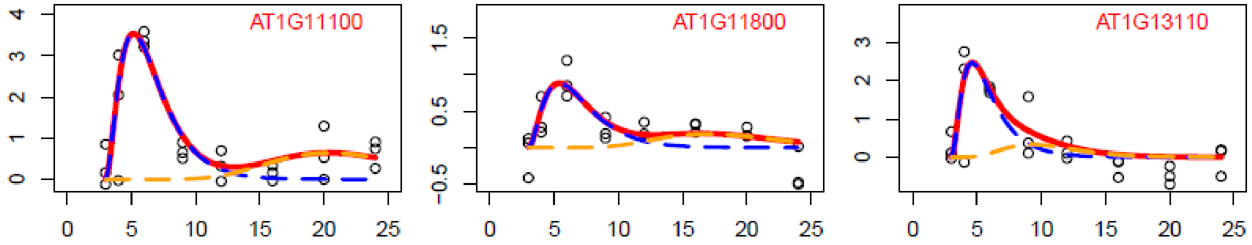


**II. Echoing genes (1366 genes)**

Class 6. 1366 genes. (first peak time) ≤ 10 hpi AND

0 < (second peak level) * 0.5 < (first peak level) ≤ (second peak level) * 4 AND

(first peak level) ≥ 0.5

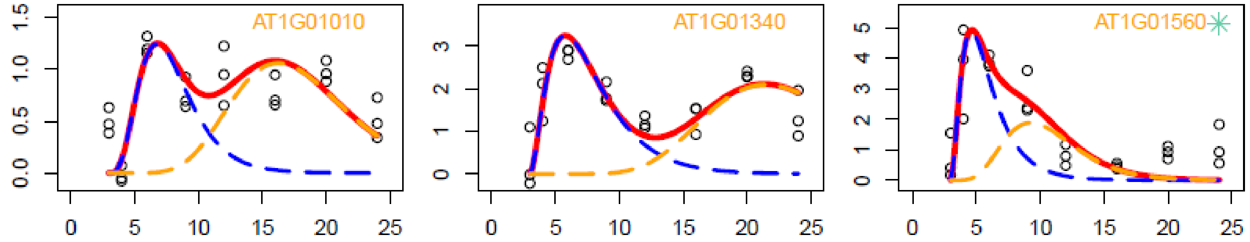


Note that the model for AT1G13210 (with a green dot next to the gene name) was fit to the data up to 16 hpi, and thus the model does not capture the third peak trend after 16 hpi.

**III. NACP-specific genes (227 genes)**

Class 1. 38 genes. (first peak level) ≤ 0 AND |(first peak level)| * 4 ≤ (second peak level)

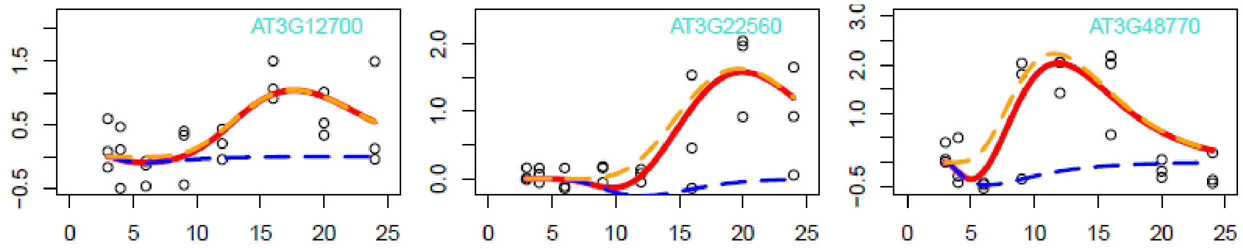


Class 7. 137 genes. (first peak time) ≤ 10 hpi AND

0 < (first peak level) ≤ (second peak level) * 0.5 AND (second peak level) ≥ 0.5

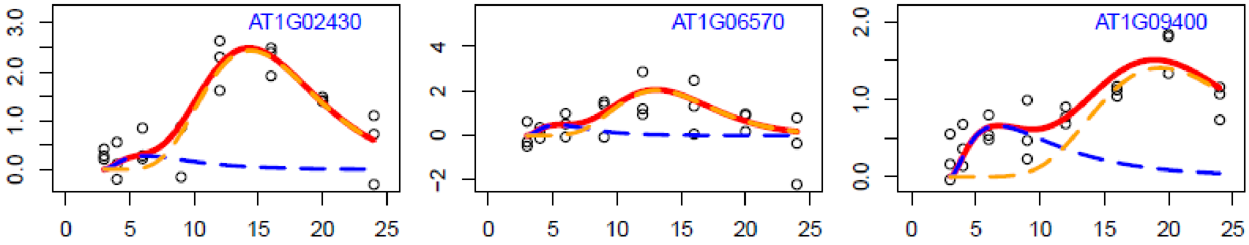


Class 8. 52 genes. (first peak time) > 10 hpi AND
{ (first peak level) ≥ 0.5 OR (second peak level) ≥ 0.5 } AND (first peak level) > 0

Since ACP cells are mostly dead by 9 hpi, occurrence of the first peak later than 10 hpi indicates that there was no detectable ACP response. Therefore, these genes were classified as NACP-specific genes.

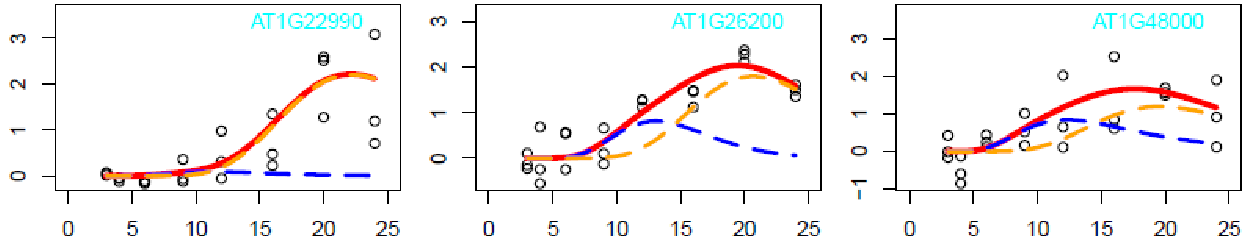


### IV. Unclassified genes (82 genes)

Class 2. 79 genes. (first peak level) ≤ 0 AND 0 < (second peak level) ≤ |(first peak level)| ∗ 4

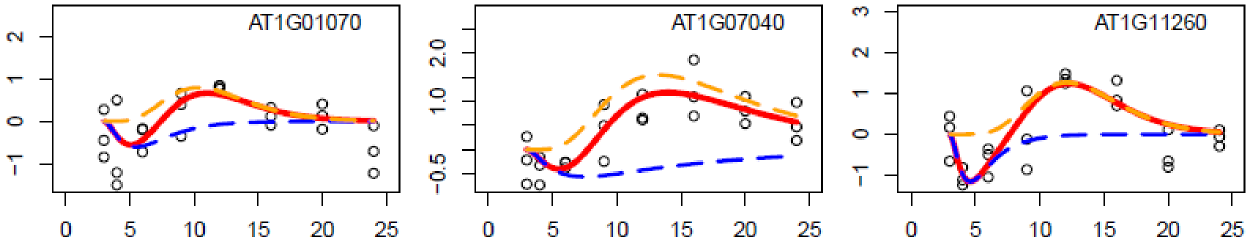


Class 3. 3 genes. (first peak level) ≤ 0 AND (second peak level) ≤ 0

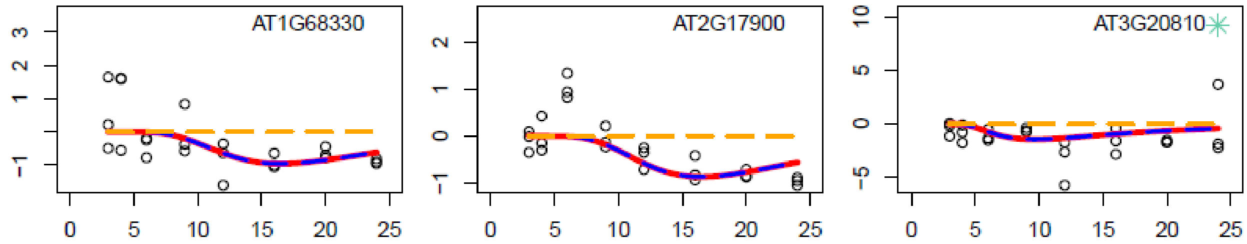


Class 4. 0 genes. 0 < (first peak level) < 0.5 AND (second peak level) < 0.5

**Fig. S8. Time-scaling relationships between the first- and second-peak responses**

A. The distribution of the ratio of peak times, second peak (NACP) / first peak (ACP), across 1366 Echoing genes.

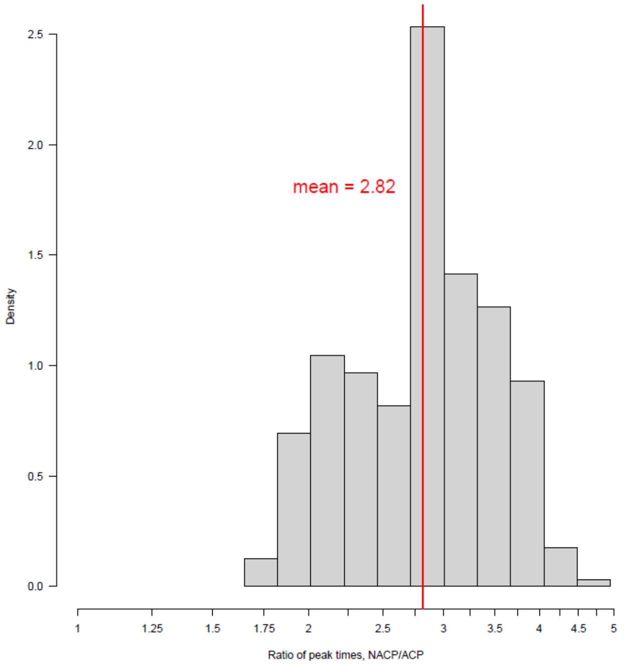


B. The distribution of the ratio of peak width at 75% peak level, second peak (NACP) / first peak (ACP), across 1366 Echoing genes.

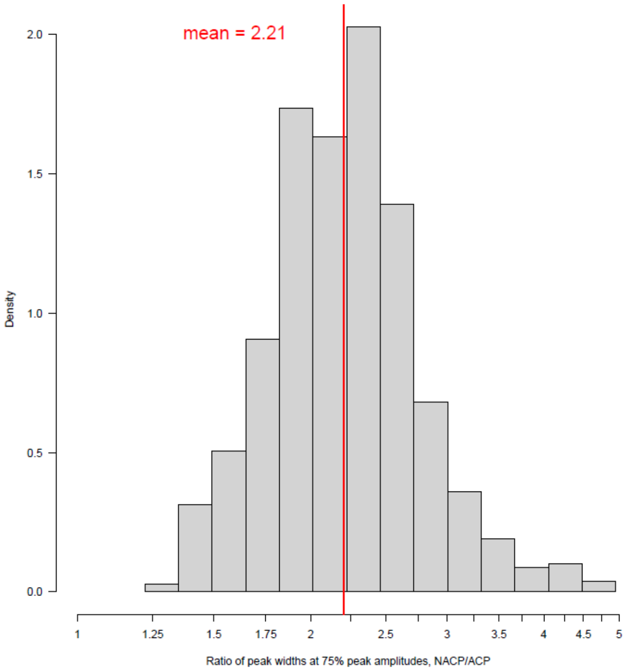

**Fig. S9. Heatmaps of genes with consolidated TF binding sites, organized along the peak times of the first- and second-peak responses.**

Members of each of the TF families, NAC1s, NAC2s, WRKYs, PEARs, and HSFs were consolidated. Otherwise, it is the same as Fig. 6A.

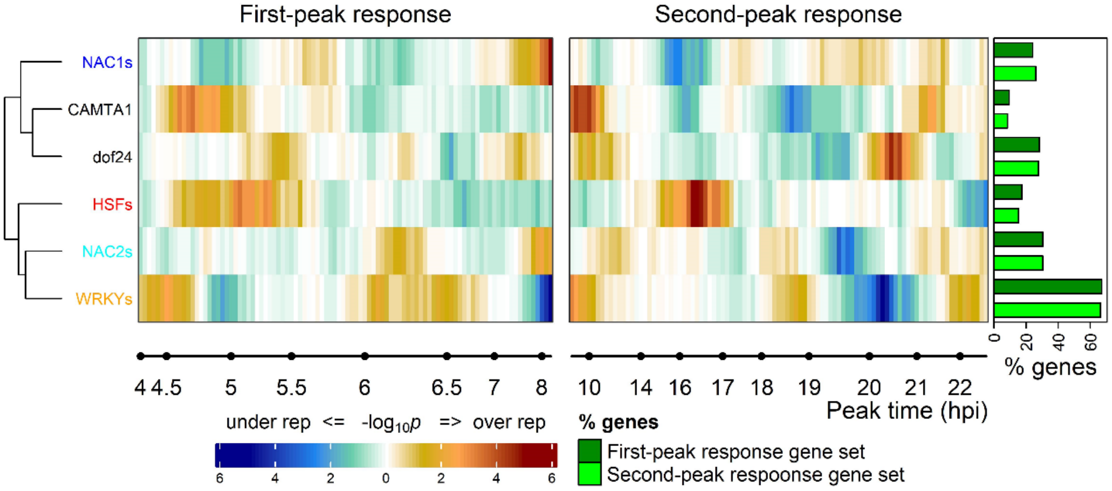

**Fig. S10. WRKY genes have binding sites for various TFs.**

The numbers of binding sites (BS No.) for 304 TFs which had at least one binding site in the 35 *WRKY* genes in Fig. 7, are shown in a heatmap for 35 *WRKY* genes. The order and the font colors of 35 *WRKY* genes are the same as those in Fig. 7. Among 304 TFs, WRKYs are indicated by red bars on the bottom of the heatmap.

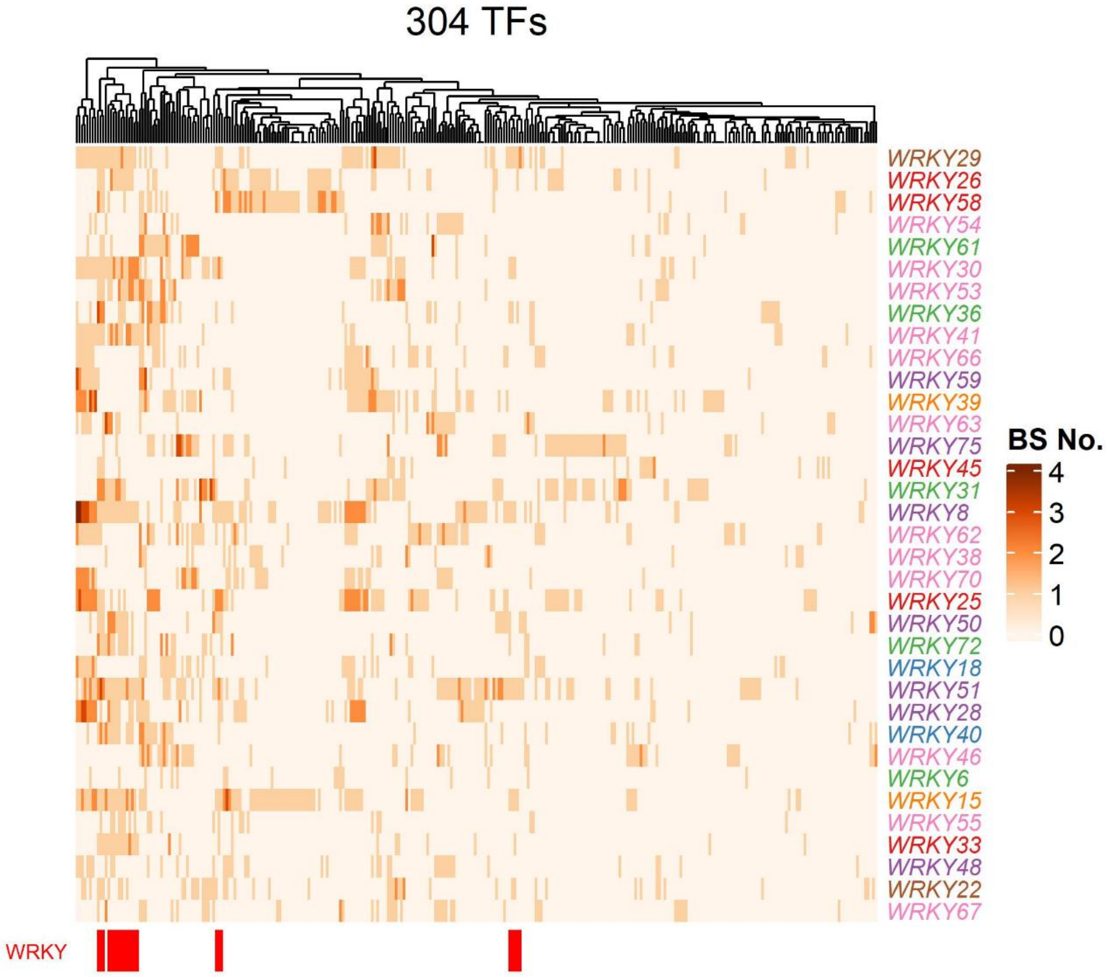

**Fig. S11. Core immune transcriptome response and responses specific to different elicitor types.**

The Venn diagram shows the number of genes specific and shared among 1684 "ACP_up" genes (ACP-specific and Echoing genes), 1658 "NACP_up" genes (Echoing and NACP-specific genes), and 2572 "flg22_up" genes. The flg22_up genes were selected for *Storey's q* $< 0.05$, $\log_2$ {(WT flg22-treated) / (*fls2* flg22 treated)} $> 1$ at at least two consecutive time points, using the GLM-NB models (Hillmer *et al.*, 2017).

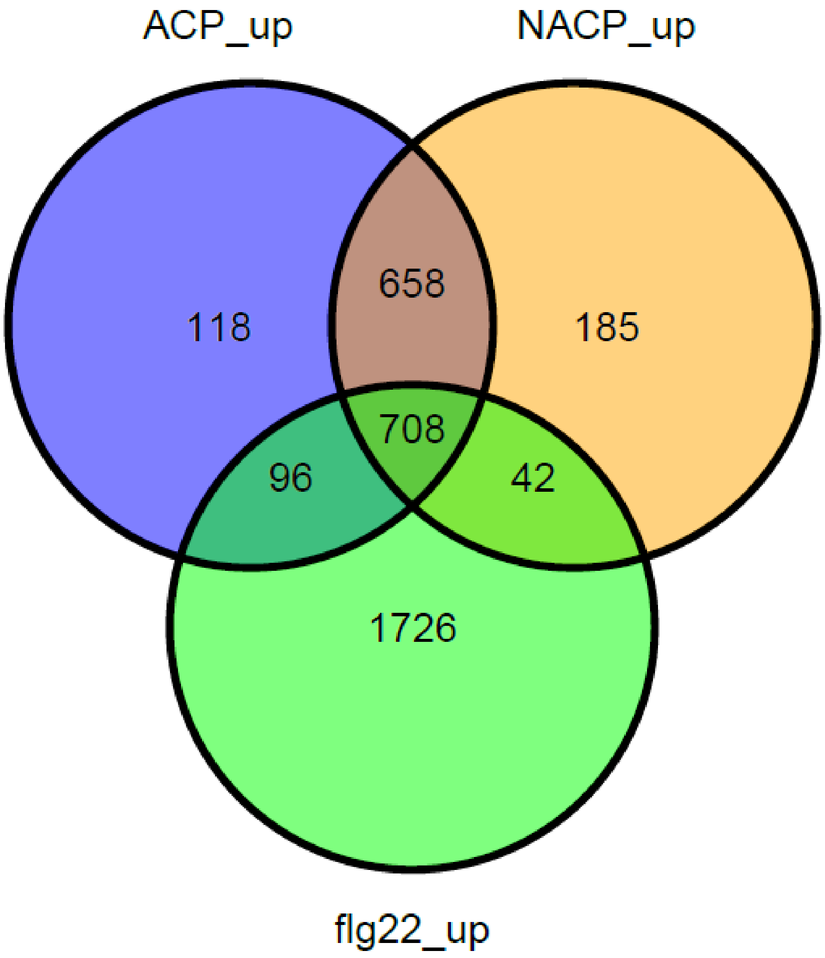
