## Supplementary figures and images for "Dynamic decomposition of transcriptome responses during plant effector-triggered immunity revealed conserved responses in two distinct cell populations"

### Fig. S3

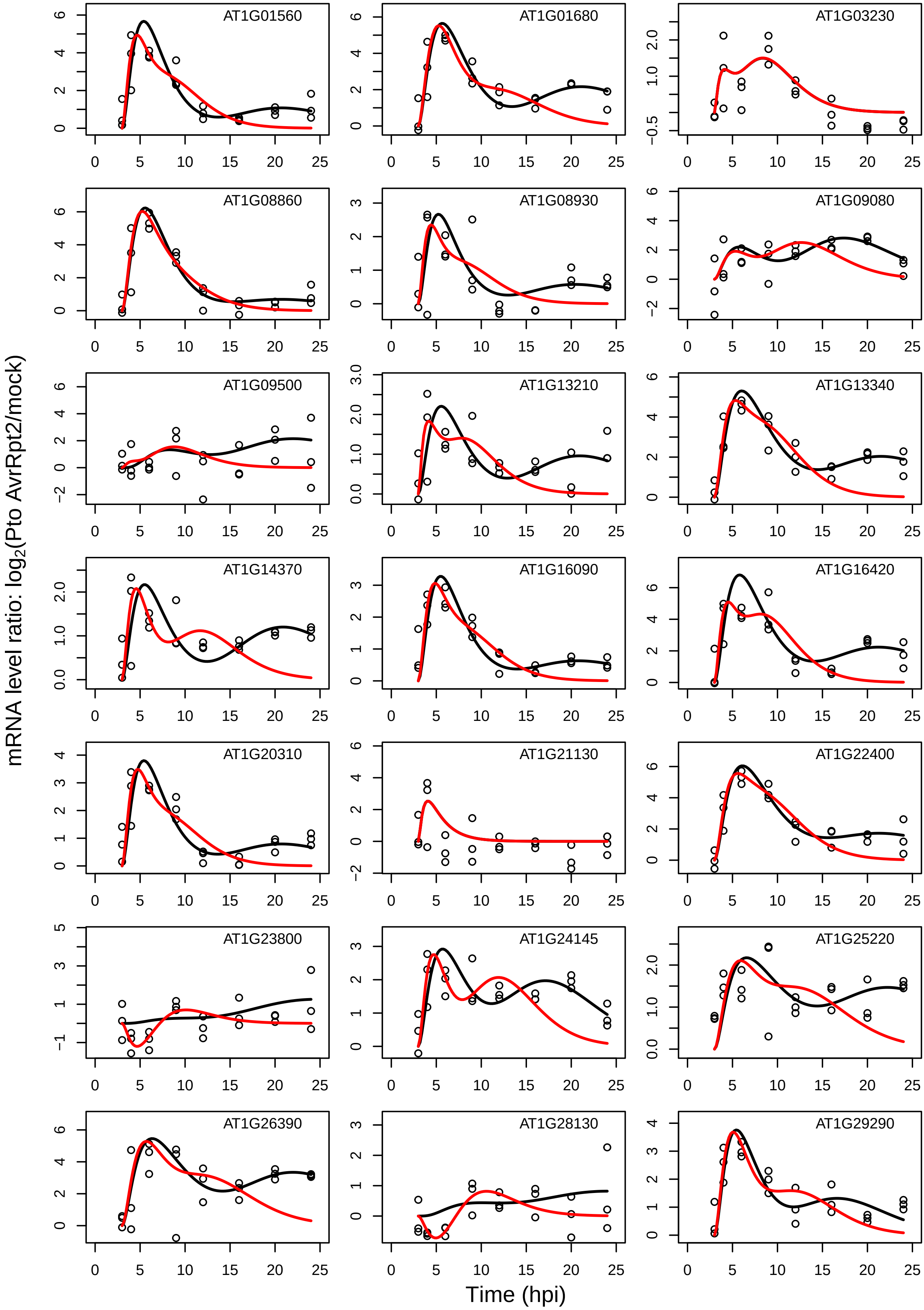

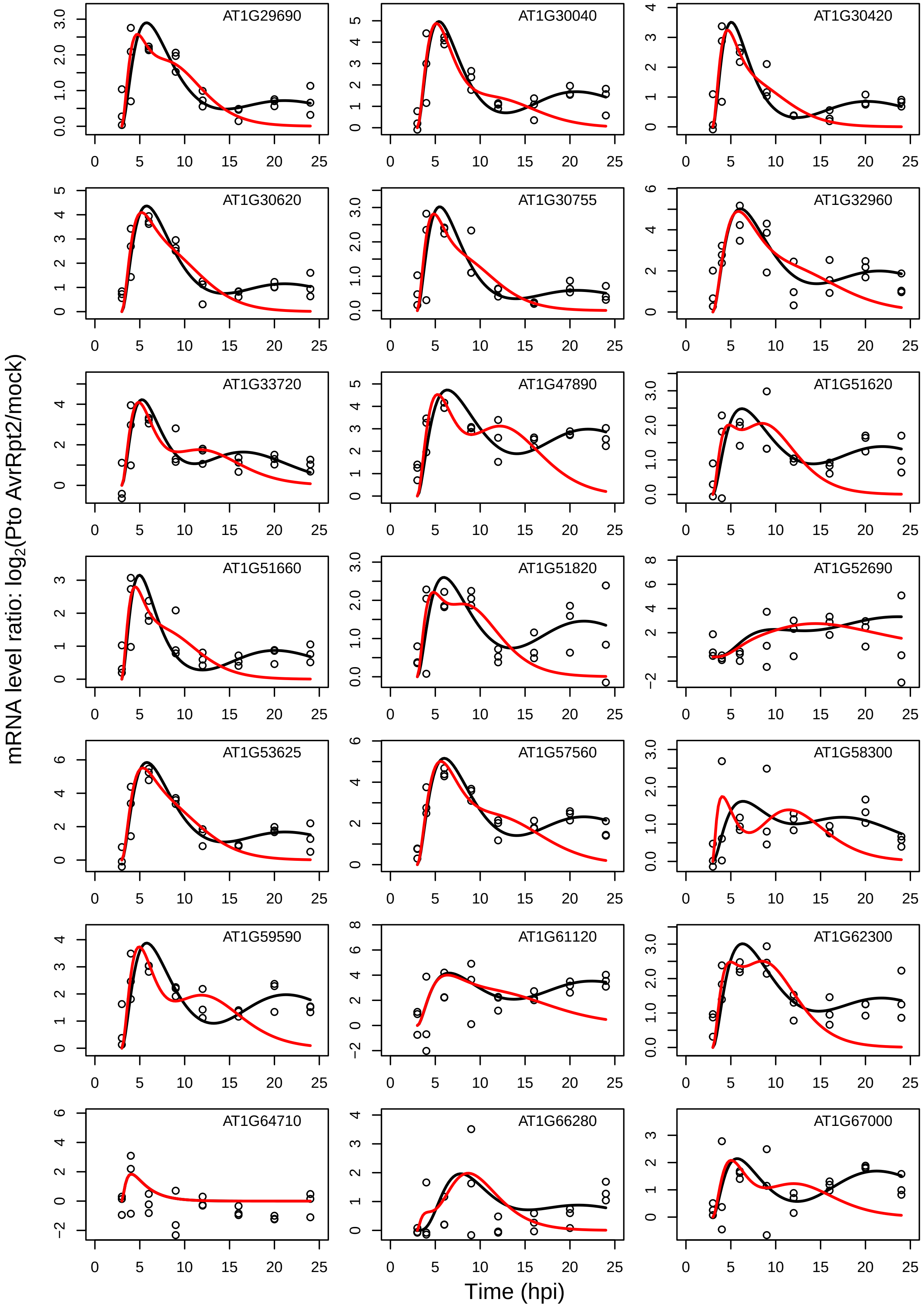

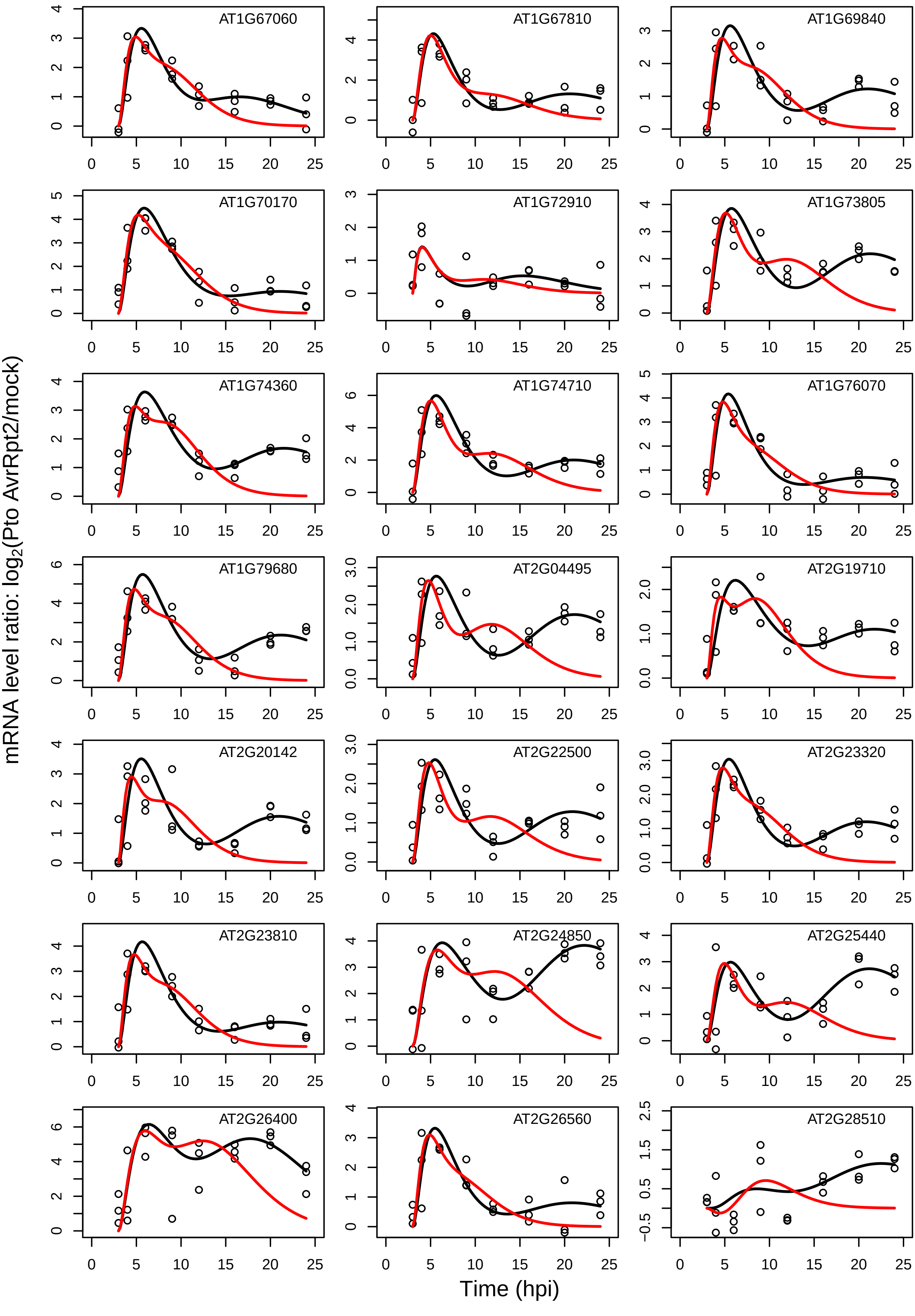

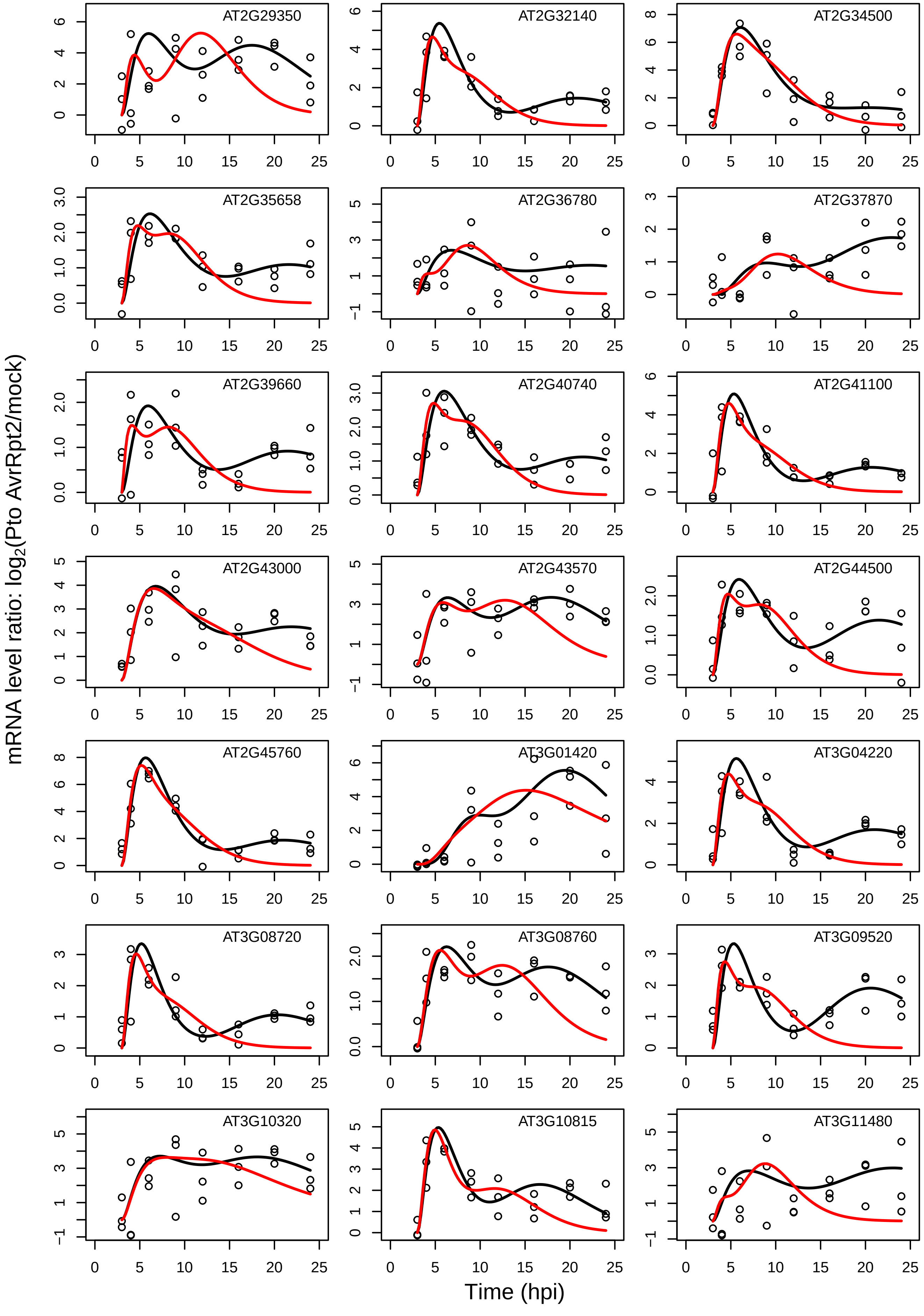

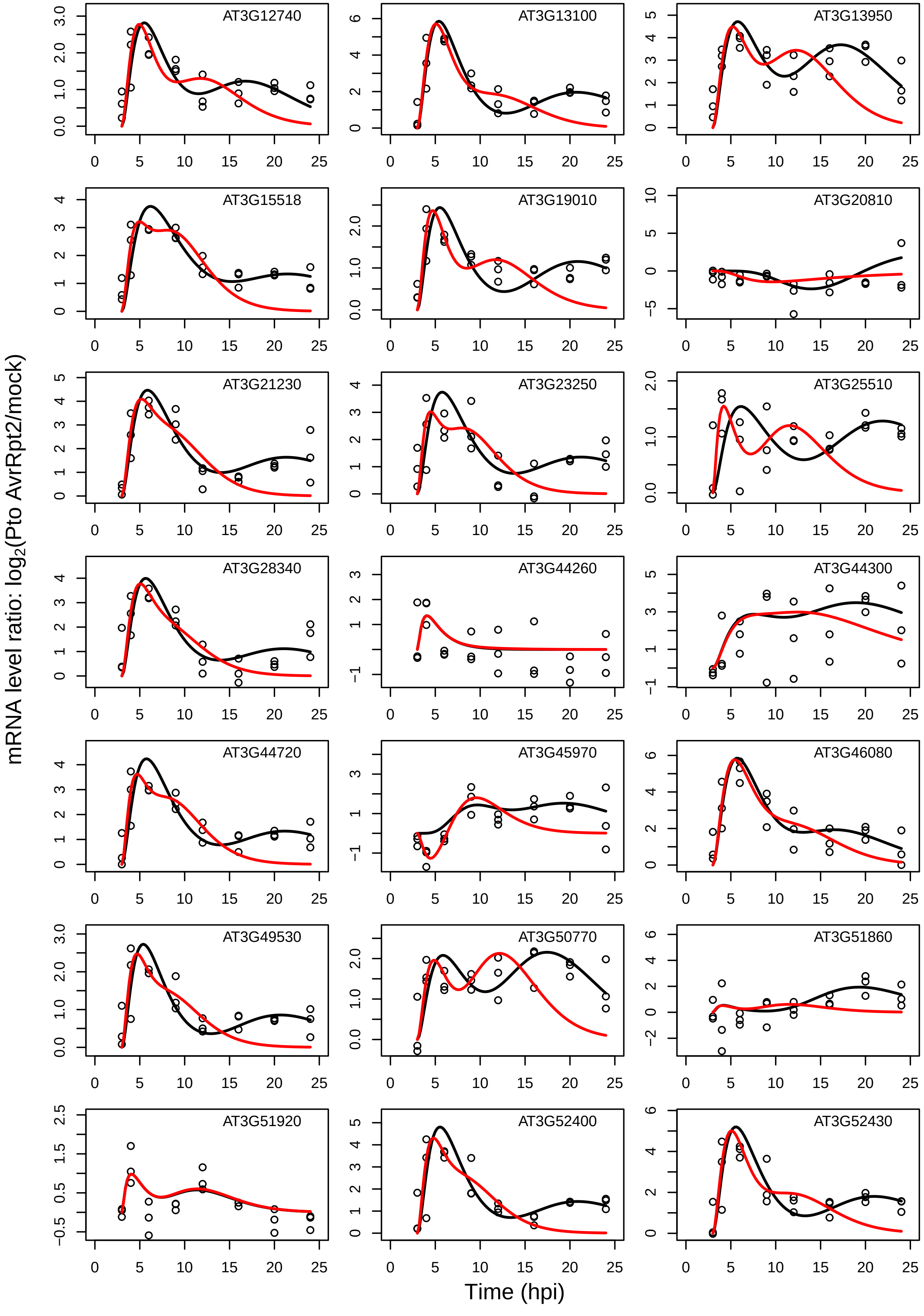

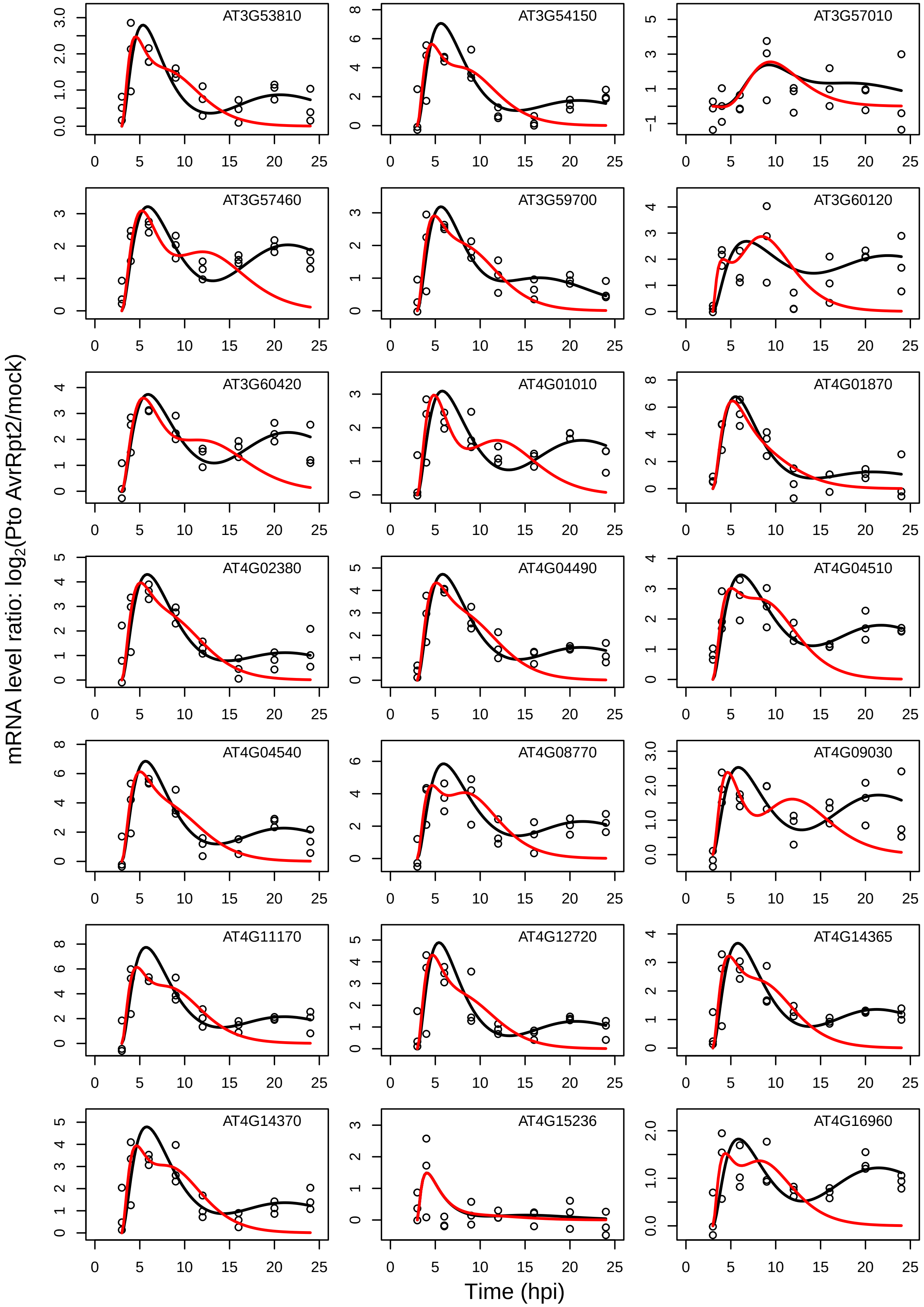

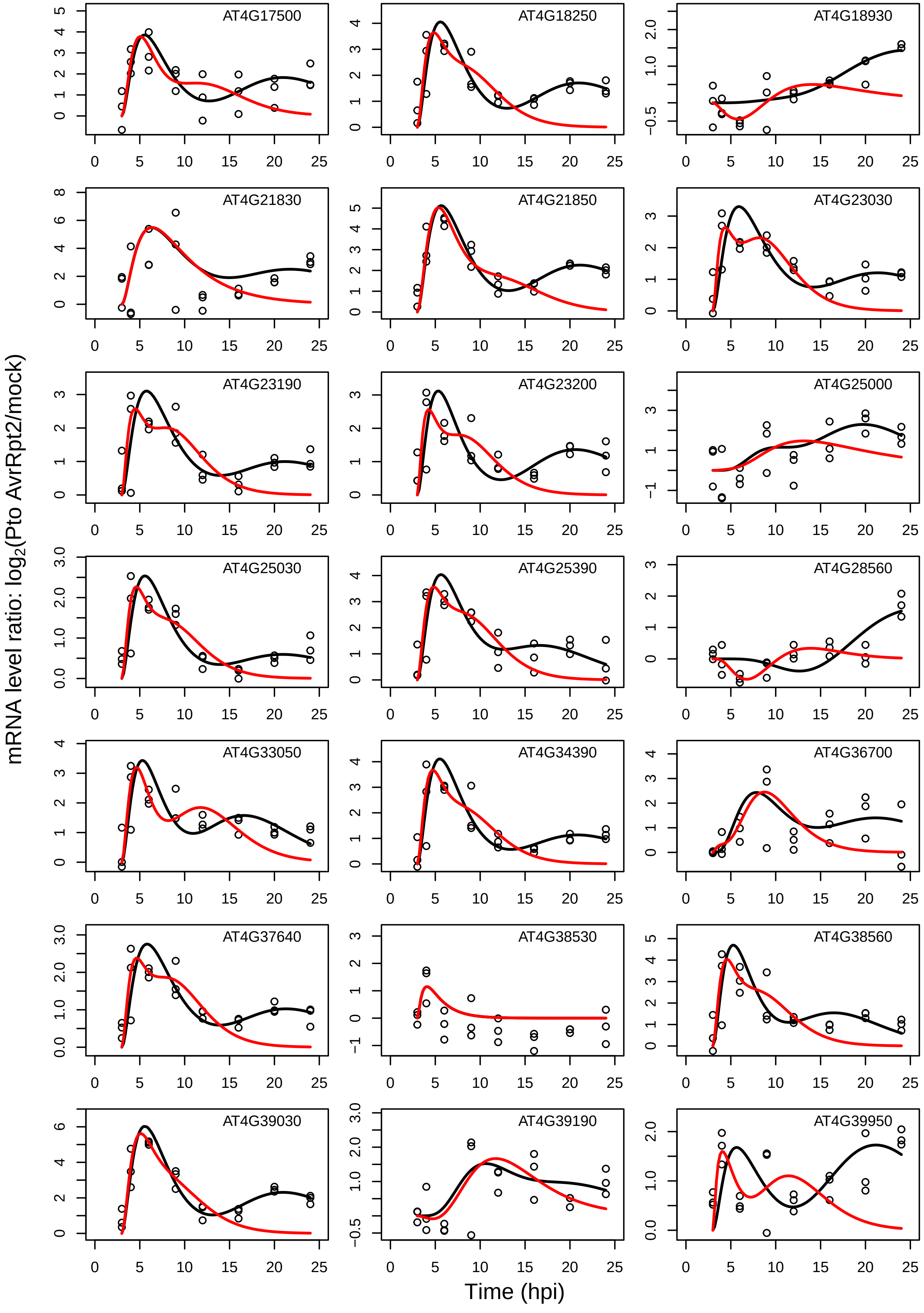

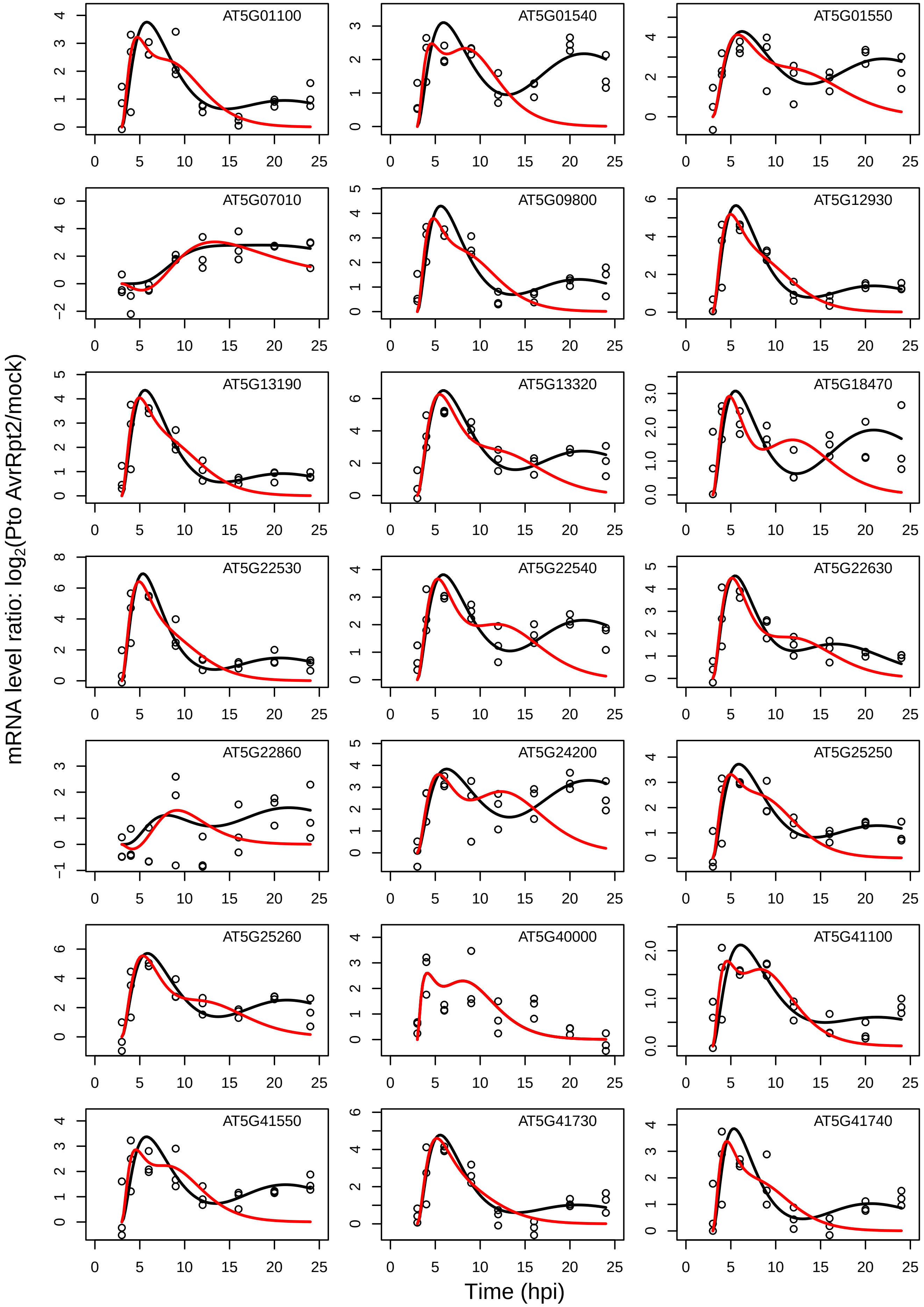

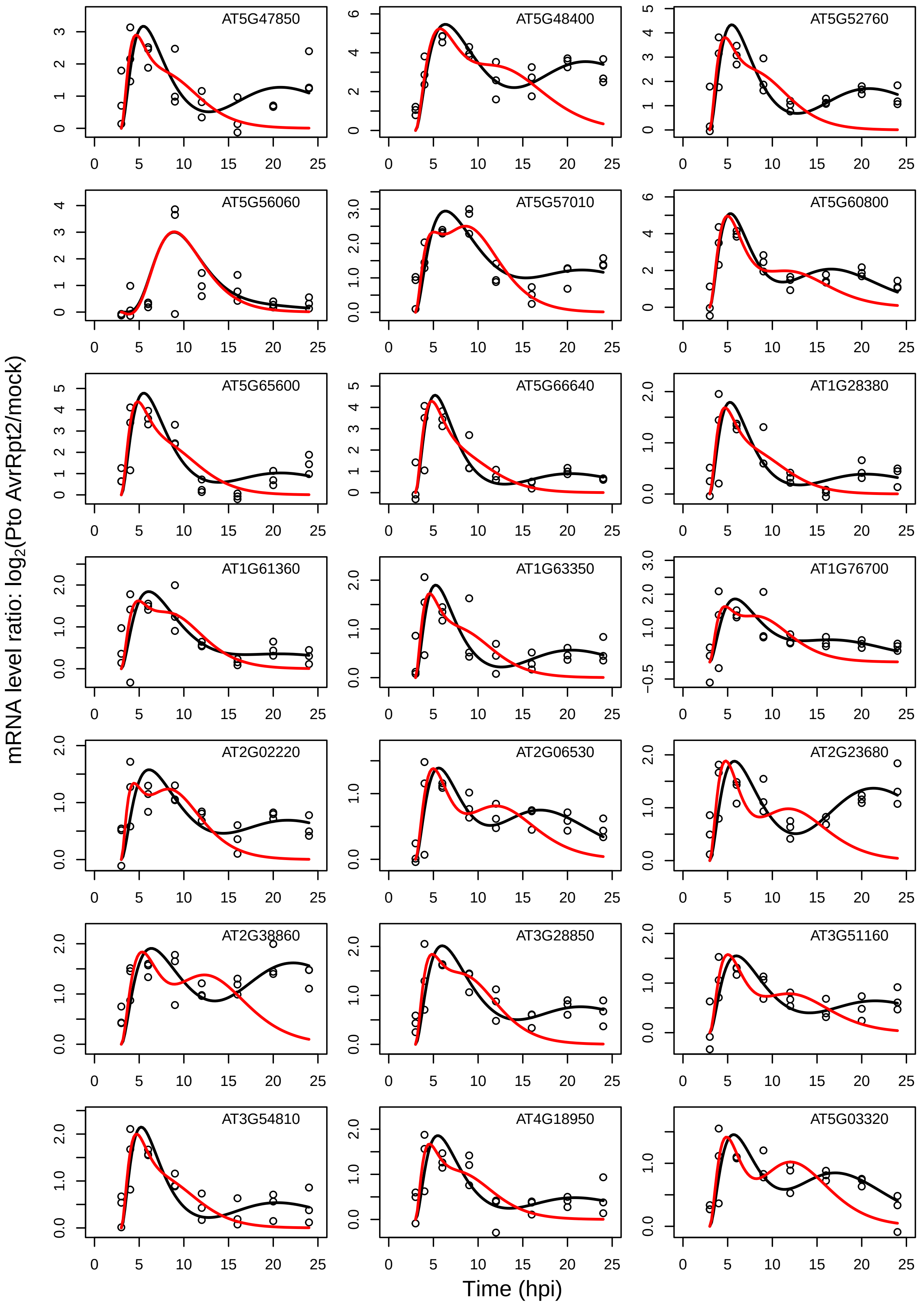

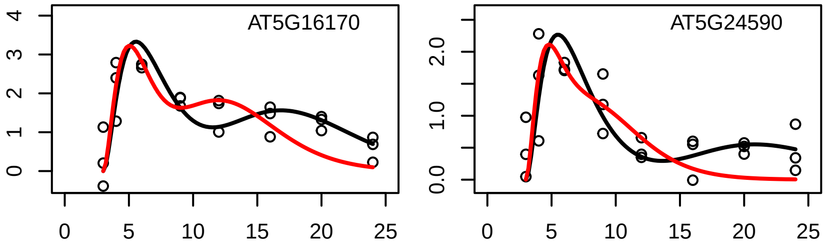
