## Supplementary material for "Dynamic decomposition of transcriptome responses during plant effector-triggered immunity revealed conserved responses in two distinct cell populations": Fig. S5

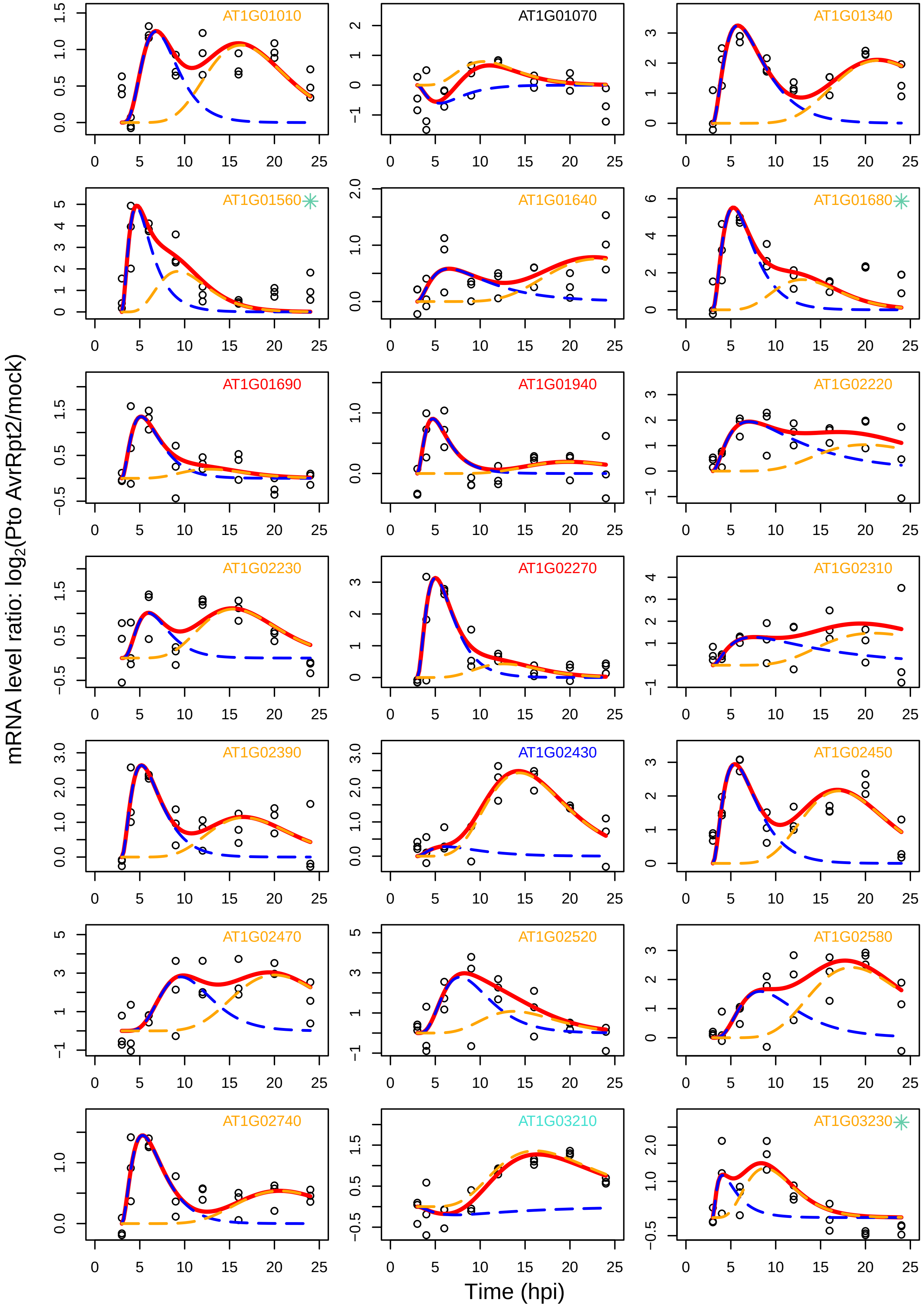
AT1G01010
AT1G01070
AT1G01340
AT1G01560
AT1G01640
AT1G01680
AT1G01690
AT1G01940
AT1G02220
AT1G02230
AT1G02270
AT1G02310
AT1G02390
AT1G02430
AT1G02450
AT1G02470
AT1G02520
AT1G02580
AT1G02740
AT1G03210
AT1G03230
mRNA level ratio: log2(Pto AvrRpt2/mock)
Time (hpi)

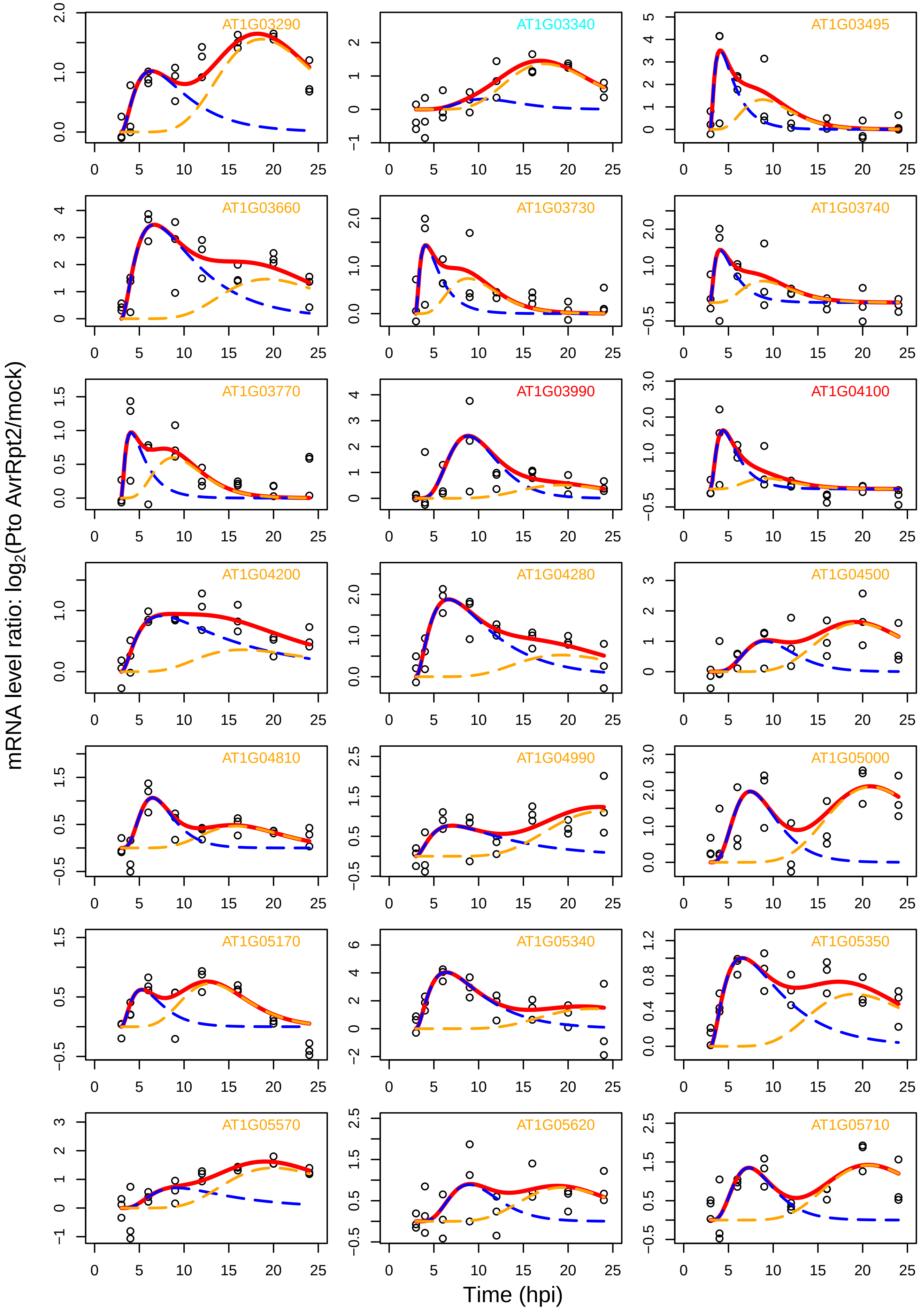
mRNA level ratio: log2(Pto AvrRpt2/mock)
Time (hpi)
AT1G03290
AT1G03340
AT1G03495
AT1G03660
AT1G03730
AT1G03740
AT1G03770
AT1G03990
AT1G04100
AT1G04200
AT1G04280
AT1G04500
AT1G04810
AT1G04990
AT1G05000
AT1G05170
AT1G05340
AT1G05350
AT1G05570
AT1G05620
AT1G05710

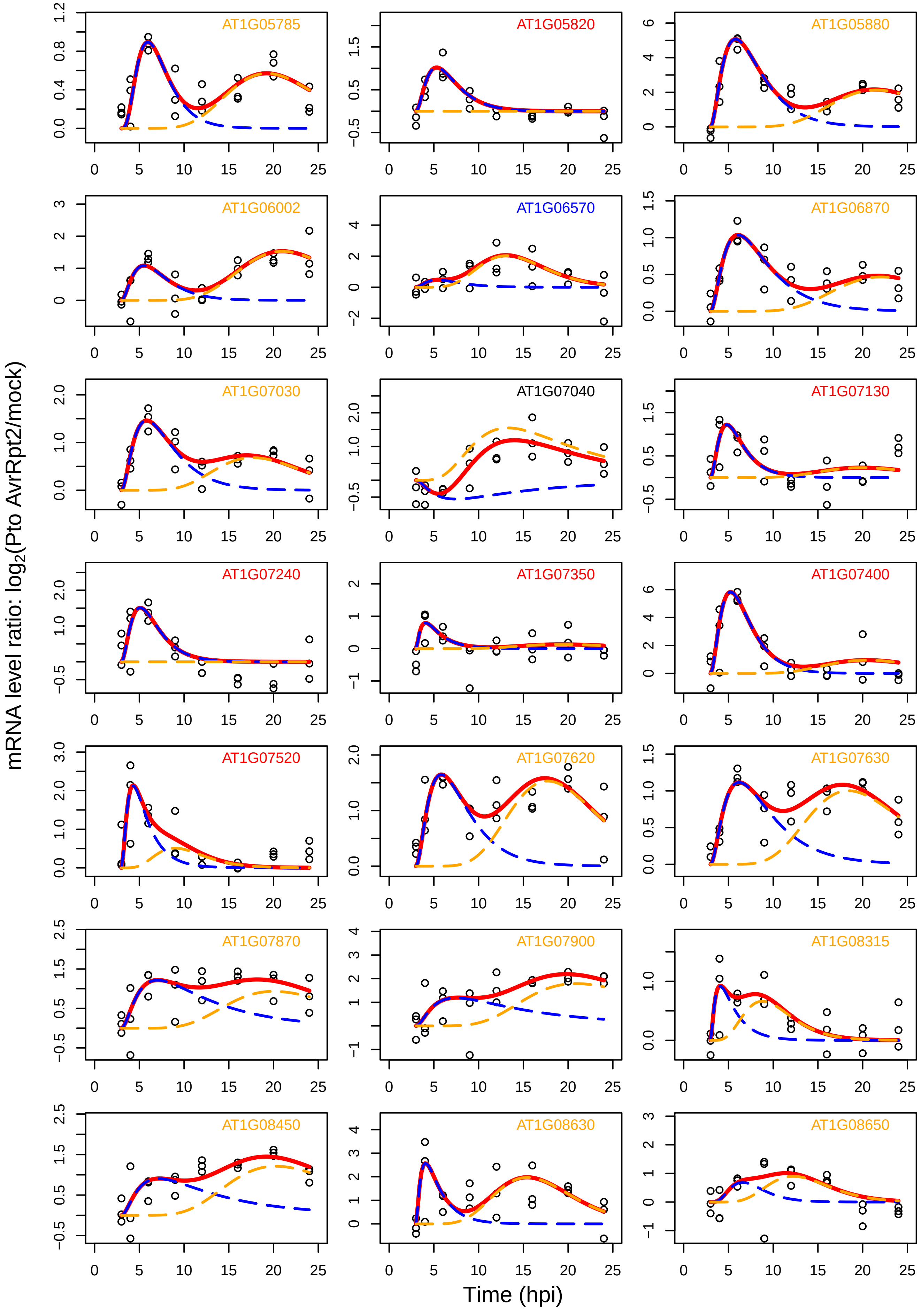

mRNA level ratio: log2(Pto AvrRpt2/mock)
Time (hpi)
AT1G05785
AT1G05820
AT1G05880
AT1G06002
AT1G06570
AT1G06870
AT1G07030
AT1G07040
AT1G07130
AT1G07240
AT1G07350
AT1G07400
AT1G07520
AT1G07620
AT1G07630
AT1G07870
AT1G07900
AT1G08315
AT1G08450
AT1G08630
AT1G08650

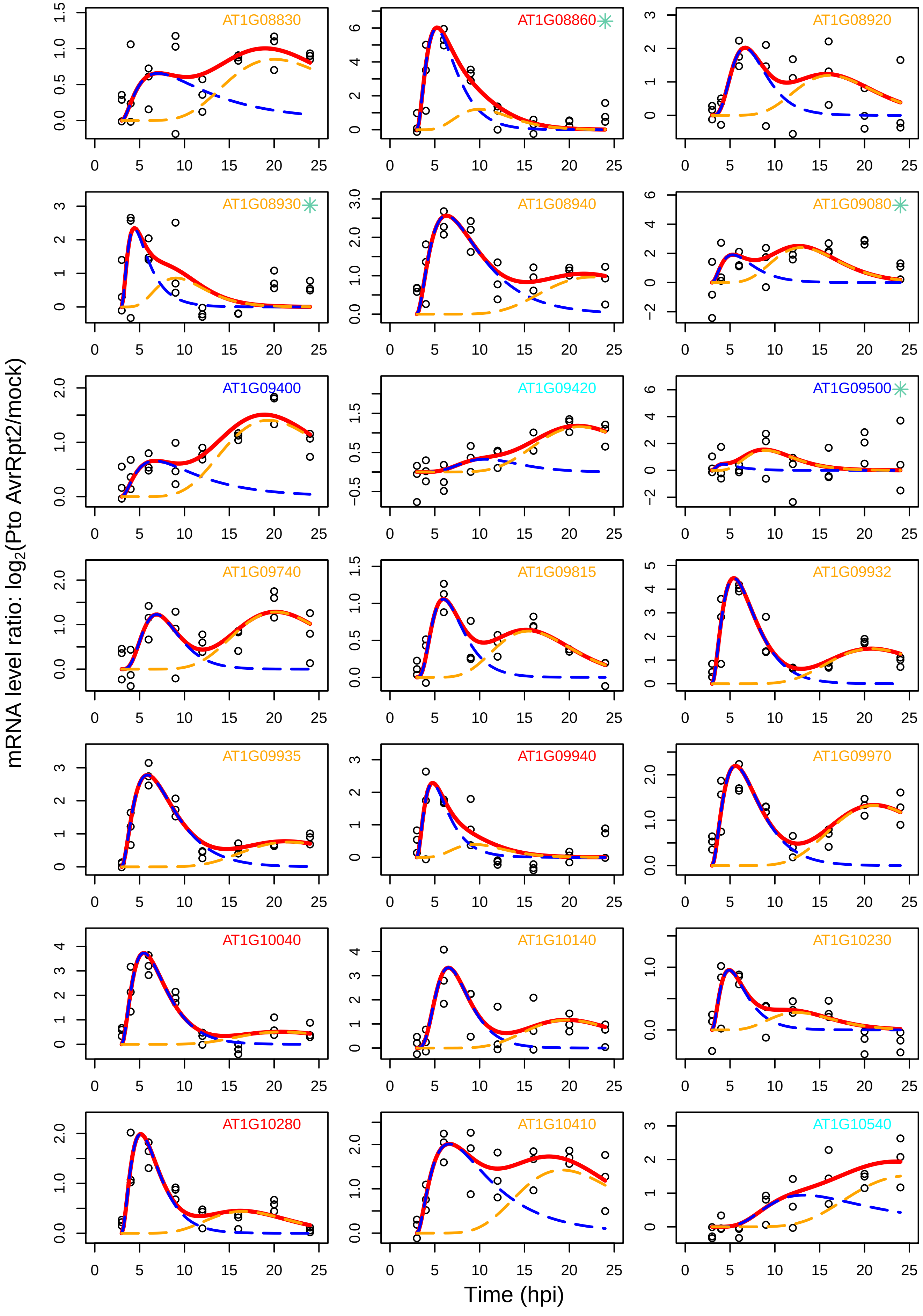

AT1G08830
AT1G08860
AT1G08920
AT1G08930
AT1G08940
AT1G09080
AT1G09400
AT1G09420
AT1G09500
AT1G09740
AT1G09815
AT1G09932
AT1G09935
AT1G09940
AT1G09970
AT1G10040
AT1G10140
AT1G10230
AT1G10280
AT1G10410
AT1G10540
mRNA level ratio: $\log_2$(Pto AvrRpt2/mock)
Time (hpi)

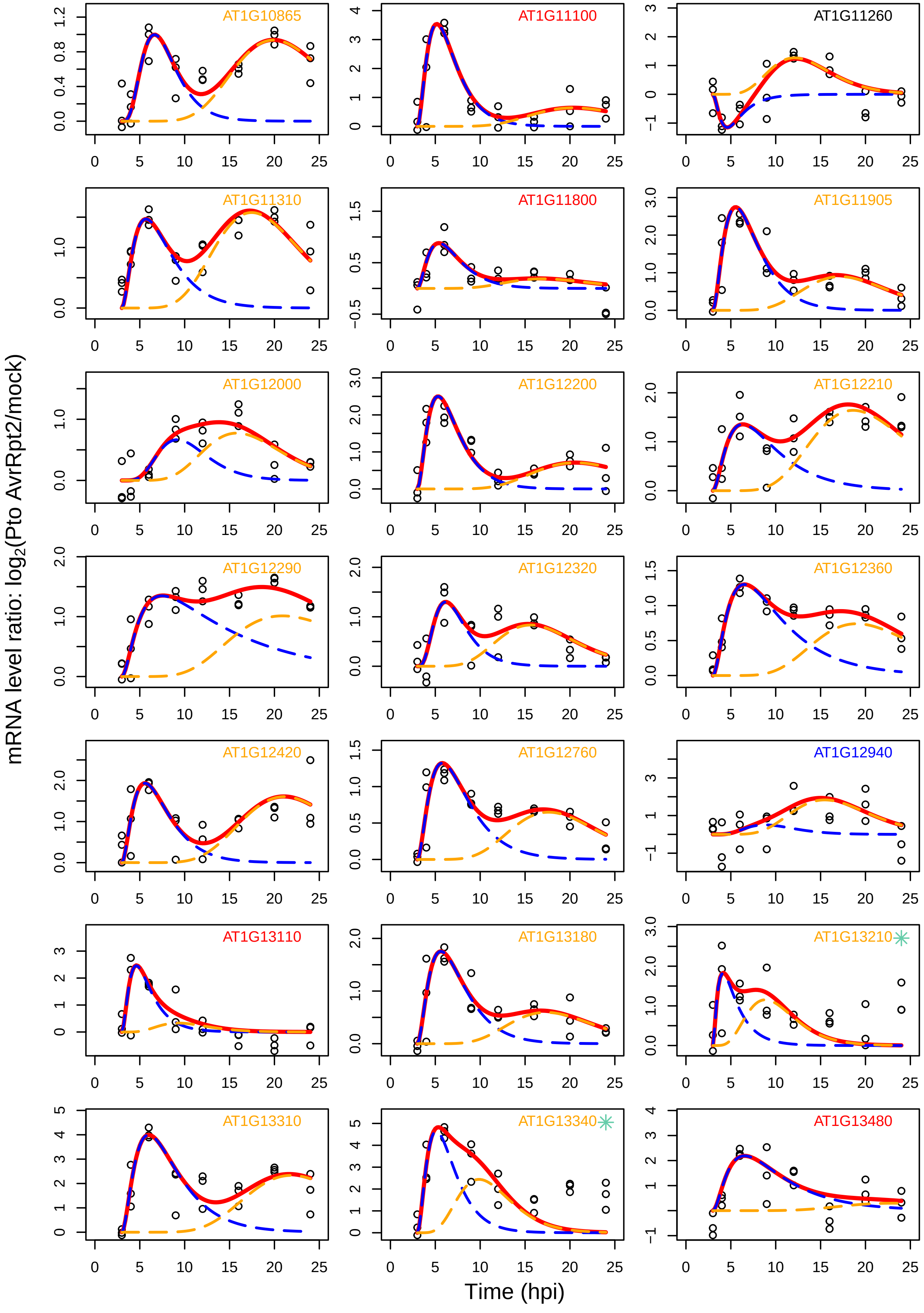
AT1G10865
AT1G11100
AT1G11260
AT1G11310
AT1G11800
AT1G11905
AT1G12000
AT1G12200
AT1G12210
AT1G12290
AT1G12320
AT1G12360
AT1G12420
AT1G12760
AT1G12940
AT1G13110
AT1G13180
AT1G13210
AT1G13310
AT1G13340
AT1G13480
mRNA level ratio: $log_2$(Pto AvrRpt2/mock)
Time (hpi)

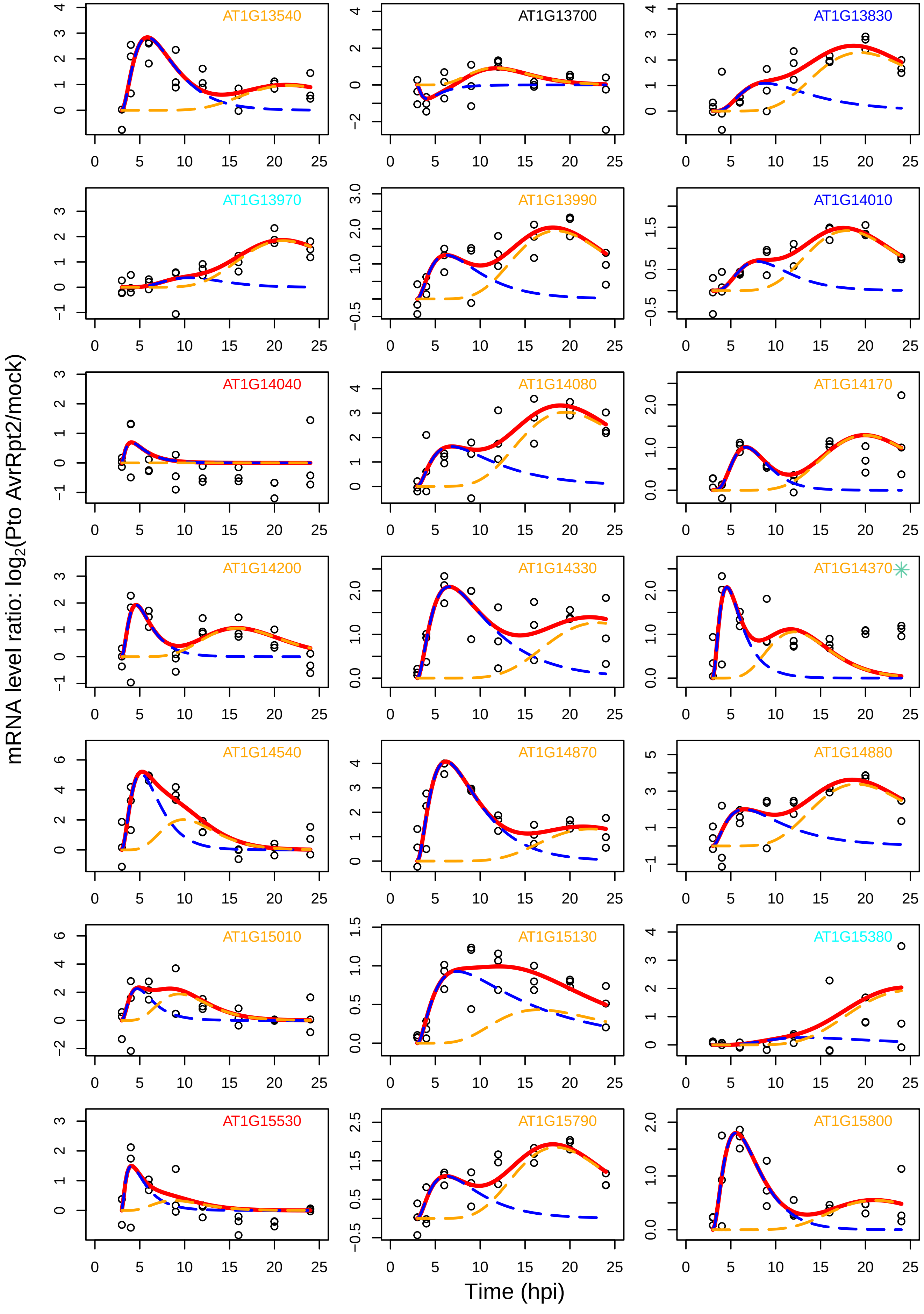
AT1G13540
AT1G13700
AT1G13830
AT1G13970
AT1G13990
AT1G14010
AT1G14040
AT1G14080
AT1G14170
AT1G14200
AT1G14330
AT1G14370
AT1G14540
AT1G14870
AT1G14880
AT1G15010
AT1G15130
AT1G15380
AT1G15530
AT1G15790
AT1G15800
mRNA level ratio: log2(Pto AvrRpt2/mock)
Time (hpi)

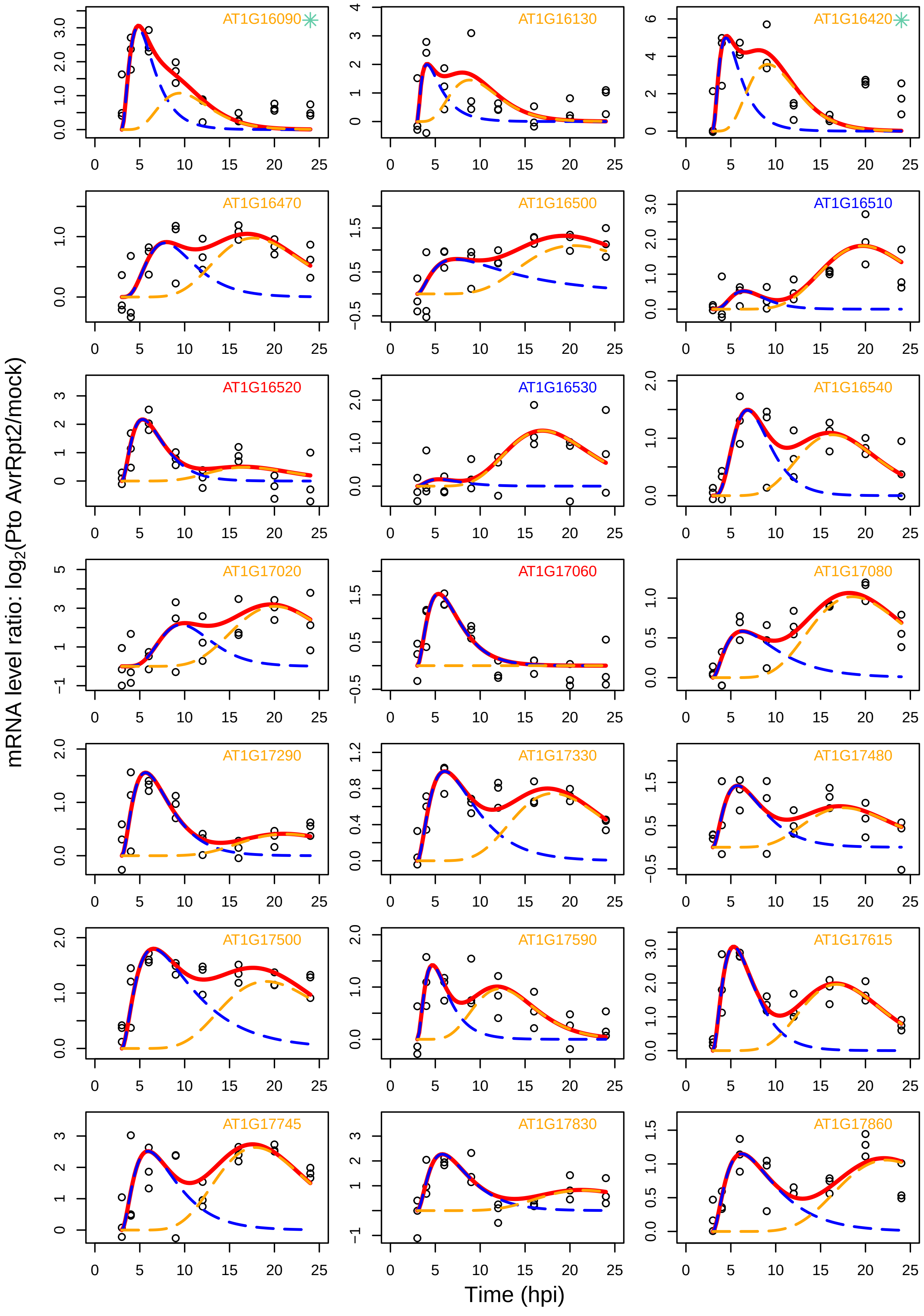
AT1G16090
AT1G16130
AT1G16420
AT1G16470
AT1G16500
AT1G16510
AT1G16520
AT1G16530
AT1G16540
AT1G17020
AT1G17060
AT1G17080
AT1G17290
AT1G17330
AT1G17480
AT1G17500
AT1G17590
AT1G17615
AT1G17745
AT1G17830
AT1G17860
mRNA level ratio: log2(Pto AvrRpt2/mock)
Time (hpi)

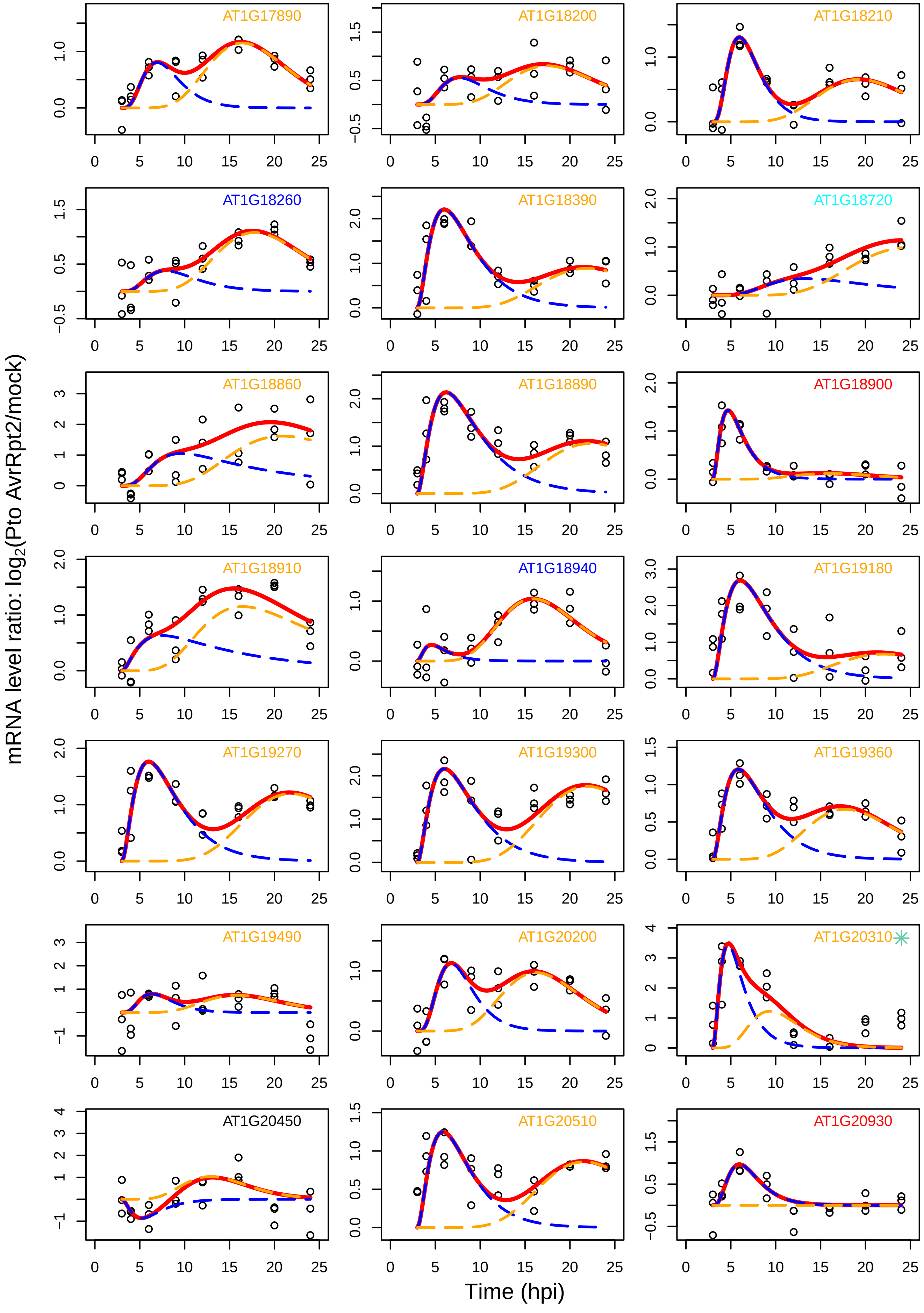
AT1G17890
AT1G18200
AT1G18210
AT1G18260
AT1G18390
AT1G18720
AT1G18860
AT1G18890
AT1G18900
AT1G18910
AT1G18940
AT1G19180
AT1G19270
AT1G19300
AT1G19360
AT1G19490
AT1G20200
AT1G20310
AT1G20450
AT1G20510
AT1G20930
mRNA level ratio: $\log_2$(Pto AvrRpt2/mock)
Time (hpi)

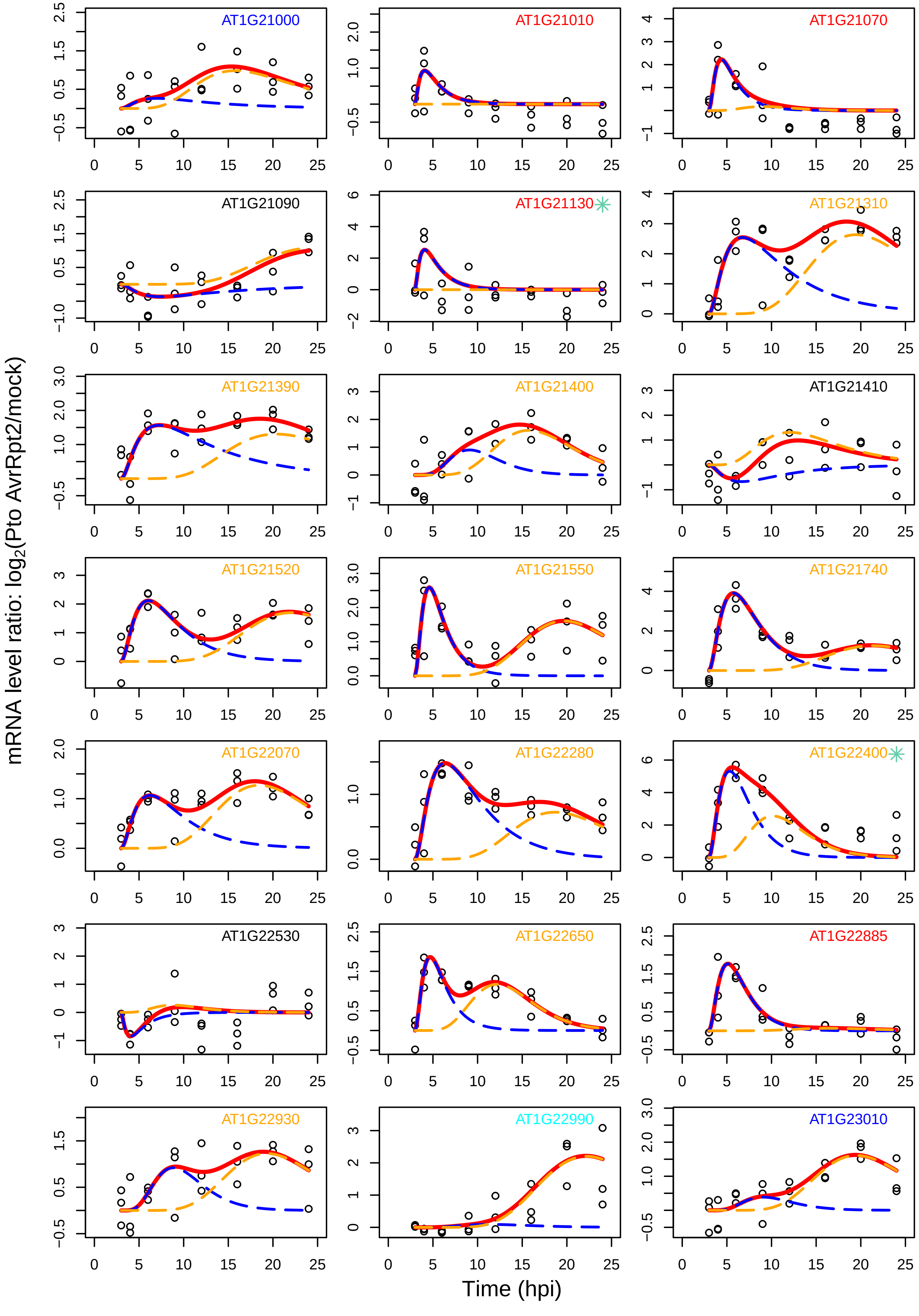
AT1G21000
AT1G21010
AT1G21070
AT1G21090
AT1G21130
AT1G21310
AT1G21390
AT1G21400
AT1G21410
AT1G21520
AT1G21550
AT1G21740
AT1G22070
AT1G22280
AT1G22400
AT1G22530
AT1G22650
AT1G22885
AT1G22930
AT1G22990
AT1G23010
mRNA level ratio: log2(Pto AvrRpt2/mock)
Time (hpi)

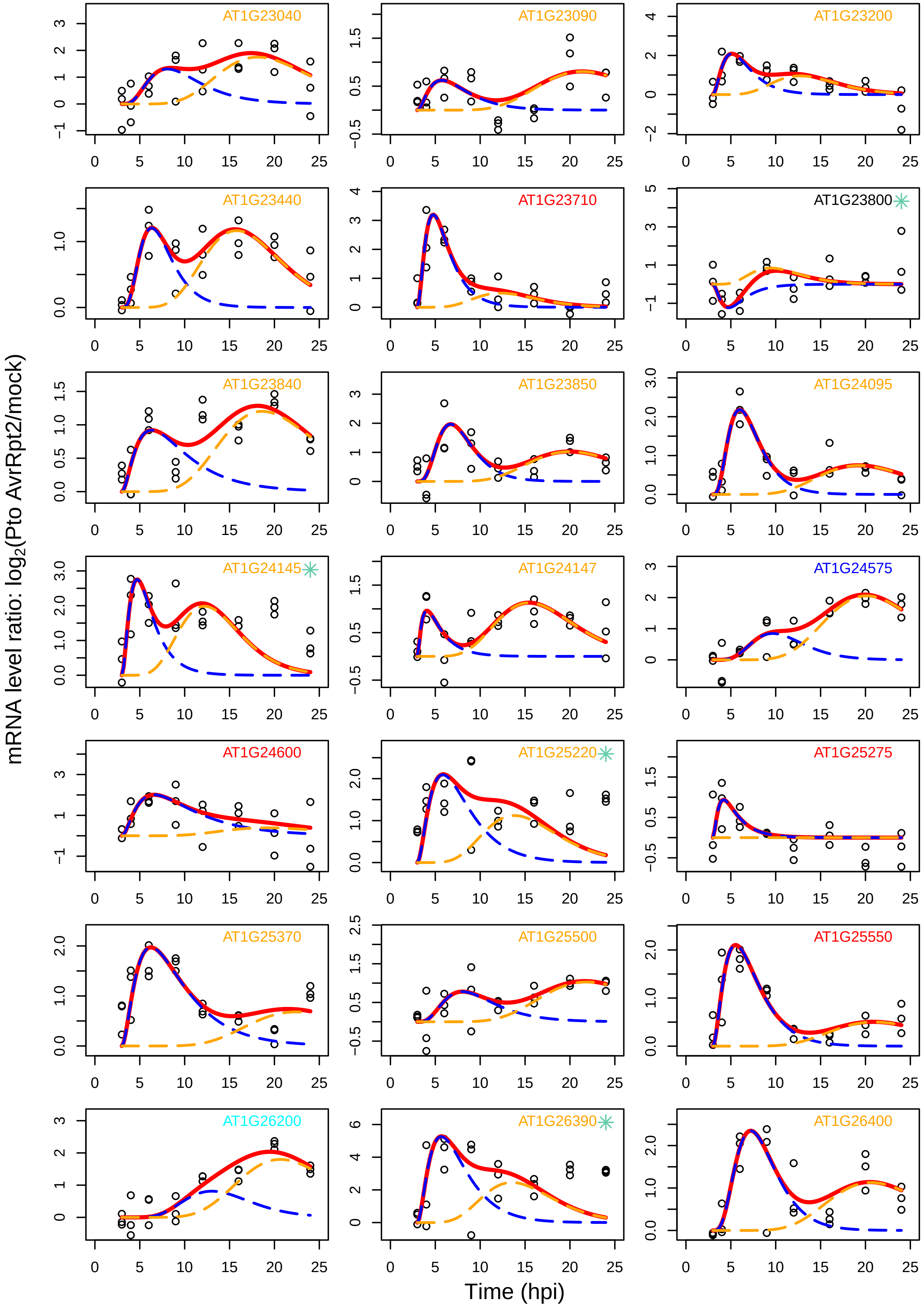

AT1G23040
AT1G23090
AT1G23200
AT1G23440
AT1G23710
AT1G23800
AT1G23840
AT1G23850
AT1G24095
AT1G24145
AT1G24147
AT1G24575
AT1G24600
AT1G25220
AT1G25275
AT1G25370
AT1G25500
AT1G25550
AT1G26200
AT1G26390
AT1G26400
mRNA level ratio: $log_2$(Pto AvrRpt2/mock)
Time (hpi)

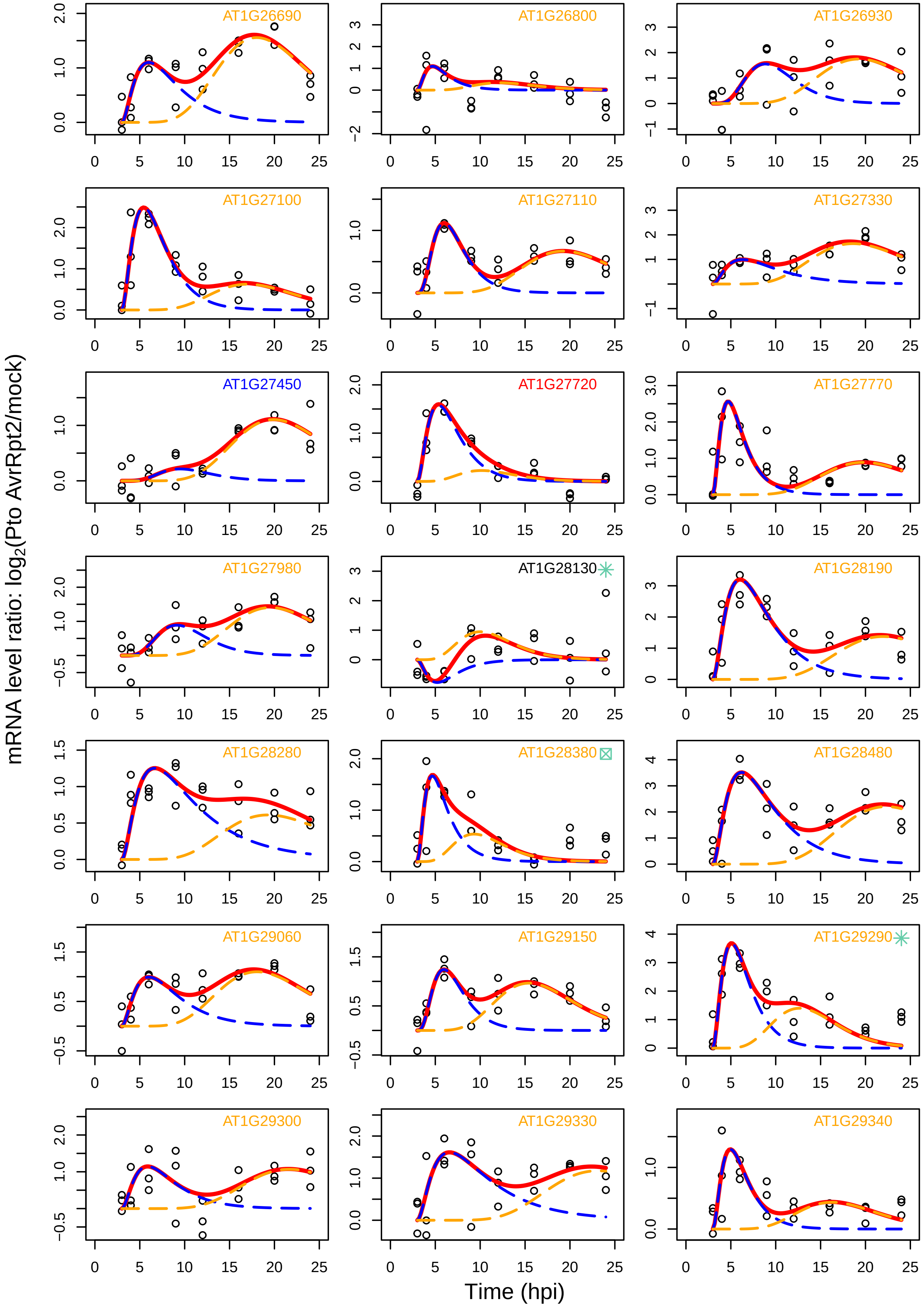
AT1G26690
AT1G26800
AT1G26930
AT1G27100
AT1G27110
AT1G27330
AT1G27450
AT1G27720
AT1G27770
AT1G27980
AT1G28130
AT1G28190
AT1G28280
AT1G28380
AT1G28480
AT1G29060
AT1G29150
AT1G29290
AT1G29300
AT1G29330
AT1G29340
mRNA level ratio: log2(Pto AvrRpt2/mock)
Time (hpi)

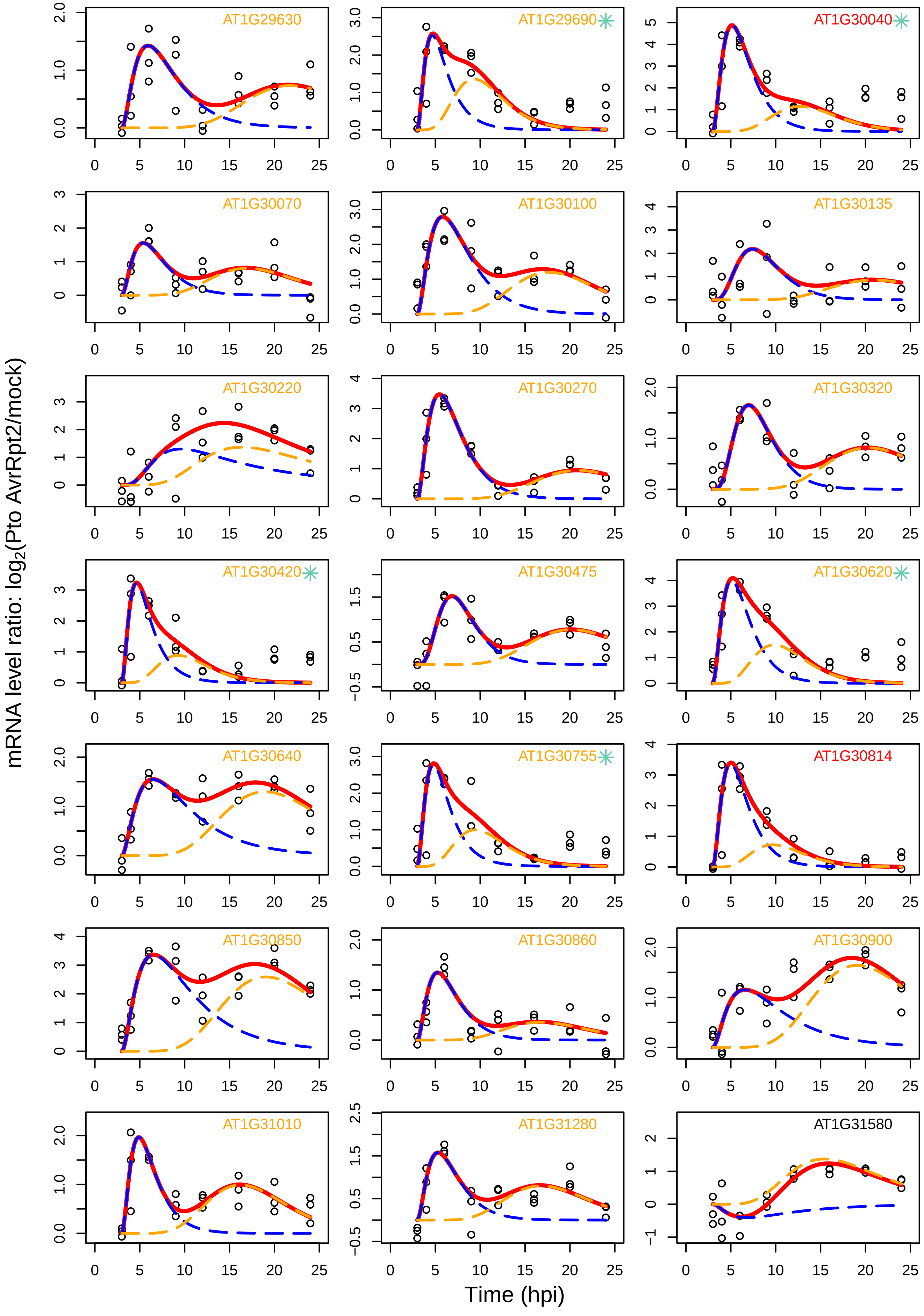

AT1G29630
AT1G29690
AT1G30040
AT1G30070
AT1G30100
AT1G30135
AT1G30220
AT1G30270
AT1G30320
AT1G30420
AT1G30475
AT1G30620
AT1G30640
AT1G30755
AT1G30814
AT1G30850
AT1G30860
AT1G30900
AT1G31010
AT1G31280
AT1G31580
mRNA level ratio: $\log_2$(Pto AvrRpt2/mock)
Time (hpi)

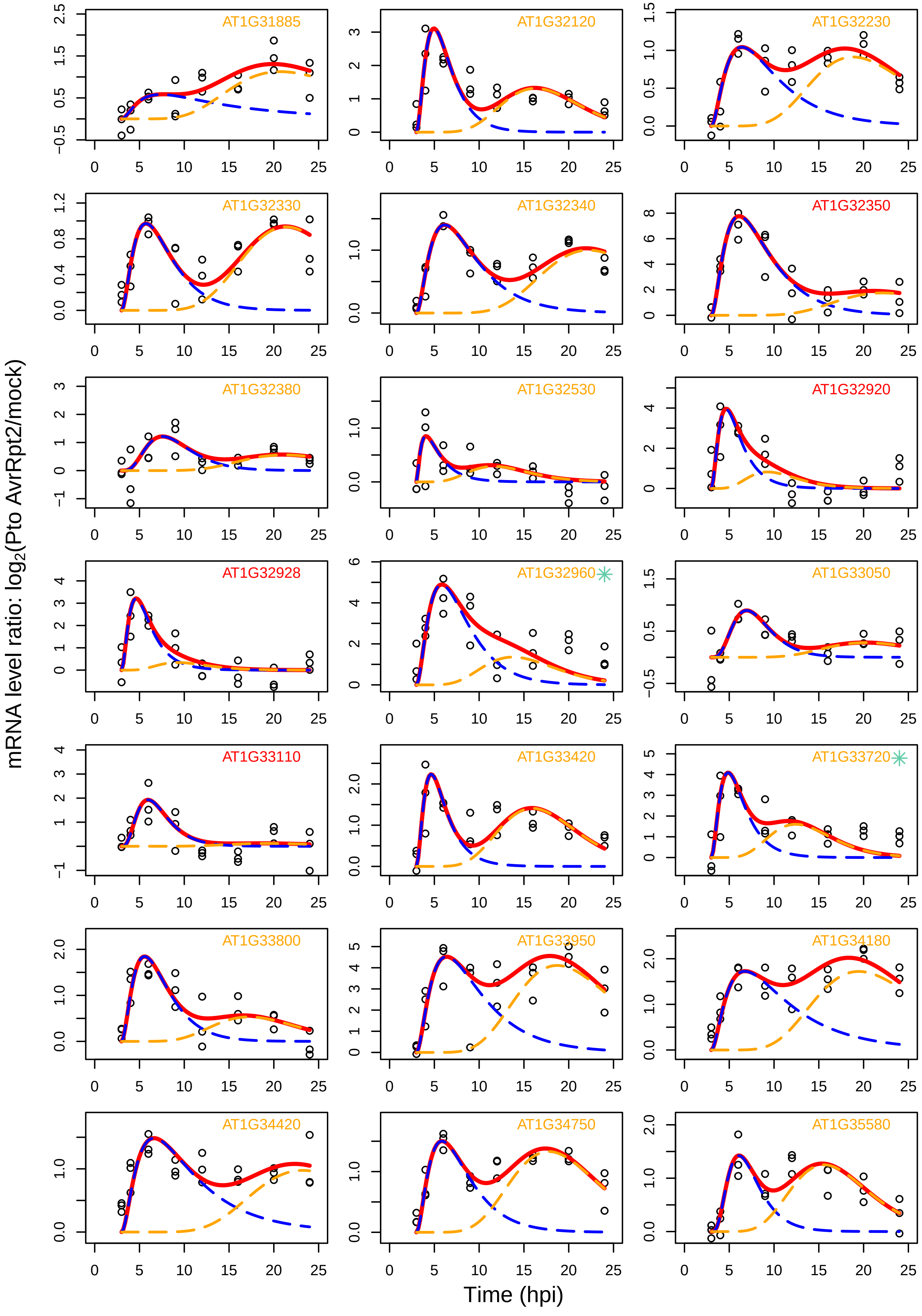

mRNA level ratio: $\log_2$(Pto AvrRpt2/mock)
Time (hpi)
AT1G31885
AT1G32120
AT1G32230
AT1G32330
AT1G32340
AT1G32350
AT1G32380
AT1G32530
AT1G32920
AT1G32928
AT1G32960
AT1G33050
AT1G33110
AT1G33420
AT1G33720
AT1G33800
AT1G33950
AT1G34180
AT1G34420
AT1G34750
AT1G35580

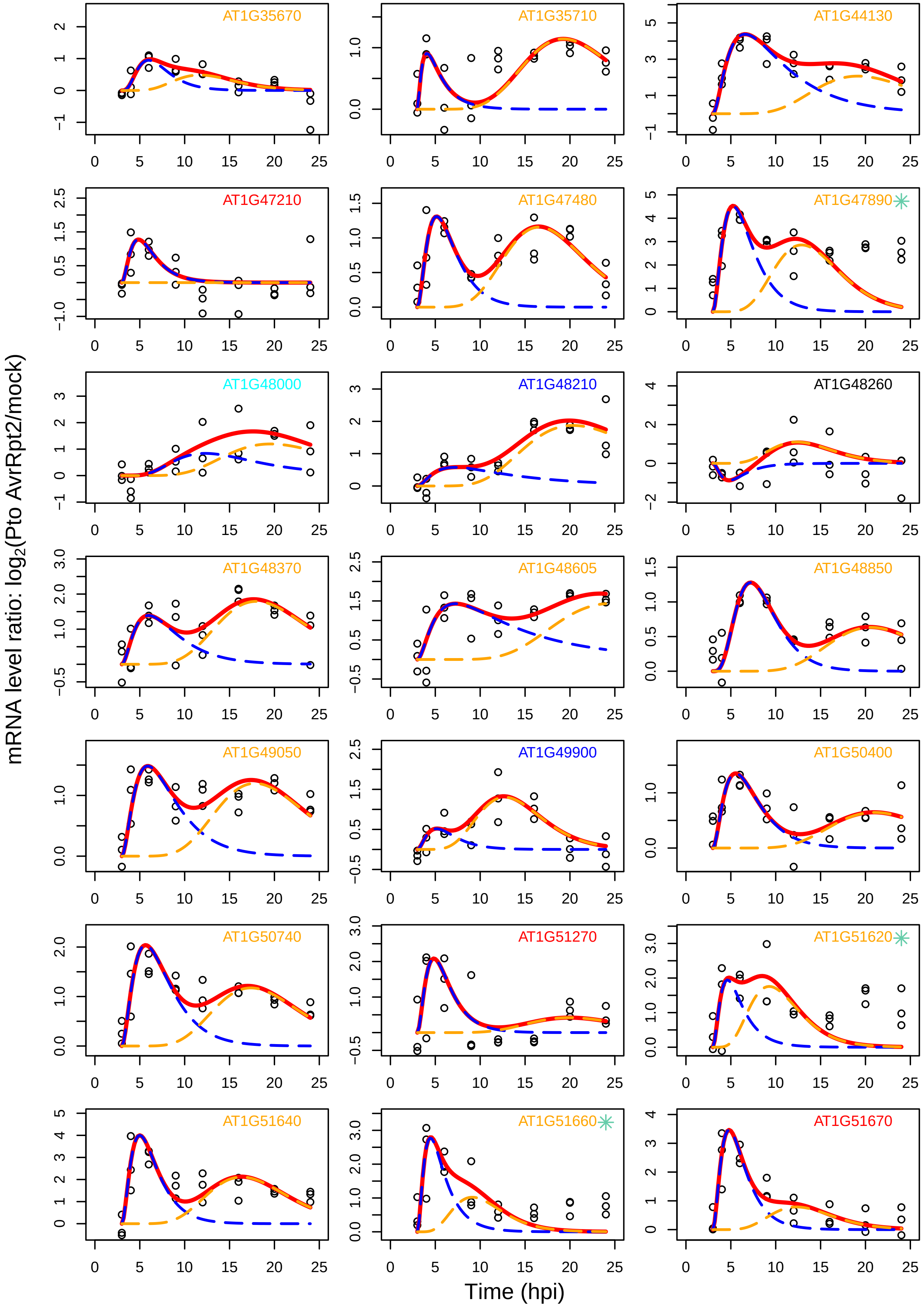

AT1G35670
AT1G35710
AT1G44130
AT1G47210
AT1G47480
AT1G47890
AT1G48000
AT1G48210
AT1G48260
AT1G48370
AT1G48605
AT1G48850
AT1G49050
AT1G49900
AT1G50400
AT1G50740
AT1G51270
AT1G51620
AT1G51640
AT1G51660
AT1G51670
mRNA level ratio: log2(Pto AvrRpt2/mock)
Time (hpi)

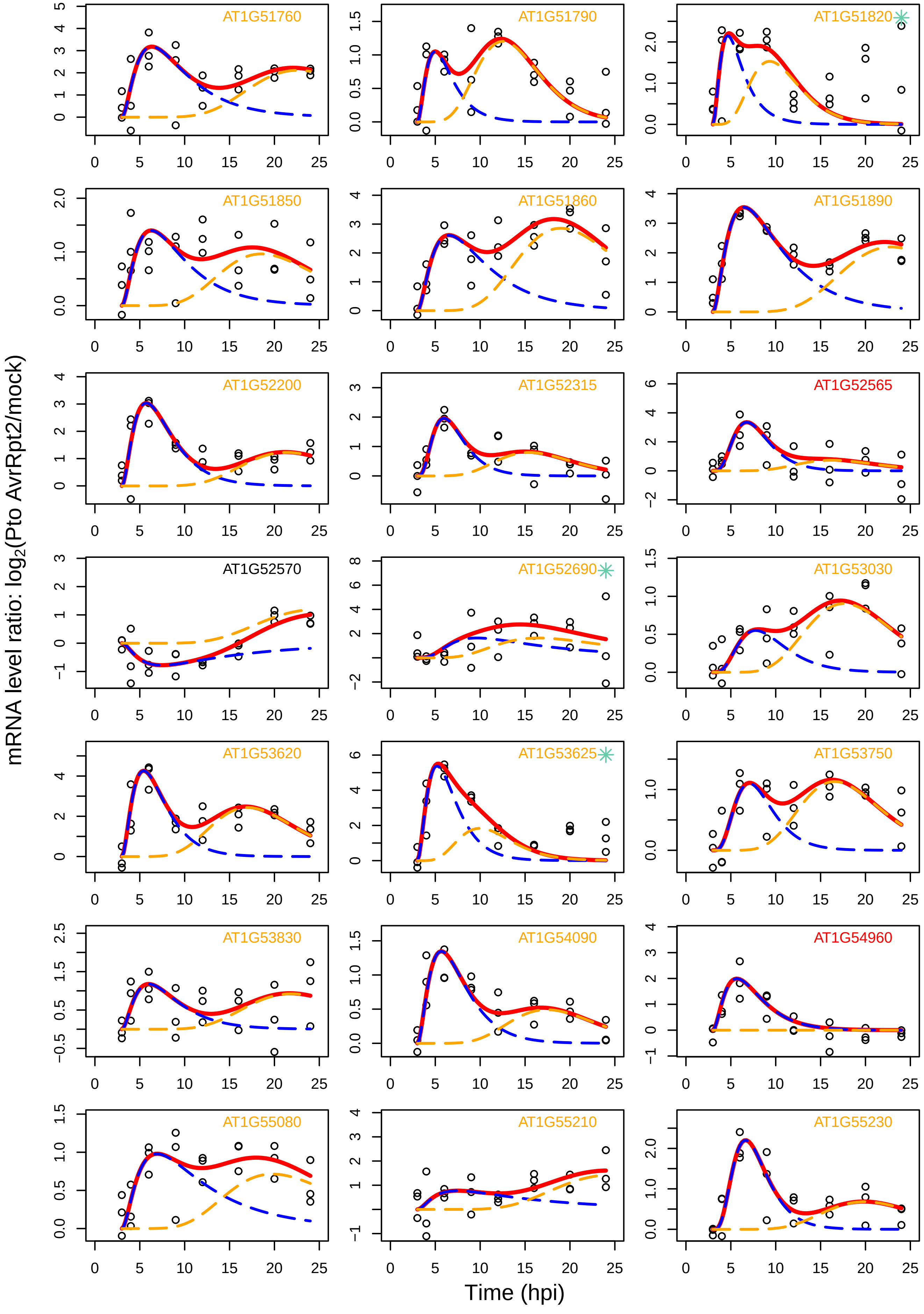
AT1G51760
AT1G51790
AT1G51820
AT1G51850
AT1G51860
AT1G51890
AT1G52200
AT1G52315
AT1G52565
AT1G52570
AT1G52690
AT1G53030
AT1G53620
AT1G53625
AT1G53750
AT1G53830
AT1G54090
AT1G54960
AT1G55080
AT1G55210
AT1G55230
mRNA level ratio: log2(Pto AvrRpt2/mock)
Time (hpi)

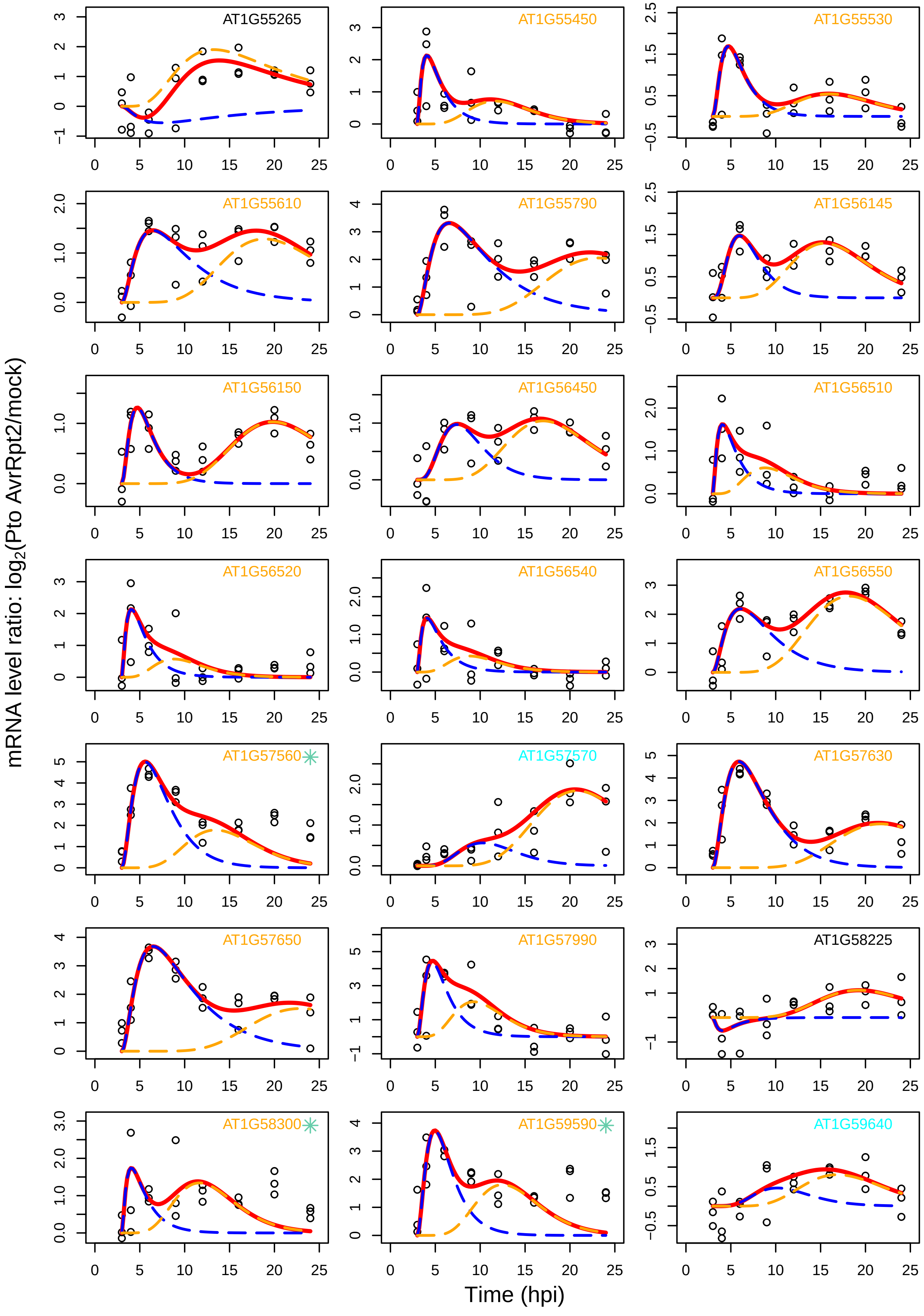

mRNA level ratio: log2(Pto AvrRpt2/mock)
Time (hpi)
AT1G55265
AT1G55450
AT1G55530
AT1G55610
AT1G55790
AT1G56145
AT1G56150
AT1G56450
AT1G56510
AT1G56520
AT1G56540
AT1G56550
AT1G57560
AT1G57570
AT1G57630
AT1G57650
AT1G57990
AT1G58225
AT1G58300
AT1G59590
AT1G59640

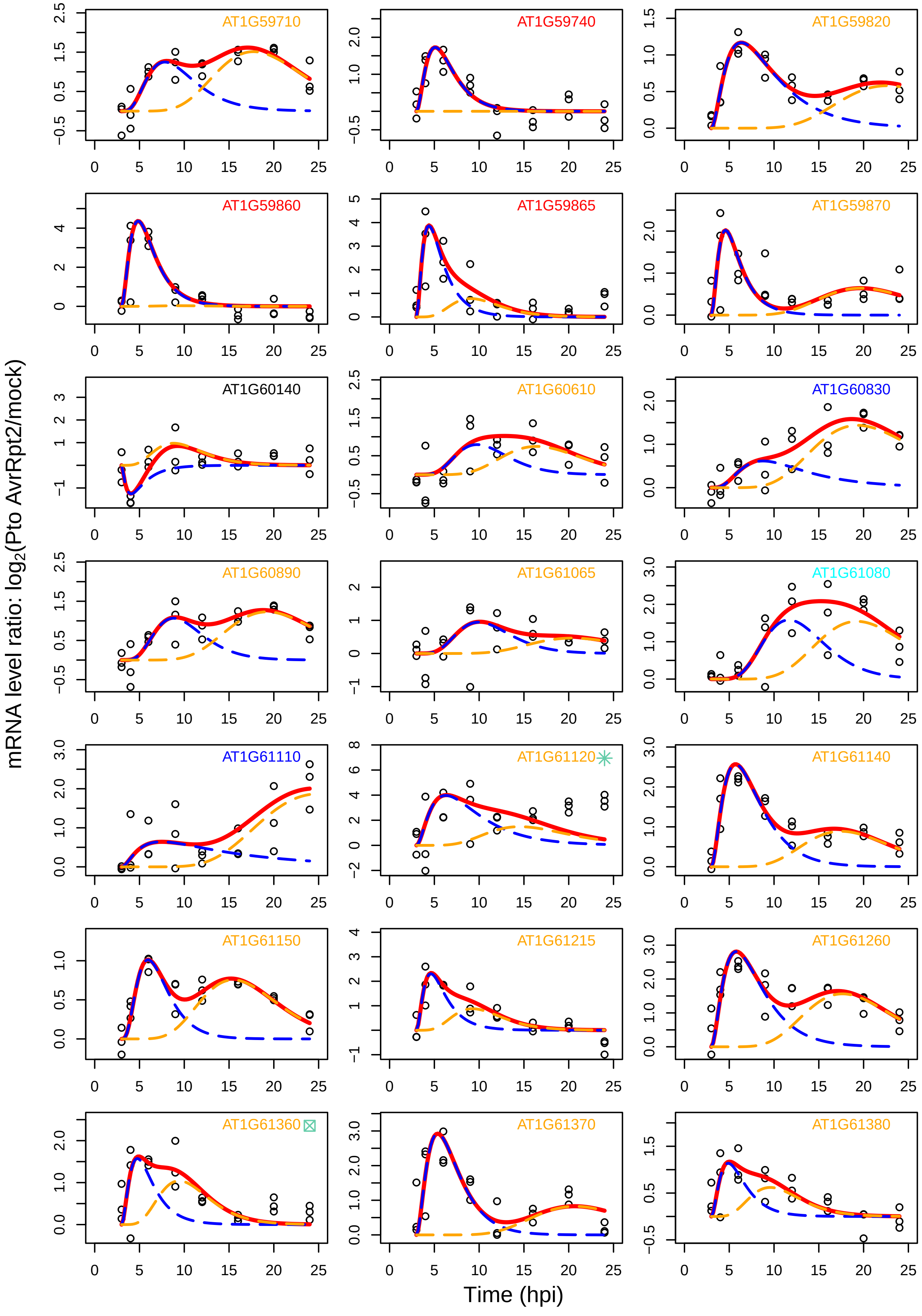
AT1G59710
AT1G59740
AT1G59820
AT1G59860
AT1G59865
AT1G59870
AT1G60140
AT1G60610
AT1G60830
AT1G60890
AT1G61065
AT1G61080
AT1G61110
AT1G61120
AT1G61140
AT1G61150
AT1G61215
AT1G61260
AT1G61360
AT1G61370
AT1G61380
mRNA level ratio: log2(Pto AvrRpt2/mock)
Time (hpi)

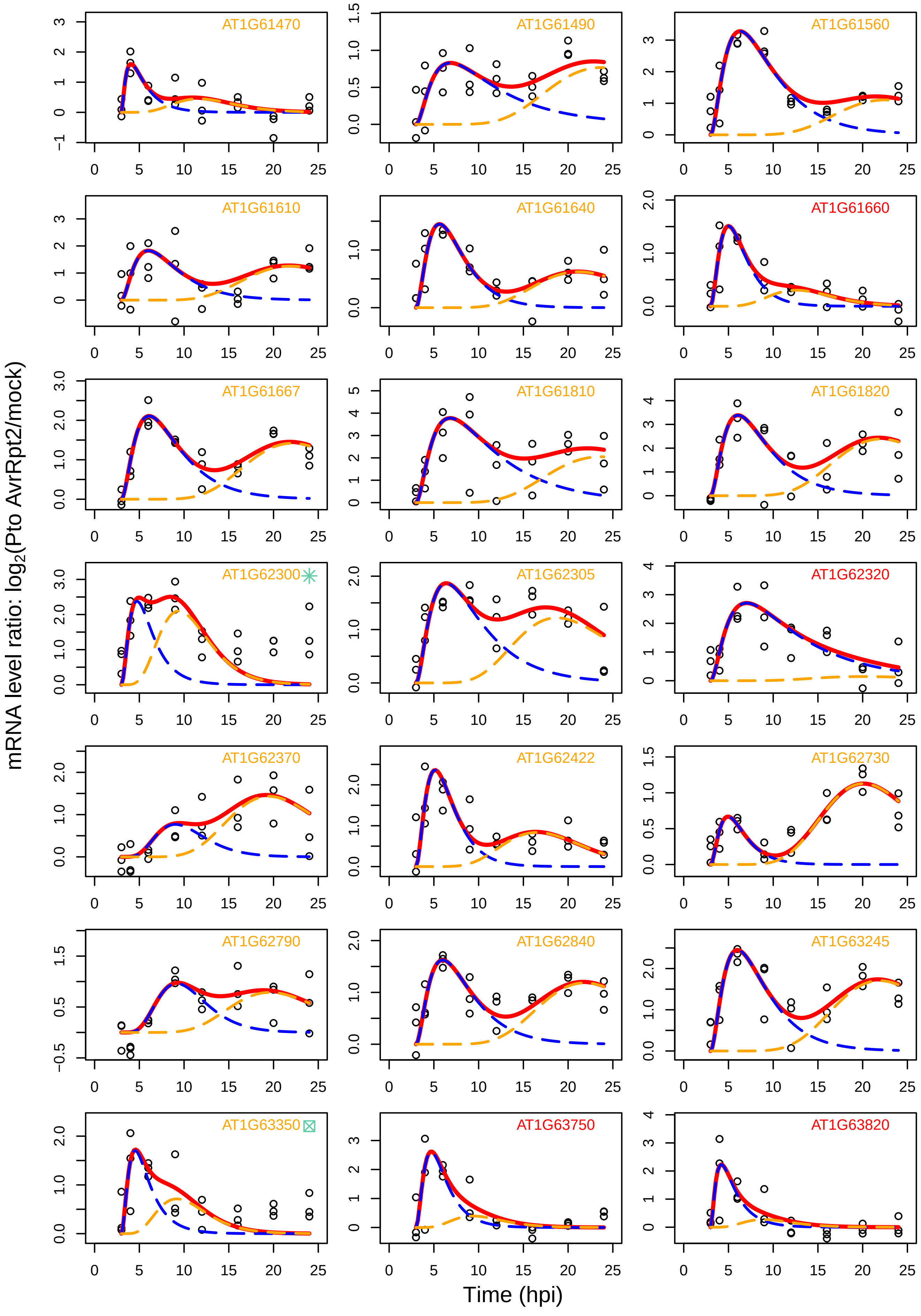
AT1G61470
AT1G61490
AT1G61560
AT1G61610
AT1G61640
AT1G61660
AT1G61667
AT1G61810
AT1G61820
AT1G62300
AT1G62305
AT1G62320
AT1G62370
AT1G62422
AT1G62730
AT1G62790
AT1G62840
AT1G63245
AT1G63350
AT1G63750
AT1G63820
mRNA level ratio: log2(Pto AvrRpt2/mock)
Time (hpi)

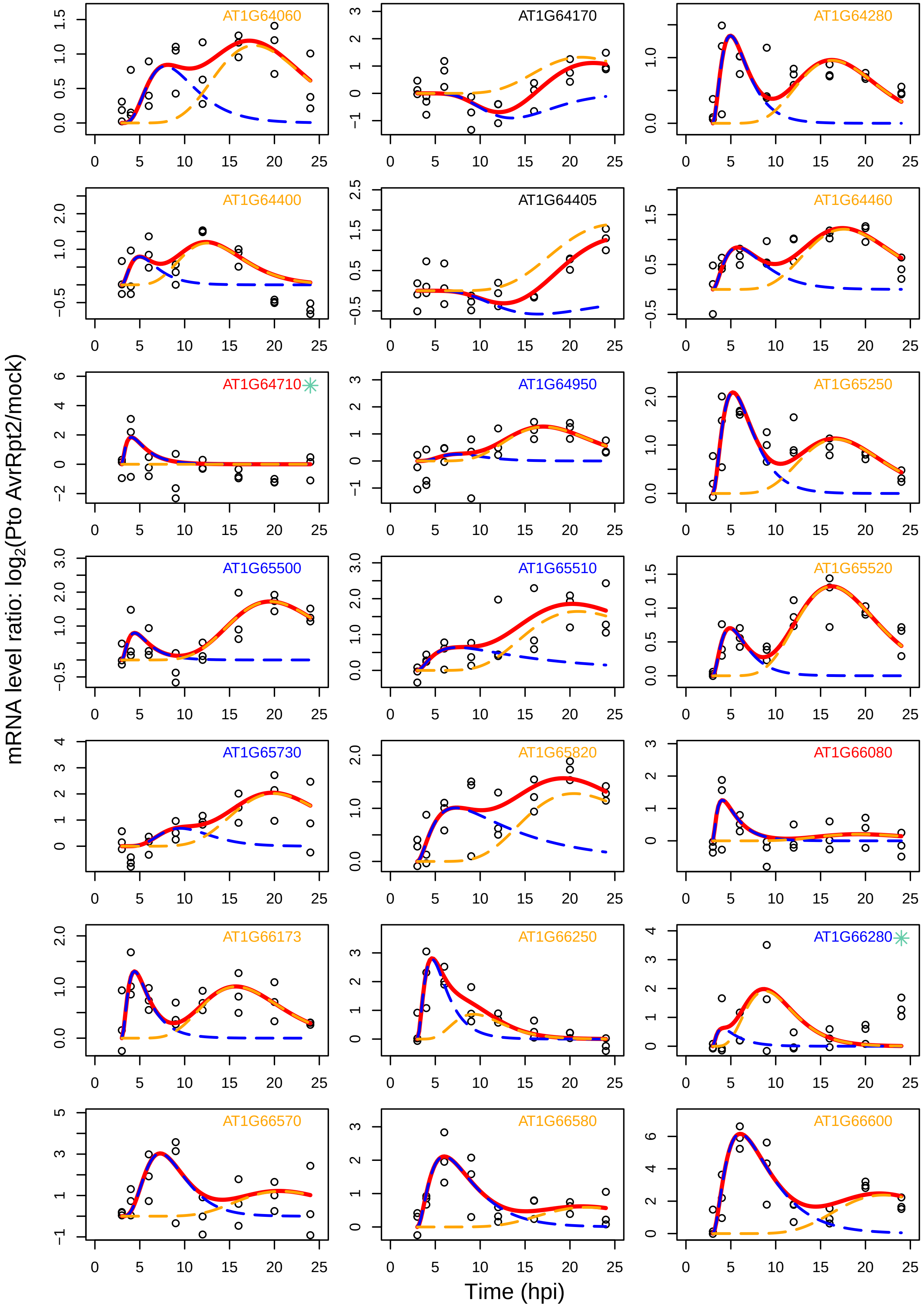
AT1G64060
AT1G64170
AT1G64280
AT1G64400
AT1G64405
AT1G64460
AT1G64710
AT1G64950
AT1G65250
AT1G65500
AT1G65510
AT1G65520
AT1G65730
AT1G65820
AT1G66080
AT1G66173
AT1G66250
AT1G66280
AT1G66570
AT1G66580
AT1G66600
mRNA level ratio: log2(Pto AvrRpt2/mock)
Time (hpi)

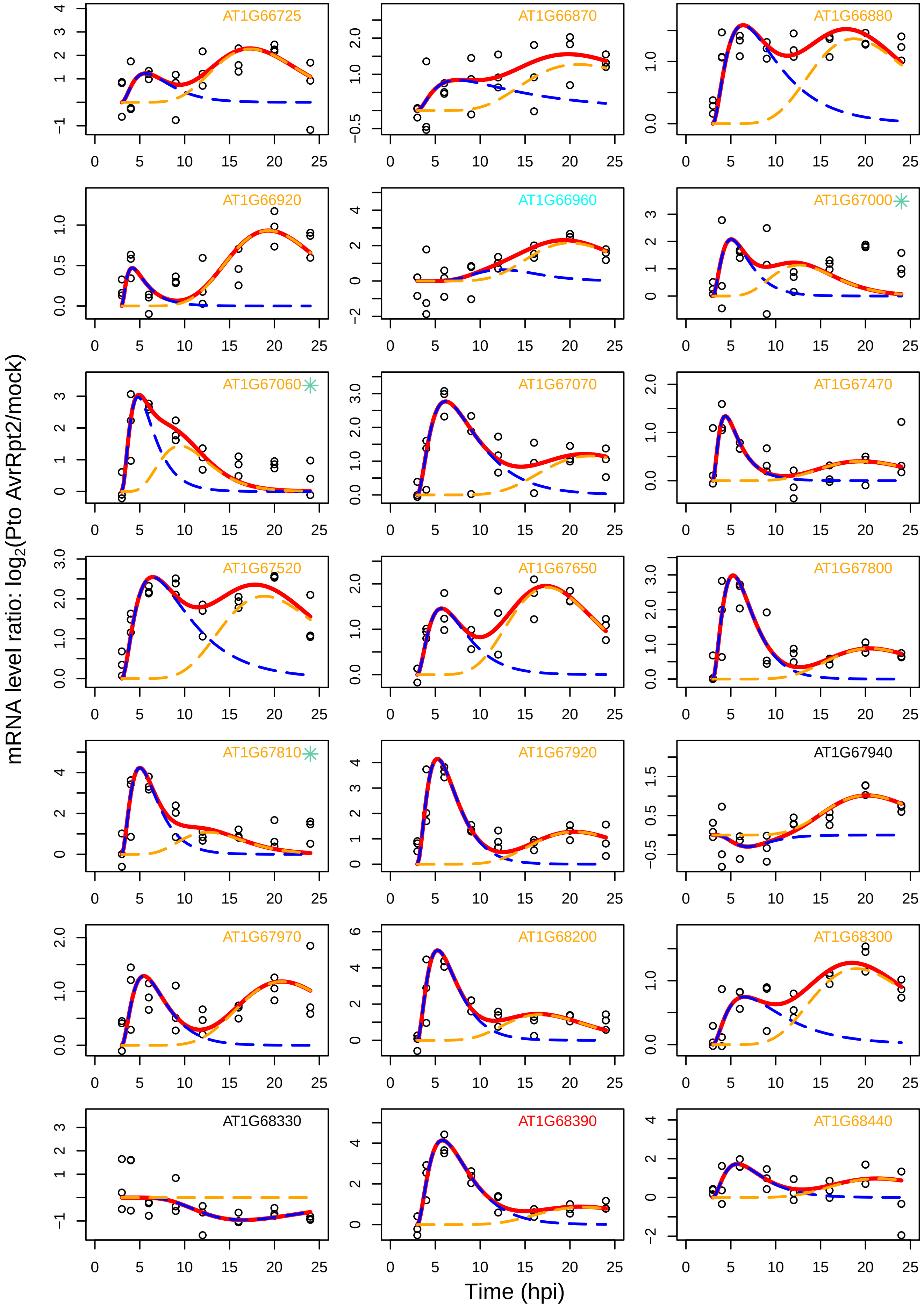

mRNA level ratio: $log_2$(Pto AvrRpt2/mock)
Time (hpi)
AT1G66725
AT1G66870
AT1G66880
AT1G66920
AT1G66960
AT1G67000
AT1G67060
AT1G67070
AT1G67470
AT1G67520
AT1G67650
AT1G67800
AT1G67810
AT1G67920
AT1G67940
AT1G67970
AT1G68200
AT1G68300
AT1G68330
AT1G68390
AT1G68440

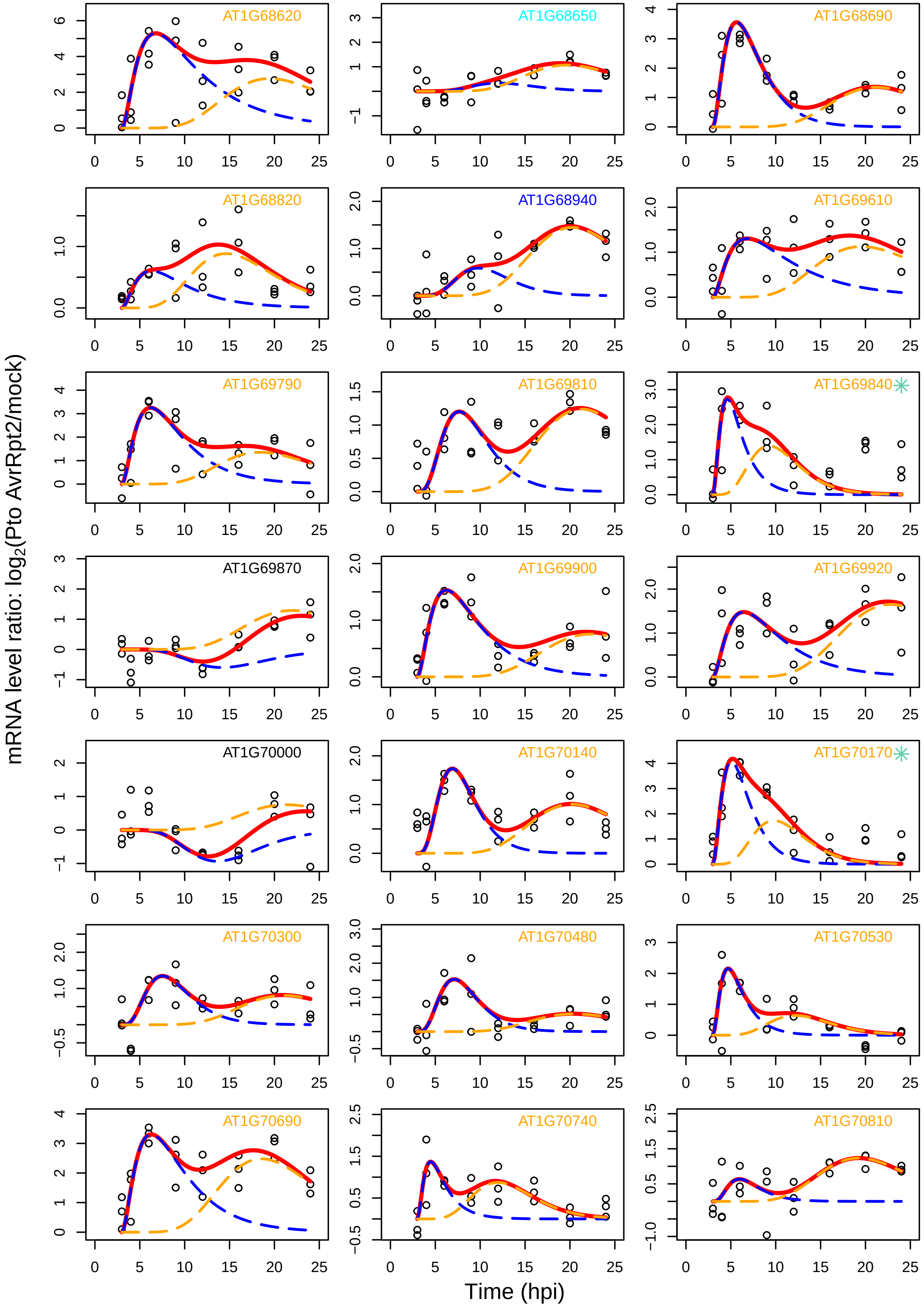

AT1G68620
AT1G68650
AT1G68690
AT1G68820
AT1G68940
AT1G69610
AT1G69790
AT1G69810
AT1G69840
AT1G69870
AT1G69900
AT1G69920
AT1G70000
AT1G70140
AT1G70170
AT1G70300
AT1G70480
AT1G70530
AT1G70690
AT1G70740
AT1G70810
mRNA level ratio: log2(Pto AvrRpt2/mock)
Time (hpi)

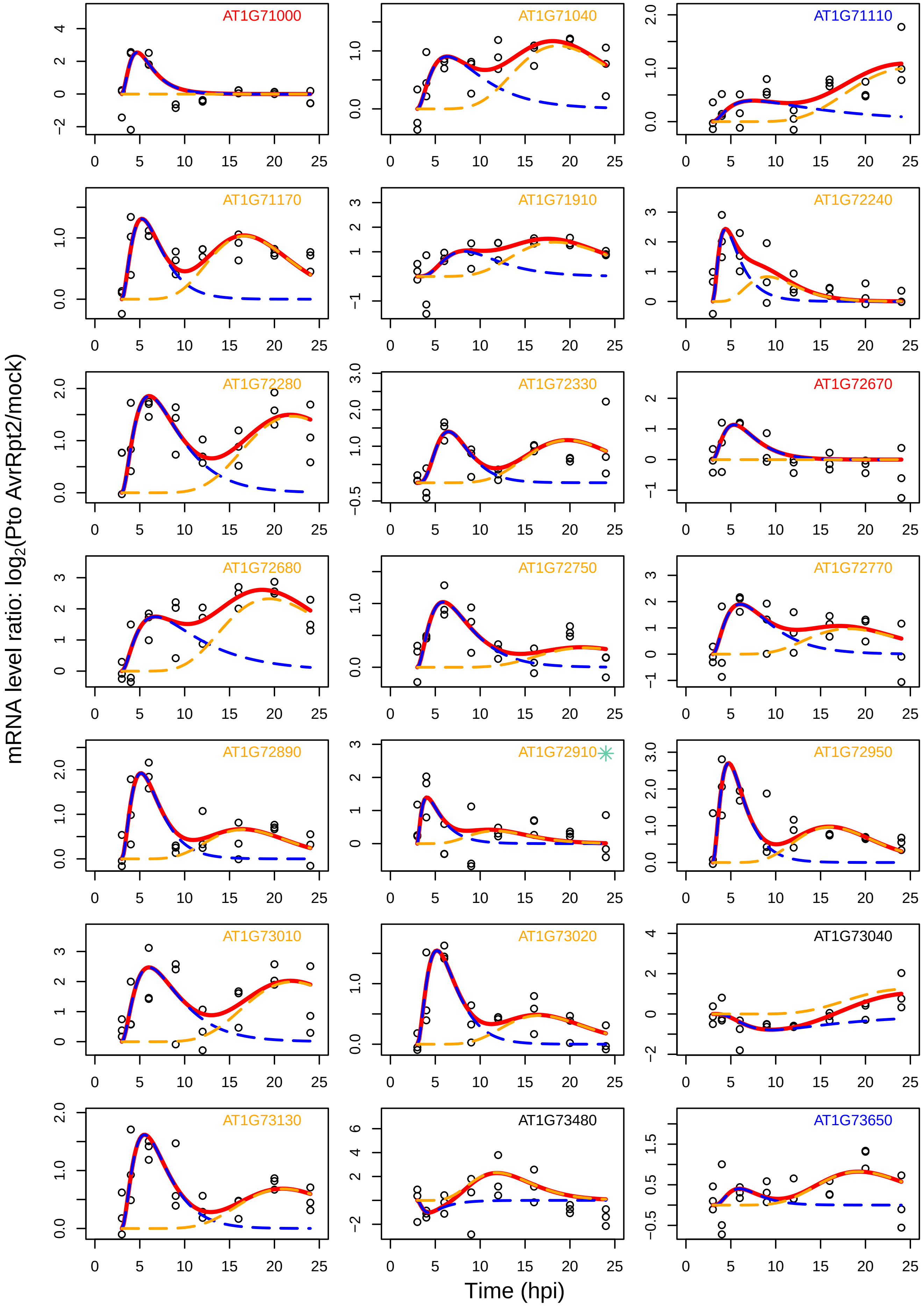
AT1G71000
AT1G71040
AT1G71110
AT1G71170
AT1G71910
AT1G72240
AT1G72280
AT1G72330
AT1G72670
AT1G72680
AT1G72750
AT1G72770
AT1G72890
AT1G72910
AT1G72950
AT1G73010
AT1G73020
AT1G73040
AT1G73130
AT1G73480
AT1G73650
mRNA level ratio: log2(Pto AvrRpt2/mock)
Time (hpi)

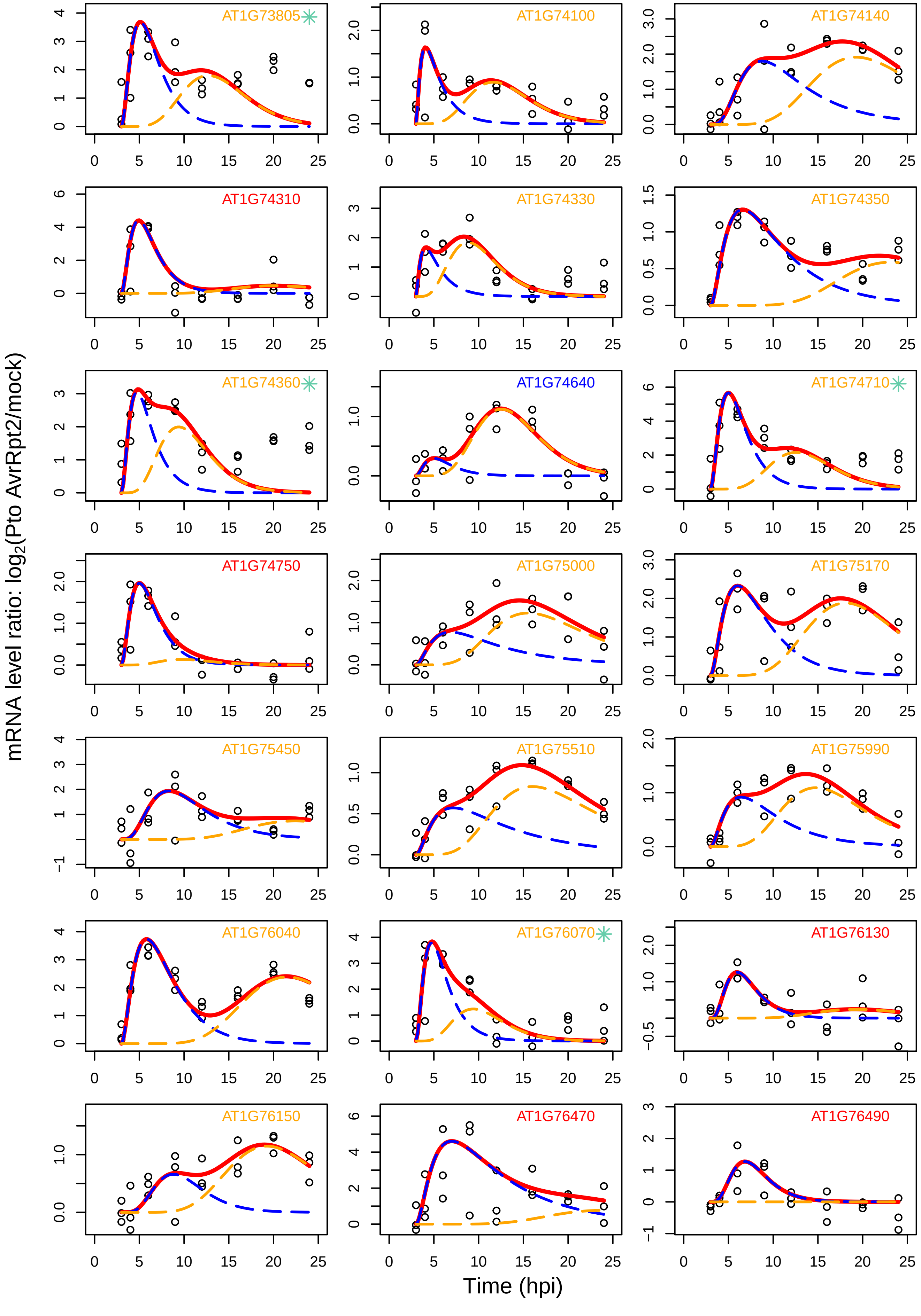

AT1G73805
AT1G74100
AT1G74140
AT1G74310
AT1G74330
AT1G74350
AT1G74360
AT1G74640
AT1G74710
AT1G74750
AT1G75000
AT1G75170
AT1G75450
AT1G75510
AT1G75990
AT1G76040
AT1G76070
AT1G76130
AT1G76150
AT1G76470
AT1G76490
mRNA level ratio: log2(Pto AvrRpt2/mock)
Time (hpi)

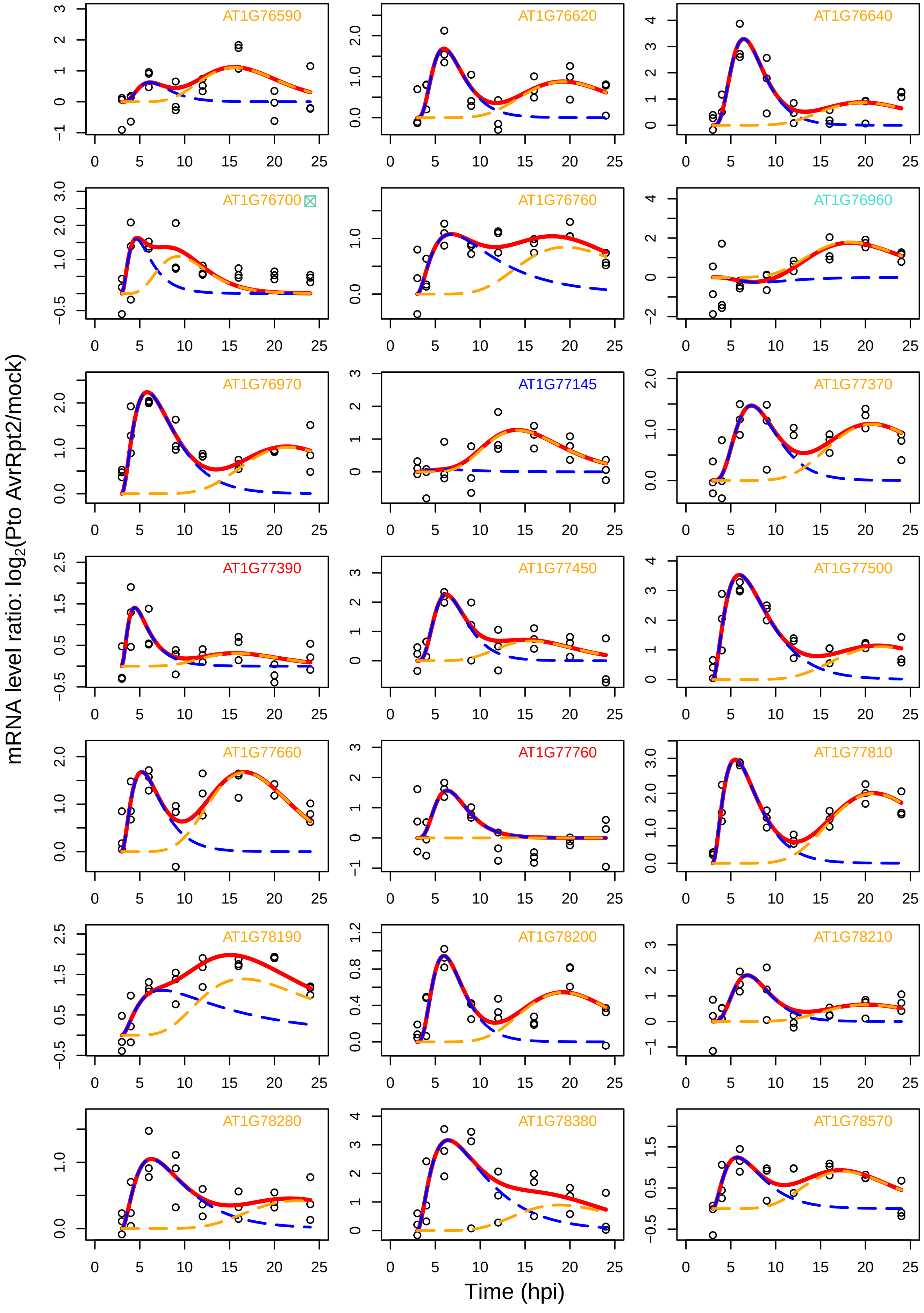
AT1G76590
AT1G76620
AT1G76640
AT1G76700
AT1G76760
AT1G76960
AT1G76970
AT1G77145
AT1G77370
AT1G77390
AT1G77450
AT1G77500
AT1G77660
AT1G77760
AT1G77810
AT1G78190
AT1G78200
AT1G78210
AT1G78280
AT1G78380
AT1G78570
mRNA level ratio: $log_2$(Pto AvrRpt2/mock)
Time (hpi)

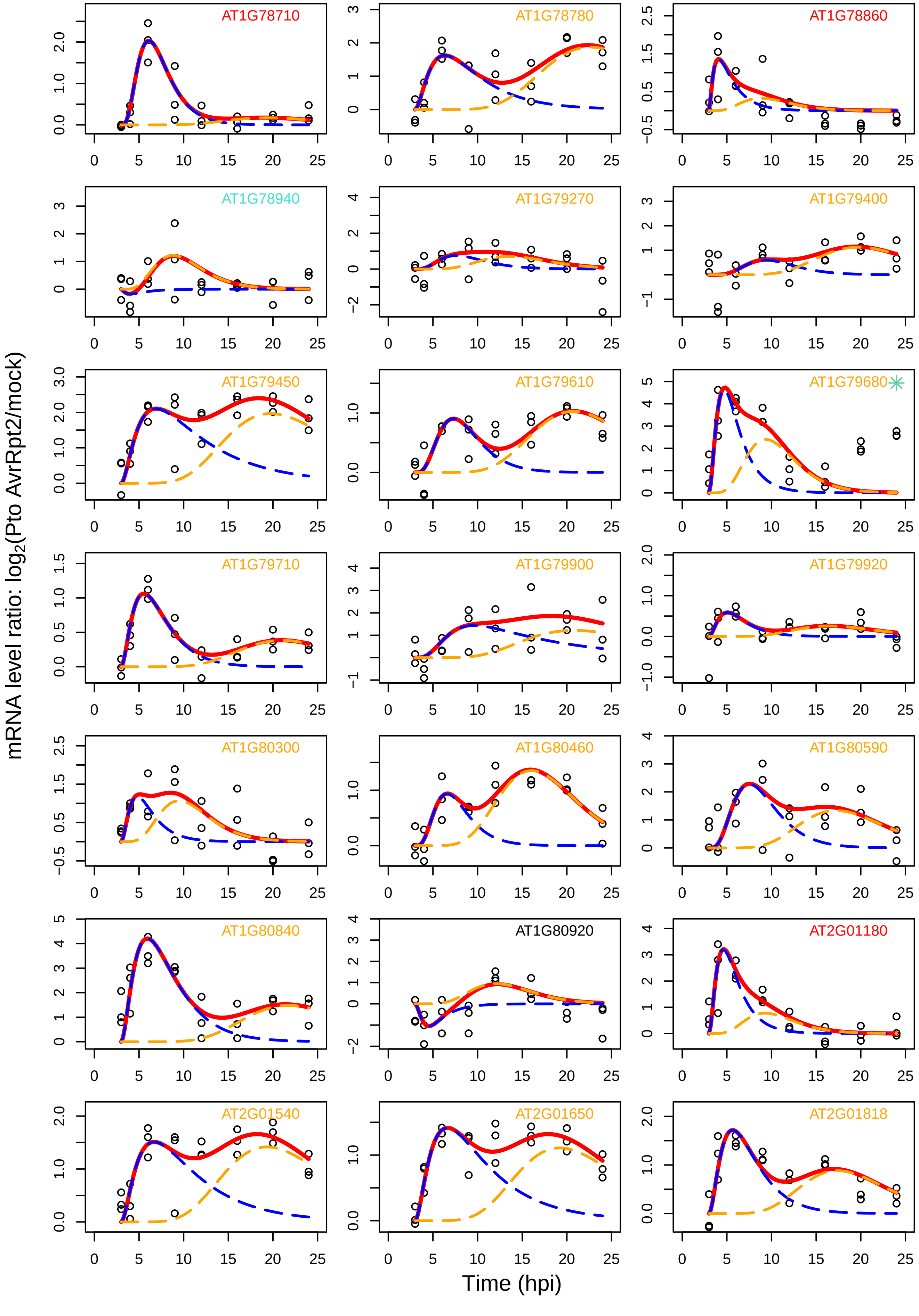

AT1G78710
AT1G78780
AT1G78860
AT1G78940
AT1G79270
AT1G79400
AT1G79450
AT1G79610
AT1G79680
AT1G79710
AT1G79900
AT1G79920
AT1G80300
AT1G80460
AT1G80590
AT1G80840
AT1G80920
AT2G01180
AT2G01540
AT2G01650
AT2G01818
mRNA level ratio: log2(Pto AvrRpt2/mock)
Time (hpi)

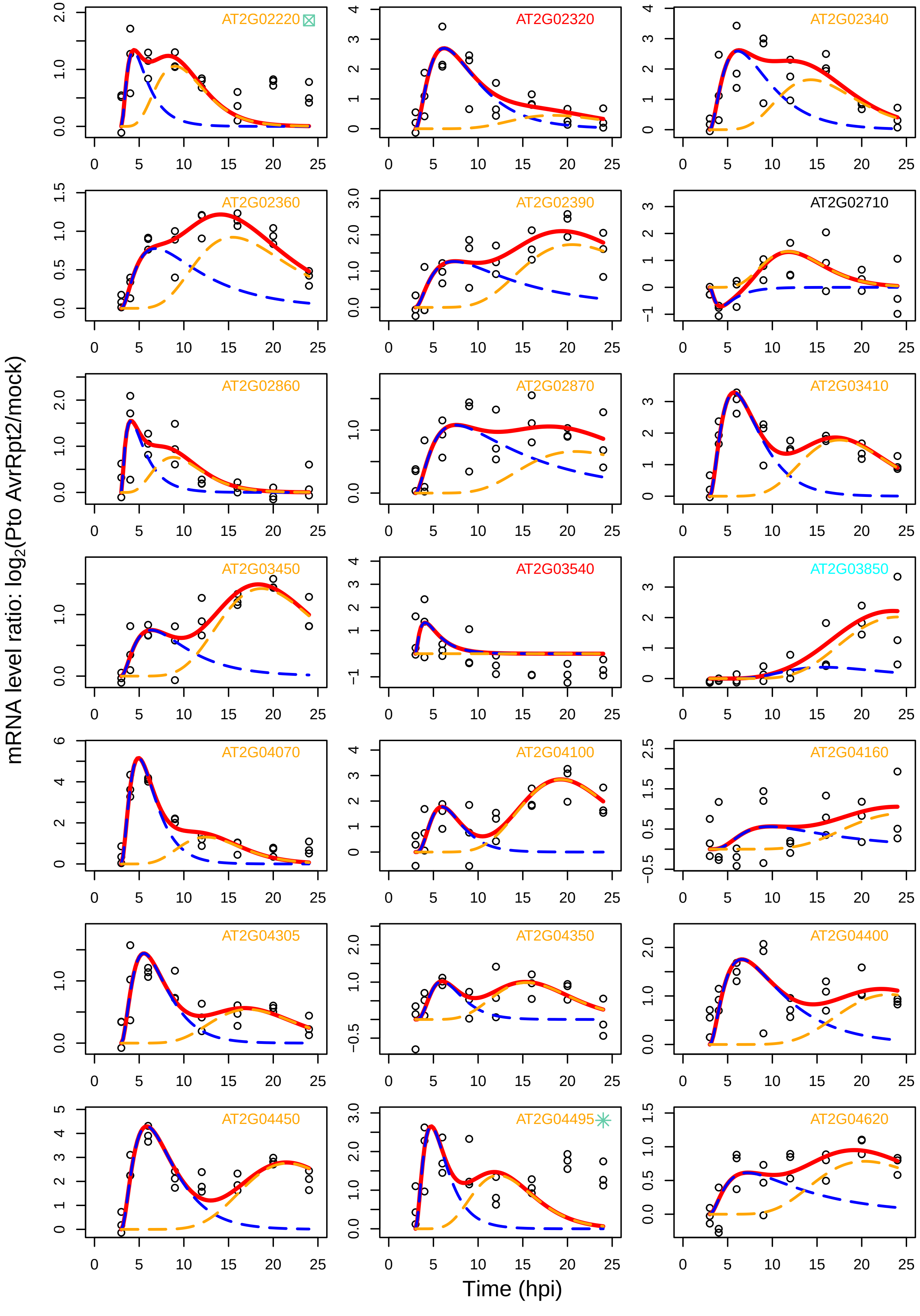
AT2G02220
AT2G02320
AT2G02340
AT2G02360
AT2G02390
AT2G02710
AT2G02860
AT2G02870
AT2G03410
AT2G03450
AT2G03540
AT2G03850
AT2G04070
AT2G04100
AT2G04160
AT2G04305
AT2G04350
AT2G04400
AT2G04450
AT2G04495
AT2G04620
mRNA level ratio: log2(Pto AvrRpt2/mock)
Time (hpi)

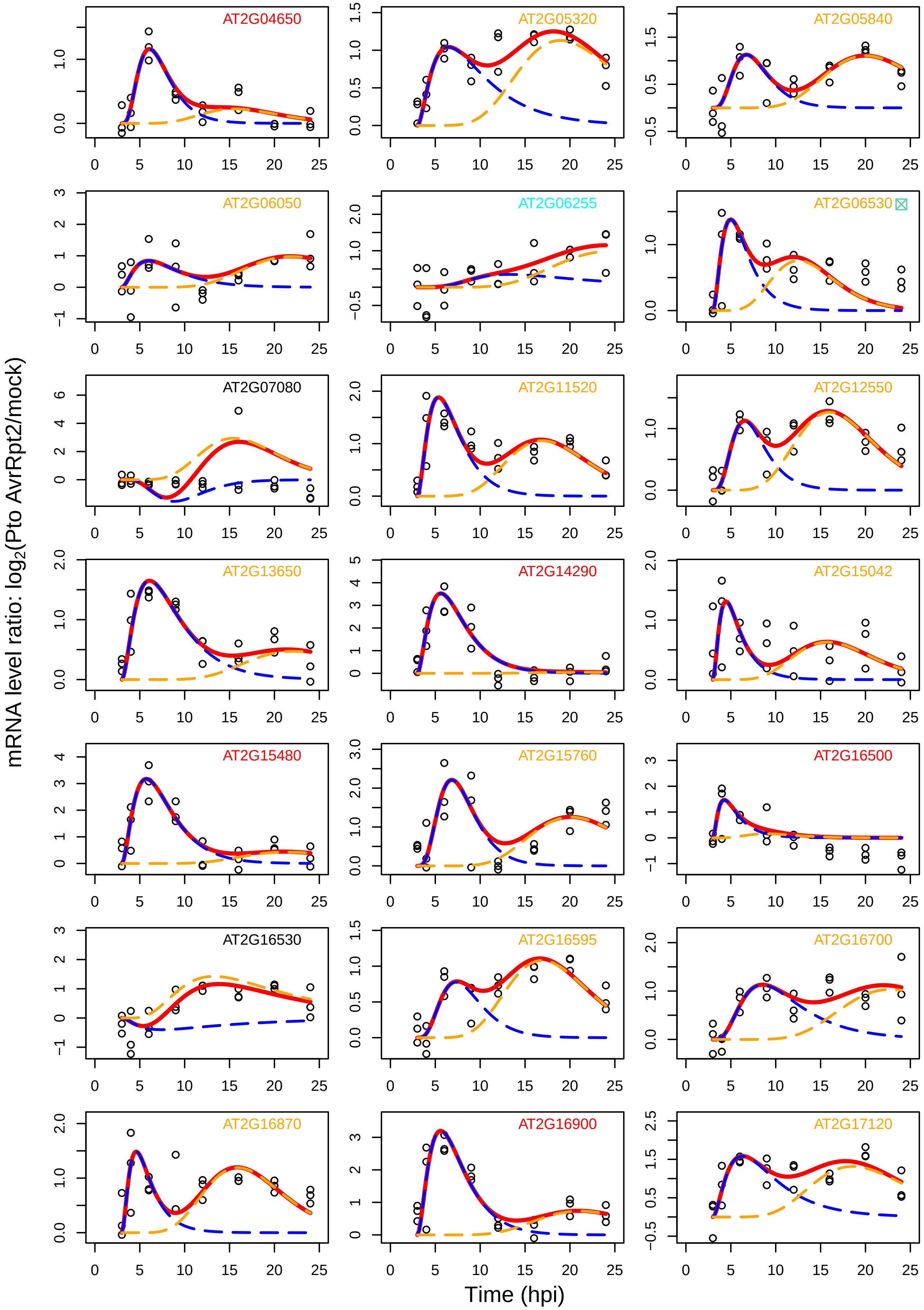
AT2G04650
AT2G05320
AT2G05840
AT2G06050
AT2G06255
AT2G06530
AT2G07080
AT2G11520
AT2G12550
AT2G13650
AT2G14290
AT2G15042
AT2G15480
AT2G15760
AT2G16500
AT2G16530
AT2G16595
AT2G16700
AT2G16870
AT2G16900
AT2G17120
mRNA level ratio: $\log_2$(Pto AvrRpt2/mock)
Time (hpi)

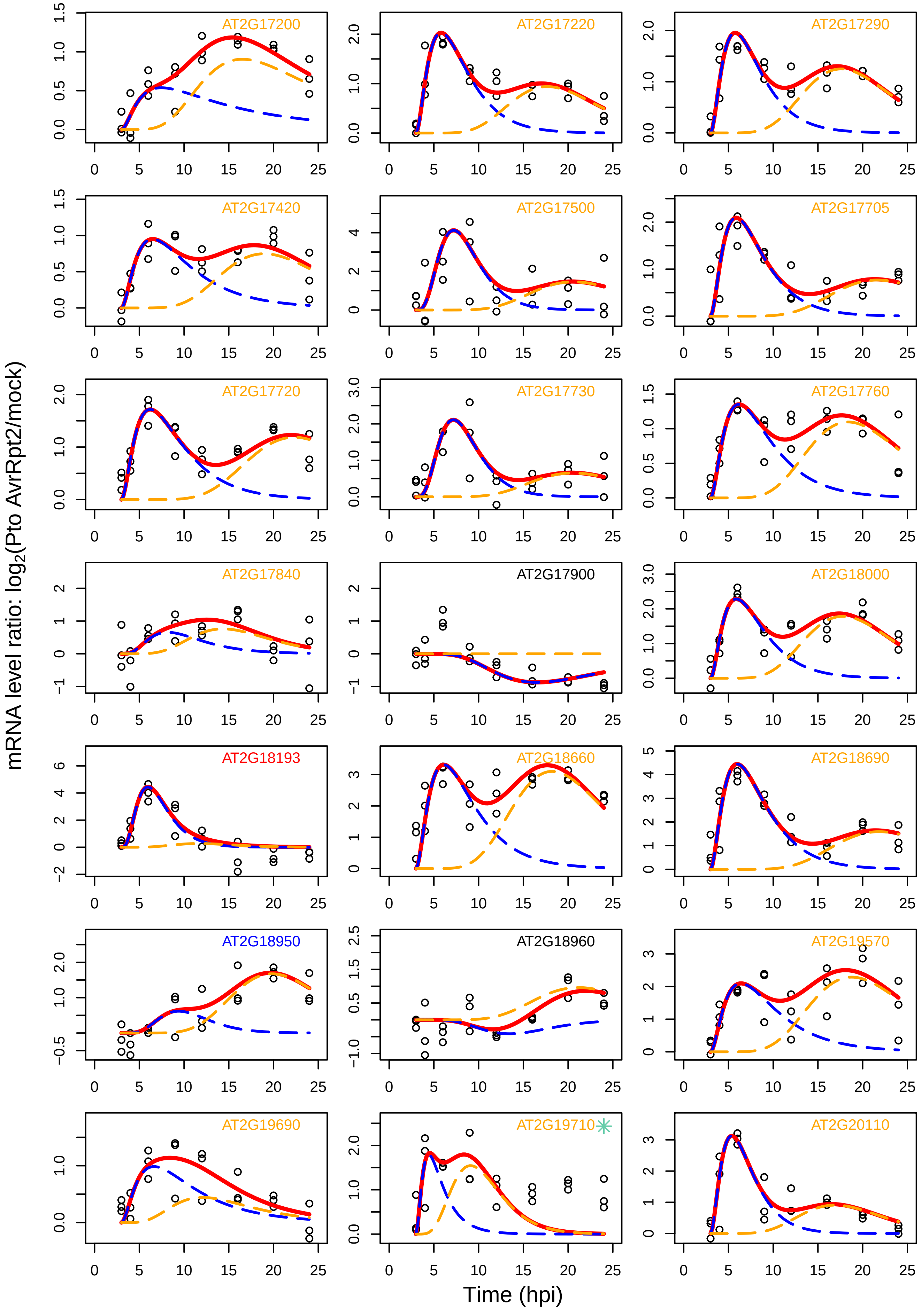

AT2G17200
AT2G17220
AT2G17290
AT2G17420
AT2G17500
AT2G17705
AT2G17720
AT2G17730
AT2G17760
AT2G17840
AT2G17900
AT2G18000
AT2G18193
AT2G18660
AT2G18690
AT2G18950
AT2G18960
AT2G19570
AT2G19690
AT2G19710
AT2G20110
mRNA level ratio: log2(Pto AvrRpt2/mock)
Time (hpi)

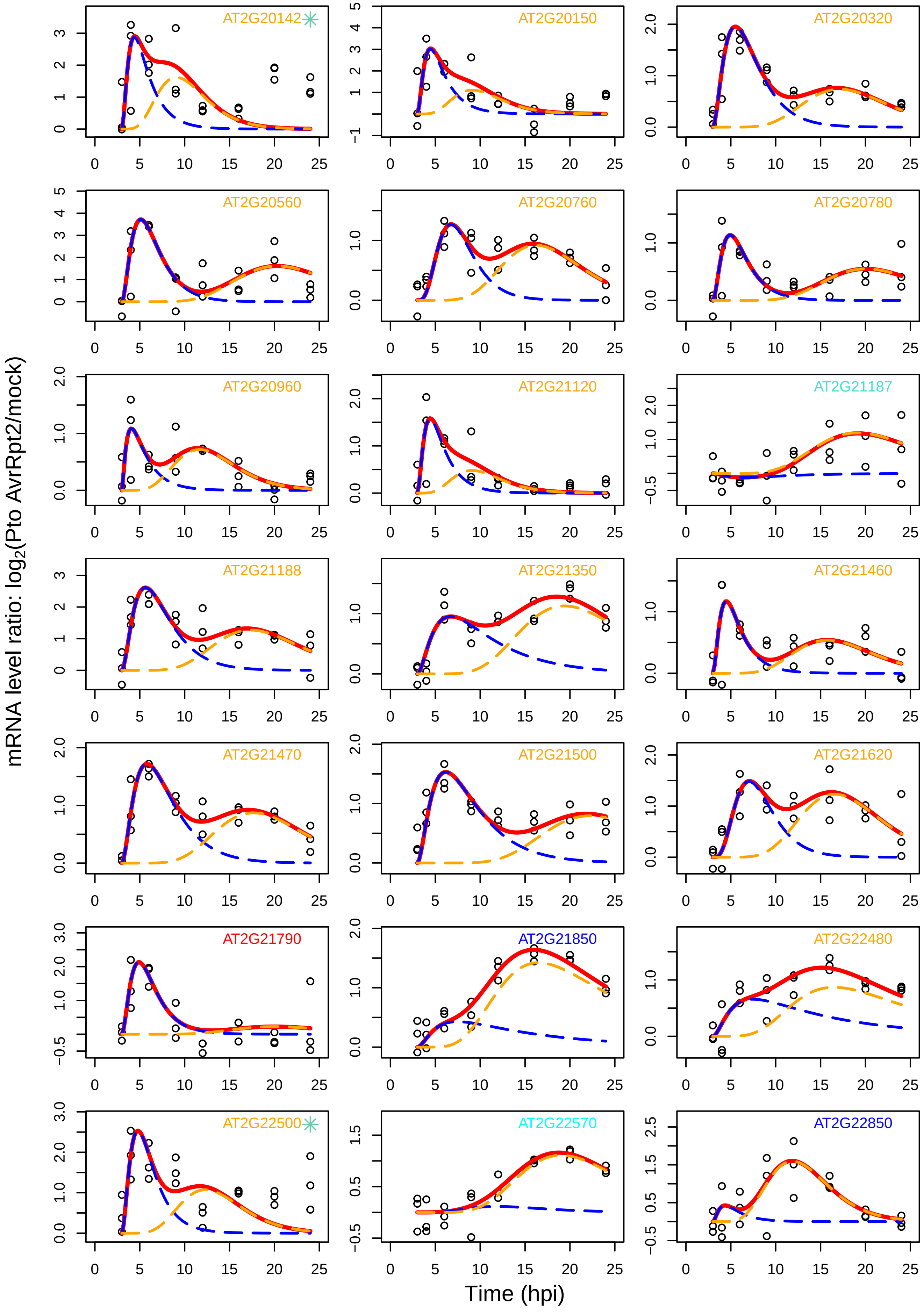

AT2G20142
AT2G20150
AT2G20320
AT2G20560
AT2G20760
AT2G20780
AT2G20960
AT2G21120
AT2G21187
AT2G21188
AT2G21350
AT2G21460
AT2G21470
AT2G21500
AT2G21620
AT2G21790
AT2G21850
AT2G22480
AT2G22500
AT2G22570
AT2G22850
mRNA level ratio: log2(Pto AvrRpt2/mock)
Time (hpi)

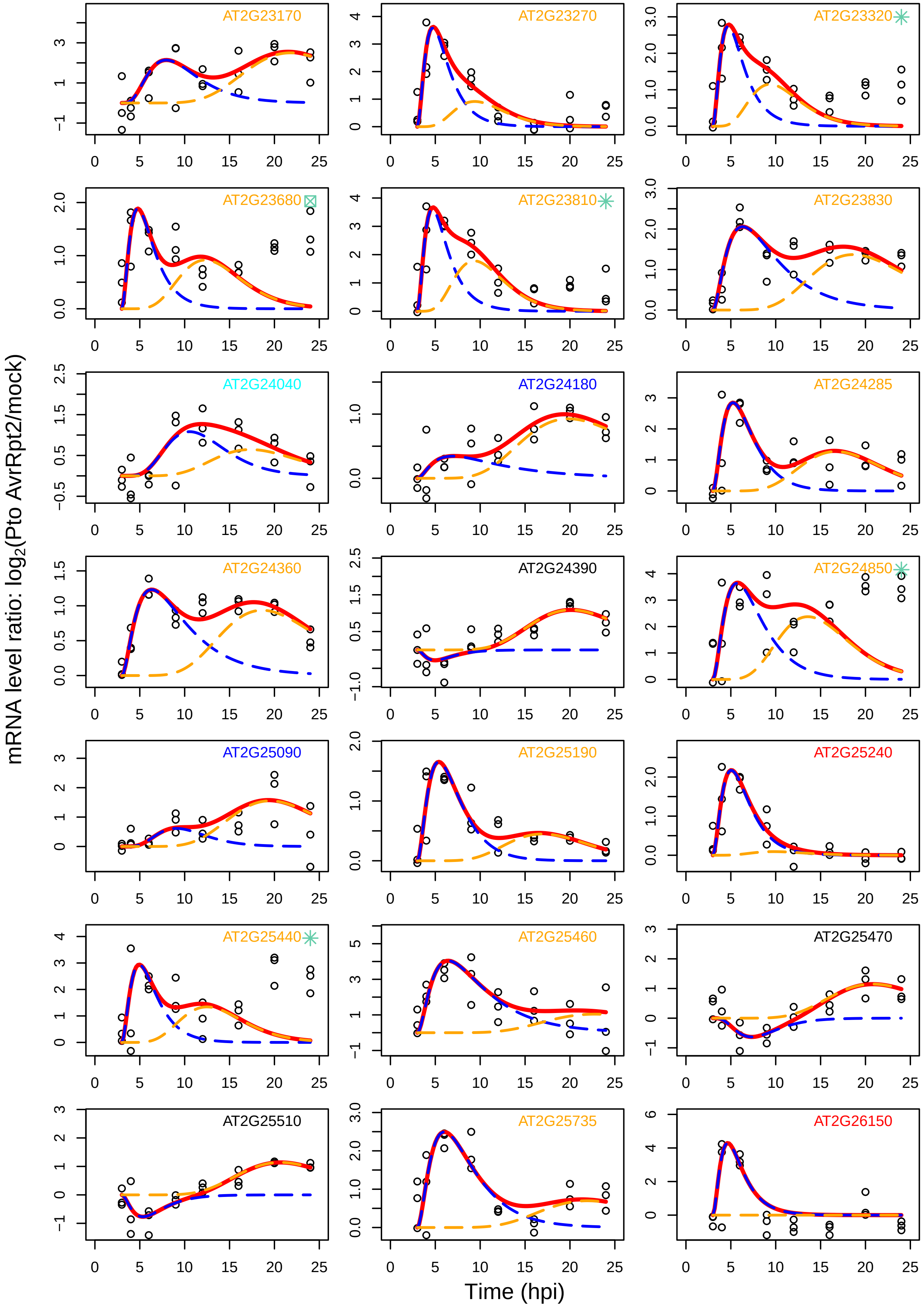
AT2G23170
AT2G23270
AT2G23320
AT2G23680
AT2G23810
AT2G23830
AT2G24040
AT2G24180
AT2G24285
AT2G24360
AT2G24390
AT2G24850
AT2G25090
AT2G25190
AT2G25240
AT2G25440
AT2G25460
AT2G25470
AT2G25510
AT2G25735
AT2G26150
mRNA level ratio: log2(Pto AvrRpt2/mock)
Time (hpi)

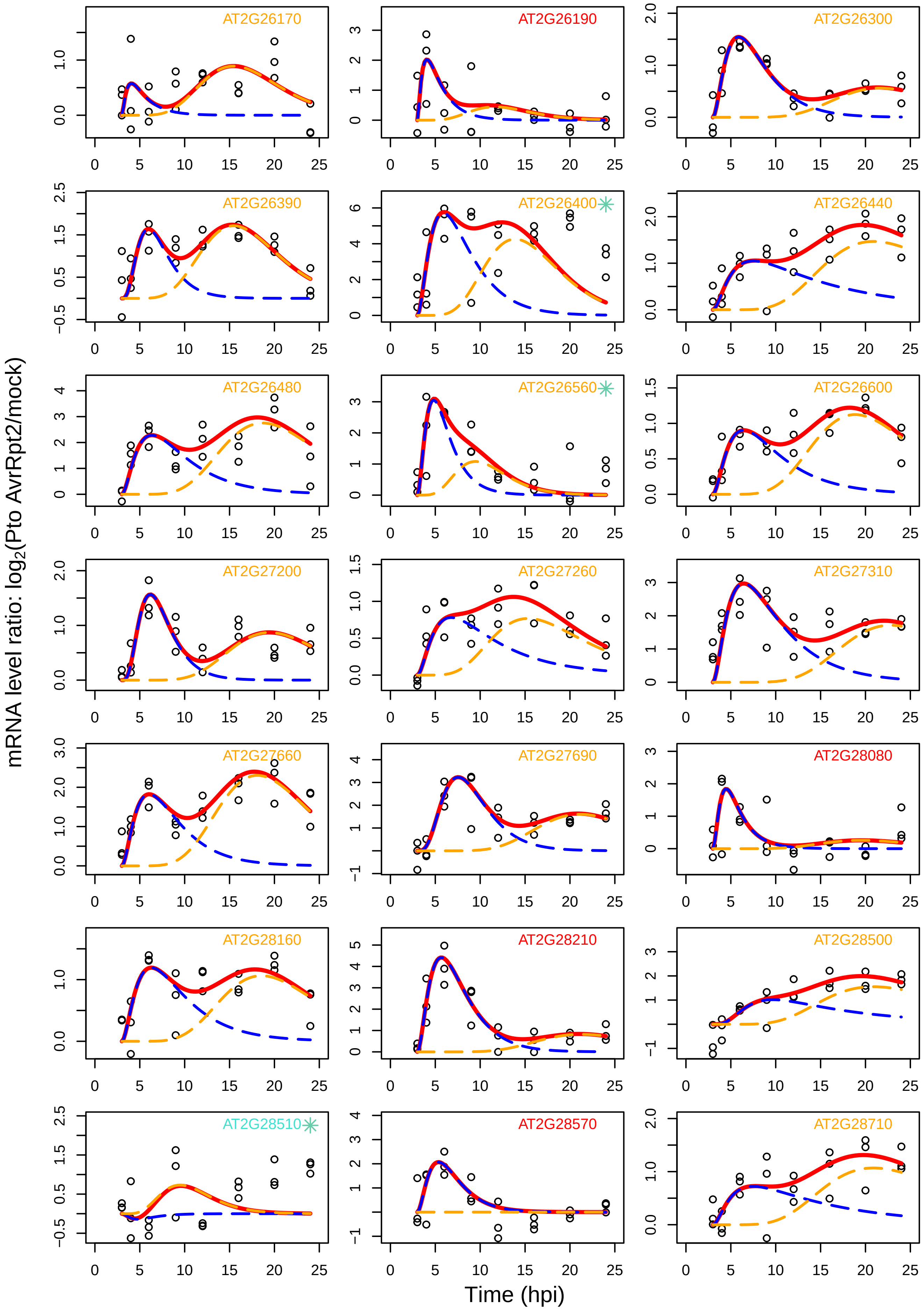
AT2G26170
AT2G26190
AT2G26300
AT2G26390
AT2G26400
AT2G26440
AT2G26480
AT2G26560
AT2G26600
AT2G27200
AT2G27260
AT2G27310
AT2G27660
AT2G27690
AT2G28080
AT2G28160
AT2G28210
AT2G28500
AT2G28510
AT2G28570
AT2G28710
mRNA level ratio: log2(Pto AvrRpt2/mock)
Time (hpi)

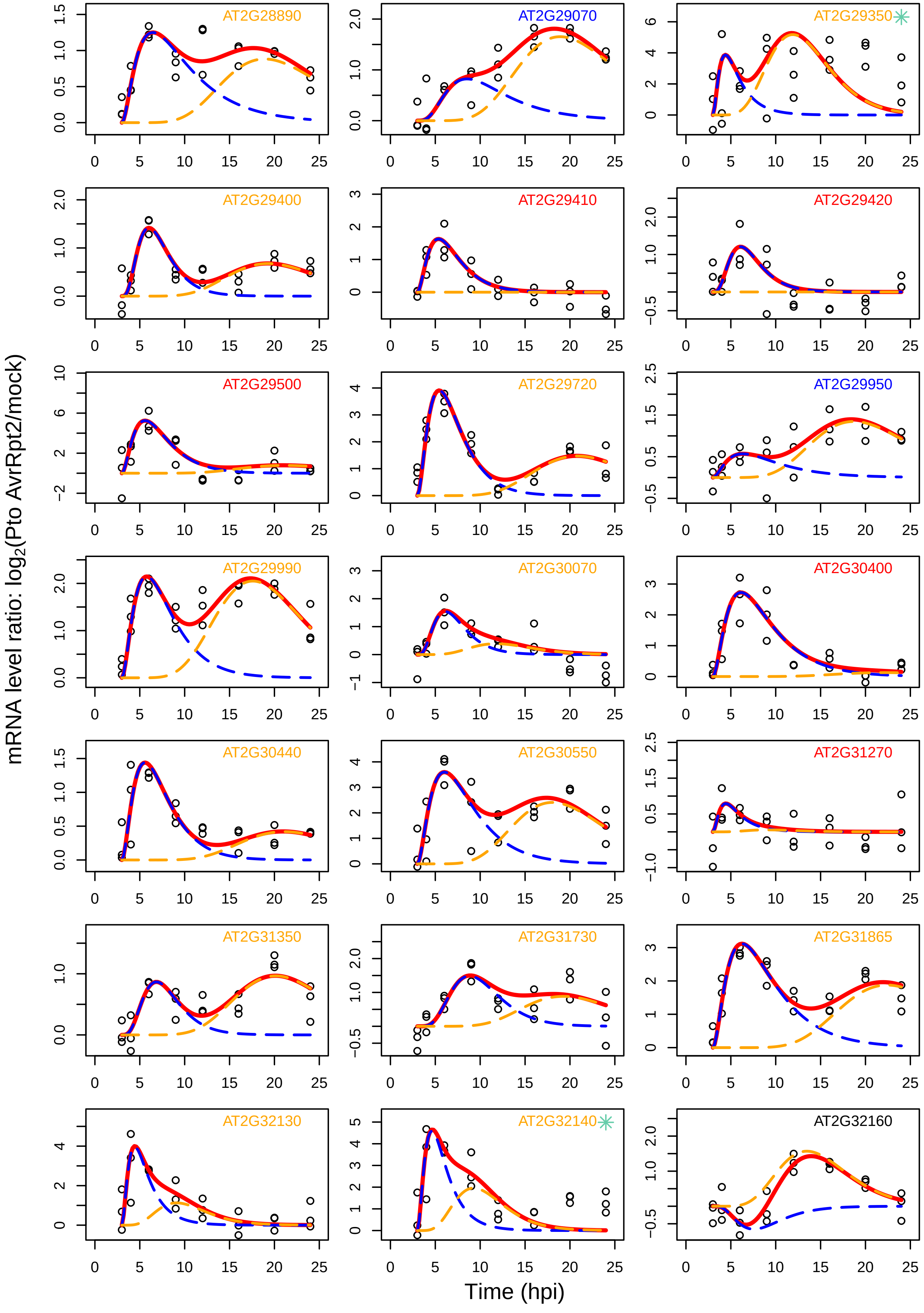
AT2G28890
AT2G29070
AT2G29350
AT2G29400
AT2G29410
AT2G29420
AT2G29500
AT2G29720
AT2G29950
AT2G29990
AT2G30070
AT2G30400
AT2G30440
AT2G30550
AT2G31270
AT2G31350
AT2G31730
AT2G31865
AT2G32130
AT2G32140
AT2G32160
mRNA level ratio: log2(Pto AvrRpt2/mock)
Time (hpi)

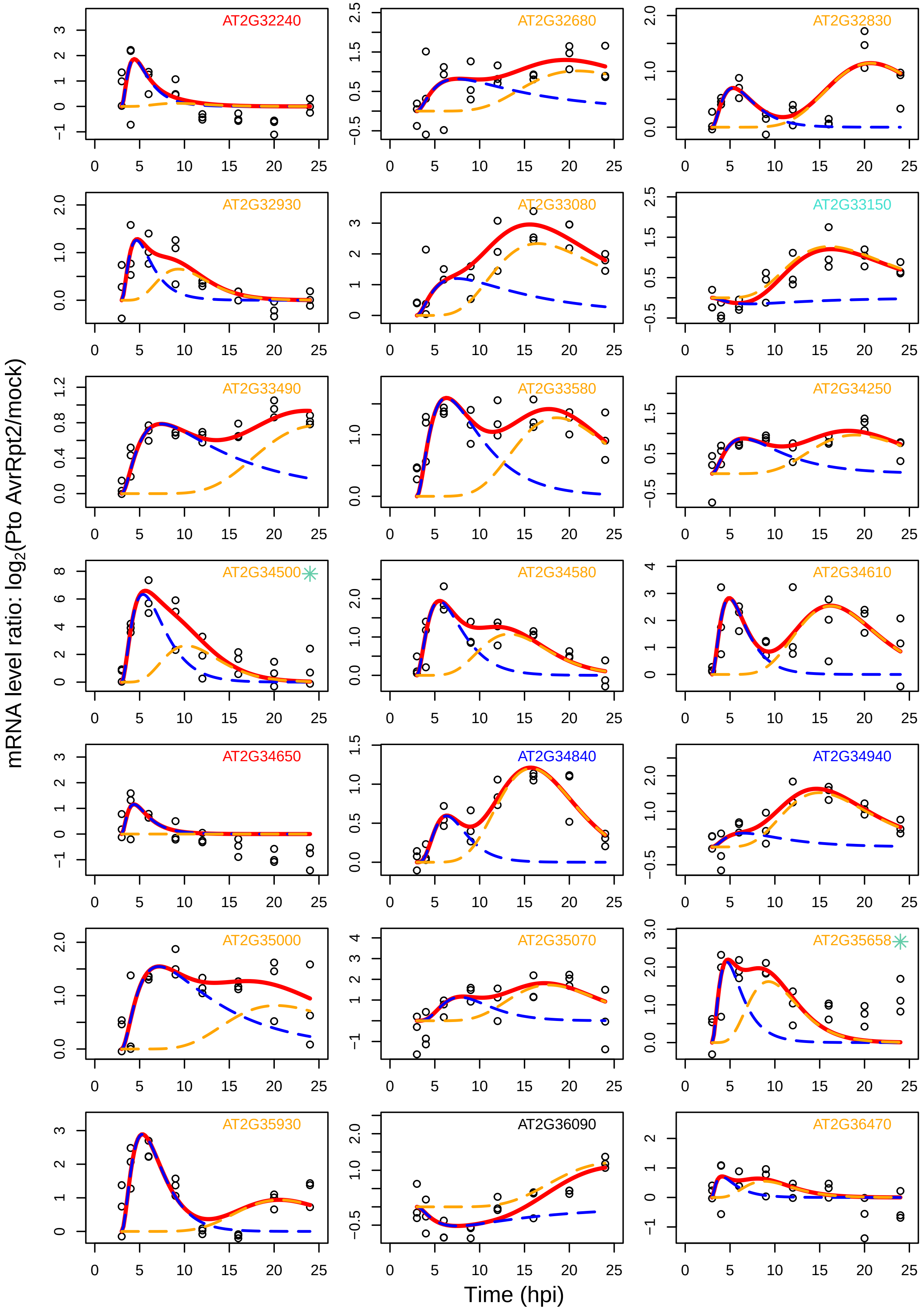

AT2G32240
AT2G32680
AT2G32830
AT2G32930
AT2G33080
AT2G33150
AT2G33490
AT2G33580
AT2G34250
AT2G34500
AT2G34580
AT2G34610
AT2G34650
AT2G34840
AT2G34940
AT2G35000
AT2G35070
AT2G35658
AT2G35930
AT2G36090
AT2G36470
mRNA level ratio: $log_2$(Pto AvrRpt2/mock)
Time (hpi)

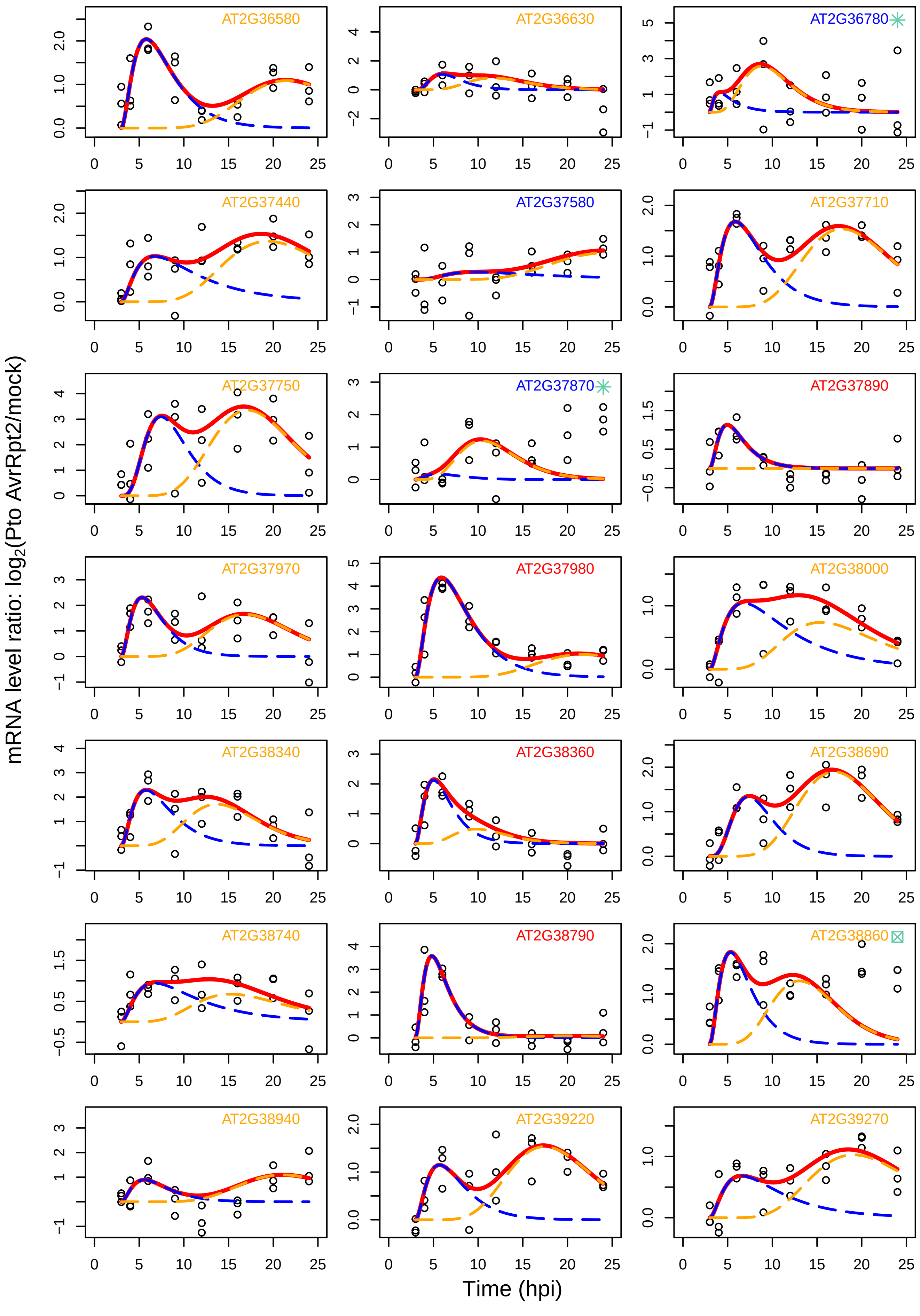
AT2G36580
AT2G36630
AT2G36780
AT2G37440
AT2G37580
AT2G37710
AT2G37750
AT2G37870
AT2G37890
AT2G37970
AT2G37980
AT2G38000
AT2G38340
AT2G38360
AT2G38690
AT2G38740
AT2G38790
AT2G38860
AT2G38940
AT2G39220
AT2G39270
mRNA level ratio: log2(Pto AvrRpt2/mock)
Time (hpi)

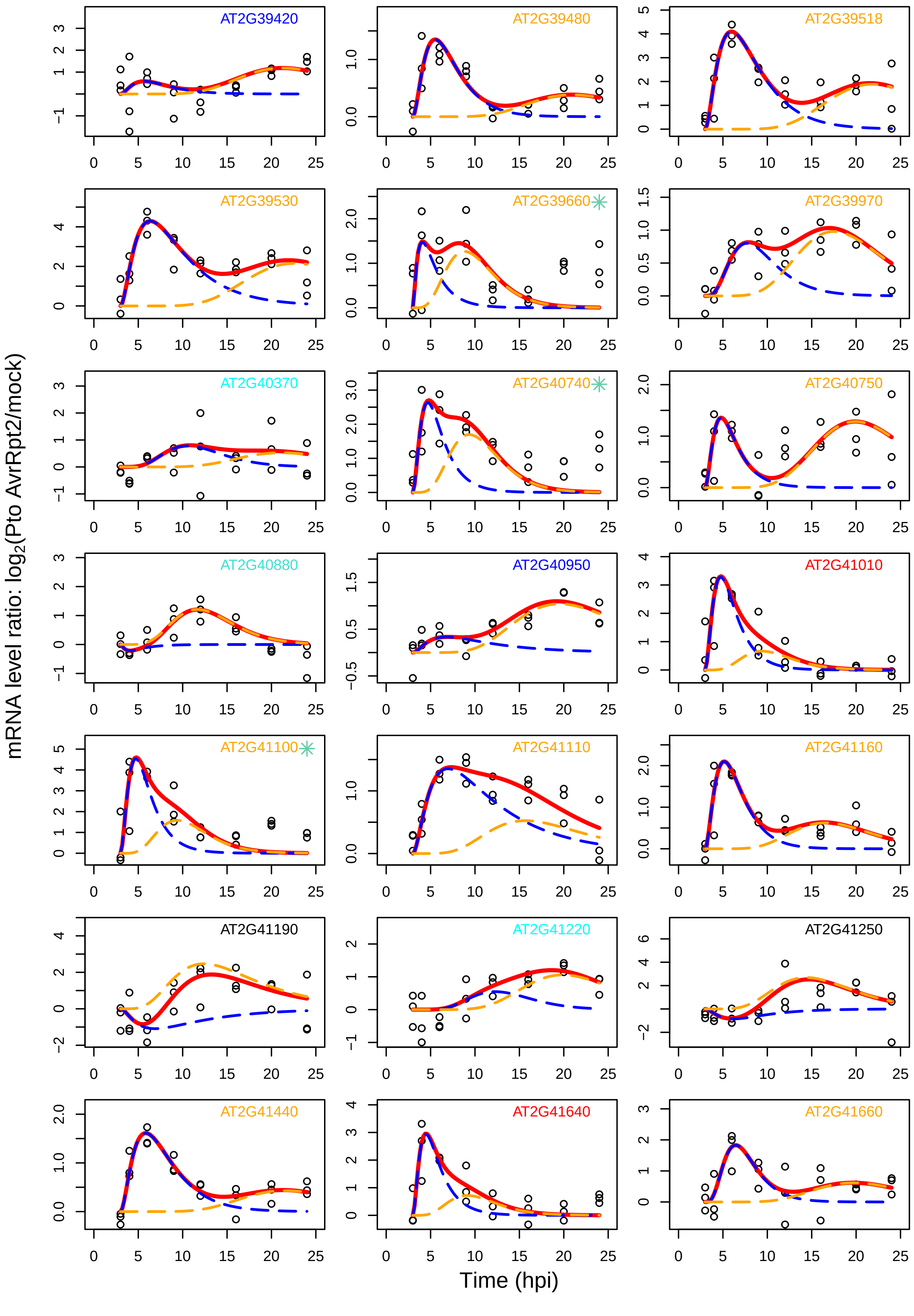
AT2G39420
AT2G39480
AT2G39518
AT2G39530
AT2G39660
AT2G39970
AT2G40370
AT2G40740
AT2G40750
AT2G40880
AT2G40950
AT2G41010
AT2G41100
AT2G41110
AT2G41160
AT2G41190
AT2G41220
AT2G41250
AT2G41440
AT2G41640
AT2G41660
mRNA level ratio: log2(Pto AvrRpt2/mock)
Time (hpi)

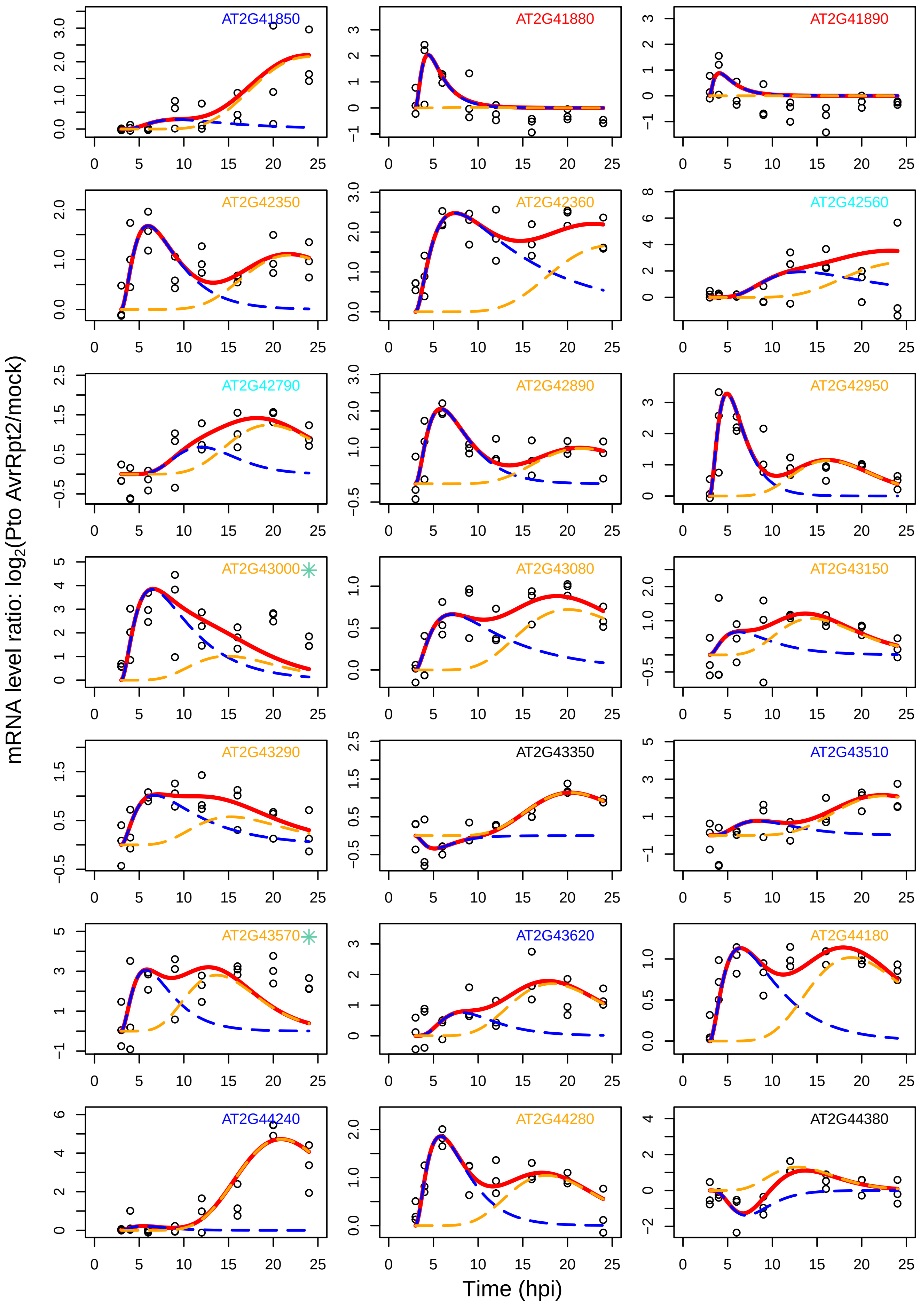
AT2G41850
AT2G41880
AT2G41890
AT2G42350
AT2G42360
AT2G42560
AT2G42790
AT2G42890
AT2G42950
AT2G43000
AT2G43080
AT2G43150
AT2G43290
AT2G43350
AT2G43510
AT2G43570
AT2G43620
AT2G44180
AT2G44240
AT2G44280
AT2G44380
mRNA level ratio: log2(Pto AvrRpt2/mock)
Time (hpi)

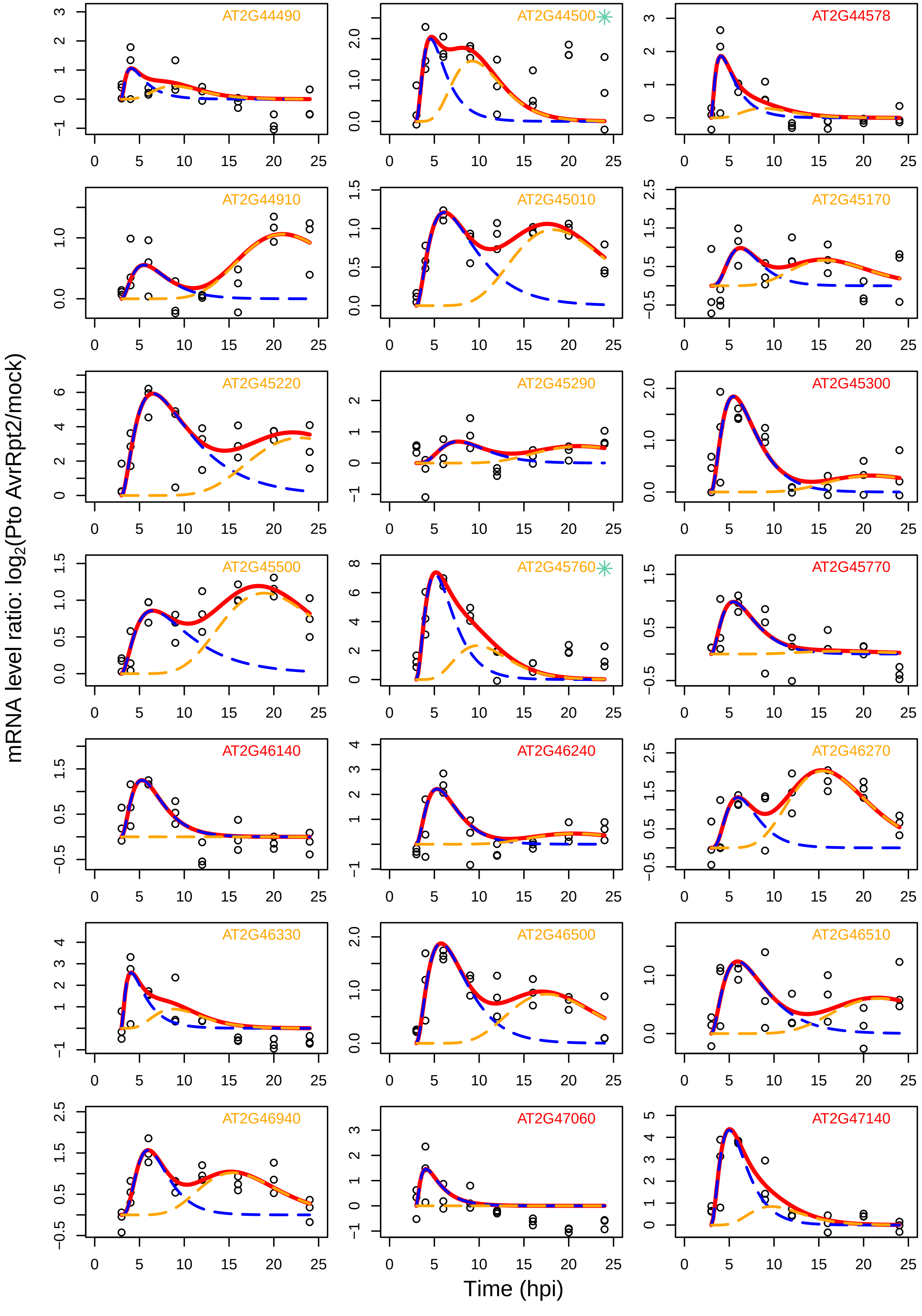

mRNA level ratio: log2(Pto AvrRpt2/mock)
Time (hpi)
AT2G44490
AT2G44500
AT2G44578
AT2G44910
AT2G45010
AT2G45170
AT2G45220
AT2G45290
AT2G45300
AT2G45500
AT2G45760
AT2G45770
AT2G46140
AT2G46240
AT2G46270
AT2G46330
AT2G46500
AT2G46510
AT2G46940
AT2G47060
AT2G47140

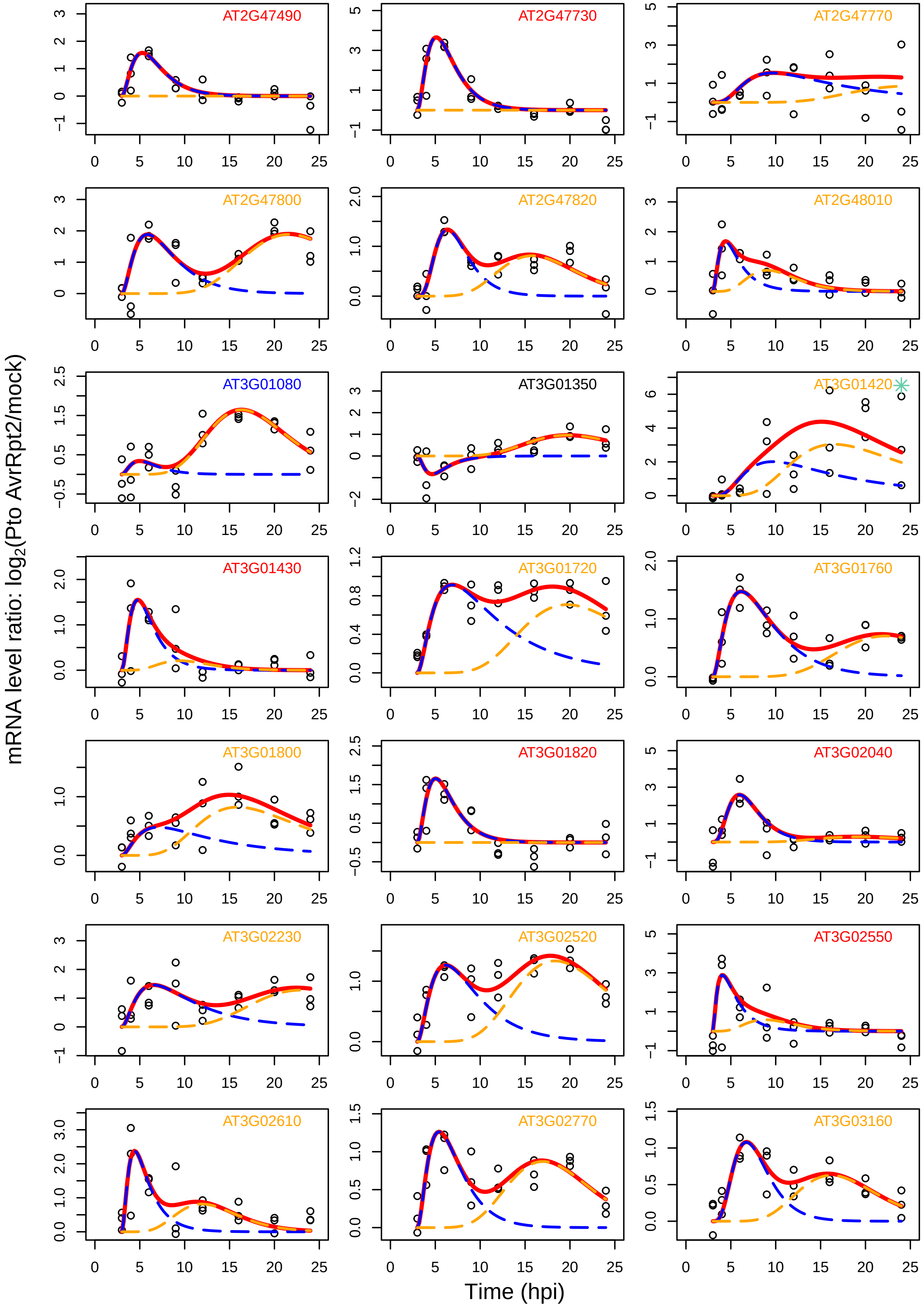

mRNA level ratio: $log_2$(Pto AvrRpt2/mock)
Time (hpi)
AT2G47490
AT2G47730
AT2G47770
AT2G47800
AT2G47820
AT2G48010
AT3G01080
AT3G01350
AT3G01420
AT3G01430
AT3G01720
AT3G01760
AT3G01800
AT3G01820
AT3G02040
AT3G02230
AT3G02520
AT3G02550
AT3G02610
AT3G02770
AT3G03160

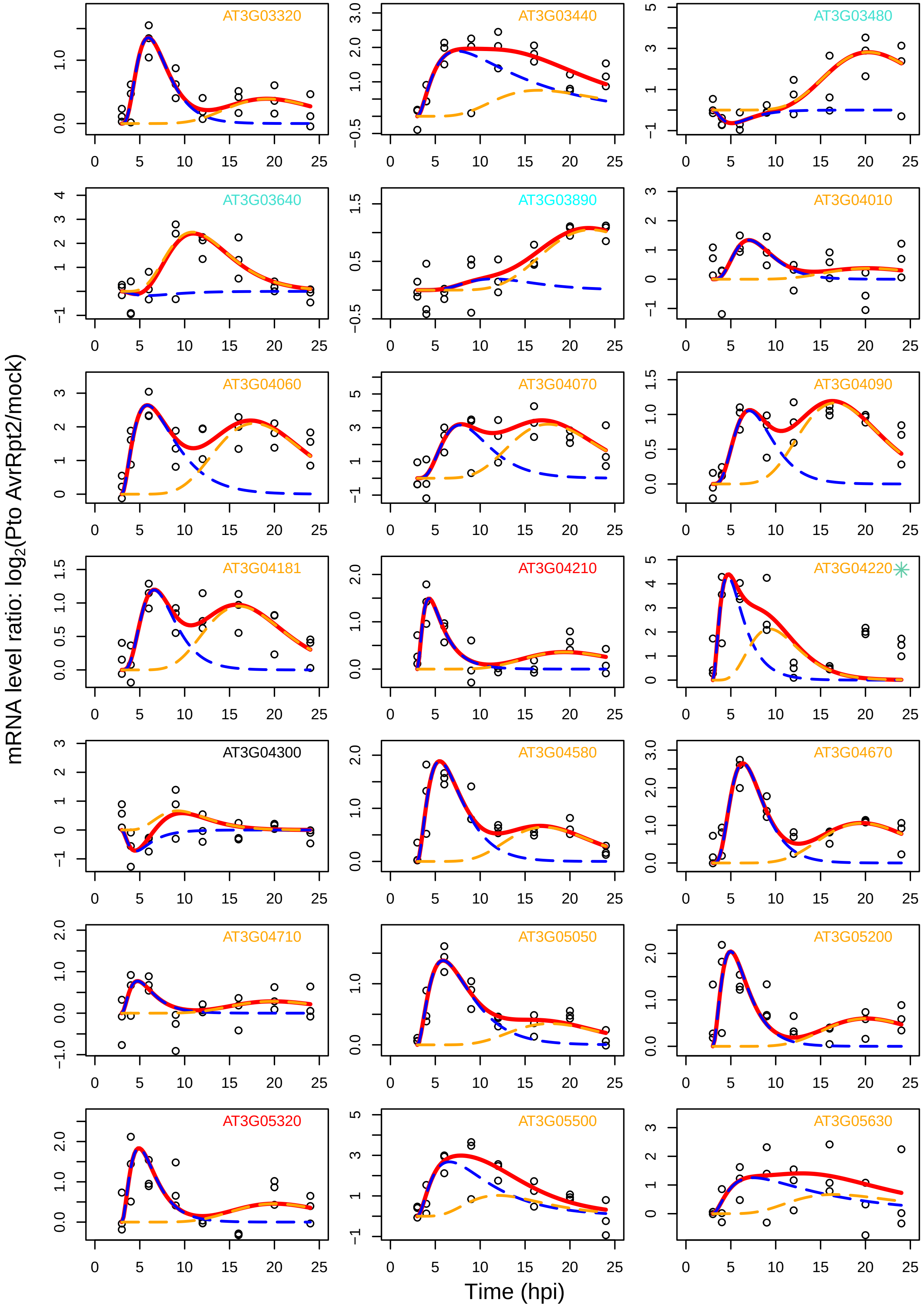

AT3G03320
AT3G03440
AT3G03480
AT3G03640
AT3G03890
AT3G04010
AT3G04060
AT3G04070
AT3G04090
AT3G04181
AT3G04210
AT3G04220
AT3G04300
AT3G04580
AT3G04670
AT3G04710
AT3G05050
AT3G05200
AT3G05320
AT3G05500
AT3G05630
mRNA level ratio: log2(Pto AvrRpt2/mock)
Time (hpi)

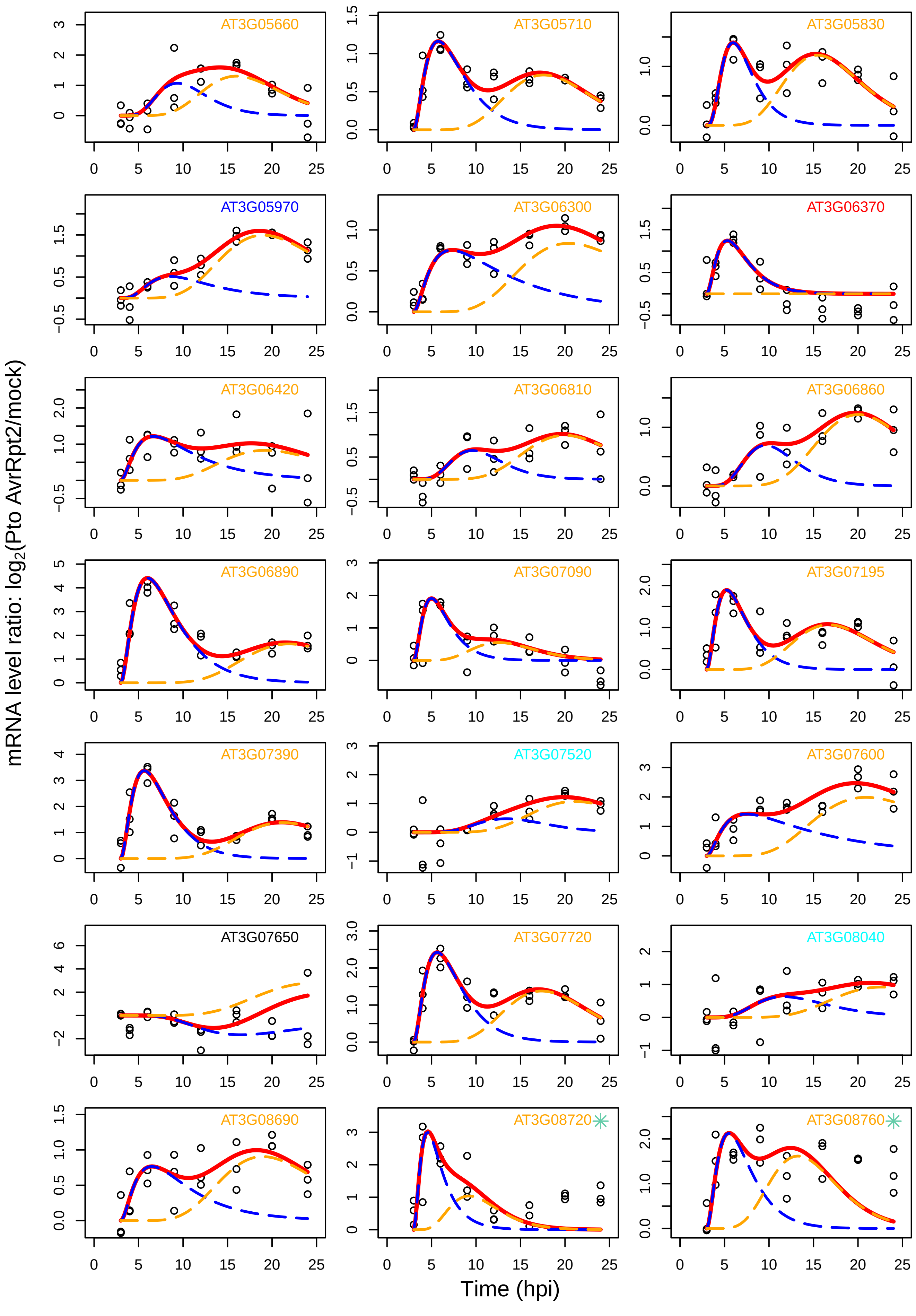

mRNA level ratio: log2(Pto AvrRpt2/mock)
Time (hpi)
AT3G05660
AT3G05710
AT3G05830
AT3G05970
AT3G06300
AT3G06370
AT3G06420
AT3G06810
AT3G06860
AT3G06890
AT3G07090
AT3G07195
AT3G07390
AT3G07520
AT3G07600
AT3G07650
AT3G07720
AT3G08040
AT3G08690
AT3G08720
AT3G08760

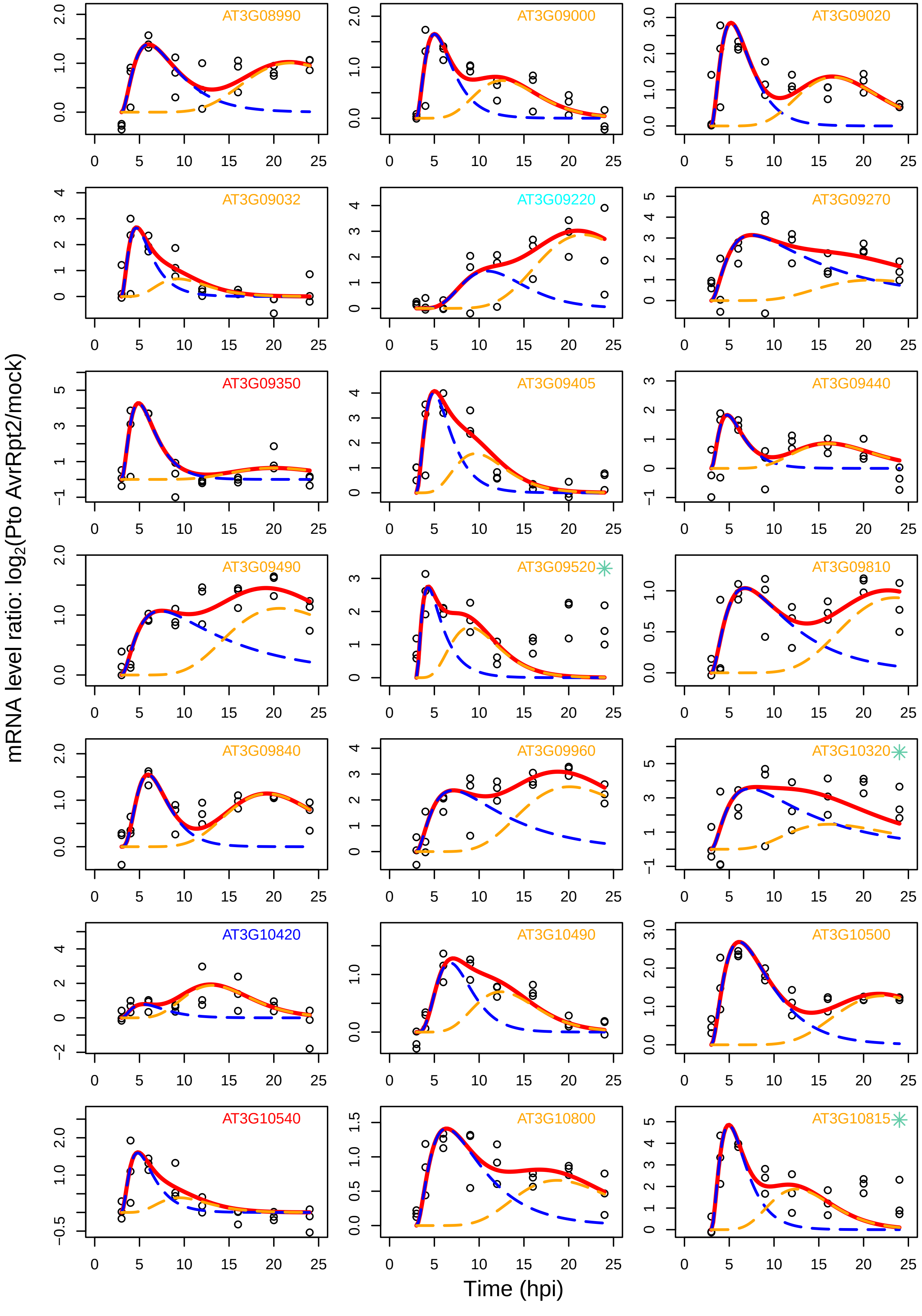

mRNA level ratio: log2(Pto AvrRpt2/mock)
Time (hpi)
AT3G08990
AT3G09000
AT3G09020
AT3G09032
AT3G09220
AT3G09270
AT3G09350
AT3G09405
AT3G09440
AT3G09490
AT3G09520
AT3G09810
AT3G09840
AT3G09960
AT3G10320
AT3G10420
AT3G10490
AT3G10500
AT3G10540
AT3G10800
AT3G10815

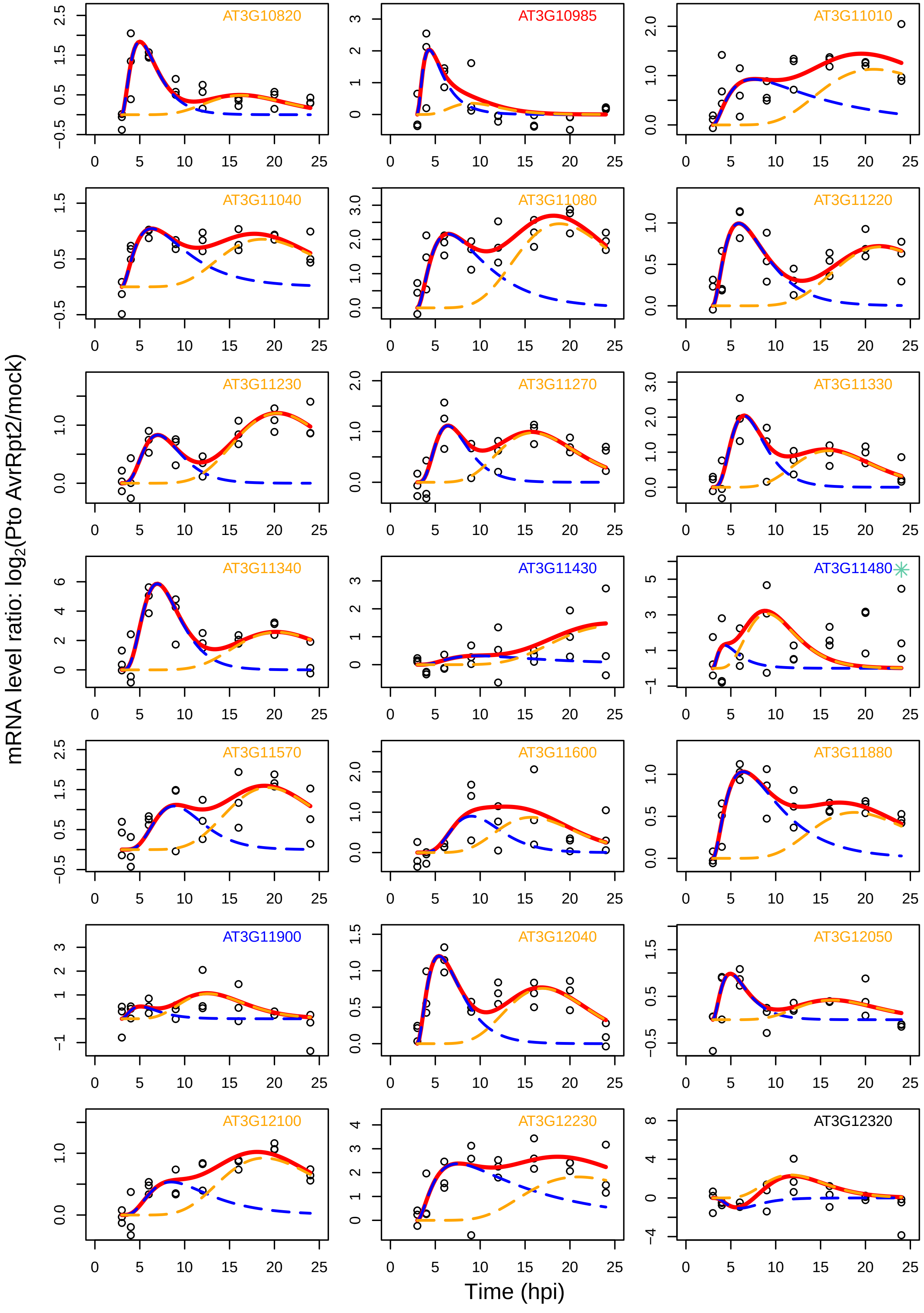
AT3G10820
AT3G10985
AT3G11010
AT3G11040
AT3G11080
AT3G11220
AT3G11230
AT3G11270
AT3G11330
AT3G11340
AT3G11430
AT3G11480
AT3G11570
AT3G11600
AT3G11880
AT3G11900
AT3G12040
AT3G12050
AT3G12100
AT3G12230
AT3G12320
mRNA level ratio: log$_2$(Pto AvrRpt2/mock)
Time (hpi)

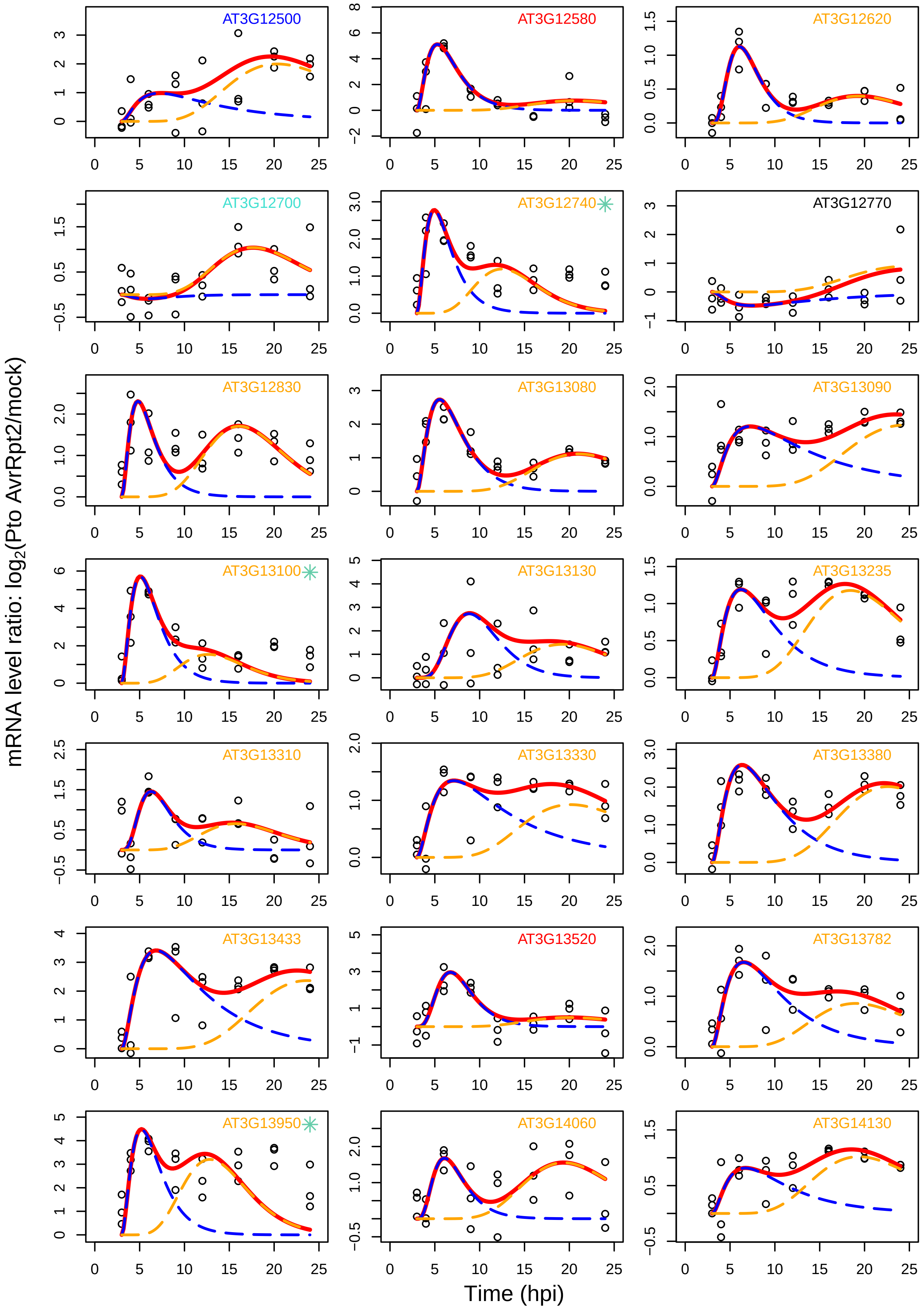
AT3G12500
AT3G12580
AT3G12620
AT3G12700
AT3G12740
AT3G12770
AT3G12830
AT3G13080
AT3G13090
AT3G13100
AT3G13130
AT3G13235
AT3G13310
AT3G13330
AT3G13380
AT3G13433
AT3G13520
AT3G13782
AT3G13950
AT3G14060
AT3G14130
mRNA level ratio: log$_2$(Pto AvrRpt2/mock)
Time (hpi)

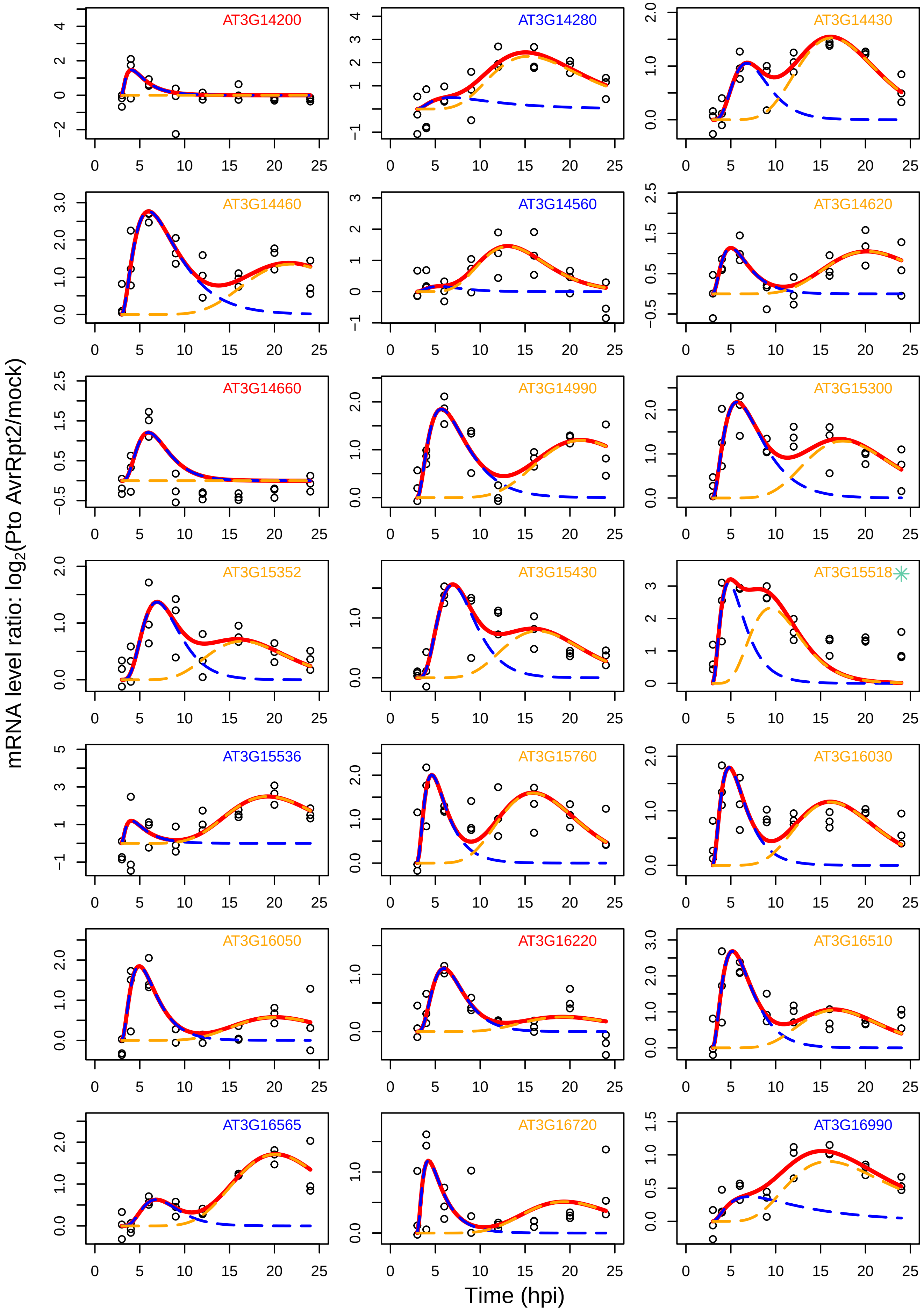

AT3G14200
AT3G14280
AT3G14430
AT3G14460
AT3G14560
AT3G14620
AT3G14660
AT3G14990
AT3G15300
AT3G15352
AT3G15430
AT3G15518
AT3G15536
AT3G15760
AT3G16030
AT3G16050
AT3G16220
AT3G16510
AT3G16565
AT3G16720
AT3G16990
mRNA level ratio: log2(Pto AvrRpt2/mock)
Time (hpi)

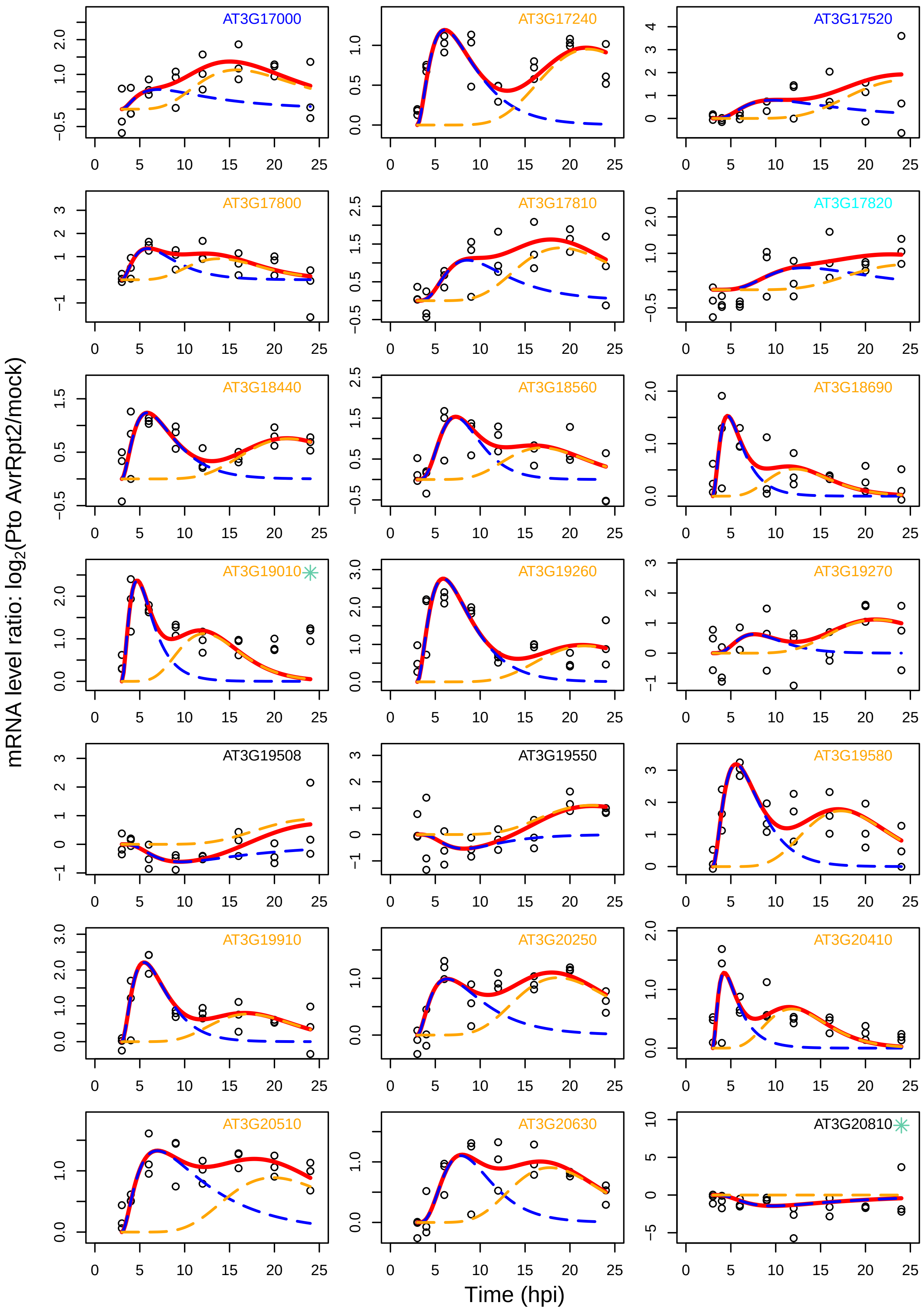
AT3G17000
AT3G17240
AT3G17520
AT3G17800
AT3G17810
AT3G17820
AT3G18440
AT3G18560
AT3G18690
AT3G19010
AT3G19260
AT3G19270
AT3G19508
AT3G19550
AT3G19580
AT3G19910
AT3G20250
AT3G20410
AT3G20510
AT3G20630
AT3G20810
mRNA level ratio: $\log_2$(Pto AvrRpt2/mock)
Time (hpi)

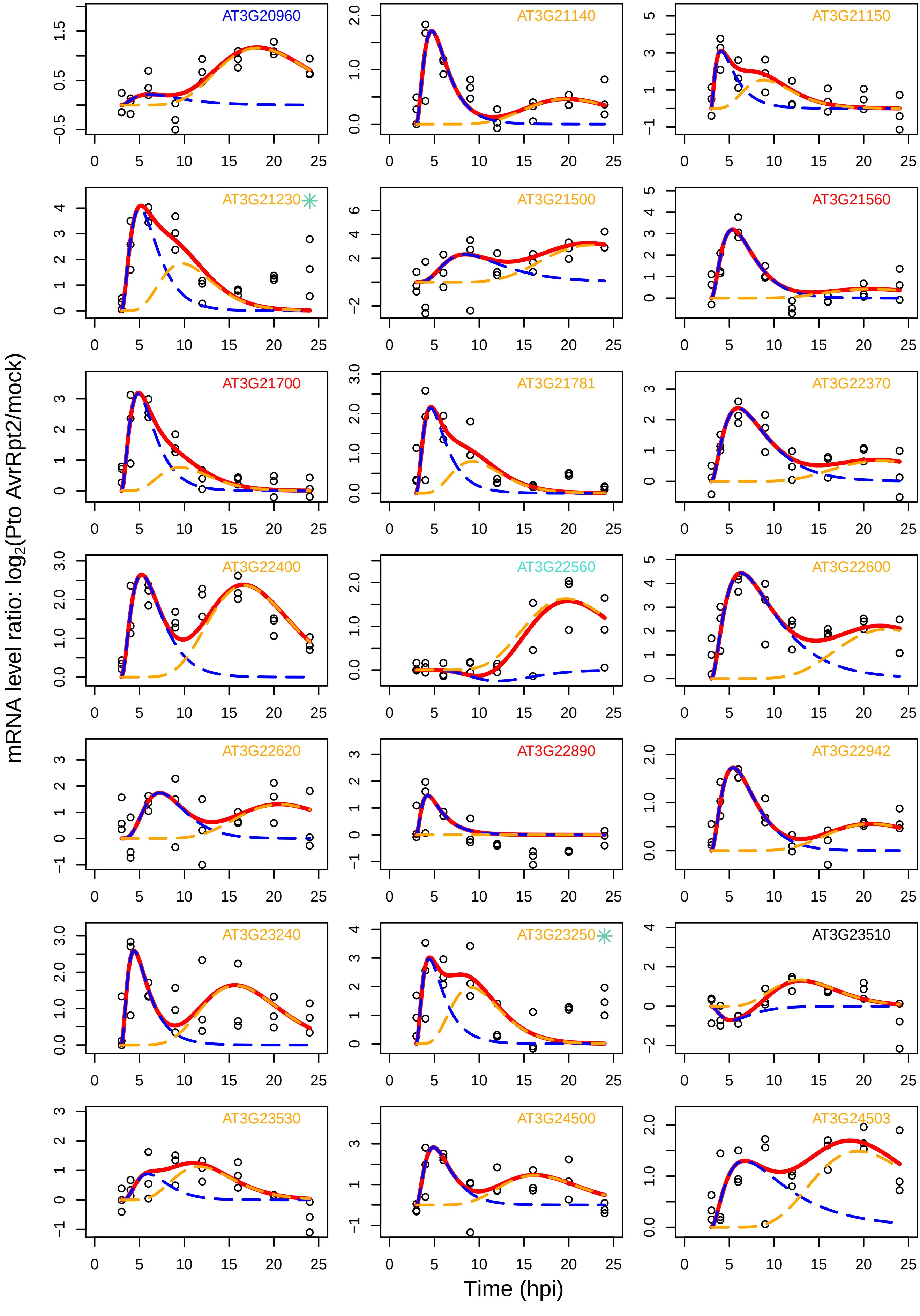

AT3G20960
AT3G21140
AT3G21150
AT3G21230
AT3G21500
AT3G21560
AT3G21700
AT3G21781
AT3G22370
AT3G22400
AT3G22560
AT3G22600
AT3G22620
AT3G22890
AT3G22942
AT3G23240
AT3G23250
AT3G23510
AT3G23530
AT3G24500
AT3G24503
mRNA level ratio: log2(Pto AvrRpt2/mock)
Time (hpi)

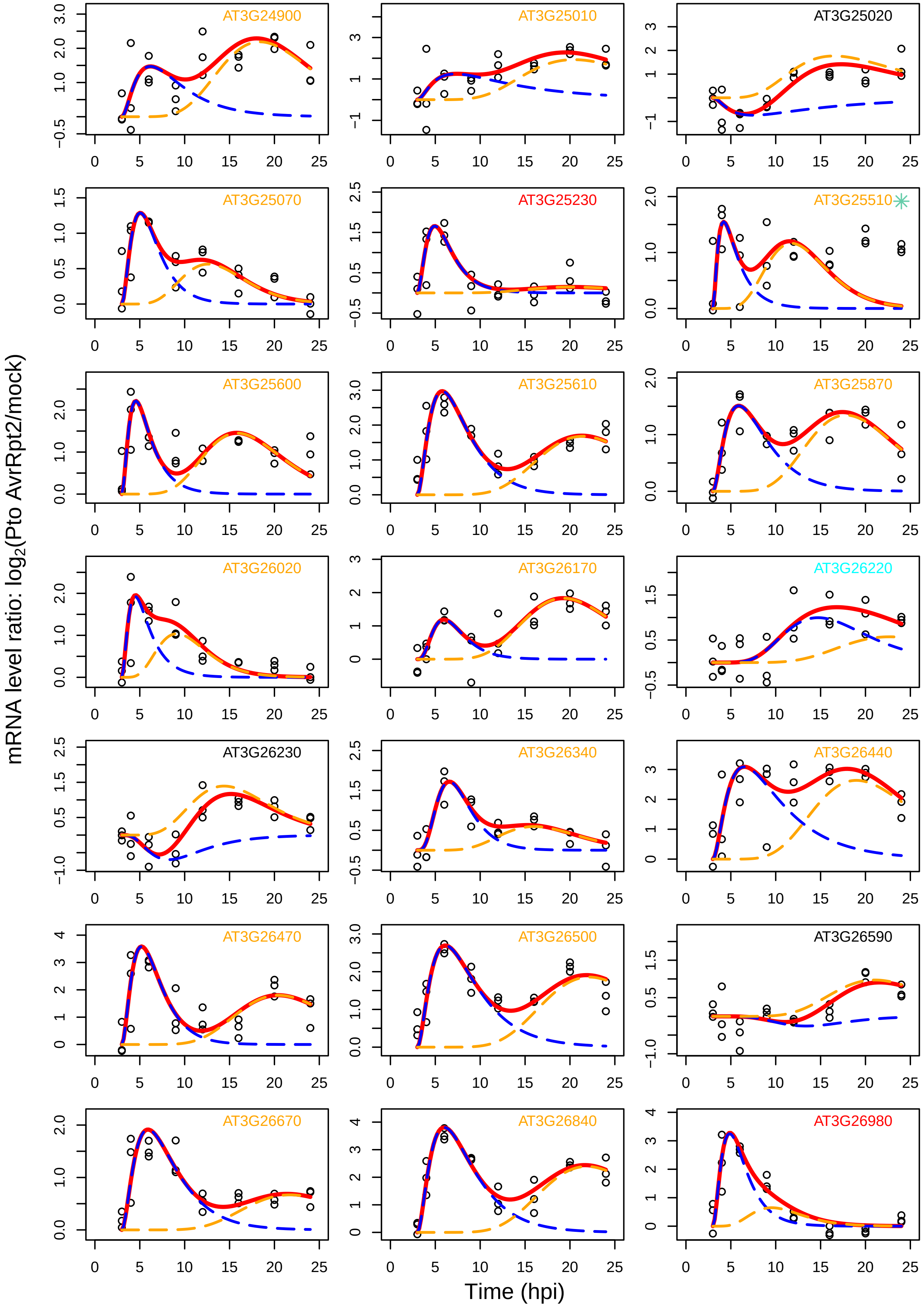

AT3G24900
AT3G25010
AT3G25020
AT3G25070
AT3G25230
AT3G25510
AT3G25600
AT3G25610
AT3G25870
AT3G26020
AT3G26170
AT3G26220
AT3G26230
AT3G26340
AT3G26440
AT3G26470
AT3G26500
AT3G26590
AT3G26670
AT3G26840
AT3G26980
mRNA level ratio: $log_2$(Pto AvrRpt2/mock)
Time (hpi)

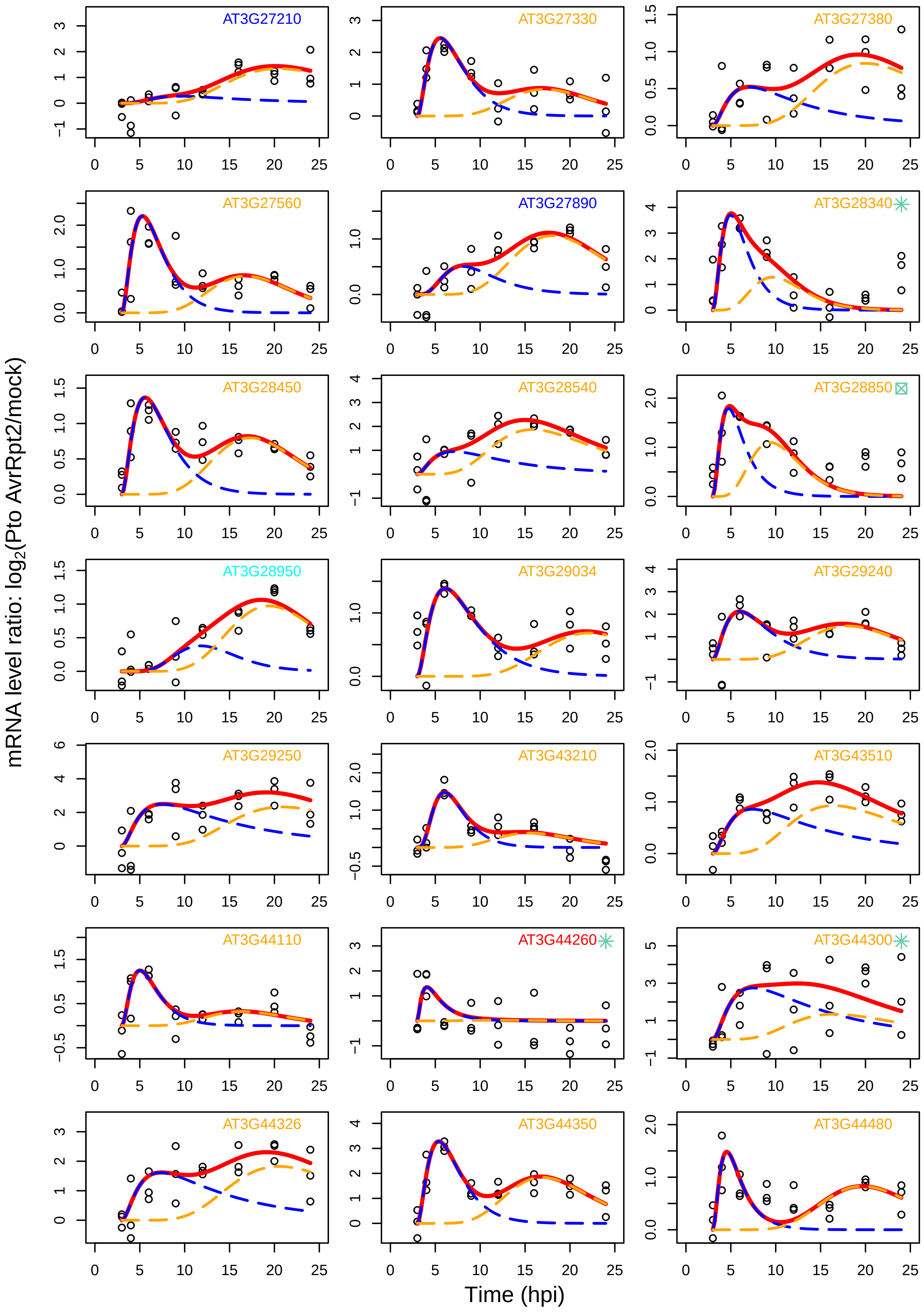

AT3G27210
AT3G27330
AT3G27380
AT3G27560
AT3G27890
AT3G28340
AT3G28450
AT3G28540
AT3G28850
AT3G28950
AT3G29034
AT3G29240
AT3G29250
AT3G43210
AT3G43510
AT3G44110
AT3G44260
AT3G44300
AT3G44326
AT3G44350
AT3G44480
mRNA level ratio: log2(Pto AvrRpt2/mock)
Time (hpi)

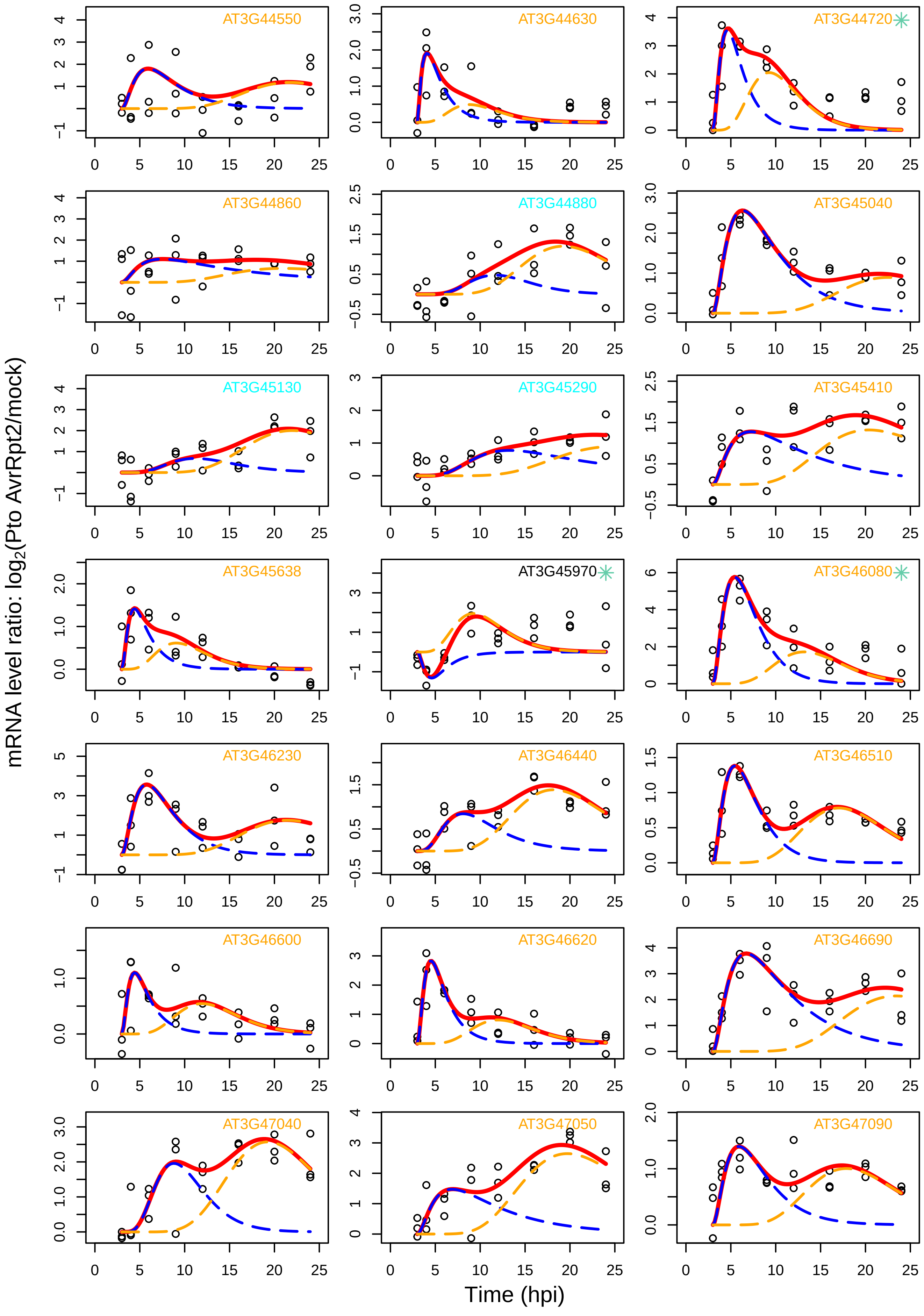
AT3G44550
AT3G44630
AT3G44720
AT3G44860
AT3G44880
AT3G45040
AT3G45130
AT3G45290
AT3G45410
AT3G45638
AT3G45970
AT3G46080
AT3G46230
AT3G46440
AT3G46510
AT3G46600
AT3G46620
AT3G46690
AT3G47040
AT3G47050
AT3G47090
mRNA level ratio: $log_2$(Pto AvrRpt2/mock)
Time (hpi)

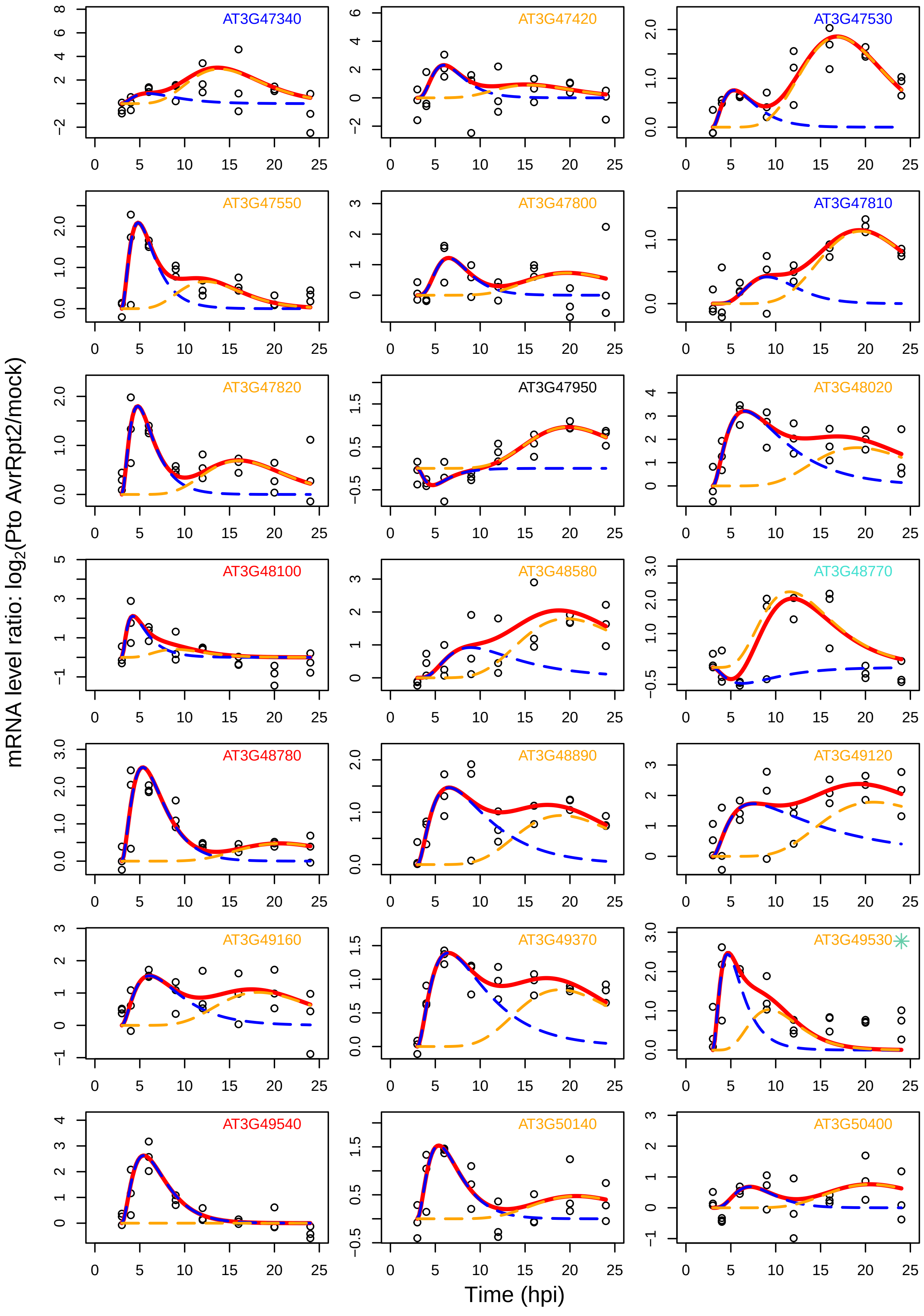

AT3G47340
AT3G47420
AT3G47530
AT3G47550
AT3G47800
AT3G47810
AT3G47820
AT3G47950
AT3G48020
AT3G48100
AT3G48580
AT3G48770
AT3G48780
AT3G48890
AT3G49120
AT3G49160
AT3G49370
AT3G49530
AT3G49540
AT3G50140
AT3G50400
mRNA level ratio: $log_2$(Pto AvrRpt2/mock)
Time (hpi)

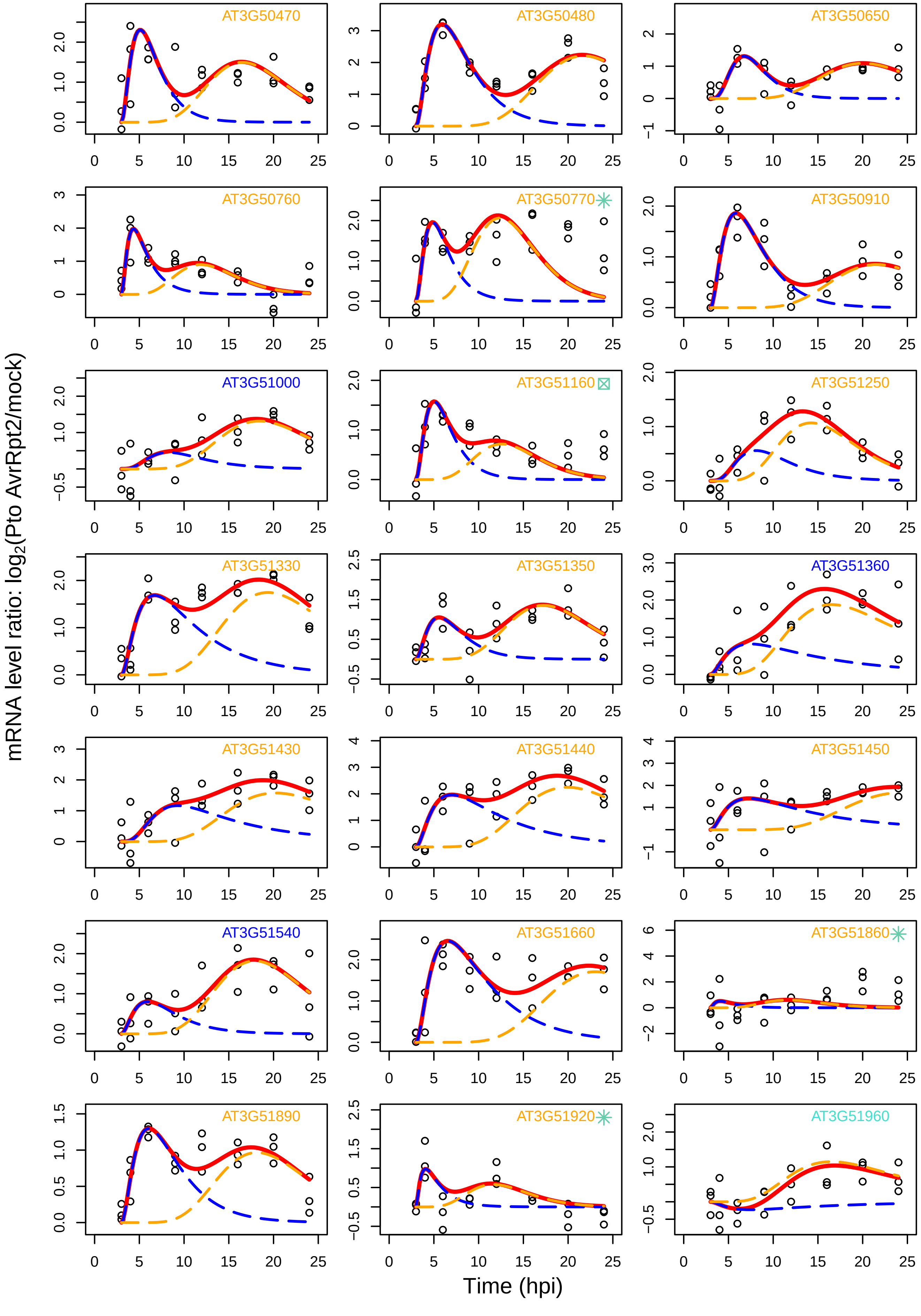
AT3G50470
AT3G50480
AT3G50650
AT3G50760
AT3G50770
AT3G50910
AT3G51000
AT3G51160
AT3G51250
AT3G51330
AT3G51350
AT3G51360
AT3G51430
AT3G51440
AT3G51450
AT3G51540
AT3G51660
AT3G51860
AT3G51890
AT3G51920
AT3G51960
mRNA level ratio: $log_2$(Pto AvrRpt2/mock)
Time (hpi)

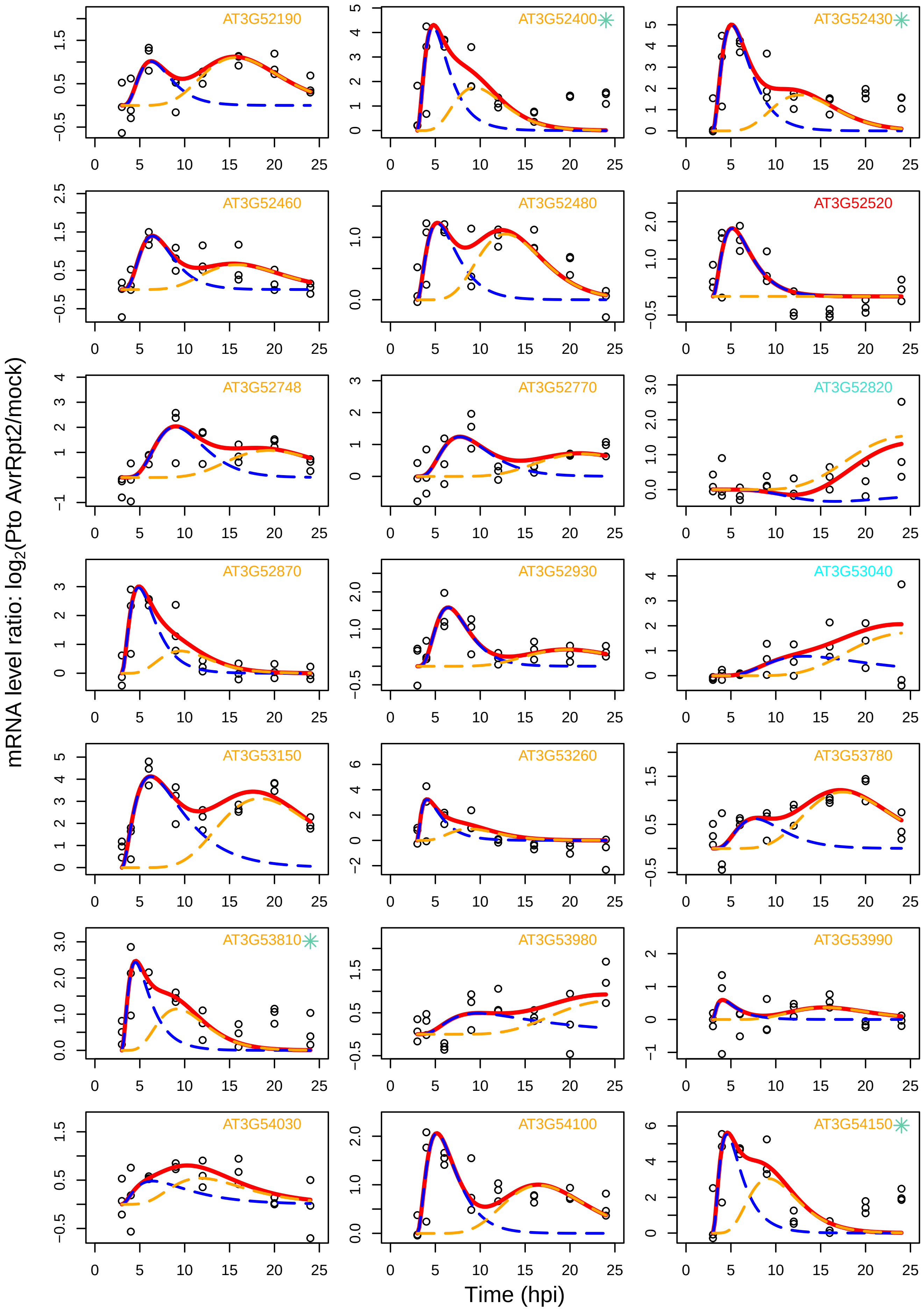

AT3G52190
AT3G52400
AT3G52430
AT3G52460
AT3G52480
AT3G52520
AT3G52748
AT3G52770
AT3G52820
AT3G52870
AT3G52930
AT3G53040
AT3G53150
AT3G53260
AT3G53780
AT3G53810
AT3G53980
AT3G53990
AT3G54030
AT3G54100
AT3G54150
mRNA level ratio: log2(Pto AvrRpt2/mock)
Time (hpi)

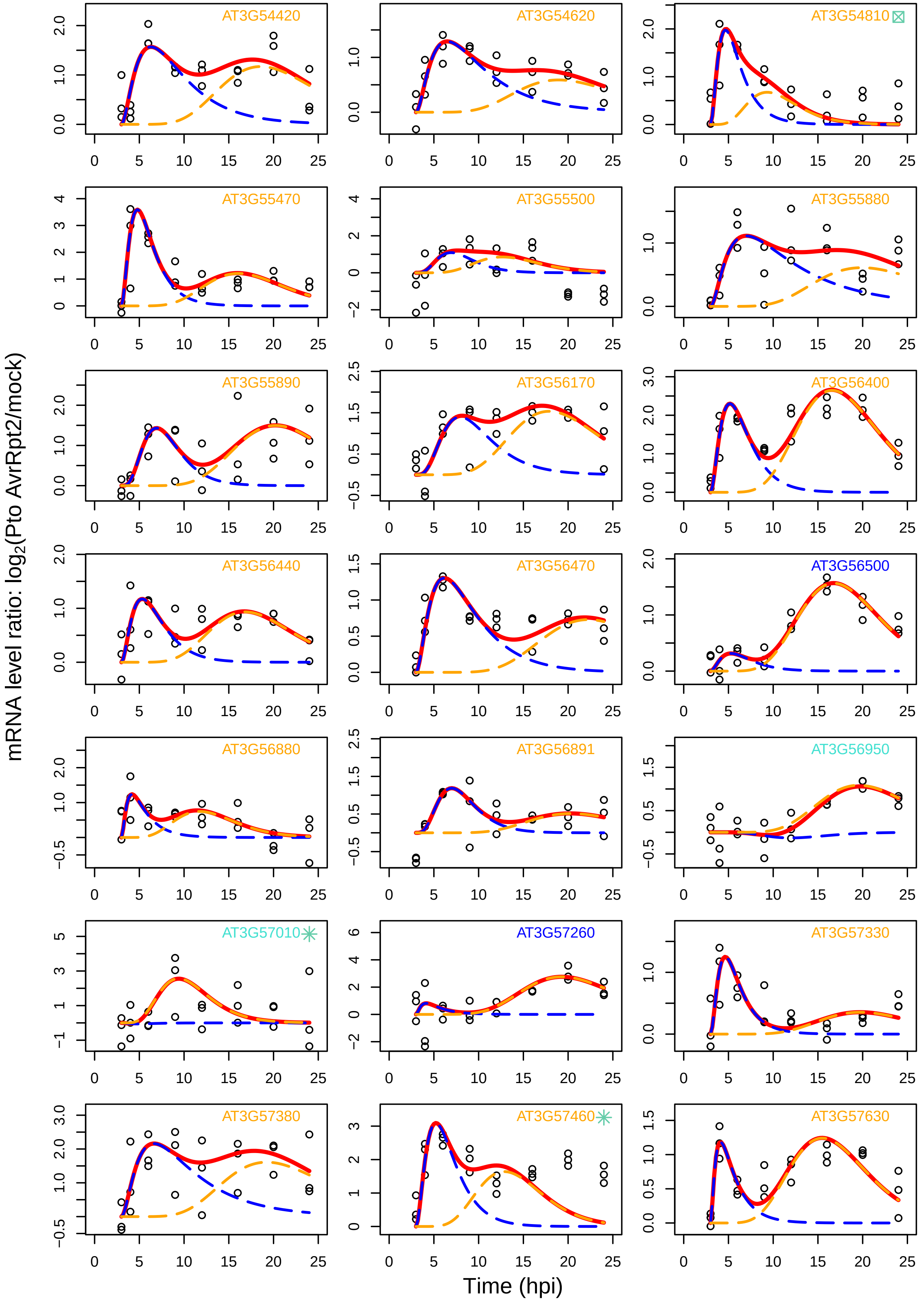

mRNA level ratio: $log_2$(Pto AvrRpt2/mock)
Time (hpi)
AT3G54420
AT3G54620
AT3G54810
AT3G55470
AT3G55500
AT3G55880
AT3G55890
AT3G56170
AT3G56400
AT3G56440
AT3G56470
AT3G56500
AT3G56880
AT3G56891
AT3G56950
AT3G57010
AT3G57260
AT3G57330
AT3G57380
AT3G57460
AT3G57630

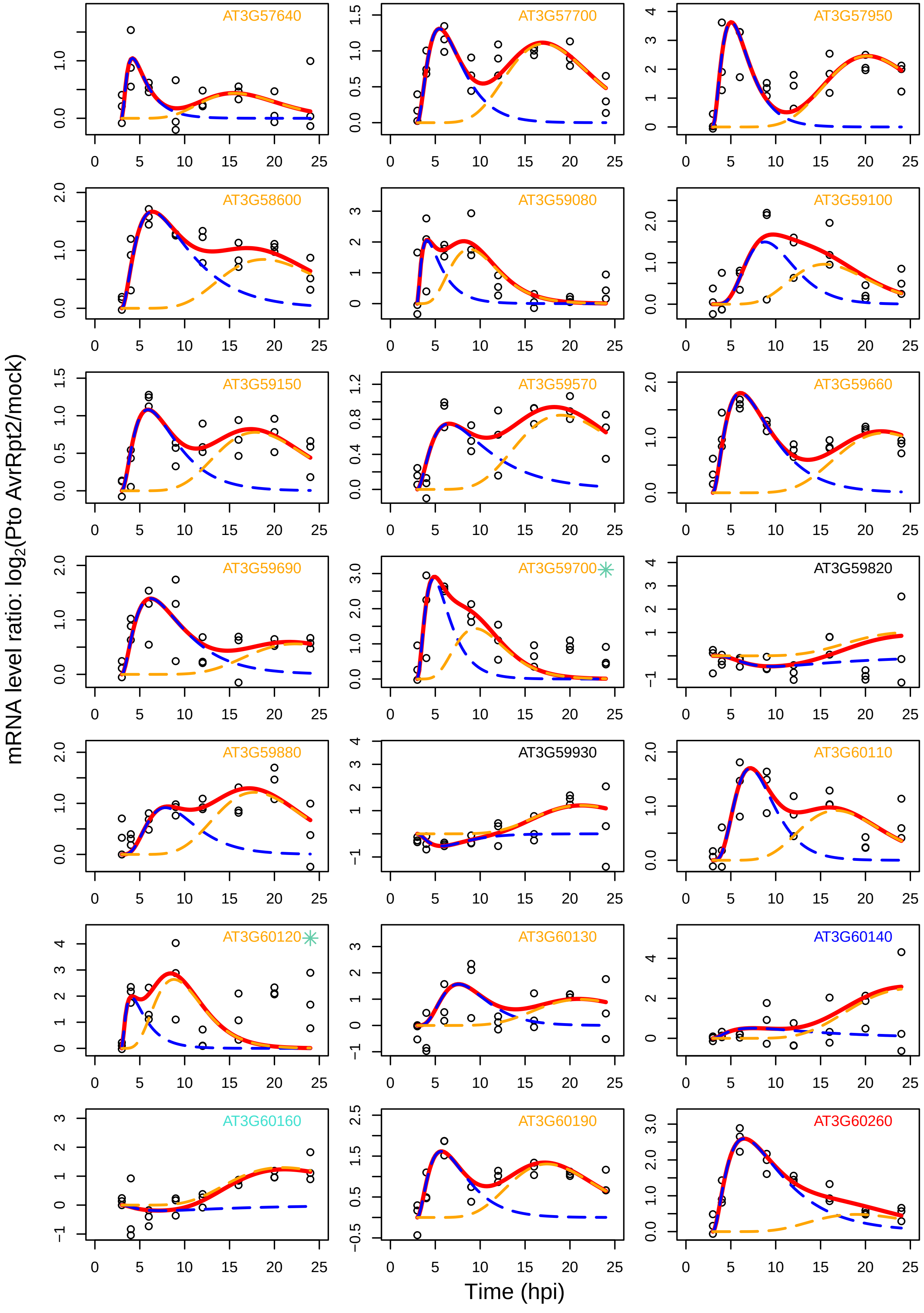

AT3G57640
AT3G57700
AT3G57950
AT3G58600
AT3G59080
AT3G59100
AT3G59150
AT3G59570
AT3G59660
AT3G59690
AT3G59700
AT3G59820
AT3G59880
AT3G59930
AT3G60110
AT3G60120
AT3G60130
AT3G60140
AT3G60160
AT3G60190
AT3G60260
mRNA level ratio: log2(Pto AvrRpt2/mock)
Time (hpi)

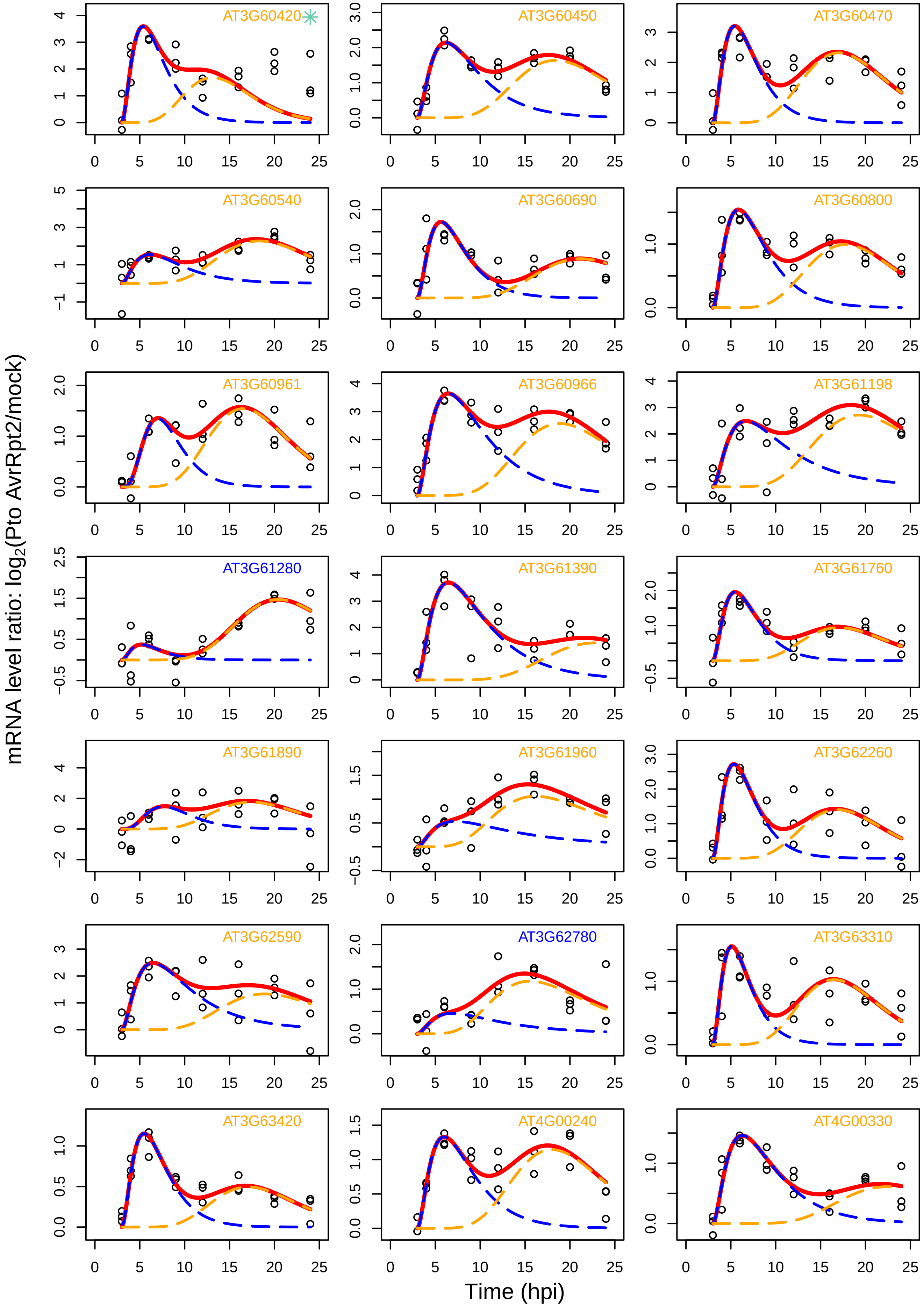

AT3G60420
AT3G60450
AT3G60470
AT3G60540
AT3G60690
AT3G60800
AT3G60961
AT3G60966
AT3G61198
AT3G61280
AT3G61390
AT3G61760
AT3G61890
AT3G61960
AT3G62260
AT3G62590
AT3G62780
AT3G63310
AT3G63420
AT4G00240
AT4G00330
mRNA level ratio: log2(Pto AvrRpt2/mock)
Time (hpi)

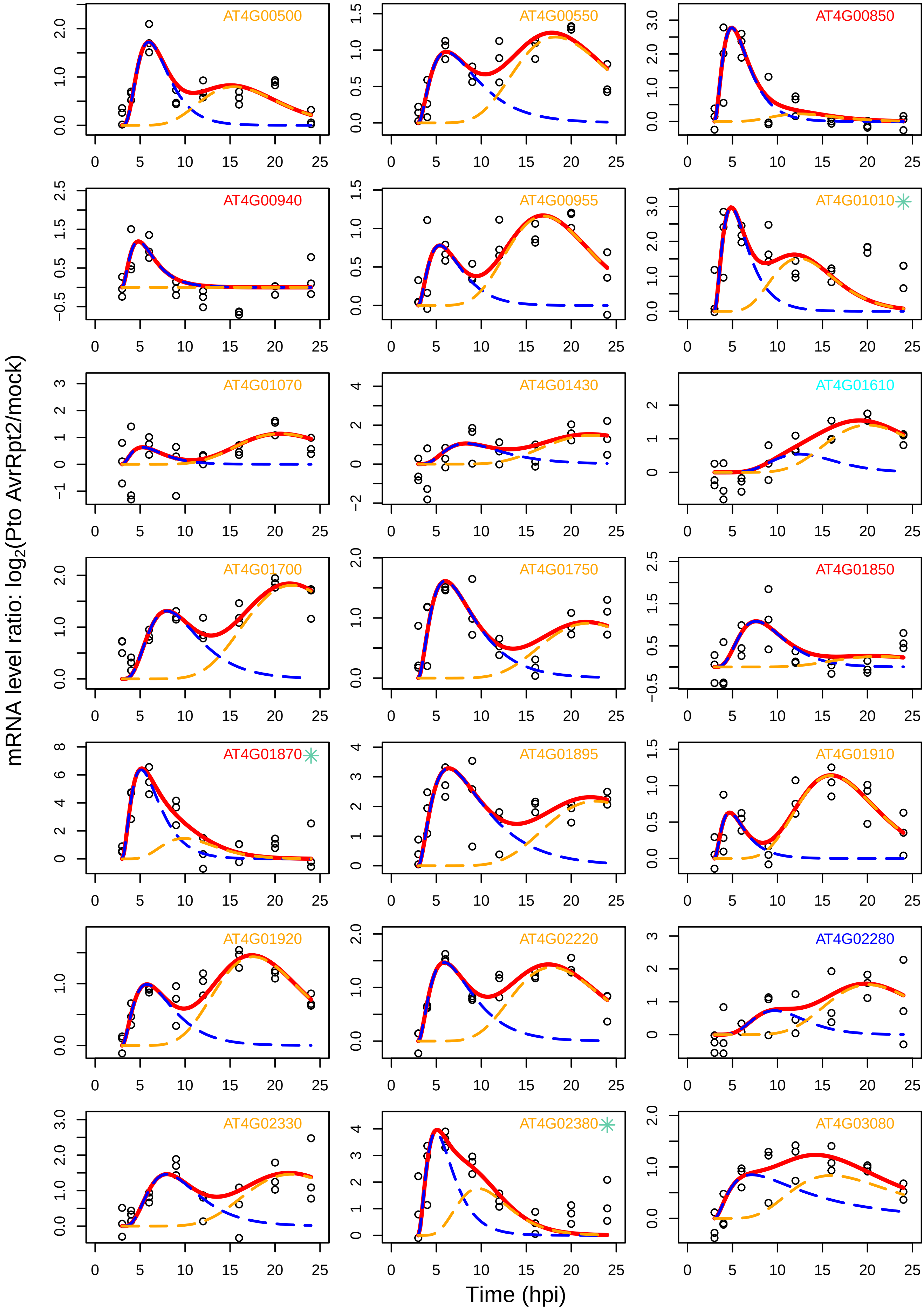

AT4G00500
AT4G00550
AT4G00850
AT4G00940
AT4G00955
AT4G01010
AT4G01070
AT4G01430
AT4G01610
AT4G01700
AT4G01750
AT4G01850
AT4G01870
AT4G01895
AT4G01910
AT4G01920
AT4G02220
AT4G02280
AT4G02330
AT4G02380
AT4G03080
mRNA level ratio: log2(Pto AvrRpt2/mock)
Time (hpi)

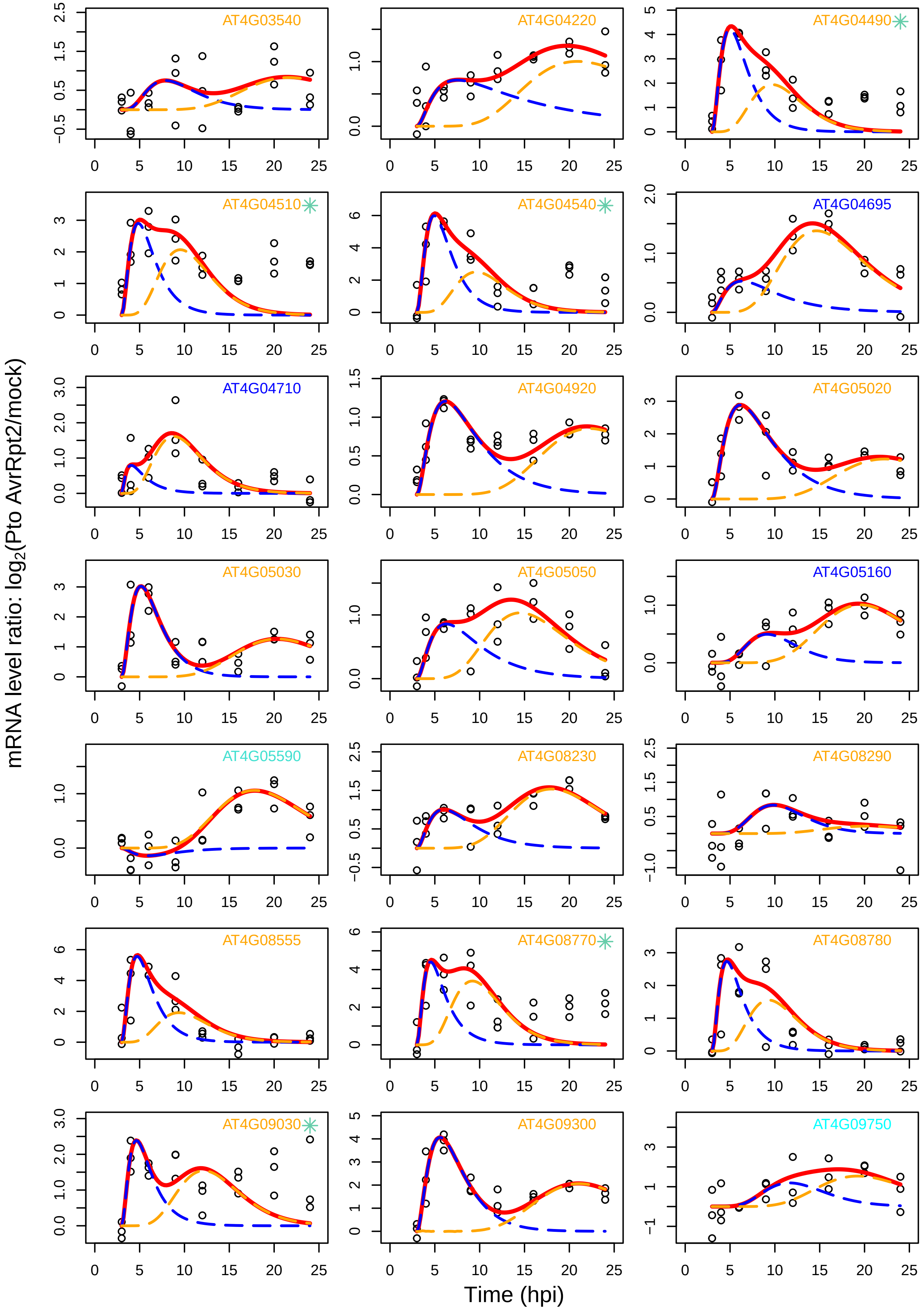
AT4G03540
AT4G04220
AT4G04490
AT4G04510
AT4G04540
AT4G04695
AT4G04710
AT4G04920
AT4G05020
AT4G05030
AT4G05050
AT4G05160
AT4G05590
AT4G08230
AT4G08290
AT4G08555
AT4G08770
AT4G08780
AT4G09030
AT4G09300
AT4G09750
mRNA level ratio: log2(Pto AvrRpt2/mock)
Time (hpi)

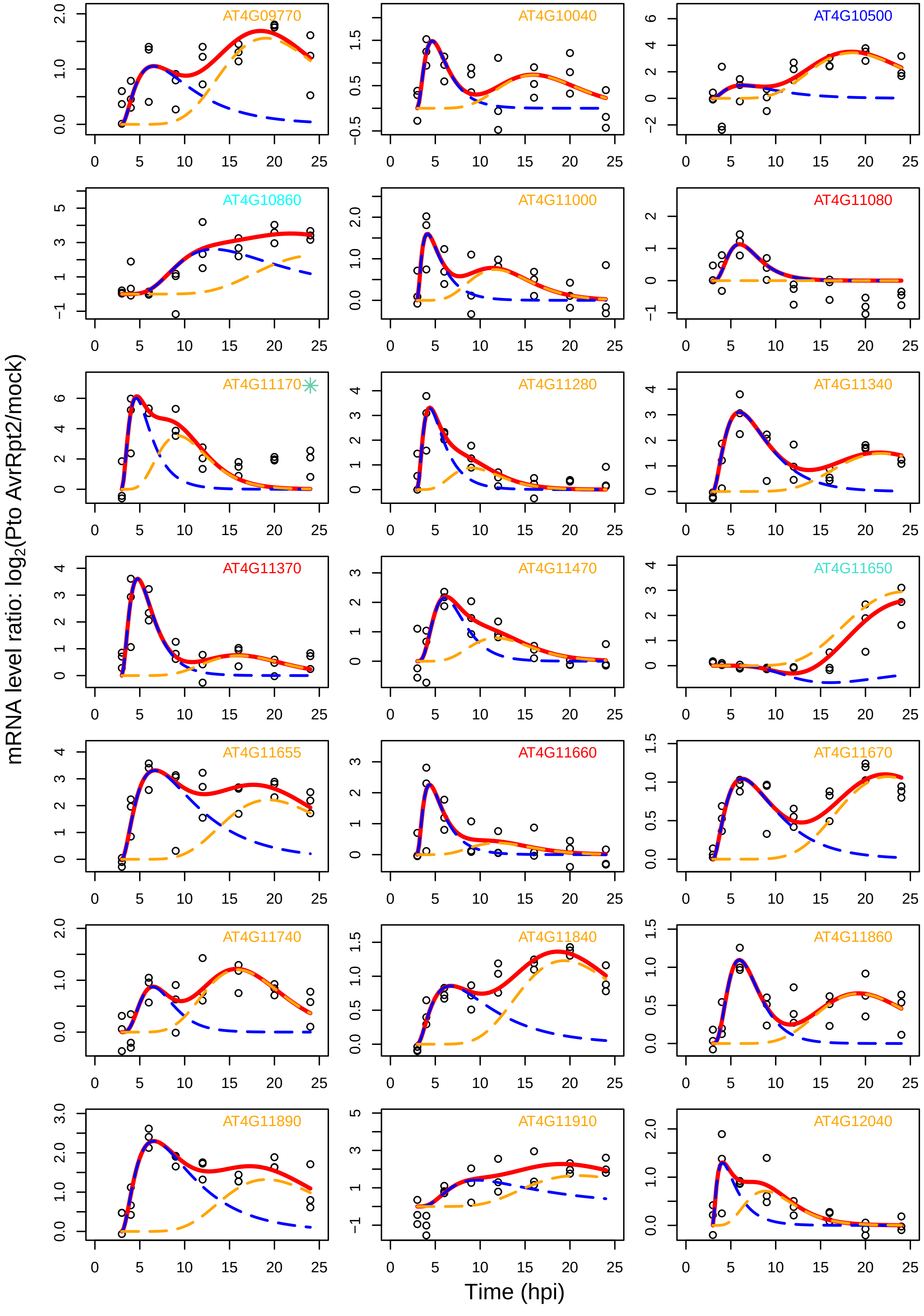
AT4G09770
AT4G10040
AT4G10500
AT4G10860
AT4G11000
AT4G11080
AT4G11170
AT4G11280
AT4G11340
AT4G11370
AT4G11470
AT4G11650
AT4G11655
AT4G11660
AT4G11670
AT4G11740
AT4G11840
AT4G11860
AT4G11890
AT4G11910
AT4G12040
mRNA level ratio: $\log_2$(Pto AvrRpt2/mock)
Time (hpi)

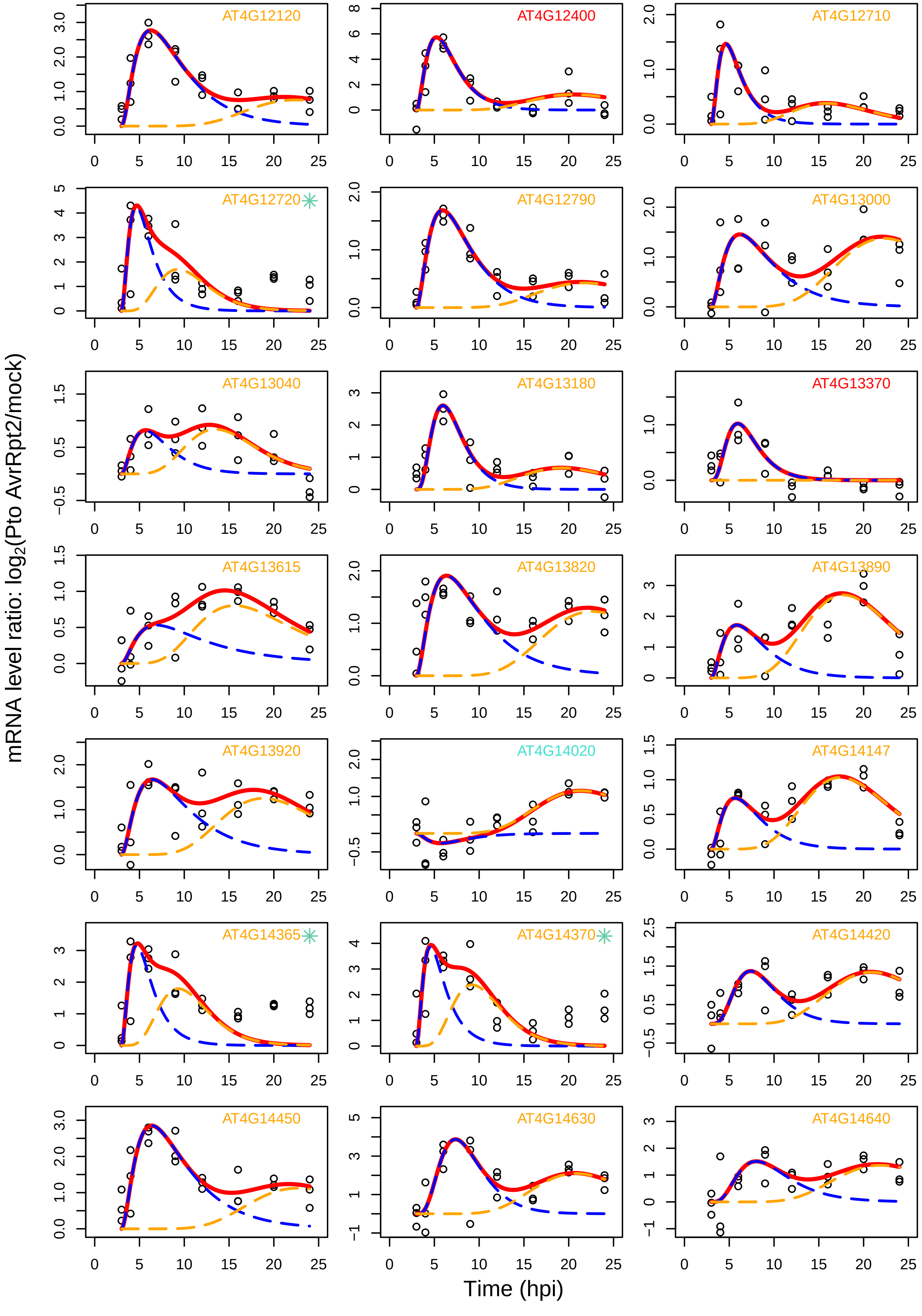
AT4G12120
AT4G12400
AT4G12710
AT4G12720
AT4G12790
AT4G13000
AT4G13040
AT4G13180
AT4G13370
AT4G13615
AT4G13820
AT4G13890
AT4G13920
AT4G14020
AT4G14147
AT4G14365
AT4G14370
AT4G14420
AT4G14450
AT4G14630
AT4G14640
mRNA level ratio: $log_2$(Pto AvrRpt2/mock)
Time (hpi)

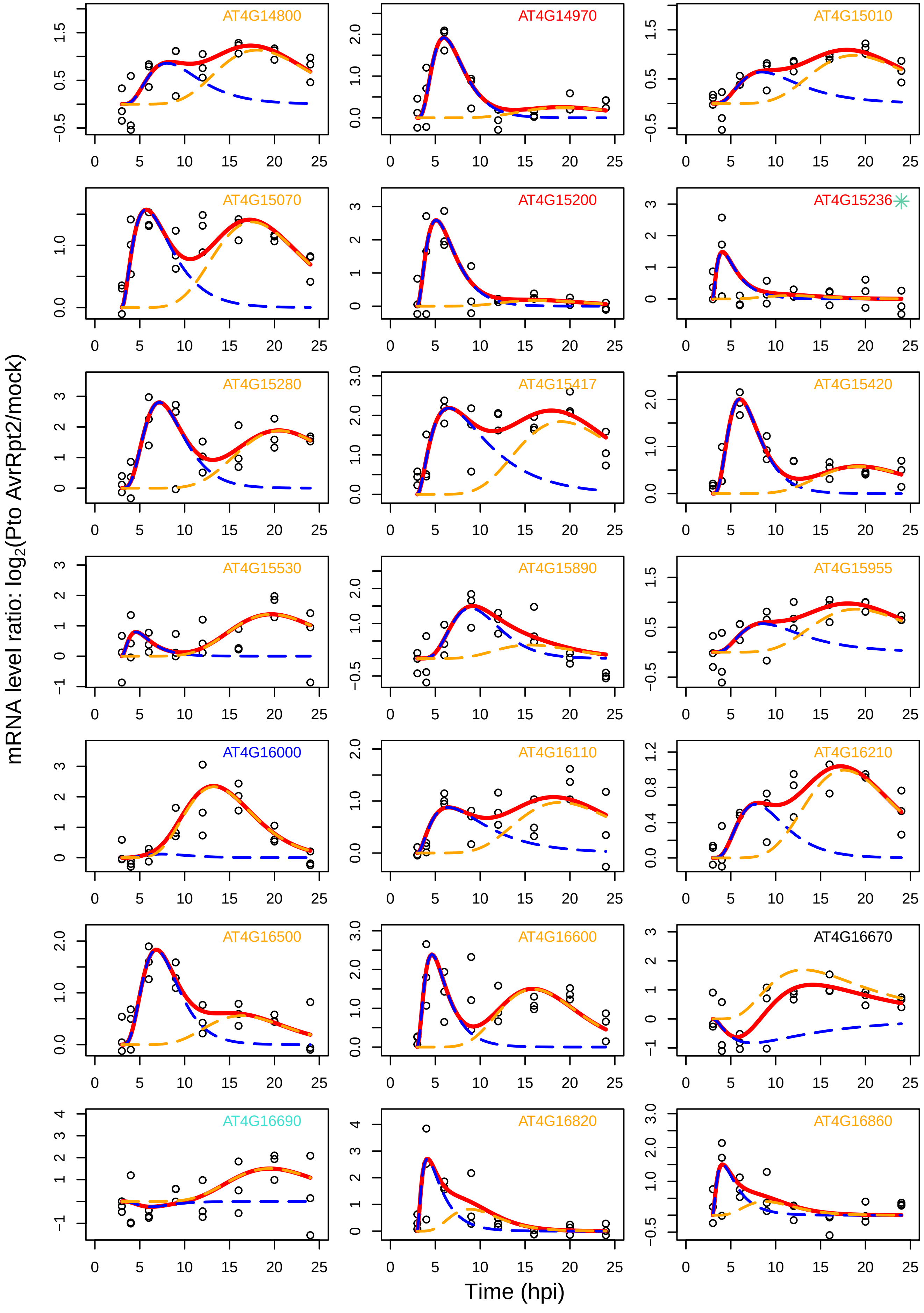
AT4G14800
AT4G14970
AT4G15010
AT4G15070
AT4G15200
AT4G15236
AT4G15280
AT4G15417
AT4G15420
AT4G15530
AT4G15890
AT4G15955
AT4G16000
AT4G16110
AT4G16210
AT4G16500
AT4G16600
AT4G16670
AT4G16690
AT4G16820
AT4G16860
mRNA level ratio: $log_2$(Pto AvrRpt2/mock)
Time (hpi)

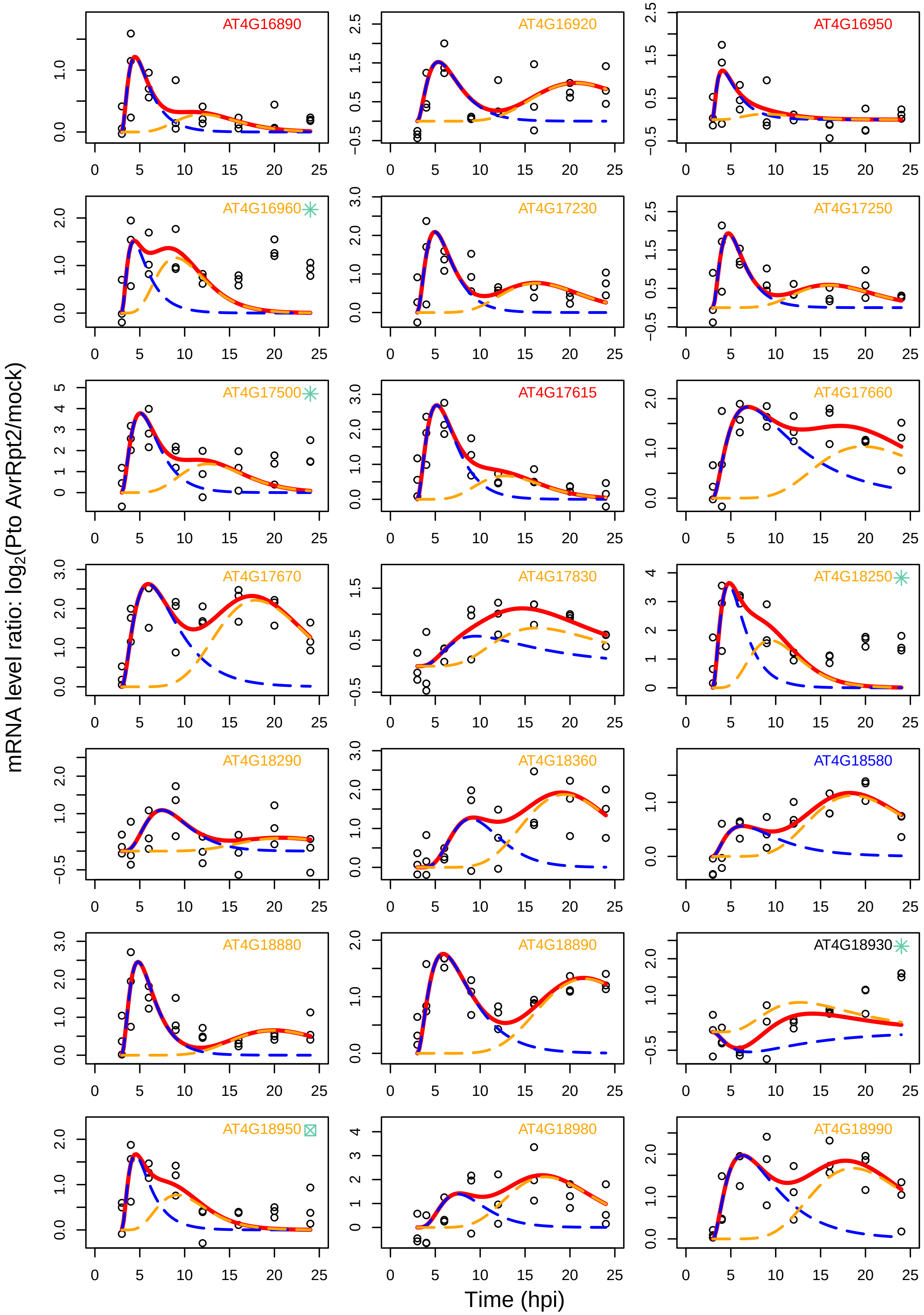
AT4G16890
AT4G16920
AT4G16950
AT4G16960
AT4G17230
AT4G17250
AT4G17500
AT4G17615
AT4G17660
AT4G17670
AT4G17830
AT4G18250
AT4G18290
AT4G18360
AT4G18580
AT4G18880
AT4G18890
AT4G18930
AT4G18950
AT4G18980
AT4G18990
mRNA level ratio: log2(Pto AvrRpt2/mock)
Time (hpi)

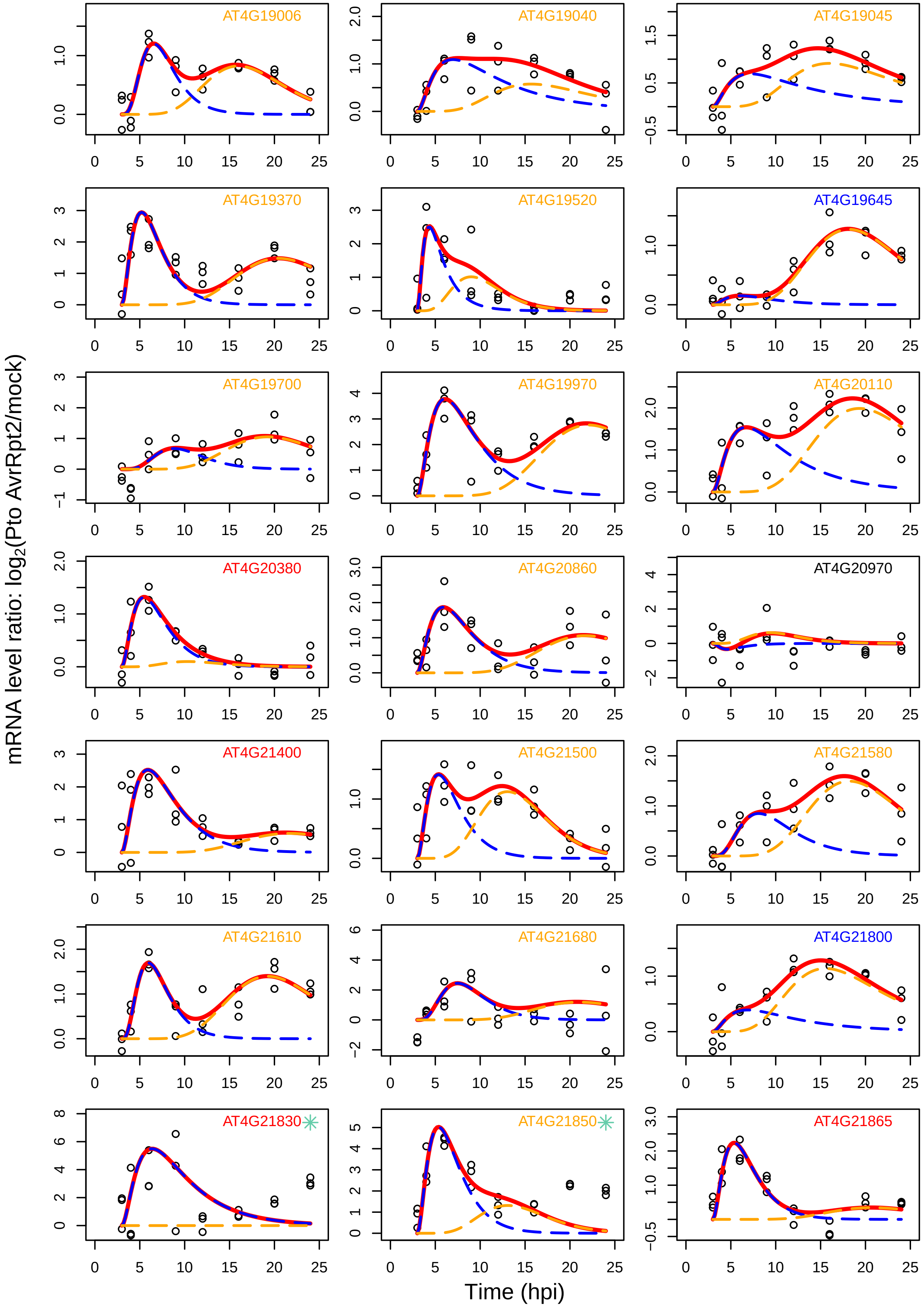
AT4G19006
AT4G19040
AT4G19045
AT4G19370
AT4G19520
AT4G19645
AT4G19700
AT4G19970
AT4G20110
AT4G20380
AT4G20860
AT4G20970
AT4G21400
AT4G21500
AT4G21580
AT4G21610
AT4G21680
AT4G21800
AT4G21830
AT4G21850
AT4G21865
mRNA level ratio: $\log_2$(Pto AvrRpt2/mock)
Time (hpi)

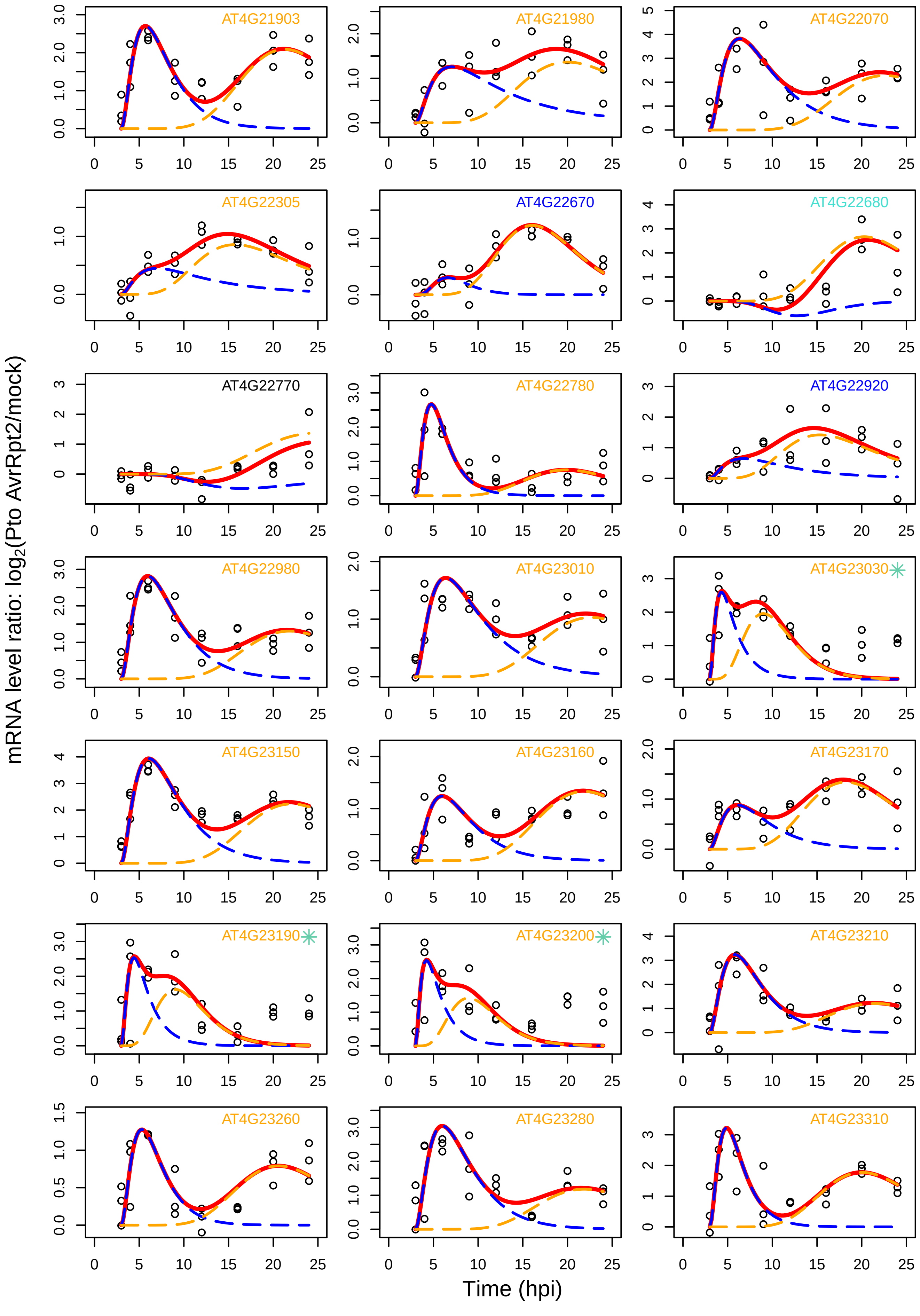
AT4G21903
AT4G21980
AT4G22070
AT4G22305
AT4G22670
AT4G22680
AT4G22770
AT4G22780
AT4G22920
AT4G22980
AT4G23010
AT4G23030
AT4G23150
AT4G23160
AT4G23170
AT4G23190
AT4G23200
AT4G23210
AT4G23260
AT4G23280
AT4G23310
mRNA level ratio: log2(Pto AvrRpt2/mock)
Time (hpi)

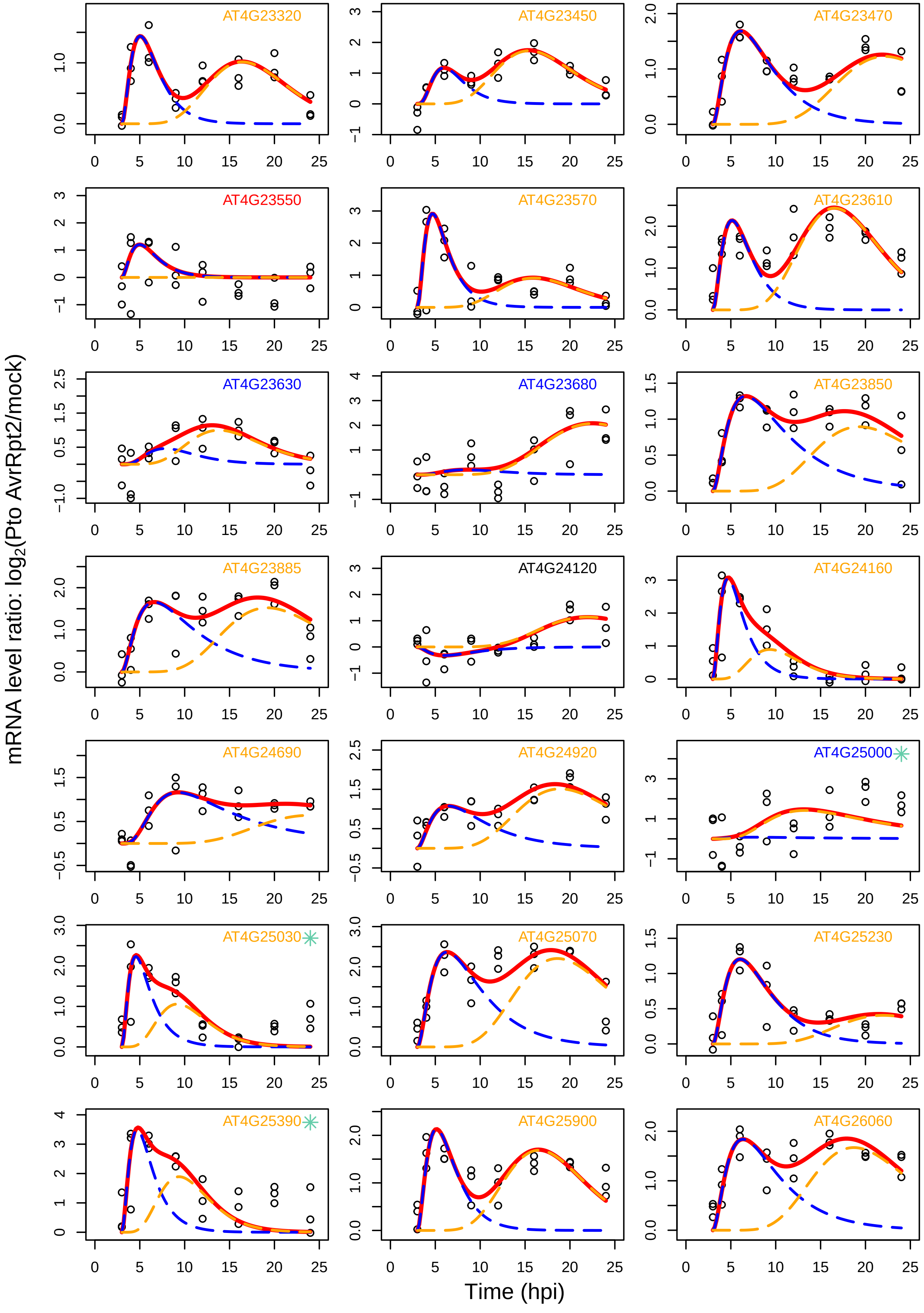
AT4G23320
AT4G23450
AT4G23470
AT4G23550
AT4G23570
AT4G23610
AT4G23630
AT4G23680
AT4G23850
AT4G23885
AT4G24120
AT4G24160
AT4G24690
AT4G24920
AT4G25000
AT4G25030
AT4G25070
AT4G25230
AT4G25390
AT4G25900
AT4G26060
mRNA level ratio: $log_2$(Pto AvrRpt2/mock)
Time (hpi)

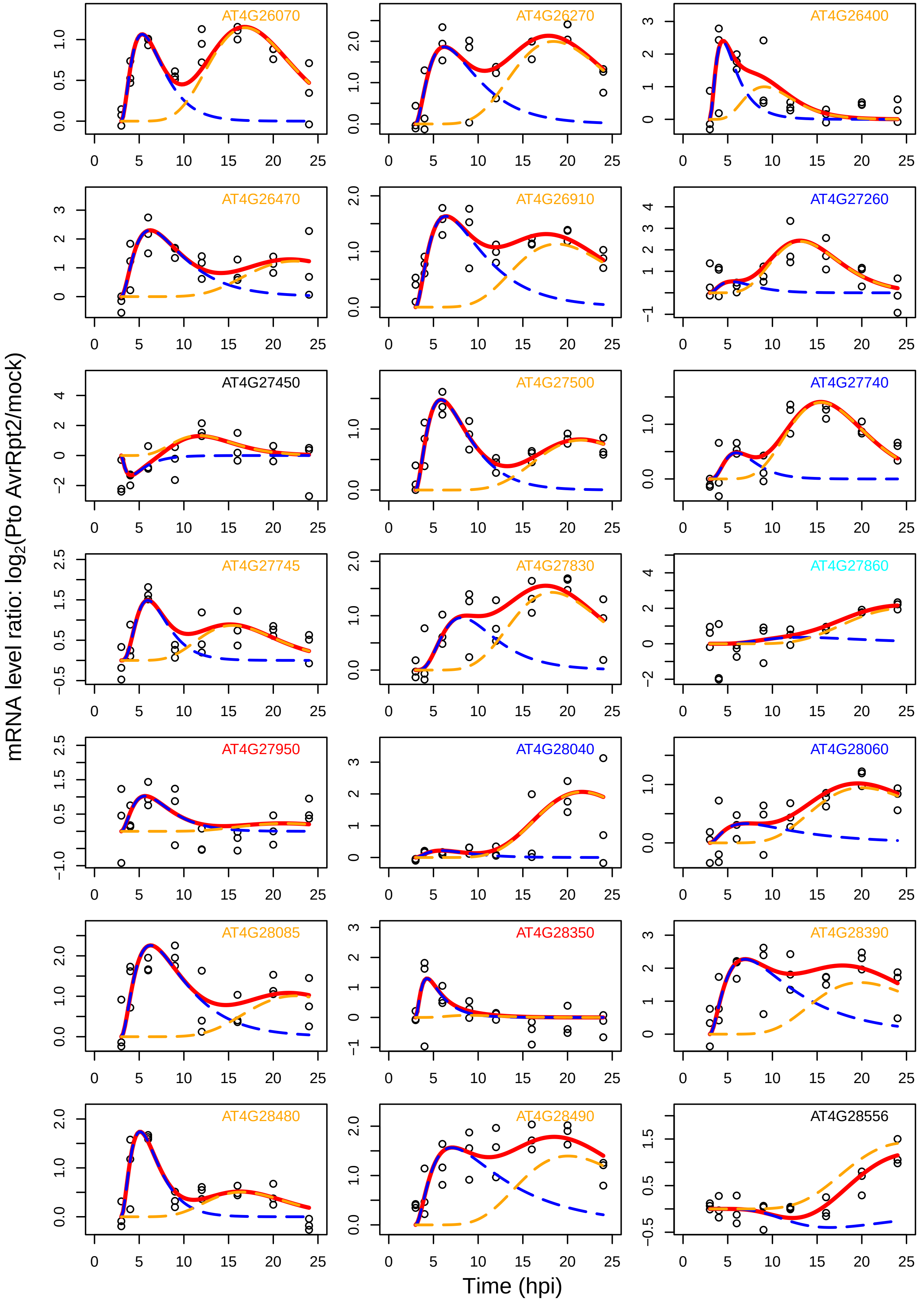

AT4G26070
AT4G26270
AT4G26400
AT4G26470
AT4G26910
AT4G27260
AT4G27450
AT4G27500
AT4G27740
AT4G27745
AT4G27830
AT4G27860
AT4G27950
AT4G28040
AT4G28060
AT4G28085
AT4G28350
AT4G28390
AT4G28480
AT4G28490
AT4G28556
mRNA level ratio: $\log_2$(Pto AvrRpt2/mock)
Time (hpi)

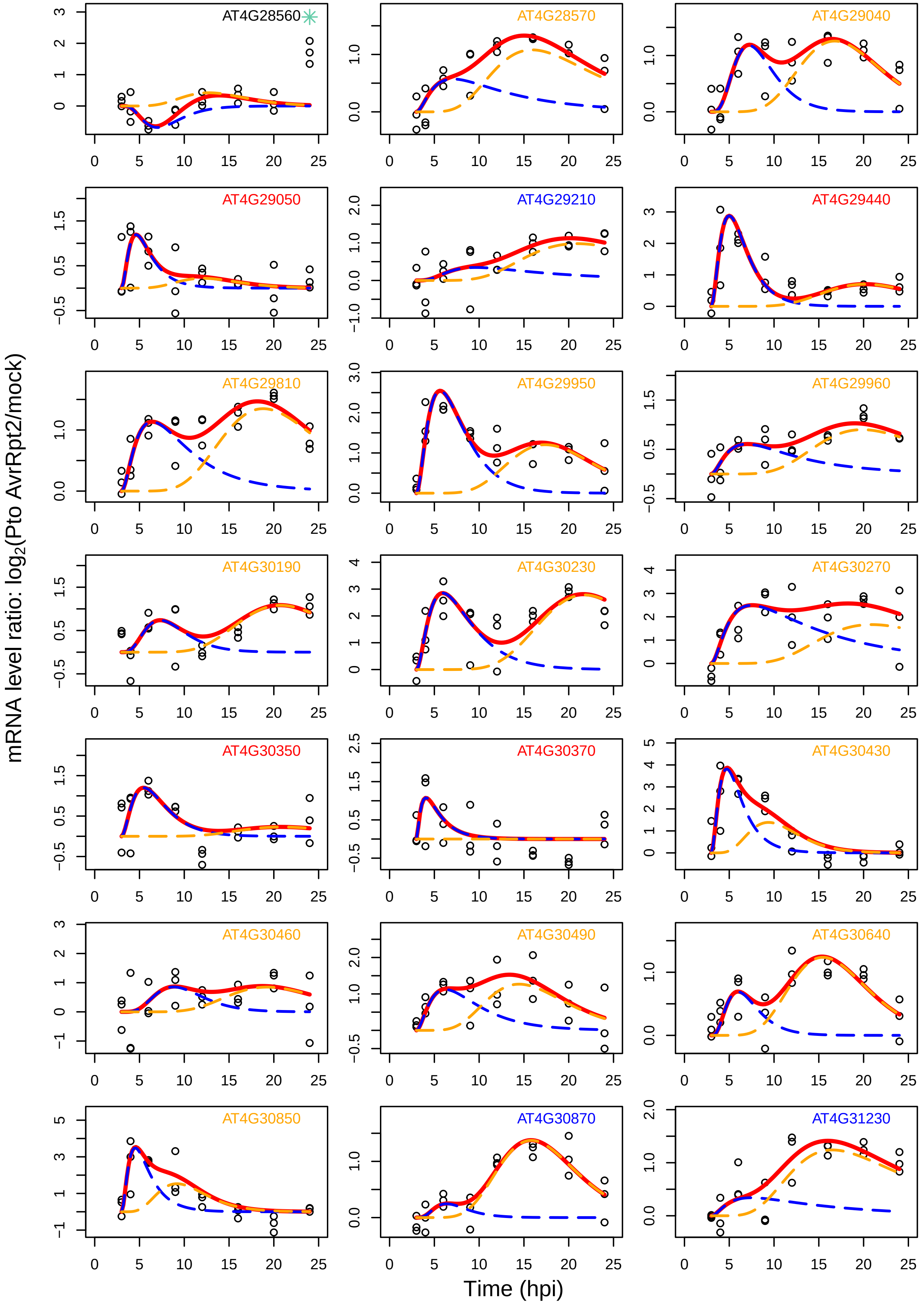
AT4G28560
AT4G28570
AT4G29040
AT4G29050
AT4G29210
AT4G29440
AT4G29810
AT4G29950
AT4G29960
AT4G30190
AT4G30230
AT4G30270
AT4G30350
AT4G30370
AT4G30430
AT4G30460
AT4G30490
AT4G30640
AT4G30850
AT4G30870
AT4G31230
mRNA level ratio: log2(Pto AvrRpt2/mock)
Time (hpi)

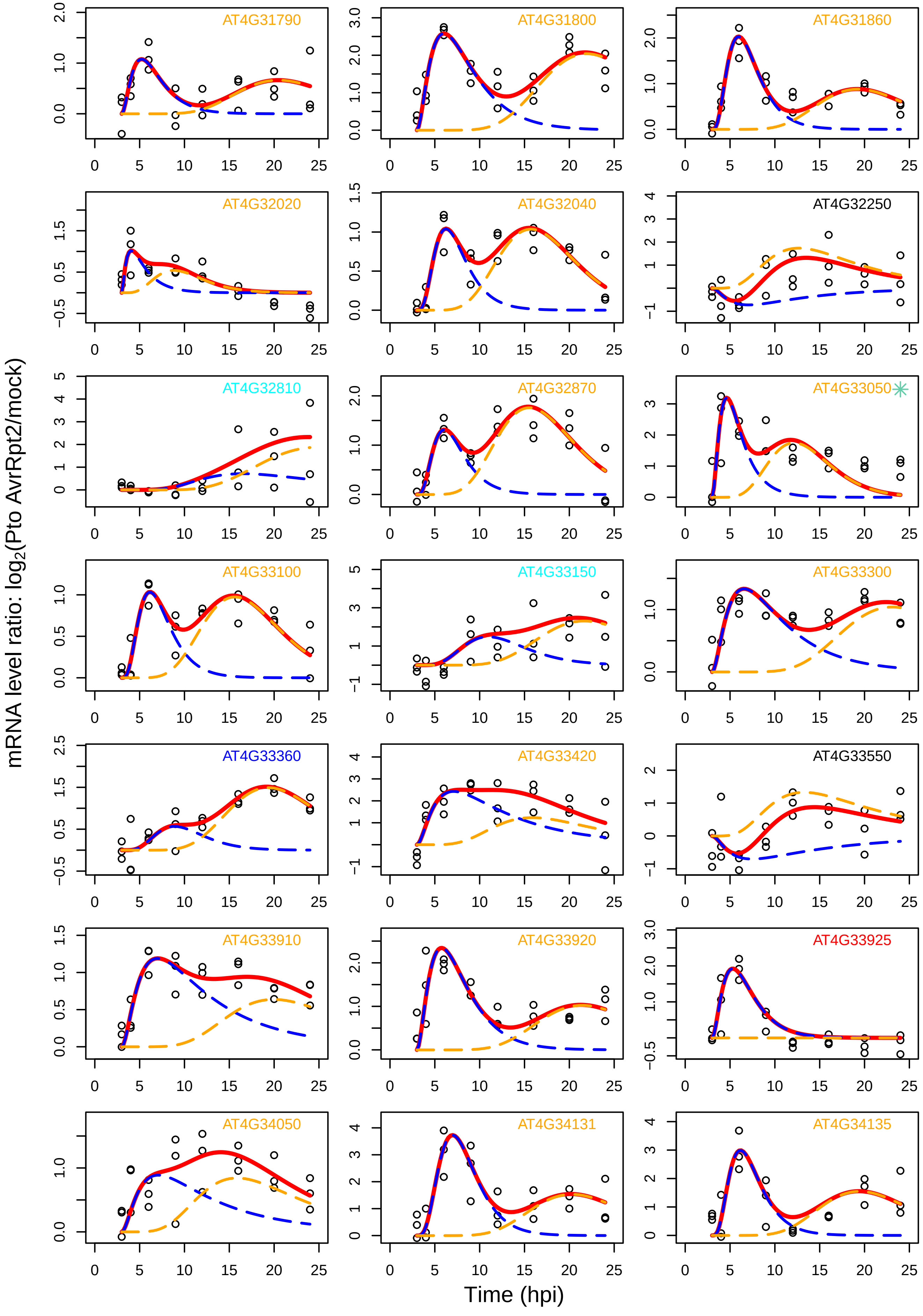
AT4G31790
AT4G31800
AT4G31860
AT4G32020
AT4G32040
AT4G32250
AT4G32810
AT4G32870
AT4G33050
AT4G33100
AT4G33150
AT4G33300
AT4G33360
AT4G33420
AT4G33550
AT4G33910
AT4G33920
AT4G33925
AT4G34050
AT4G34131
AT4G34135
mRNA level ratio: log2(Pto AvrRpt2/mock)
Time (hpi)

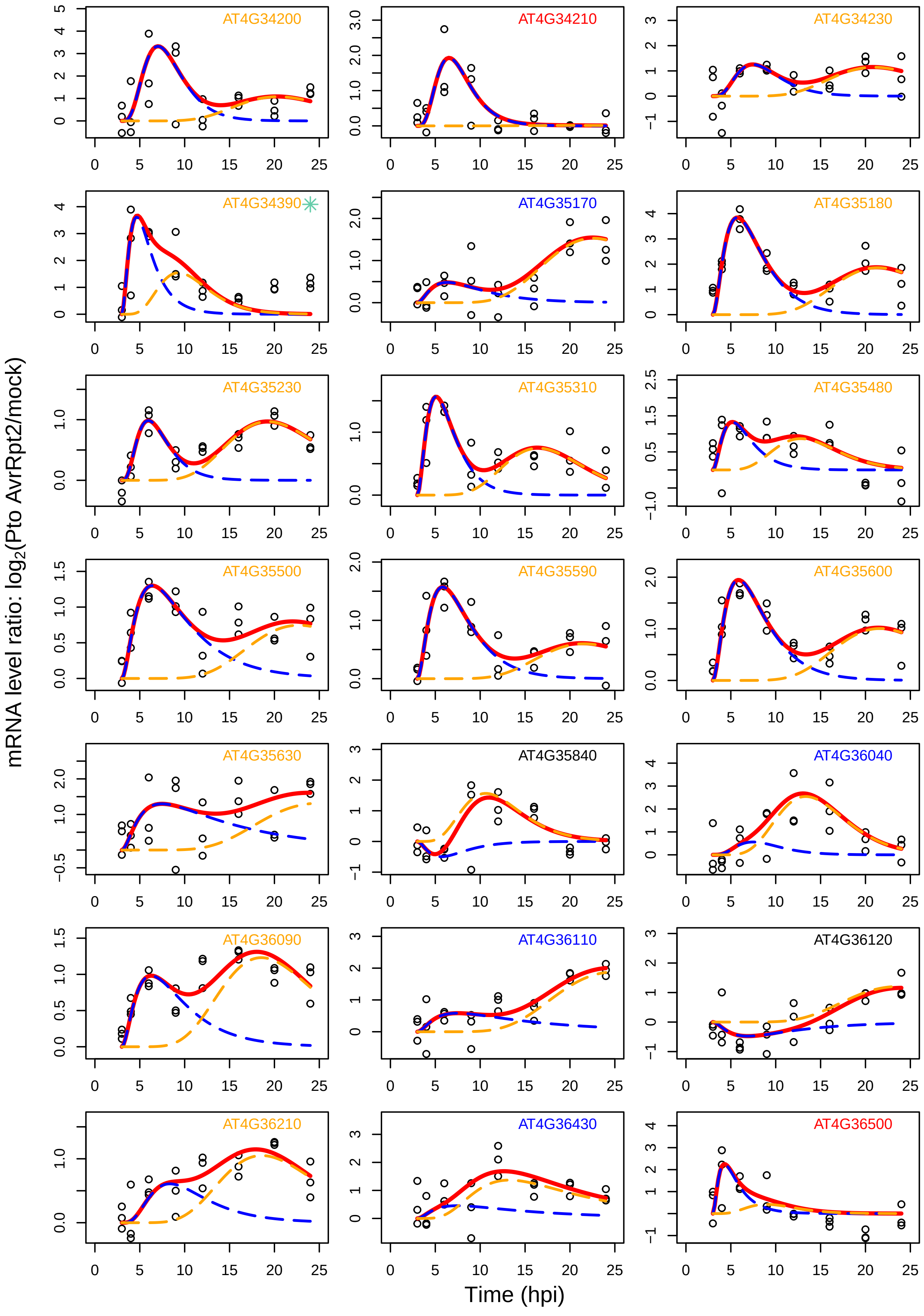
AT4G34200
AT4G34210
AT4G34230
AT4G34390
AT4G35170
AT4G35180
AT4G35230
AT4G35310
AT4G35480
AT4G35500
AT4G35590
AT4G35600
AT4G35630
AT4G35840
AT4G36040
AT4G36090
AT4G36110
AT4G36120
AT4G36210
AT4G36430
AT4G36500
mRNA level ratio: $log_2$(Pto AvrRpt2/mock)
Time (hpi)

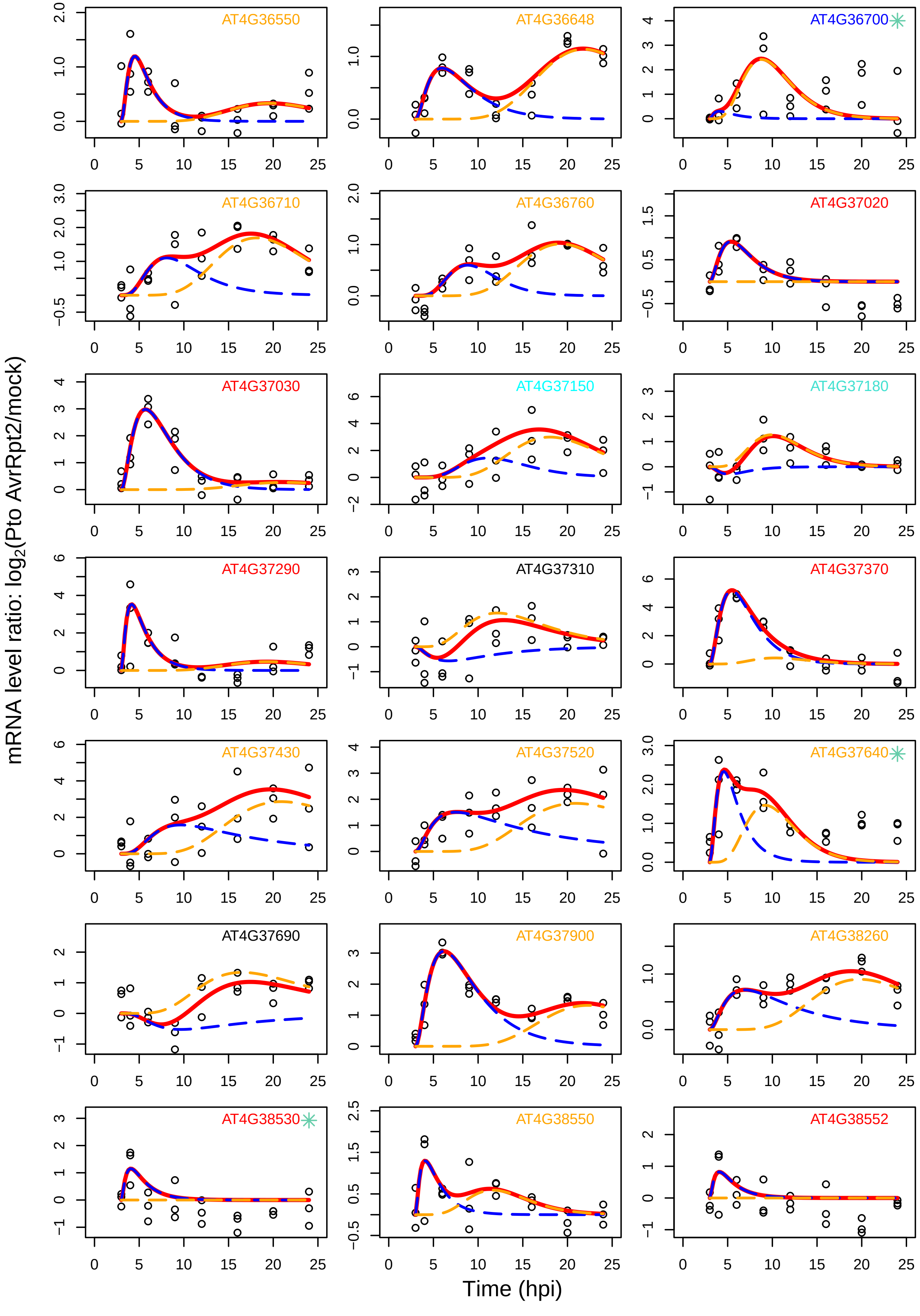
AT4G36550
AT4G36648
AT4G36700
AT4G36710
AT4G36760
AT4G37020
AT4G37030
AT4G37150
AT4G37180
AT4G37290
AT4G37310
AT4G37370
AT4G37430
AT4G37520
AT4G37640
AT4G37690
AT4G37900
AT4G38260
AT4G38530
AT4G38550
AT4G38552
mRNA level ratio: $\log_2$(Pto AvrRpt2/mock)
Time (hpi)

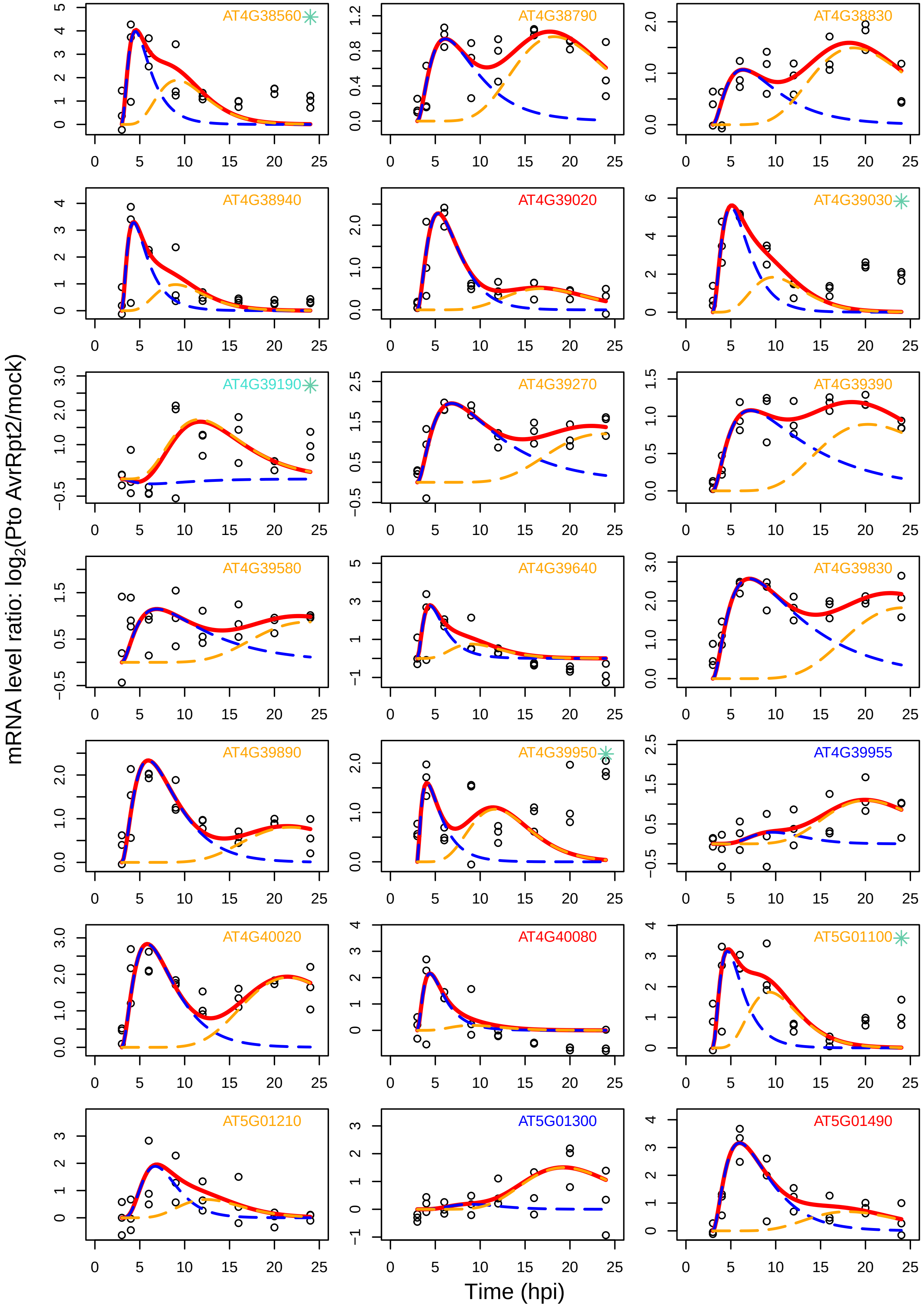

AT4G38560
AT4G38790
AT4G38830
AT4G38940
AT4G39020
AT4G39030
AT4G39190
AT4G39270
AT4G39390
AT4G39580
AT4G39640
AT4G39830
AT4G39890
AT4G39950
AT4G39955
AT4G40020
AT4G40080
AT5G01100
AT5G01210
AT5G01300
AT5G01490
mRNA level ratio: $\log_2$(Pto AvrRpt2/mock)
Time (hpi)

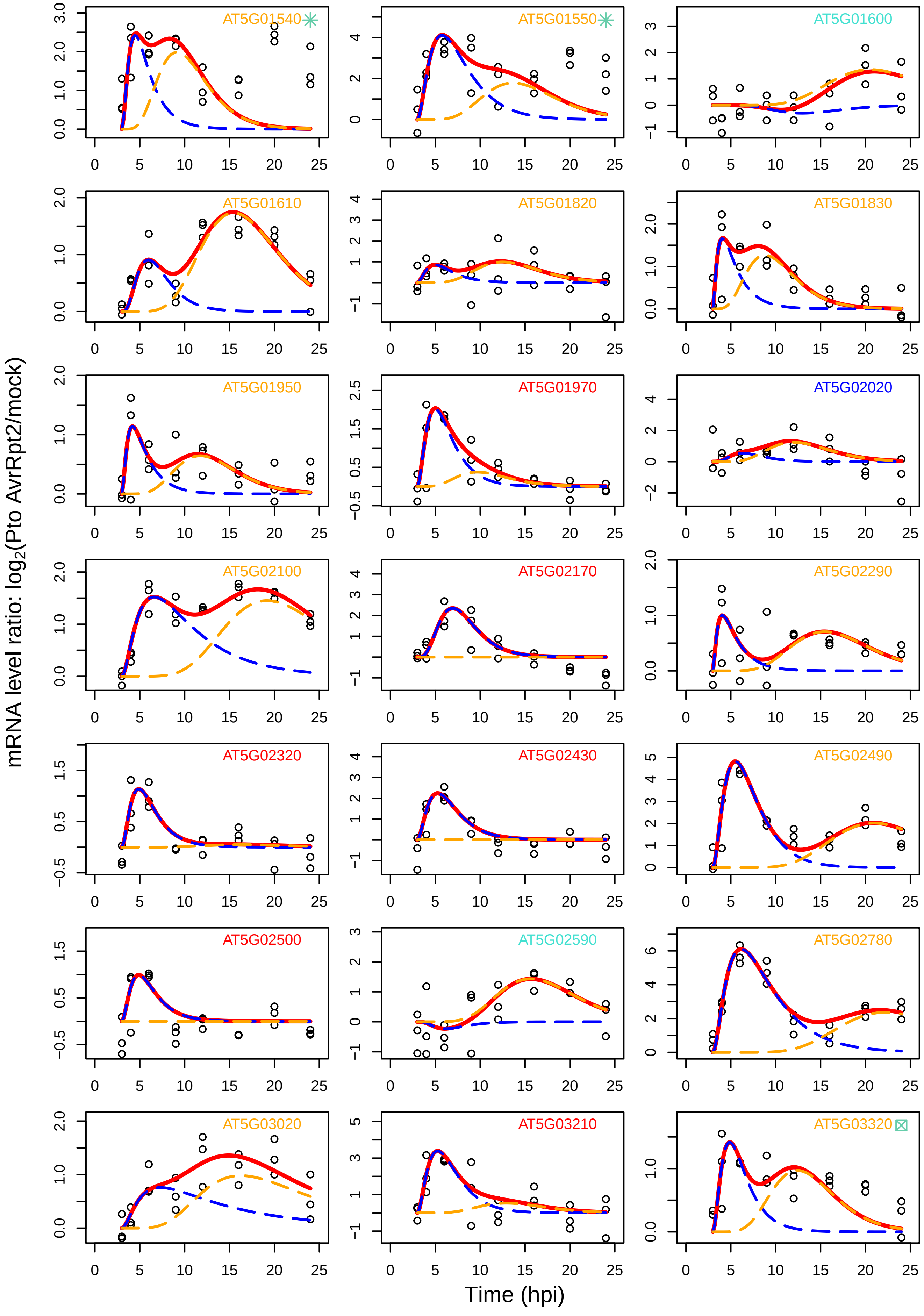

AT5G01540
AT5G01550
AT5G01600
AT5G01610
AT5G01820
AT5G01830
AT5G01950
AT5G01970
AT5G02020
AT5G02100
AT5G02170
AT5G02290
AT5G02320
AT5G02430
AT5G02490
AT5G02500
AT5G02590
AT5G02780
AT5G03020
AT5G03210
AT5G03320
mRNA level ratio: log2(Pto AvrRpt2/mock)
Time (hpi)

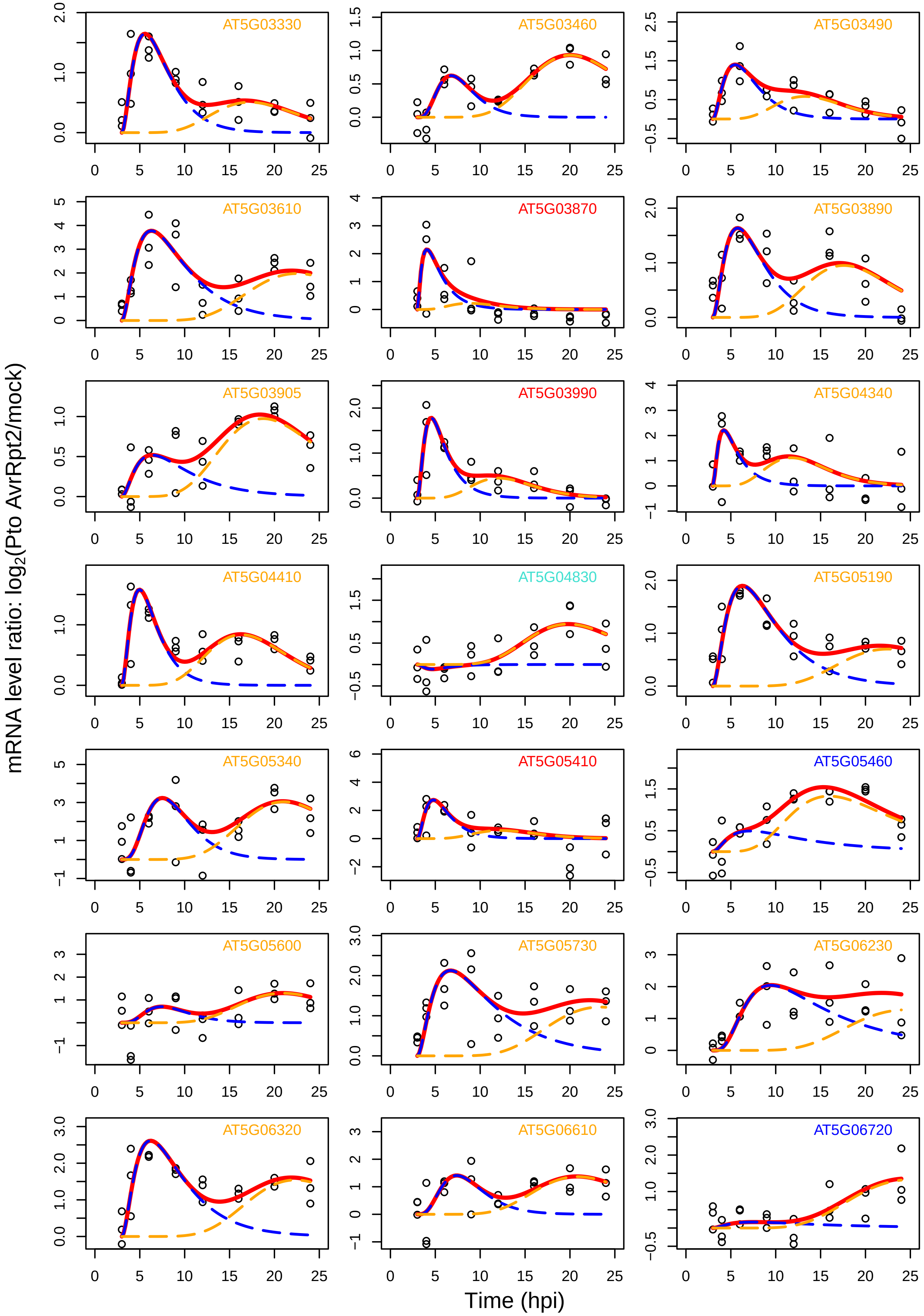

AT5G03330
AT5G03460
AT5G03490
AT5G03610
AT5G03870
AT5G03890
AT5G03905
AT5G03990
AT5G04340
AT5G04410
AT5G04830
AT5G05190
AT5G05340
AT5G05410
AT5G05460
AT5G05600
AT5G05730
AT5G06230
AT5G06320
AT5G06610
AT5G06720
mRNA level ratio: $\log_2$(Pto AvrRpt2/mock)
Time (hpi)

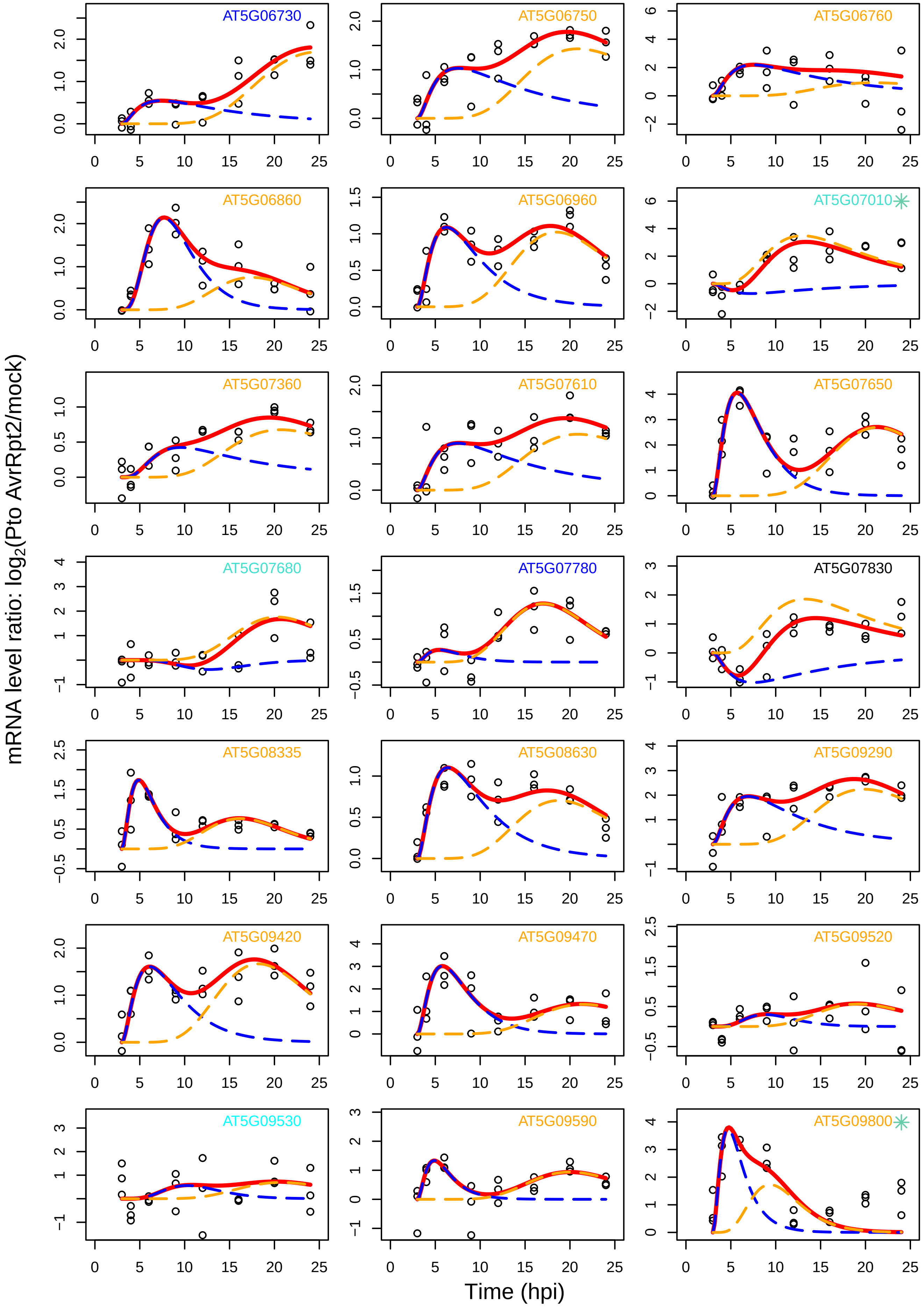
AT5G06730
AT5G06750
AT5G06760
AT5G06860
AT5G06960
AT5G07010
AT5G07360
AT5G07610
AT5G07650
AT5G07680
AT5G07780
AT5G07830
AT5G08335
AT5G08630
AT5G09290
AT5G09420
AT5G09470
AT5G09520
AT5G09530
AT5G09590
AT5G09800
mRNA level ratio: log2(Pto AvrRpt2/mock)
Time (hpi)

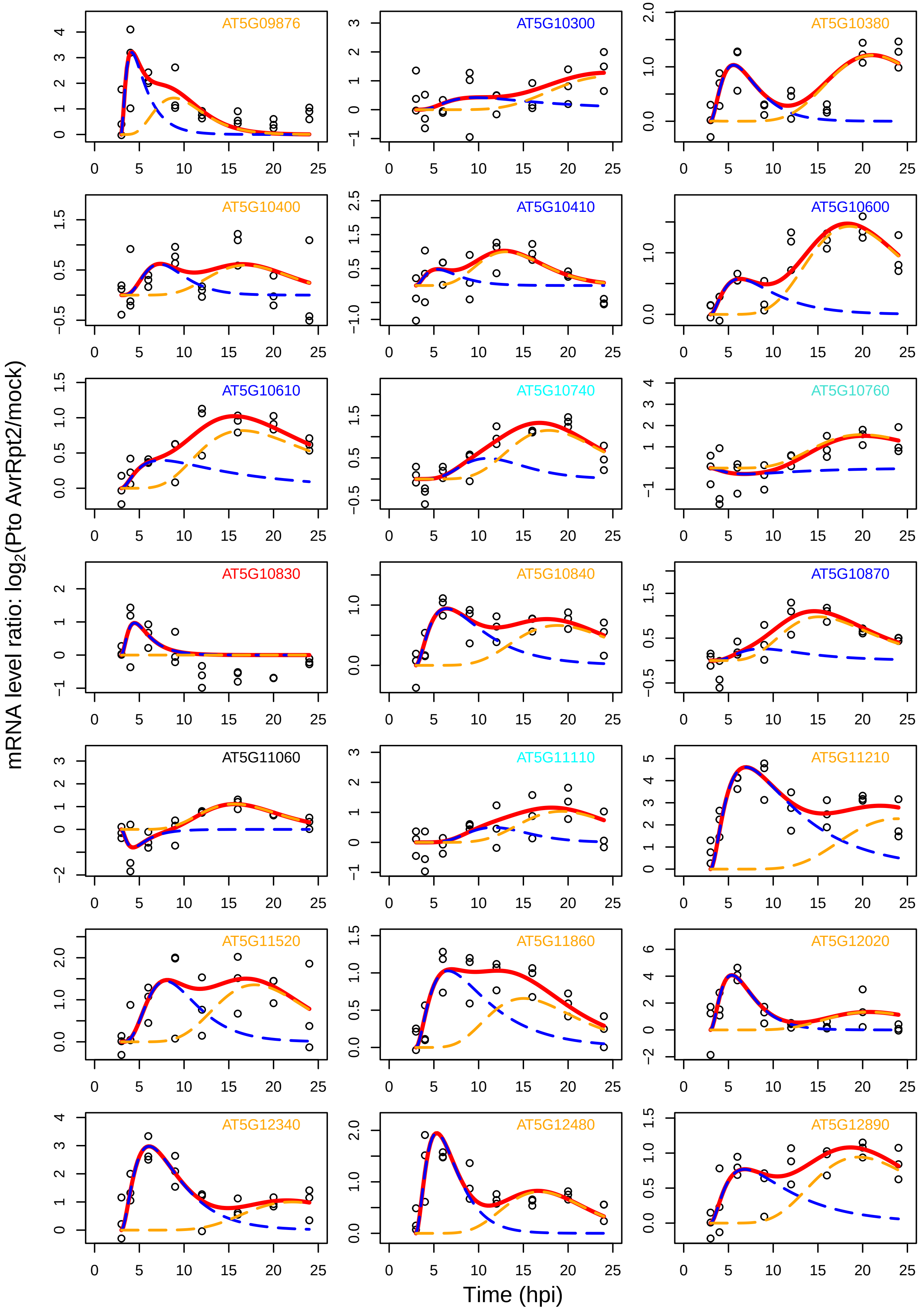

AT5G09876
AT5G10300
AT5G10380
AT5G10400
AT5G10410
AT5G10600
AT5G10610
AT5G10740
AT5G10760
AT5G10830
AT5G10840
AT5G10870
AT5G11060
AT5G11110
AT5G11210
AT5G11520
AT5G11860
AT5G12020
AT5G12340
AT5G12480
AT5G12890
mRNA level ratio: $\log_2$(Pto AvrRpt2/mock)
Time (hpi)

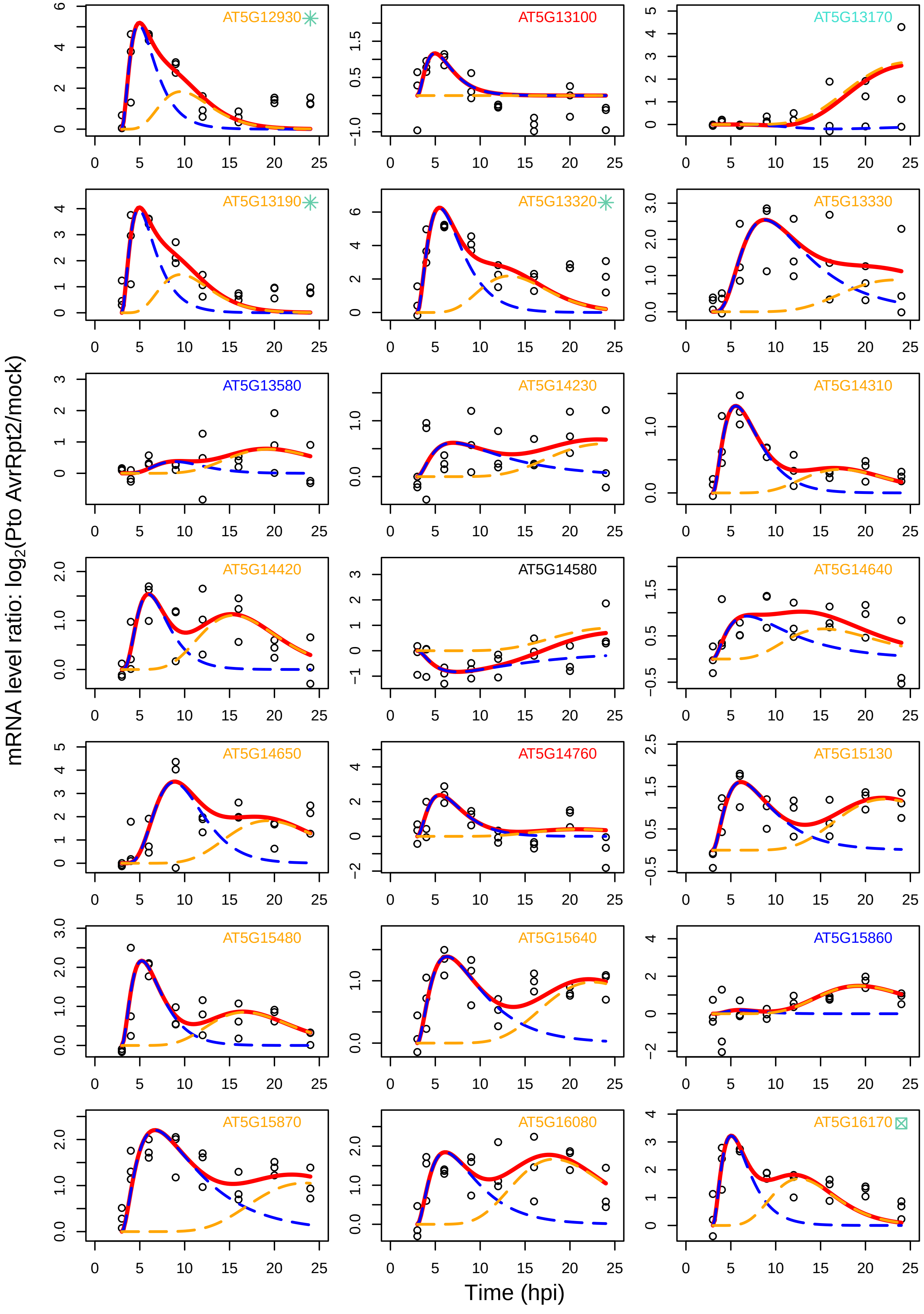

mRNA level ratio: $log_2$(Pto AvrRpt2/mock)
AT5G12930
AT5G13100
AT5G13170
AT5G13190
AT5G13320
AT5G13330
AT5G13580
AT5G14230
AT5G14310
AT5G14420
AT5G14580
AT5G14640
AT5G14650
AT5G14760
AT5G15130
AT5G15480
AT5G15640
AT5G15860
AT5G15870
AT5G16080
AT5G16170
Time (hpi)

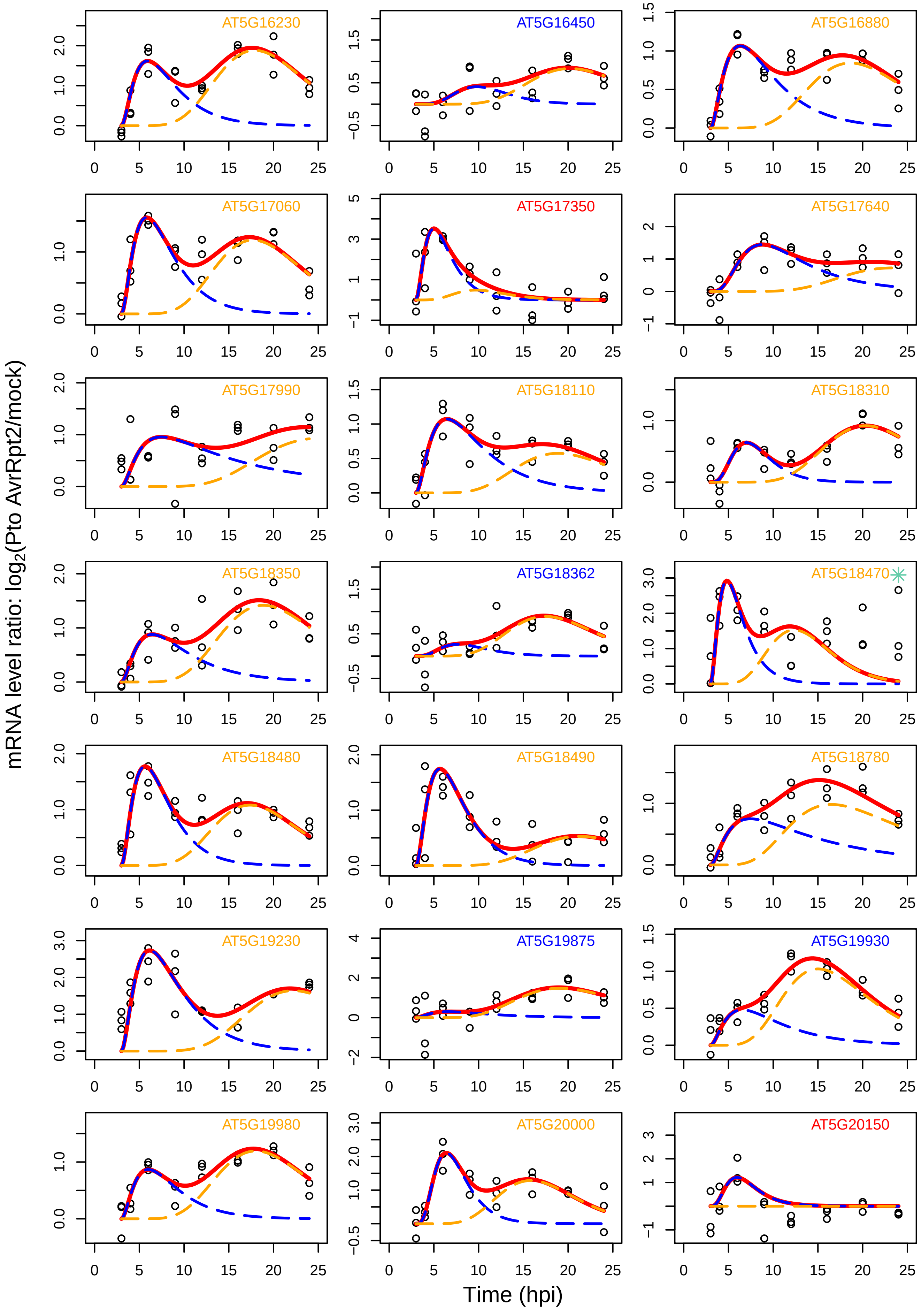

AT5G16230
AT5G16450
AT5G16880
AT5G17060
AT5G17350
AT5G17640
AT5G17990
AT5G18110
AT5G18310
AT5G18350
AT5G18362
AT5G18470
AT5G18480
AT5G18490
AT5G18780
AT5G19230
AT5G19875
AT5G19930
AT5G19980
AT5G20000
AT5G20150
mRNA level ratio: log2(Pto AvrRpt2/mock)
Time (hpi)

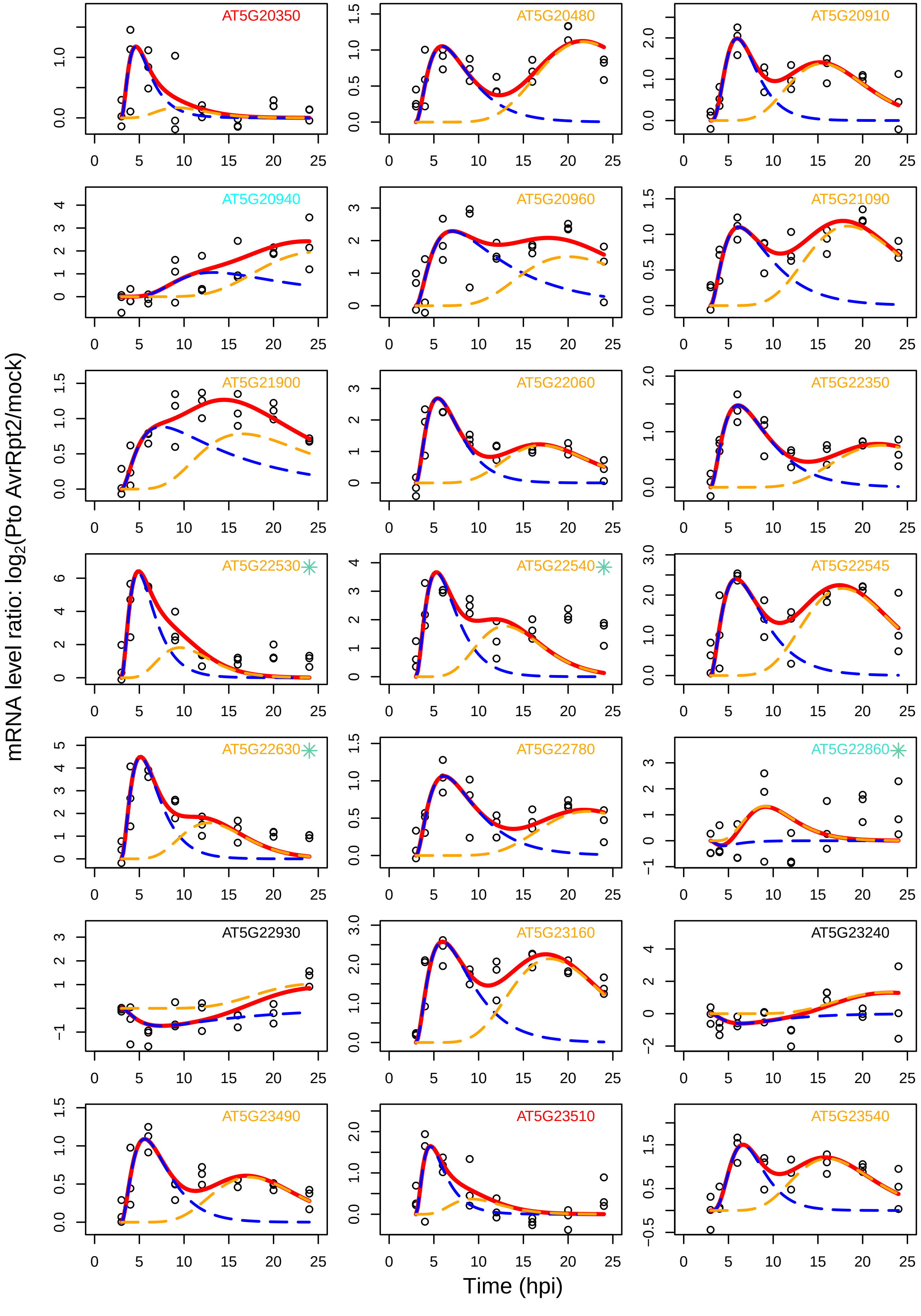

AT5G20350
AT5G20480
AT5G20910
AT5G20940
AT5G20960
AT5G21090
AT5G21900
AT5G22060
AT5G22350
AT5G22530
AT5G22540
AT5G22545
AT5G22630
AT5G22780
AT5G22860
AT5G22930
AT5G23160
AT5G23240
AT5G23490
AT5G23510
AT5G23540
mRNA level ratio: log2(Pto AvrRpt2/mock)
Time (hpi)

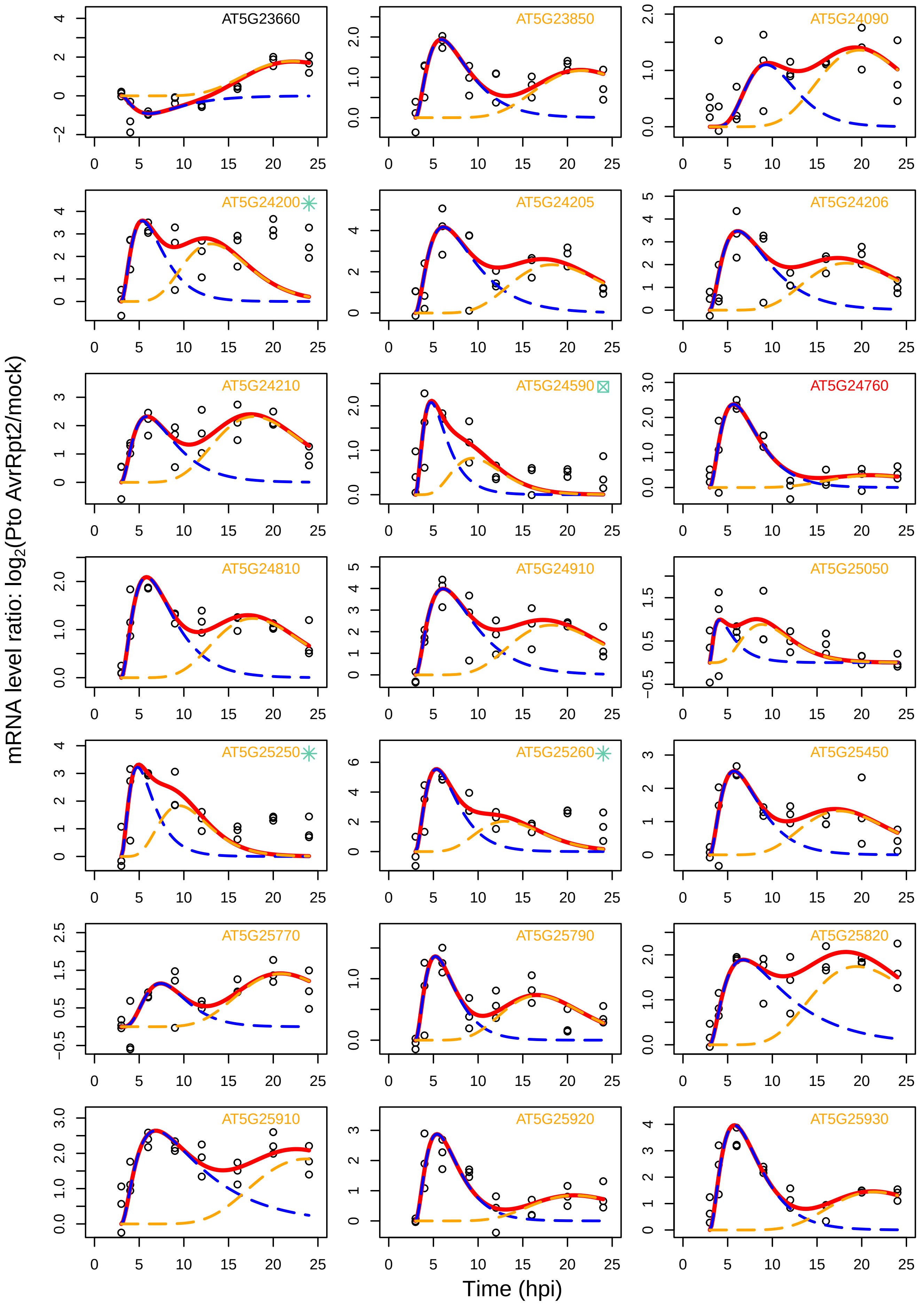

AT5G23660
AT5G23850
AT5G24090
AT5G24200
AT5G24205
AT5G24206
AT5G24210
AT5G24590
AT5G24760
AT5G24810
AT5G24910
AT5G25050
AT5G25250
AT5G25260
AT5G25450
AT5G25770
AT5G25790
AT5G25820
AT5G25910
AT5G25920
AT5G25930
mRNA level ratio: log2(Pto AvrRpt2/mock)
Time (hpi)

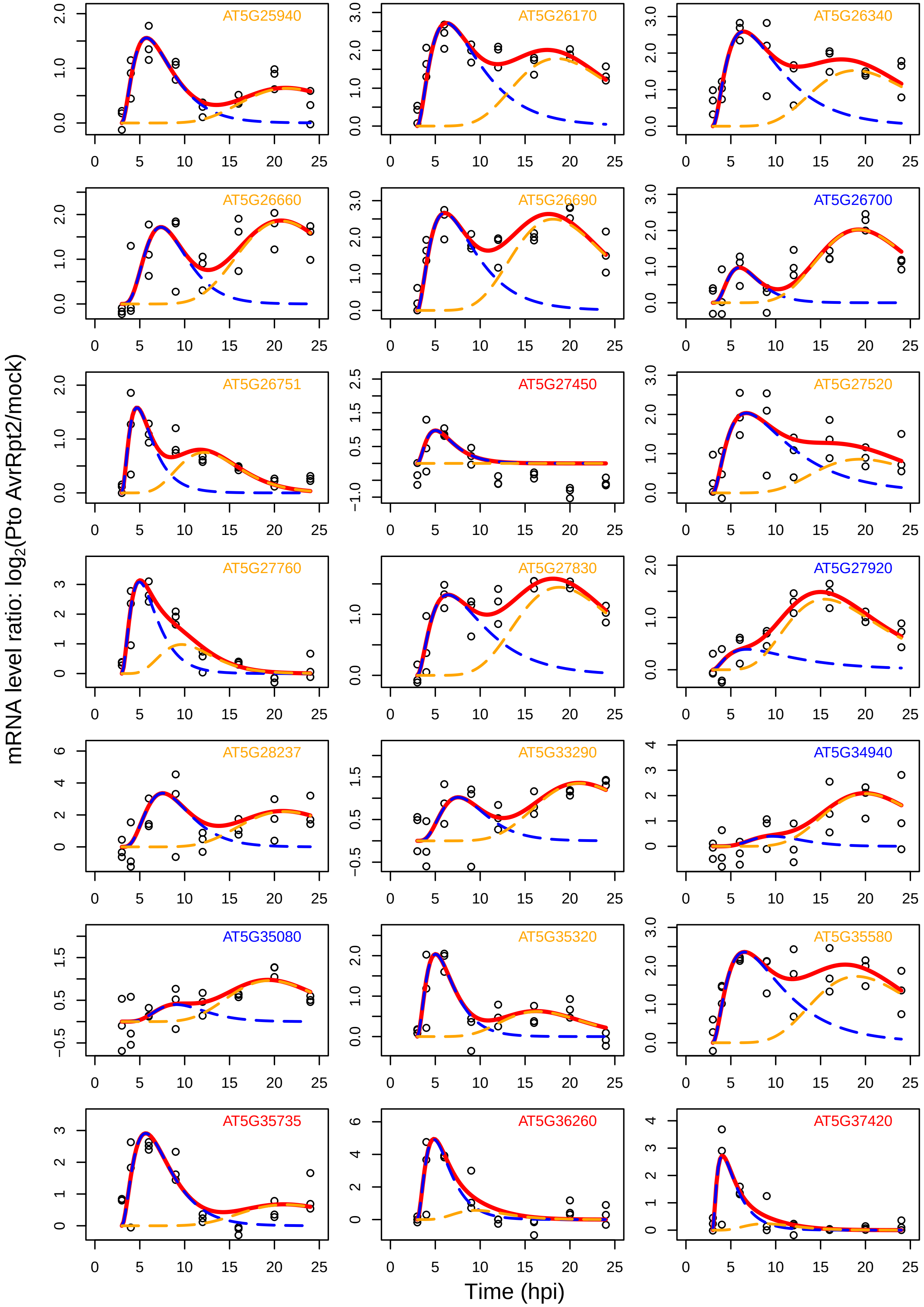

AT5G25940
AT5G26170
AT5G26340
AT5G26660
AT5G26690
AT5G26700
AT5G26751
AT5G27450
AT5G27520
AT5G27760
AT5G27830
AT5G27920
AT5G28237
AT5G33290
AT5G34940
AT5G35080
AT5G35320
AT5G35580
AT5G35735
AT5G36260
AT5G37420
mRNA level ratio: log2(Pto AvrRpt2/mock)
Time (hpi)

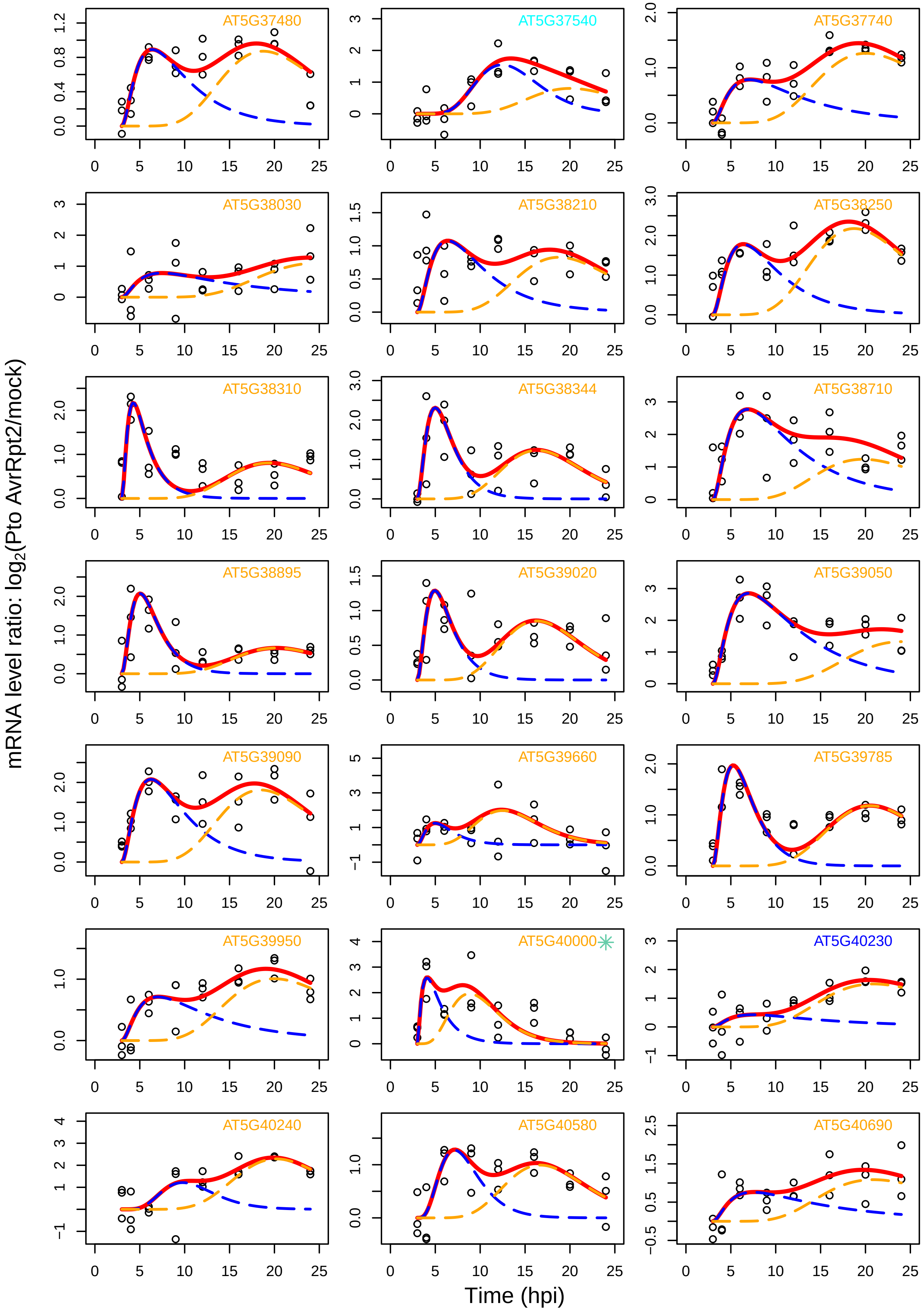
mRNA level ratio: log2(Pto AvrRpt2/mock)
Time (hpi)
AT5G37480
AT5G37540
AT5G37740
AT5G38030
AT5G38210
AT5G38250
AT5G38310
AT5G38344
AT5G38710
AT5G38895
AT5G39020
AT5G39050
AT5G39090
AT5G39660
AT5G39785
AT5G39950
AT5G40000
AT5G40230
AT5G40240
AT5G40580
AT5G40690

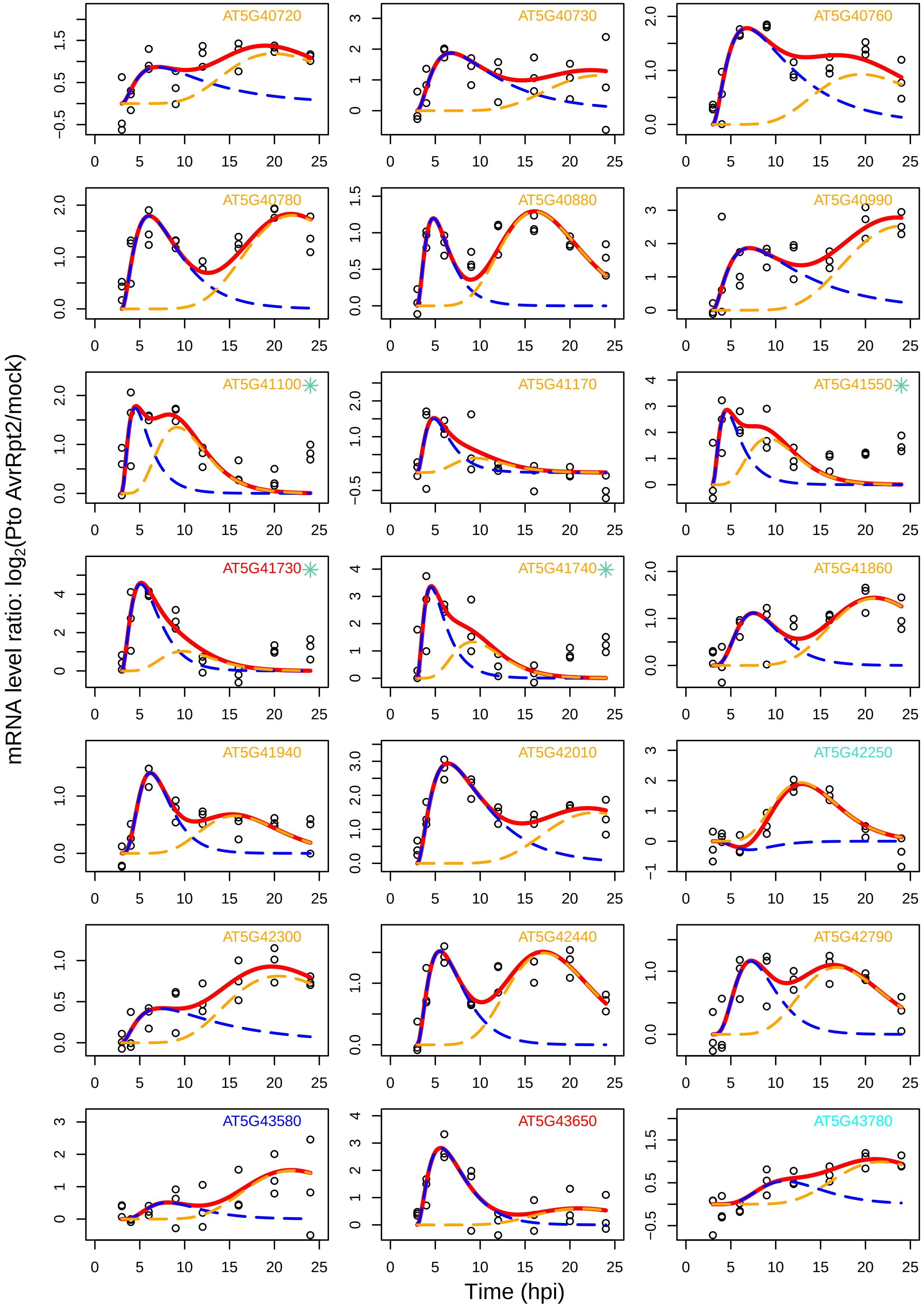

AT5G40720
AT5G40730
AT5G40760
AT5G40780
AT5G40880
AT5G40990
AT5G41100
AT5G41170
AT5G41550
AT5G41730
AT5G41740
AT5G41860
AT5G41940
AT5G42010
AT5G42250
AT5G42300
AT5G42440
AT5G42790
AT5G43580
AT5G43650
AT5G43780
mRNA level ratio: $\log_2$(Pto AvrRpt2/mock)
Time (hpi)

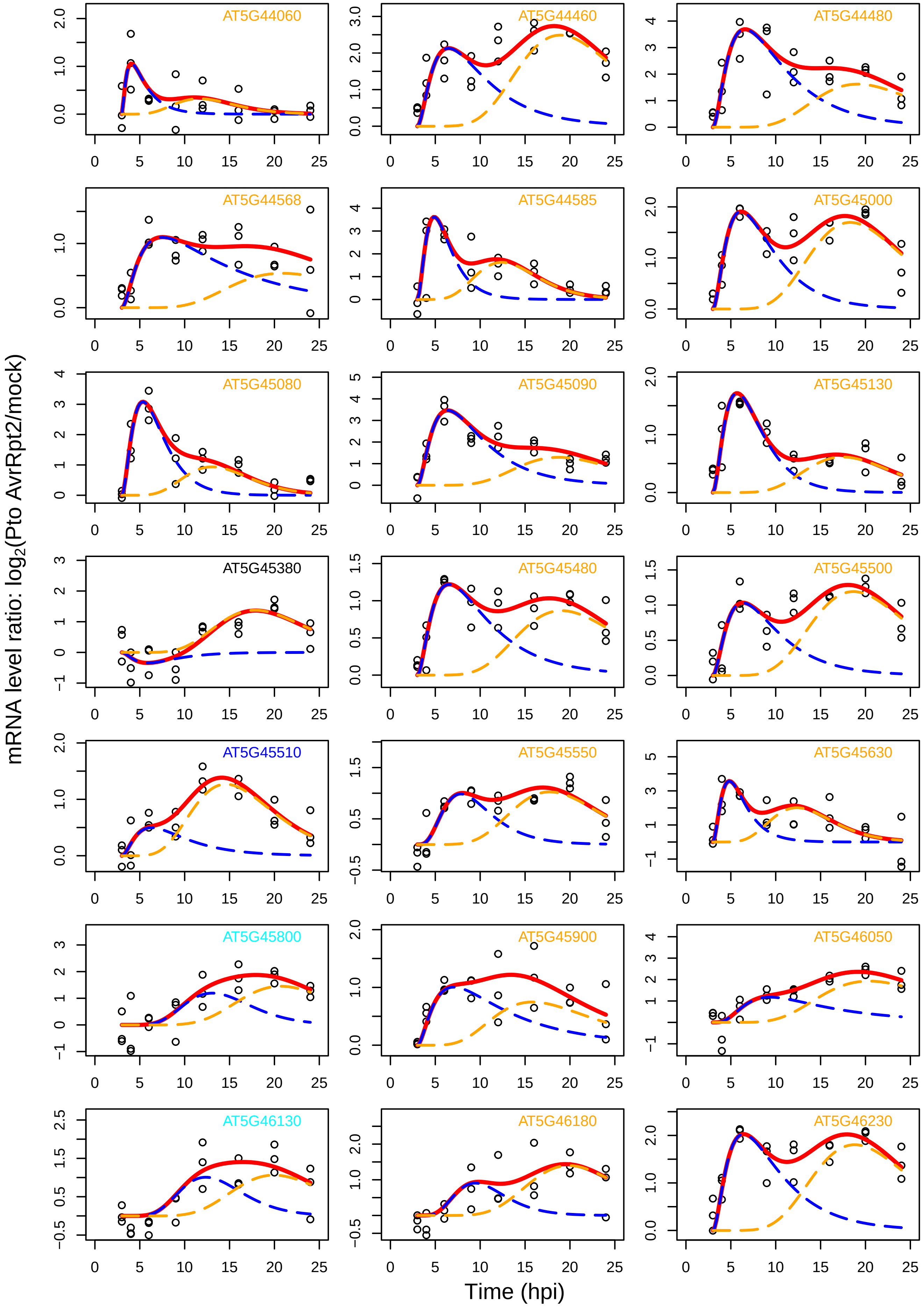

mRNA level ratio: log2(Pto AvrRpt2/mock)
Time (hpi)
AT5G44060
AT5G44460
AT5G44480
AT5G44568
AT5G44585
AT5G45000
AT5G45080
AT5G45090
AT5G45130
AT5G45380
AT5G45480
AT5G45500
AT5G45510
AT5G45550
AT5G45630
AT5G45800
AT5G45900
AT5G46050
AT5G46130
AT5G46180
AT5G46230

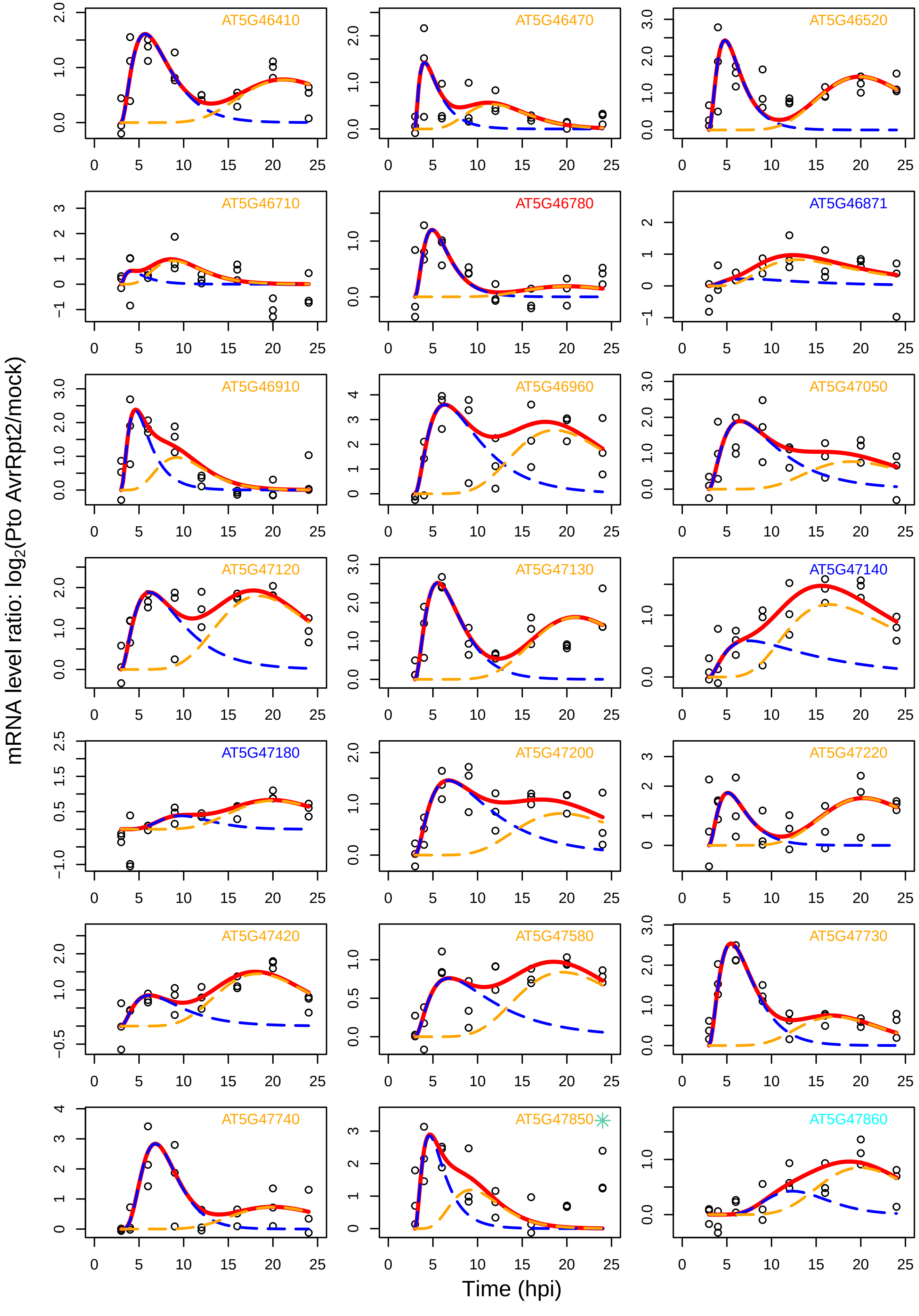
AT5G46410
AT5G46470
AT5G46520
AT5G46710
AT5G46780
AT5G46871
AT5G46910
AT5G46960
AT5G47050
AT5G47120
AT5G47130
AT5G47140
AT5G47180
AT5G47200
AT5G47220
AT5G47420
AT5G47580
AT5G47730
AT5G47740
AT5G47850
AT5G47860
mRNA level ratio: log2(Pto AvrRpt2/mock)
Time (hpi)

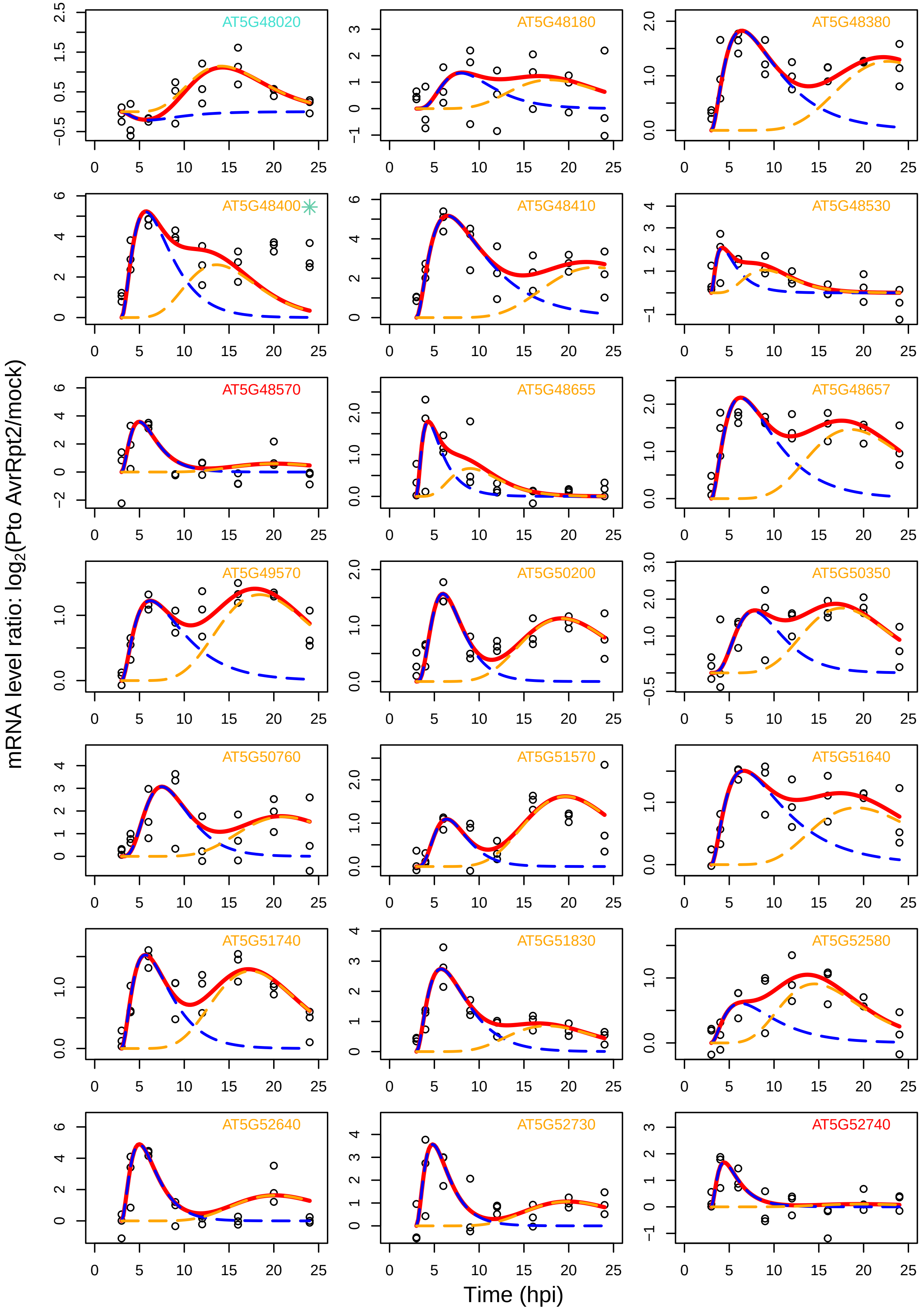
AT5G48020
AT5G48180
AT5G48380
AT5G48400
AT5G48410
AT5G48530
AT5G48570
AT5G48655
AT5G48657
AT5G49570
AT5G50200
AT5G50350
AT5G50760
AT5G51570
AT5G51640
AT5G51740
AT5G51830
AT5G52580
AT5G52640
AT5G52730
AT5G52740
mRNA level ratio: $\log_2$(Pto AvrRpt2/mock)
Time (hpi)

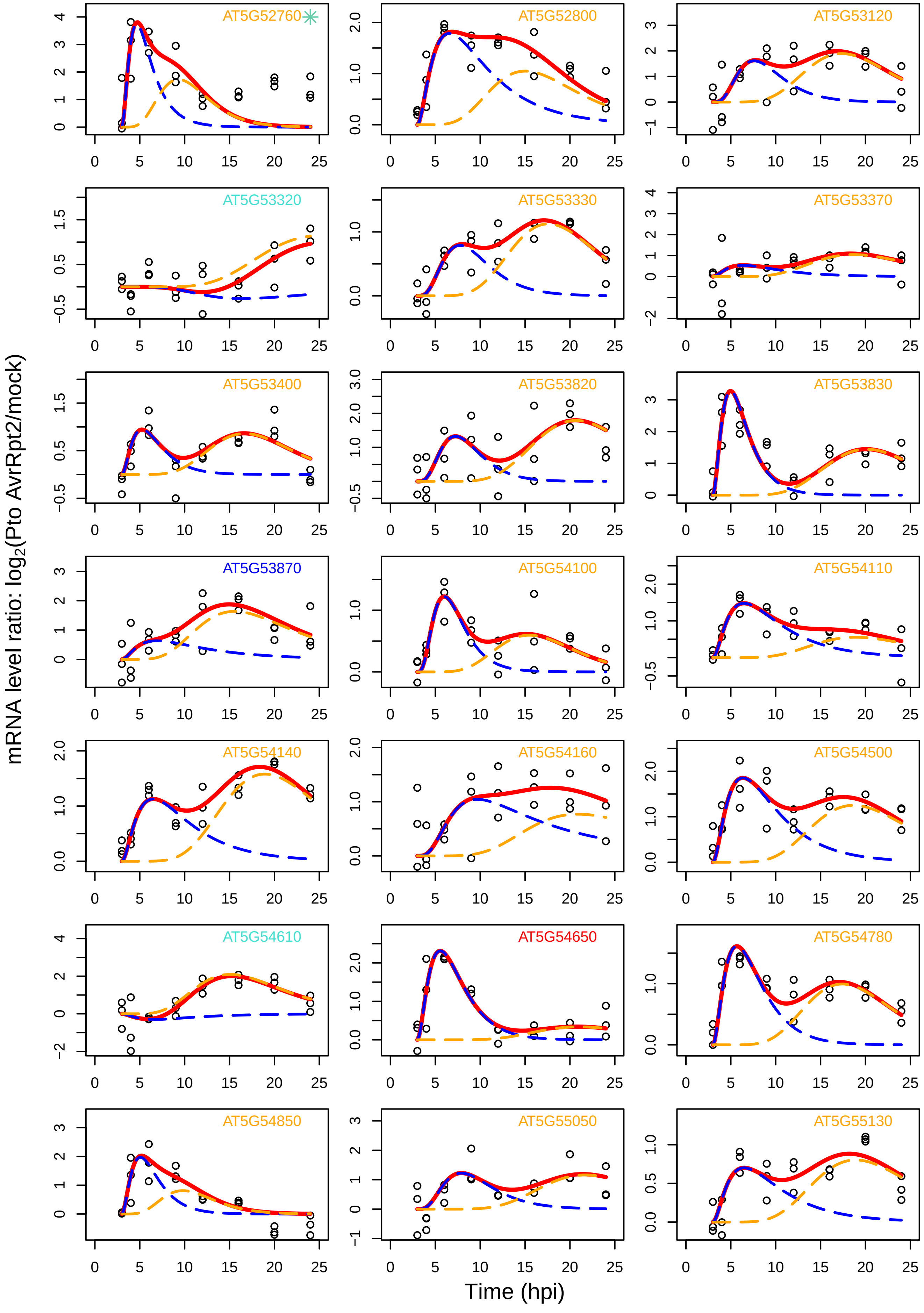

AT5G52760
AT5G52800
AT5G53120
AT5G53320
AT5G53330
AT5G53370
AT5G53400
AT5G53820
AT5G53830
AT5G53870
AT5G54100
AT5G54110
AT5G54140
AT5G54160
AT5G54500
AT5G54610
AT5G54650
AT5G54780
AT5G54850
AT5G55050
AT5G55130
mRNA level ratio: log2(Pto AvrRpt2/mock)
Time (hpi)

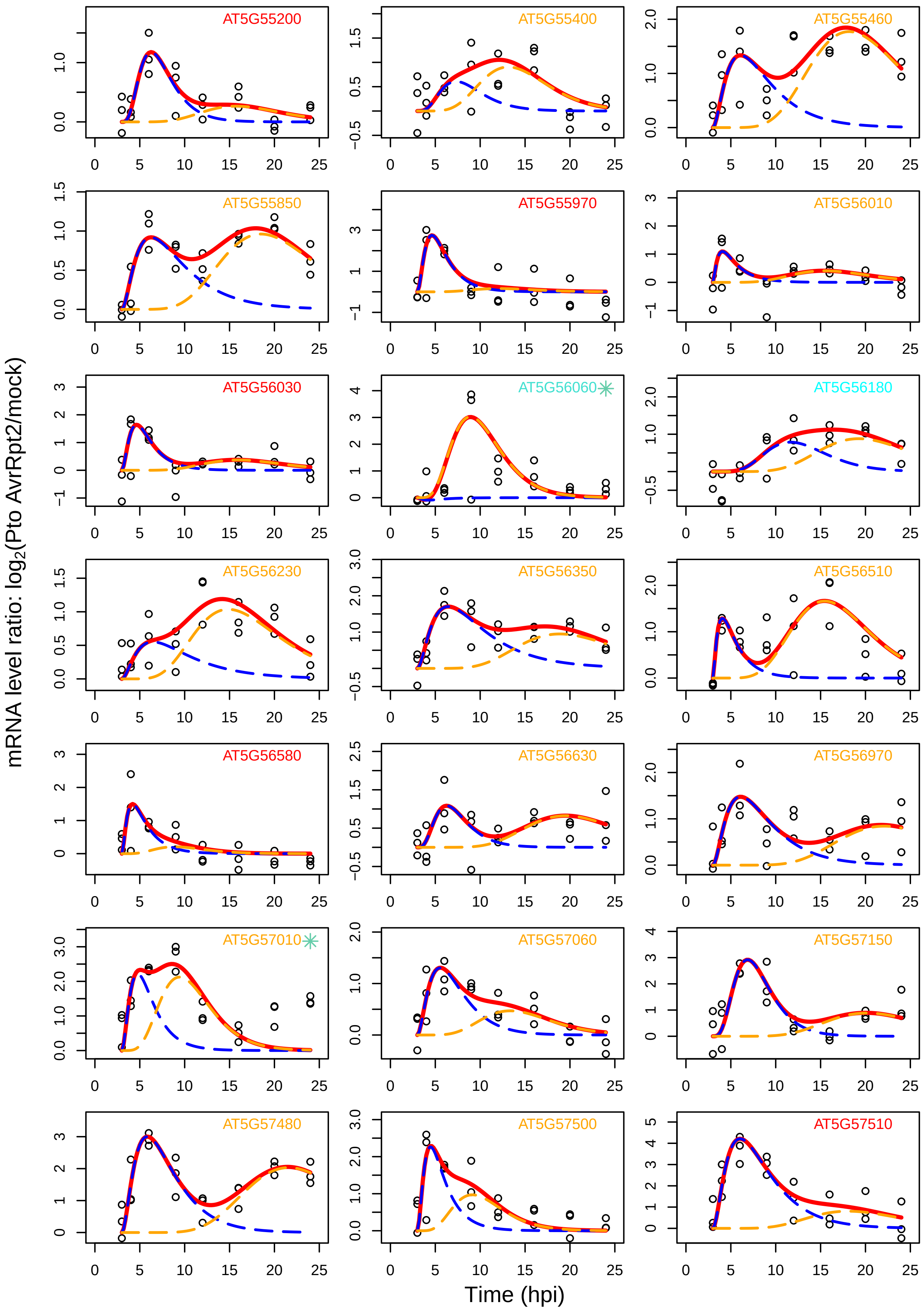

mRNA level ratio: $log_2$(Pto AvrRpt2/mock)
Time (hpi)
AT5G55200
AT5G55400
AT5G55460
AT5G55850
AT5G55970
AT5G56010
AT5G56030
AT5G56060
AT5G56180
AT5G56230
AT5G56350
AT5G56510
AT5G56580
AT5G56630
AT5G56970
AT5G57010
AT5G57060
AT5G57150
AT5G57480
AT5G57500
AT5G57510

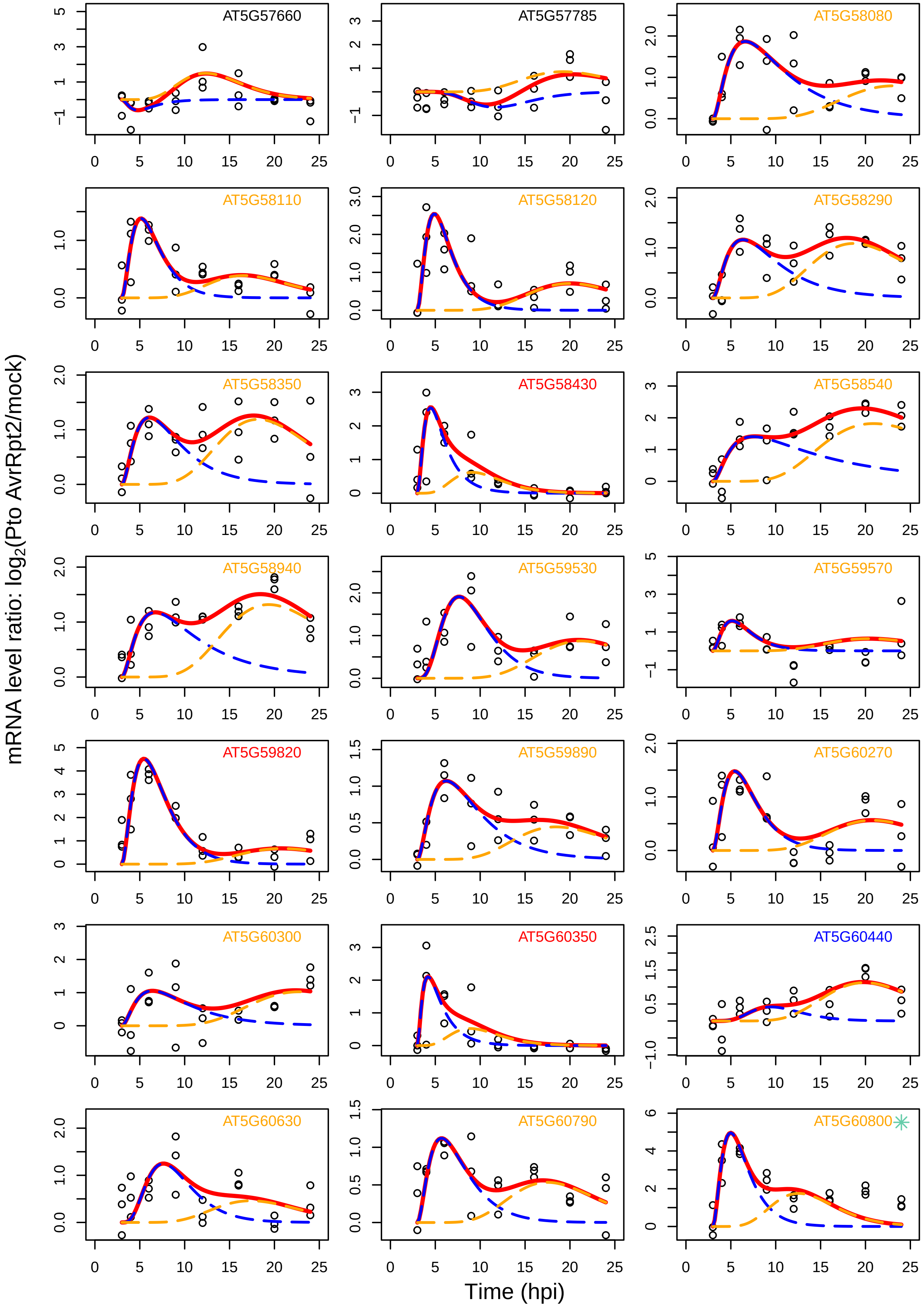

AT5G57660
AT5G57785
AT5G58080
AT5G58110
AT5G58120
AT5G58290
AT5G58350
AT5G58430
AT5G58540
AT5G58940
AT5G59530
AT5G59570
AT5G59820
AT5G59890
AT5G60270
AT5G60300
AT5G60350
AT5G60440
AT5G60630
AT5G60790
AT5G60800
mRNA level ratio: log2(Pto AvrRpt2/mock)
Time (hpi)

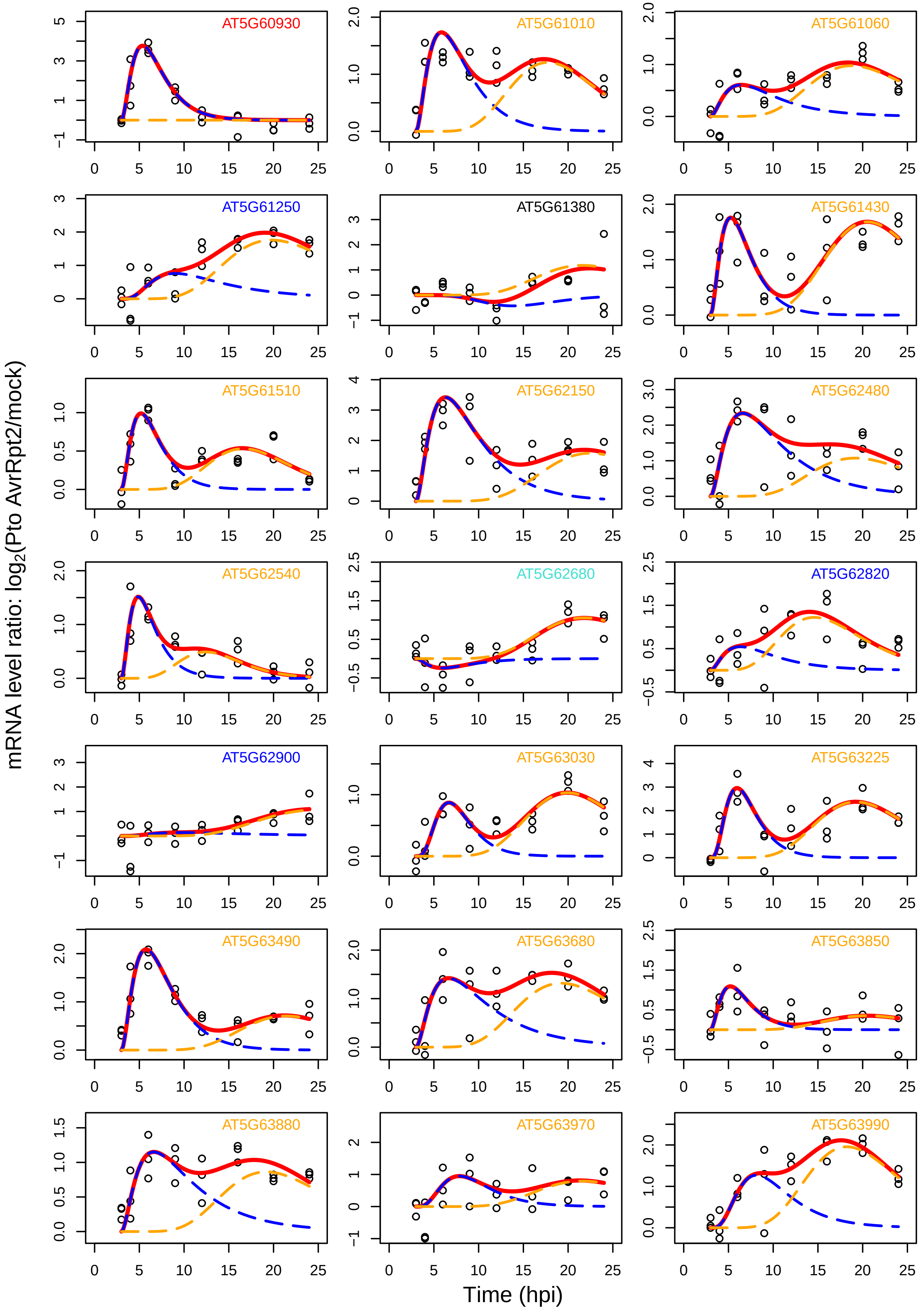
AT5G60930
AT5G61010
AT5G61060
AT5G61250
AT5G61380
AT5G61430
AT5G61510
AT5G62150
AT5G62480
AT5G62540
AT5G62680
AT5G62820
AT5G62900
AT5G63030
AT5G63225
AT5G63490
AT5G63680
AT5G63850
AT5G63880
AT5G63970
AT5G63990
mRNA level ratio: log2(Pto AvrRpt2/mock)
Time (hpi)

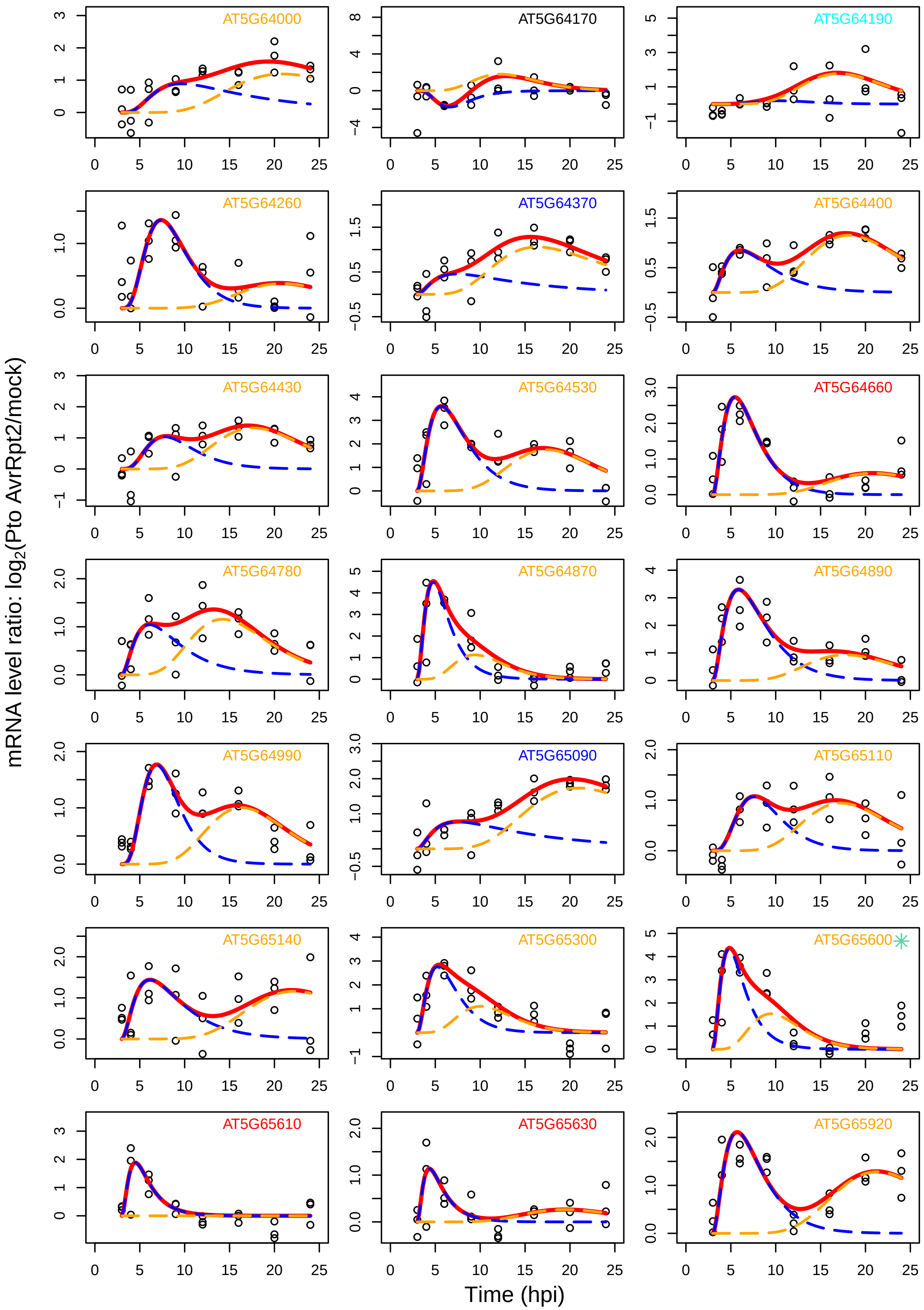

mRNA level ratio: log2(Pto AvrRpt2/mock)
Time (hpi)
AT5G64000
AT5G64170
AT5G64190
AT5G64260
AT5G64370
AT5G64400
AT5G64430
AT5G64530
AT5G64660
AT5G64780
AT5G64870
AT5G64890
AT5G64990
AT5G65090
AT5G65110
AT5G65140
AT5G65300
AT5G65600
AT5G65610
AT5G65630
AT5G65920

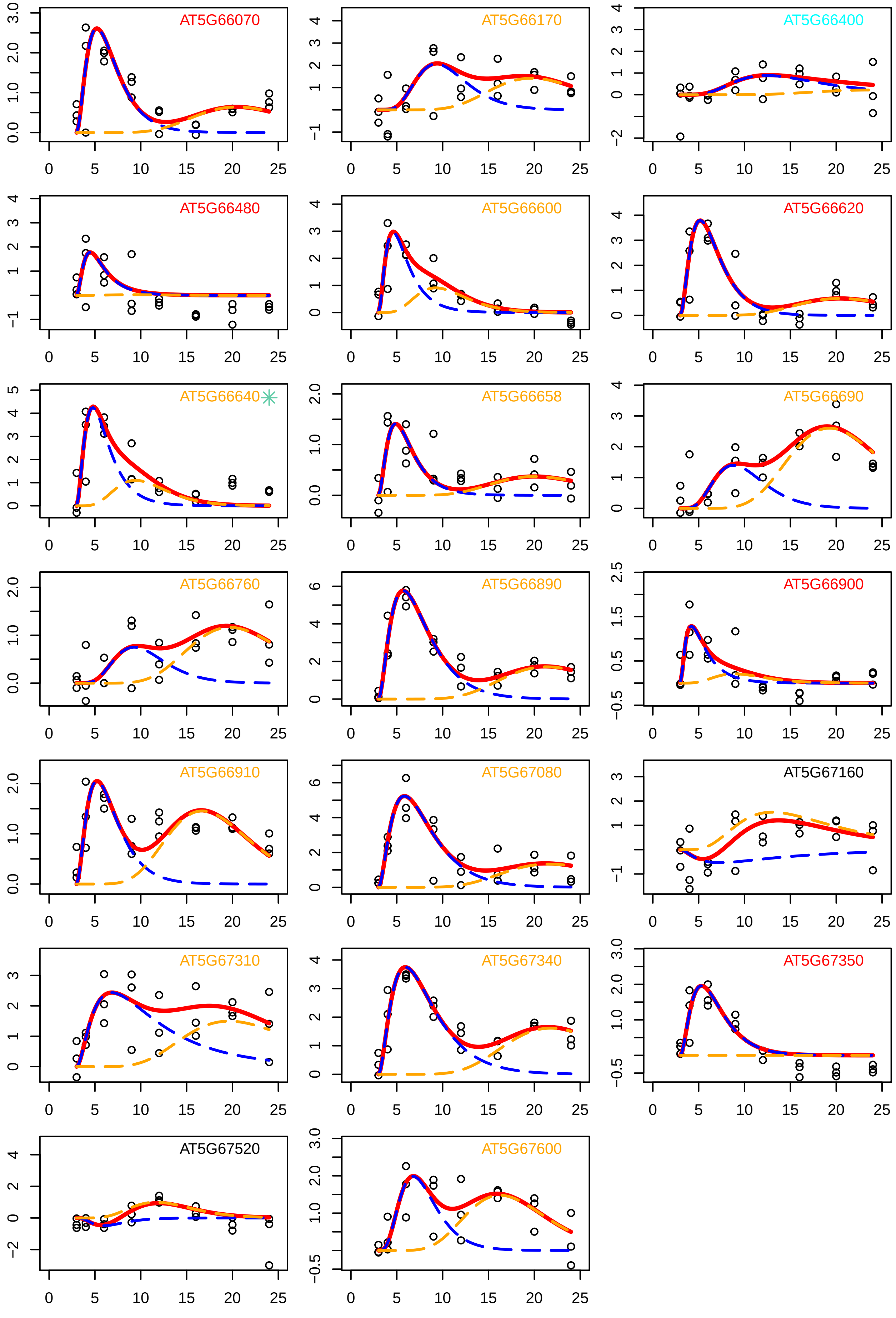
AT5G66070
AT5G66170
AT5G66400
AT5G66480
AT5G66600
AT5G66620
AT5G66640
AT5G66658
AT5G66690
AT5G66760
AT5G66890
AT5G66900
AT5G66910
AT5G67080
AT5G67160
AT5G67310
AT5G67340
AT5G67350
AT5G67520
AT5G67600
